# Extensive mitogenome divergence across the Rafflesiaceae in size and impact of horizontal gene transfer

**DOI:** 10.64898/2026.07.16.738911

**Authors:** LF Ceriotti, LM Gatica-Soria, W Tulle, Runxian Yu, Tian Bin, Zhu Renbin, Lu Zhiqiang, Yang Yongzhi, Zhou Renchao, MV Sanchez-Puerta

**Affiliations:** IBAM, Universidad Nacional de Cuyo, CONICET, Facultad de Ciencias Agrarias, Almirante Brown 500, Chacras de Coria, M5528AHB, Mendoza, Argentina; Facultad de Ciencias Exactas y Naturales, Universidad Nacional de Cuyo, Padre Jorge Contreras 1300, M5502JMA, Mendoza, Argentina; School of Life Sciences, State Key Laboratory of Biocontrol and Guangdong Provincial Key Laboratory of Plant Resources, Sun Yat-sen University, Guangzhou 510275, China; Yunnan Key Laboratory of Plateau Wetland Conservation, Restoration and Ecological Services, Southwest Forestry University, Kunming 650224, China; Shangri-La Potatso National Park Bita Lake Plateau Wetland Ecosystem Observation and Research Station of Yunnan Province, Kunming 650233, China; Laboratory of Tropical Forest Ecology, Xishuangbanna Tropical Botanical Garden, Chinese Academy of Sciences, Mengla 666303, China; Xishuangbanna Tropical Botanical Garden, Chinese Academy of Sciences, Mengla 666303, Yunnan, China; State Key Laboratory of Herbage Improvement and Grassland Agro-Ecosystems, College of Ecology, Lanzhou University, Lanzhou 730000, China

## Abstract

Horizontal gene transfer (HGT) drives organellar evolution, particularly in parasitic plants where host connections facilitate extensive DNA exchange. However, how these processes intersect with cellular machinery to reshape mitogenomic architecture remains poorly understood. Here, we investigate the mechanisms governing structural plasticity and asymmetric host-DNA integration in the extreme holoparasitic family Rafflesiaceae. By performing a comprehensive comparative analysis across all three extant genera (*Sapria*, *Rhizanthes*, and *Rafflesia*) and their *Tetrastigma* host lineage, we discovered extraordinary mitogenome size divergence, ranging from the expanded 824-kb genome of *Sapria* (40 circular chromosomes) to the streamlined 282-kb genome of *Rhizanthes* (35 circular chromosomes). Strikingly, these closely related genera display a total lack of chromosomal synteny, which we link to the ancestral loss of key recombination surveillance genes (RECX, ODB1). Furthermore, while all three genera strictly conserve an identical core of 30 protein-coding genes, host-derived HGT is highly asymmetric, ranging from minimal in *Rhizanthes* to 60% in *Sapria*. In *Sapria*, foreign tracts are sequestered into 15 predominantly non-coding circular chromosomes, a structural arrangement that aligns with the circle-mediated HGT model validated in other holoparasites. Collectively, these parallel patterns across phylogenetically distant lineages demonstrate that sorting and maintaining foreign DNA in autonomous circular blocks is a convergent architectural outcome of massive host-to-parasite genetic transfers.

**SIGNIFICANCE STATEMENT:** Horizontal gene transfer is widespread in the nuclear genome of the parasitic plant family Rafflesiaceae, but its contribution to mitochondrial genome evolution has been assessed through the analyses of a limited number of genes. By comparing complete mitochondrial genomes of the parasites and their hosts, we found that closely related species evolved dramatically different genome architectures through distinct mechanisms: one lineage accumulated large amounts of host-derived DNA, whereas another expanded through the proliferation of repetitive sequences with limited contribution from foreign DNA. These findings show that different evolutionary processes can generate profoundly divergent mitochondrial genomes even among closely related parasitic plants.

## INTRODUCTION

Horizontal Gene Transfer (HGT), the movement of DNA between different species, is a major evolutionary force, yet multiple aspects of this phenomenon are not fully understood. Among eukaryotes, numerous reports involve land plants, which have acquired genes mainly from bacteria, fungi, and viruses [Ma2022MolecularPlant]. More recent HGT events involving angiosperms result from plant-to-plant transfers, which have been increasingly identified in both their nuclear and mitochondrial genomes (Petersen *et al*., 2020; Wickell & Li, 2020; Pereira *et al*., 2023; Mariault *et al*., 2025).

A mitochondrial fusion compatibility model poses that complete mitochondria from the donor plant enter the recipient cell, fuse with the resident mitochondria, and their genomes recombine, completing the process of mitochondrial HGT (Rice *et al*., 2013). Multiple lines of evidence support this model, including the fact that large foreign mitochondrial DNA tracts are often identified and the donors are green plants, which have compatible mitochondrial fusion mechanisms, instead of bacteria or fungi that frequently interact closely with plants. Recently, a circle-mediated HGT model proposes that the foreign mitochondrial genome (mtDNA) does not necessarily recombine with the native one upon transfer (Roulet *et al*., 2024). In these cases, microhomology-mediated pathways can generate subgenomic circles from the donor mtDNA, and these foreign circular molecules can replicate autonomously in the recipient mitochondria. So far, this model has been verified in two independent lineages of holoparasitic plants, Balanophoraceae and Mitrastemonaceae (Roulet *et al*., 2024; Roulet *et al*., 2026). It is yet unclear how widespread this model is for angiosperm mtDNA and which conditions are relevant to set the stage for the circularization process.

Parasitic plants, which invade stems or roots of other plants, are excellent systems to deepen our understanding of mitochondrial HGT because they are particularly prone to receiving DNA from their host plants through cell-to-cell connections. The holoparasitic angiosperm family Rafflesiaceae (order Malpighiales) represents one of the most extreme forms of parasitism among flowering plants (Thorogood *et al*., 2021). These endoparasitic plants lack a typical vegetative body and persist within their hosts as highly reduced, mycelium-like tissues, emerging only to produce flowers. Rafflesiaceae are known for exhibiting the largest flowers and for lacking photosynthesis and likely, also their plastid genomes (Molina *et al*., 2014; Cai *et al*., 2021; Yu *et al*., 2026). It encompasses three genera of endoparasitic species that live within the roots of plants from the genus *Tetrastigma* (Vitaceae) (Thorogood *et al*., 2021). Earlier studies discovered a large number of foreign nuclear and mitochondrial genes as a result of recent and ancestral HGT events from their hosts (Xi *et al*., 2013; Cai *et al*., 2021; Guo *et al*., 2023; Molina *et al*., 2025). However, the mtDNA of angiosperms consists of >80% non-coding DNA that awaits examination to assess the extent of mitochondrial HGT and the evidence of the circle-mediated HGT model.

To date, the phylogenetic studies of Rafflesiaceae mtDNA have been restricted to the protein-coding regions of a few species of Raffleisaceae (Nickrent *et al*., 2004; Barkman *et al*., 2007; Xi *et al*., 2013; Guo *et al*., 2023). In this study, we expand this framework by analyzing the mitochondrial genomes of *Rhizanthes lowii*, two individuals of *Sapria himalayana*, and three species of *Rafflesia*, alongside host mitochondrial data comprising two complete *Tetrastigma* mtDNAs and 15 additional draft assemblies from diverse *Tetrastigma* species and other Vitaceae. We assessed the extent of foreign HGT in the non-coding mtDNA of Rafflesiaceae, and evaluated the phylogenetic origin of their mitochondrial genes inferring both recent and ancestral events. Finally, we investigated the integrity of the organellar DNA DNA replication, recombination, and repair (DNA-RRR) machinery as a putative factor driving mitochondrial genome evolution in this family.

## RESULTS

### Extensive mitochondrial genome divergence across the Rafflesiaceae

The mitochondrial genomes of representatives from the three genera within the family Rafflesiaceae exhibit striking divergence in both sequence composition and overall size (Table 1). *Sapria himalayana* possesses the largest mitogenome in the family (Figure 1), as observed in the two independently assembled individuals, which display nearly identical genome size, architecture, and sequence content (Table S1A, Figure S1, Supplementary Dataset 1). Both mtDNAs span ∼824 kb, are partitioned into 40 circular chromosomes ranging from 7,086 to 39,301 bp, and have an average GC content of 45.3% (Table 1 and S1). None of the assemblies recovered the mitochondrial chromosomes 1 and 14 (NCBI accessions OQ719965 and OQ719978, respectively) assembled by Guo et al. (Guo *et al*., 2023), even when using the same dataset. A careful examination suggests that those chromosomes are DNA sequences integrated in the nuclear genome of *Sapria himalayana* (Note S1; Figure S2).

**Figure 1.**
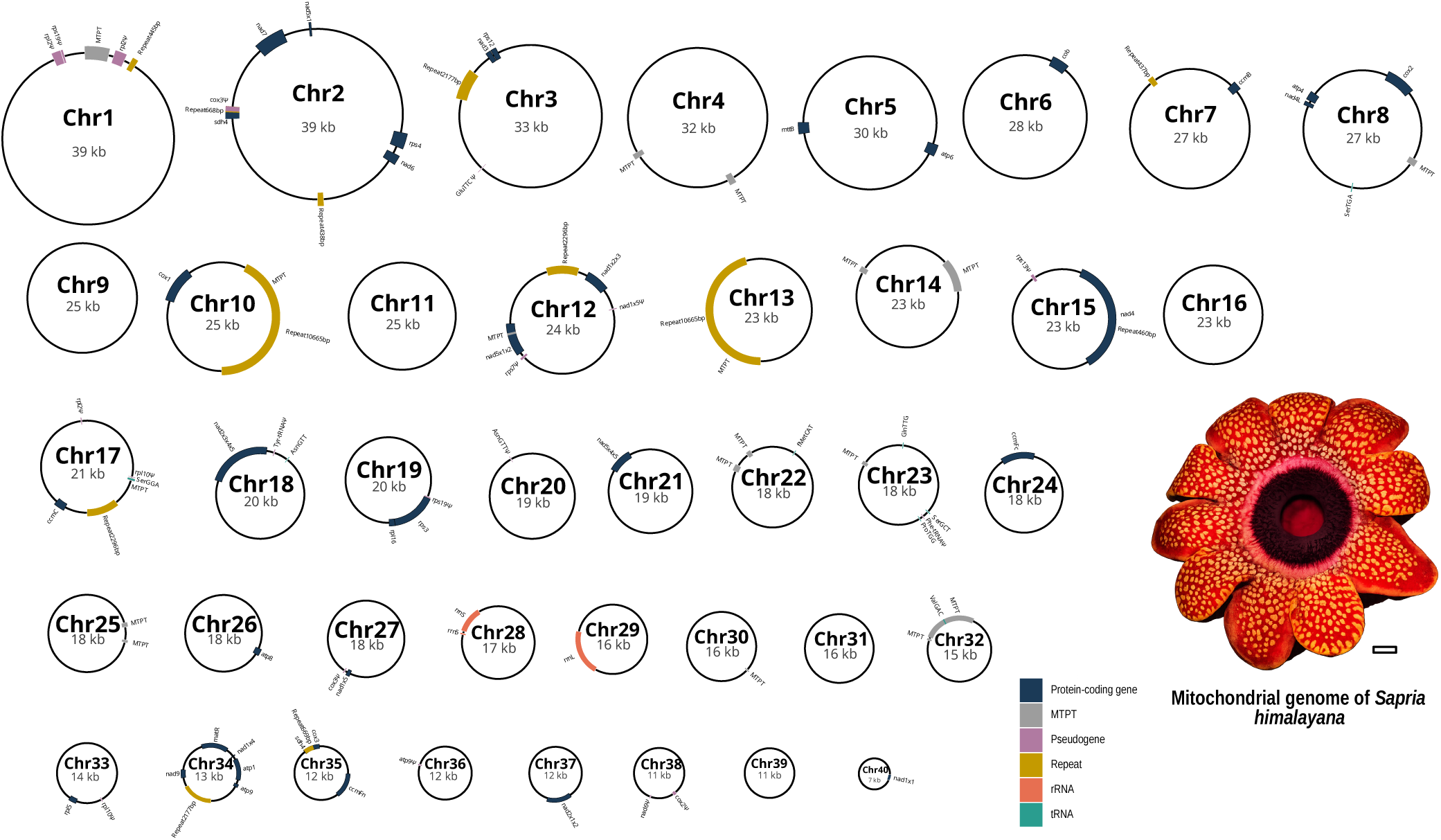
Mitogenomic architecture of the holoparasitic plant *Sapria himalayana* (Rafflesiaceae). The mitochondrial genome (mtDNA) is 824,044 bp in length and is partitioned into 40 circular chromosomes (Chr) of varying sizes. Concentric tracks depict distinct genomic and structural features, including intact protein-coding, ribosomal (rRNA), and transfer (tRNA) genes; pseudogenes >100 bp (denoted by Ψ); repetitive elements >400 bp; and plastid-derived integrants (MTPTs) >100 bp. The inset photograph displays a S. himalayana flower (photograph by Rohit Naniwadekar, sourced from iNaturalist, licensed under CC BY-SA). Scale bar, 1 cm.

**Table 1.** General features of the mitochondrial genomes of Rafflesiaceae species.

|  | *Sapria himalayana*<br>(individual 1) | *Sapria himalayana*<br>(individual 2) | *Rhizanthus lowii* | *Rafflesia arnoldii* | *Rafflesia leonardi*<sup>d</sup> | *Rafflesia tuan-mudae* |
| --- | --- | --- | --- | --- | --- | --- |
| Genome length (bp) | 824.044 | 824.038 | 282.577 | 406.461 | 393.915 | 416.026 |
| Status of mtDNA | Complete | Complete | Complete | Draft | Draft | Draft |
| Number of circular chromosomes (or * linear contigs) | 40 | 40 | 35 | 59 * | 42 * | 63 * |
| Average GC content (%) | 45.3 | 45.3 | 46.7 | 46.2 | 45.9 | 46.1 |
| Protein-coding genes (including repeats) | 30 (31) | 30 (31) | 30 (30) <sup>a</sup> | 30 (30) | 30 (33) | 30 (30) |
| rRNA-coding genes | 3 | 3 | 3 | 3 | 3 | 3 |
| tRNA-coding genes (including repeats) | 8 | 8 | 1 | 5 (6) | 5 (7) | 5 (6) |
| Group II introns | 19 | 19 | 19 | 19 | 19 | 19 |
| cis-splicing | 13 | 13 | 13 <sup>b</sup> | 13 | 13 | 13 |
| trans-splicing | 6 | 6 | 6 <sup>c</sup> | 6 | 6 | 6 |
| Group I introns | 1 | 1 | 1 | 1 | 1 | 1 |
| Repeats (bp, % of the genome) | 72,346 (8.7) | 72,707 (8.8) | 101,450 (35.9) | 31,573 (7.76) | 29,683 (7.53) | 32,815 (7.88) |
| Repeats >1 kb (unique) | 6 (3) | 6 (3) | 27 (16) | 0 | 0 | 0 |
| Chloroplast derived sequences (bp, % of the genome) | 12,428 (1.51) | 12,833 (1.55) | 555 (1.96) | 7,916 (1.95) | 6,320 (1.6) | 7,248 (1.74) |
| Foreign DNA content (bp, % of the genome) | 494,426 (60) | 494,422 (60) | 2,260 (0.8) | 14,632 (3.6) | 29,543 (7.5) | 38,274 (9.2) |

<sup>a</sup> *ccmFc* exon1 of *Rhizanthus* has a 339 bp insertion that does not shift the reading frame and has no sequence similarity in NCBI (BLASTn e-value <0.01).
<sup>b</sup> The *rps3* exons and *nad4* exons of *Rhizanthus* are in different chromosomes but the intron was considered cis-splicing, assuming that the chromosomes could be rearranged based on the multiple repeats.
<sup>c</sup> *nad5* intron 2 of *Rhizanthus* was considered a trans-splicing intron, although the distance between its exons could also be consistent with cis-splicing.
<sup>d</sup> The mitochondrial assembly of *R. leonardi* could not be corroborated because the DNaseq data is not available.

*Rhizanthes* harbors a ∼3 times smaller mtDNA (282 kb in length), exhibits a GC content of 46.7%, and is organized into 35 circular chromosomes, ranging from 2,818 to 17,967 bp (Figure S3-A). *Rhizanthes* shares the same protein-coding gene and intron content with *Sapria* comprising 30 genes, one group I intron, and 19 group II introns (13 cis-spliced and six trans-spliced) (Table 1). In contrast, the tRNA content differs markedly between the two taxa. While *Rhizanthes* encodes a single tRNA (*trnfM*), *Sapria* presents eight tRNA genes, all of which are of foreign origin (see below). Coding regions are distributed across 28 and 27 chromosomes in *Sapria* and *Rhizanthes* (Figure 1; Figure S3A), respectively, and chloroplast-derived sequences represent 1.5-2% of the genomes (Table 1).

For comparative purposes, we also assembled the draft mitochondrial genomes of two *Rafflesia* species (*R. arnoldii* and *R. tuan-mudae*; Figure S3B-C) and included the draft mtDNA of *R. leonardi* (Xi *et al*., 2013). These mtDNAs are approximately half the size of the *Sapria* mtDNA, comprising 42–63 linear contigs with a GC content of ∼46% and a total length of approximately 400 kb (Table 1; Supplementary Dataset 2). They carry the same 30 protein-coding genes as *Sapria* and *Rhizanthes* (Table 1). These three mitochondrial assemblies of *Rafflesia* species are highly similar in sequence, sharing more than 85% of their total length, with *R. arnoldii* and *R. tuan-mudae* sharing 97–98% (Table S1B).

The proportion of shared mtDNA among genera within the Rafflesiaceae is quite low considering intrafamily comparisons across angiosperms (Roulet *et al*., 2024). In fact, not a single mitochondrial chromosome is shared by these closely related genera (Figure 2) and the regions of similarity, that is homologous regions, are highly fragmented. *Rhizanthes* sequences that share similarity with *Rafflesia* and *Sapria* mtDNA are mostly restricted to genes and their surrounding regions (see below). Noticeably, the degree of mitochondrial sequence content shared across Rafflesiaceae (Figure 2) is not congruent with their established phylogenetic relationships (Pelser et al., 2019). *Rafflesia* and *Rhizanthes* share a higher proportion of homologous regions with *Sapria* (270-283 and 96 kb, respectively) than between them (76-83 kb).

**Figure 2.**
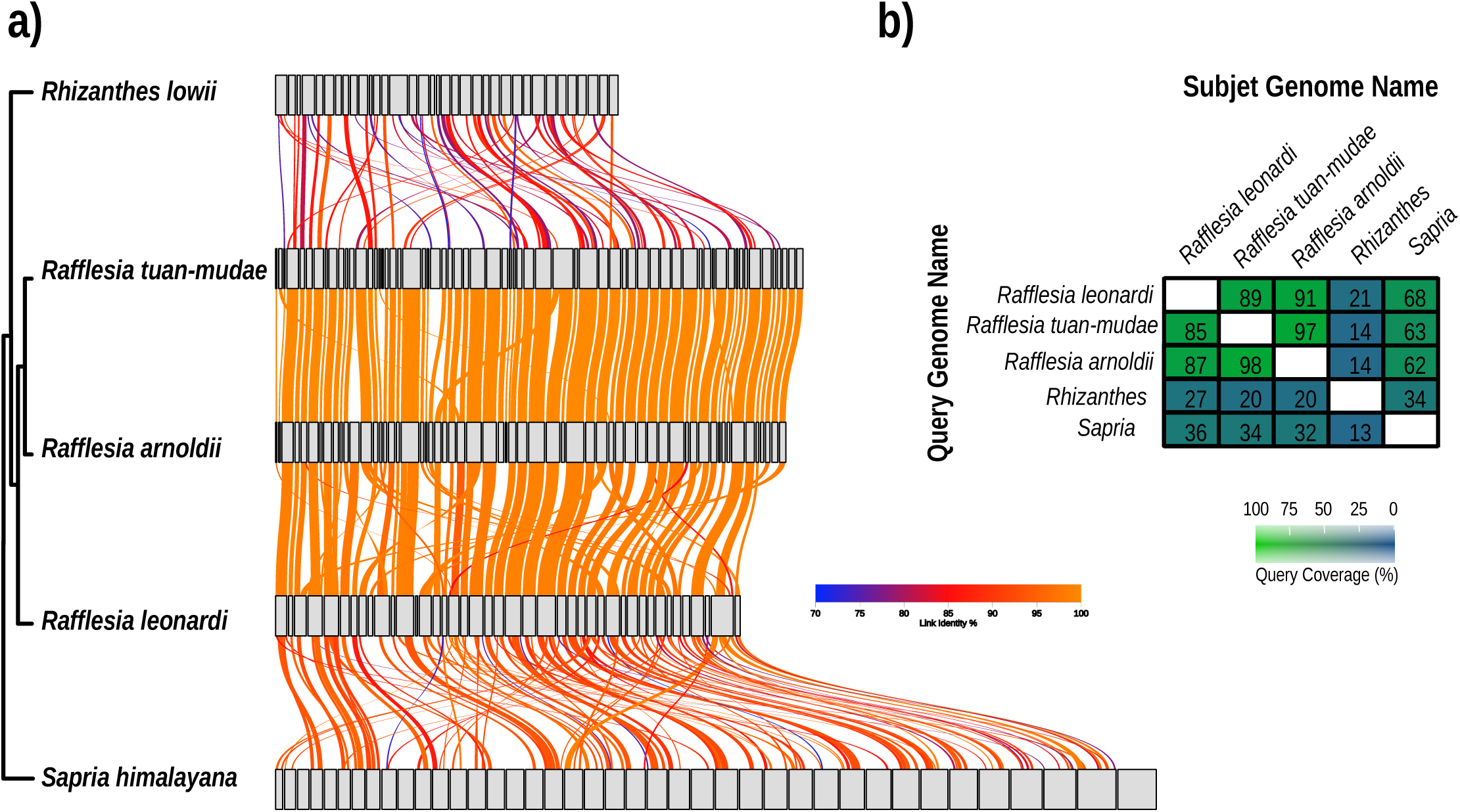
Interspecific mitochondrial genome variability in Rafflesiaceae. (a) Collinearity among mitochondrial sequences of endoparasitic Rafflesiaceae. Gray blocks represent mitochondrial chromosomes or contigs, with block width proportional to sequence length. Colored connecting lines indicate homologous regions (>1 kb, >70% identity), with color intensity reflecting percentage identity (70–100%). Mitochondrial alignments were generated using LASTZ v1.0.4 and visualized with AliTV v1.0.6. The phylogenetic topology of Rafflesiaceae was taken from Pelser *et al*. (2019). (b) Percentage of query coverage among mitochondrial genomes of Rafflesiaceae. For each species, mitochondrial chromosomes were concatenated prior to BLASTn analysis (e-value < 2 × 10^−10^).

Finally, the Rafflesiaceae exhibits contrasting repeat patterns (Figure 3A). The mtDNAs of *Sapria* and *Rafflesia* spp. contain very few repeats, encompassing less than 9% of their genomes (Figure 3). In contrast, the mtDNA of *Rhizanthes* is highly repetitive, with repeated sequences distributed across all mitochondrial chromosomes and accounting for 101 kb of the genome (Table S2A). Given that more than 25% of the mtDNA is composed of short repeats (73.5 kb), we further characterized these sequences in this species to determine whether they represent numerous independent repeat families or expansions of a limited set of motifs (Table S2B). Short repeats were clustered based on sequence similarity, defining each cluster by a centroid sequence to which all members shared ≥80% identity. The majority of short repeats (>90%) were assigned to clusters, indicating substantial sequence similarity among repeats. Notably, clusters containing ≥200 repeat copies accounted for more than 80% of all short repeat occurrences, and the five largest clusters alone comprised over half of the short-repeat content of the genome. In addition, short repeat sequences covering ∼30 kb are arranged in tandem (Table S2B). Together, these patterns indicate that the repetitive landscape of the *Rhizanthes* mtDNA is largely dominated by a small number of highly expanded repeat families.

**Figure 3.**
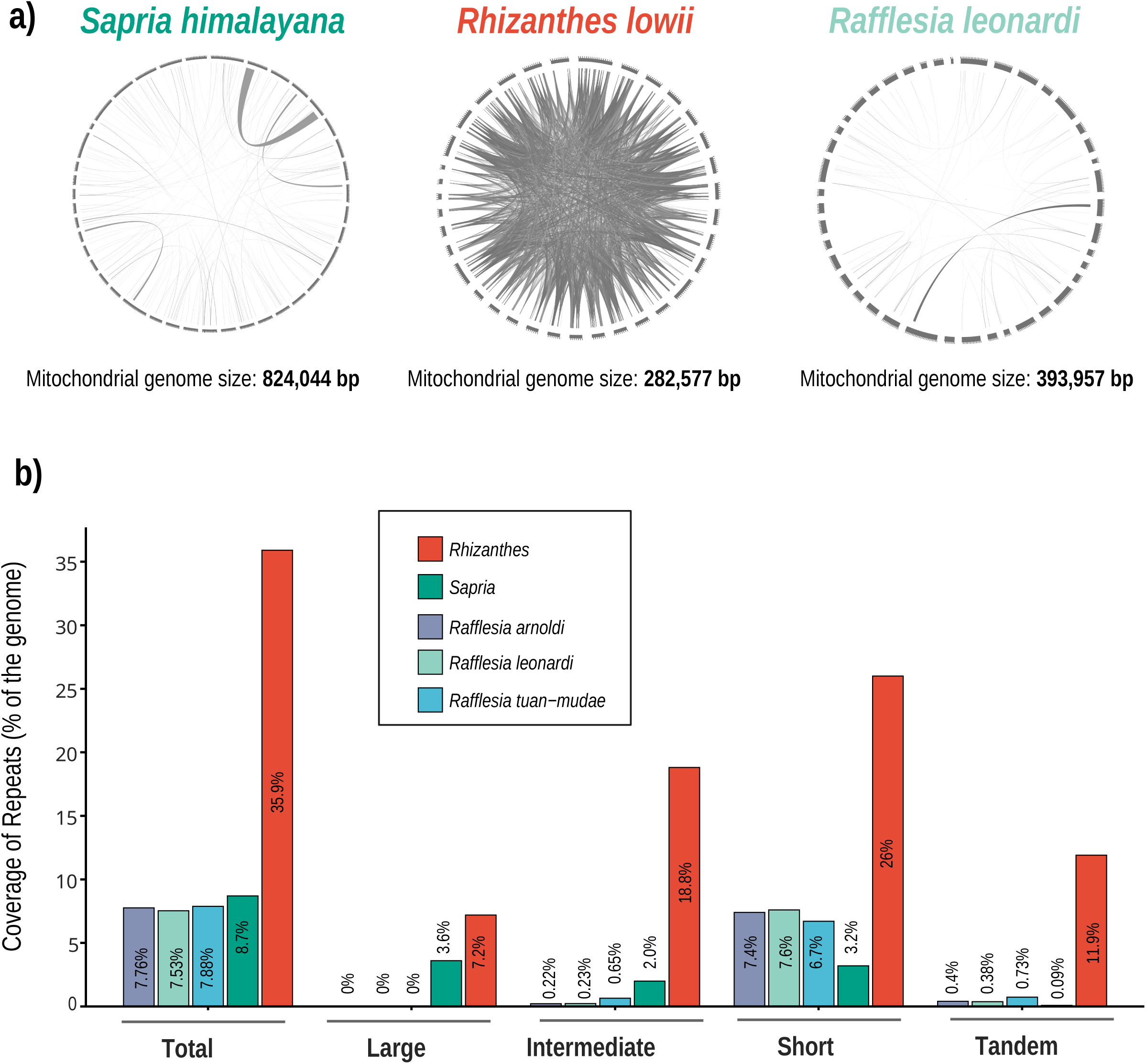
Repetitive elements in mitochondrial genomes of the Rafflesiaceae. (a) Visualization of repeats in the mitochondrial genomes of *Rhizanthes lowii*, *Sapria himalayana*, and *Rafflesia leonardi*. Only repeats ≥400 bp in length and with ≥90% nucleotide identity are shown. Intragenomic alignments were generated using LASTZ v1.0.4 and visualized with AliTV v1.0.6. (b) Proportion of repetitive sequences relative to total mitochondrial genome size across five sequenced species of Rafflesiaceae. Repeats are classified by size and structural category (Total, Large, Intermediate, Short, and Tandem repeats). Bar heights and numeric labels indicate the percentage of the mitogenome occupied by each repeat category.

### Asymmetric HGT dynamics in Rafflesiaceae mtDNAs

To unveil the impact of horizontal gene transfer from their hosts or other donors on the mtDNAs of Rafflesiaceae, we performed comparative genomics and phylogenetic analyses of intergenic and genic regions, respectively. Overall, the extent of foreign sequences vary markedly across the three Rafflesiaceae lineages, revealing highly contrasting patterns (Figure 4, Figure S4, Tables S7-S11).

**Figure 4.**
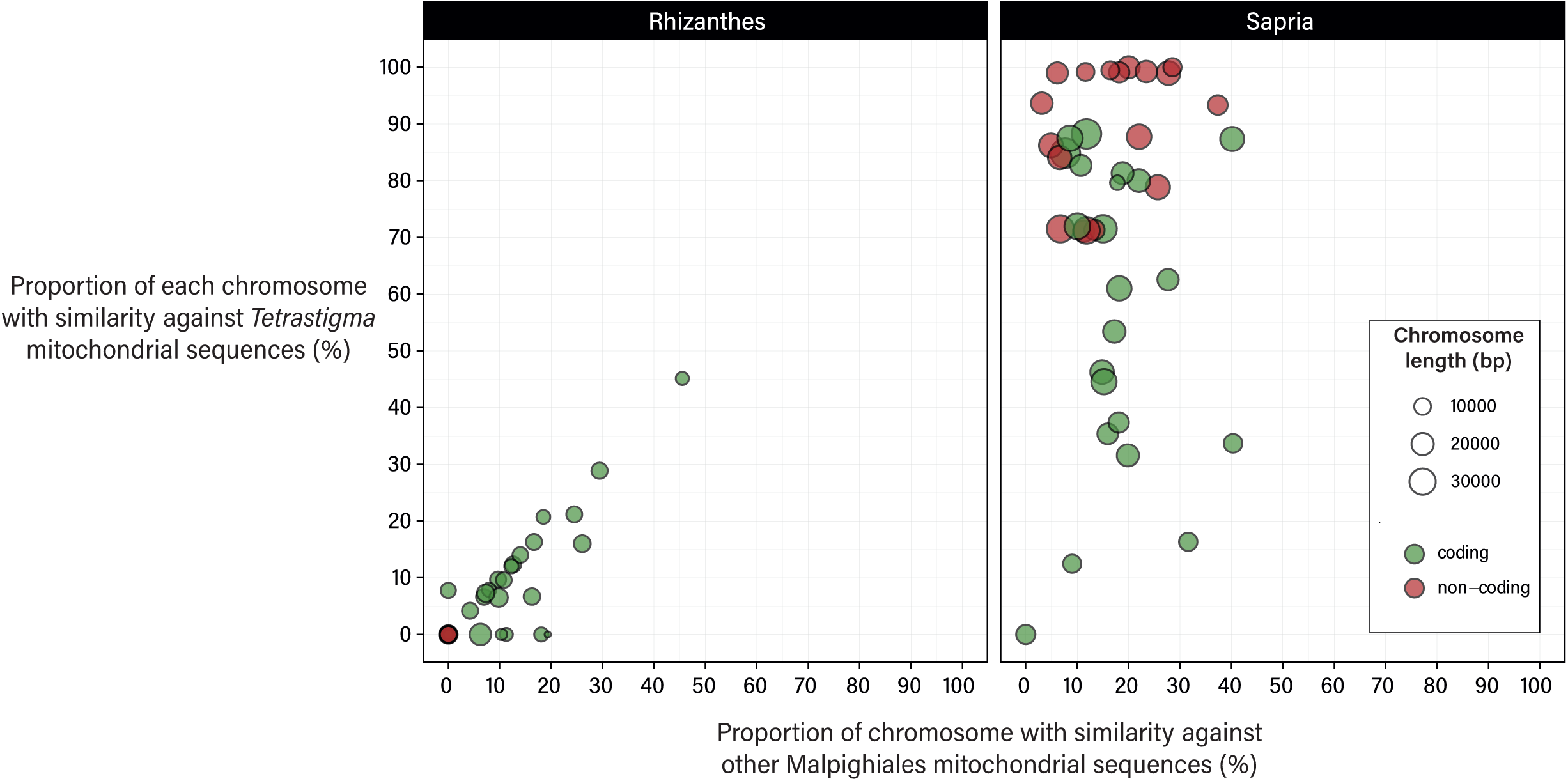
Sequence comparisons of *Rhizantes lowii* and *Sapria himalayana* mitochondrial chromosomes against mitochondrial sequences from *Tetrastigma* and other Malpighiales (excluding Rafflesiaceae). BLASTn hits >200 bp in length and >90% identity were included.

*Sapria* exhibits the highest proportion of foreign mitochondrial DNA in the family, with ∼60% of its mtDNA derived from *Tetrastigma* (Figure S4). The high-scoring BLASTn hits (identity percent >90% and >200 bp in length) of each *Sapria* mitochondrial chromosome show substantially greater coverage and identity to *Tetrastigma* than to any other taxonomic category (Table S3), including their close relatives in the order Malpighiales (Figure 4). A detailed analysis of each chromosome of *Sapria* demonstrates that the amount of sequences with affinity to *Tetrastigma* is unevenly distributed (Figure S4). Ten noncoding mitochondrial chromosomes (14, 20, 22, 23, 25, 30, 32, 36, 38, and 39) share >90% of their length with *Tetrastigma* mtDNAs and are inferred to be fully foreign; eight of these exhibit an outstanding >98% coverage with *Tetrastigma* mitochondrial DNA (Figure 4). An additional of five noncoding chromosomes (4, 9, 11, 13, and 31) share 71–88% of their sequence with *Tetrastigma* mtDNA (Figure 4).

Although most chromosomes also display BLASTn hits to taxa in the Malpighiales, these hits are shorter, more scattered, and exhibit lower sequence identity than those to *Tetrastigma* or other Vitales (Supplementary Dataset 3). In fact, when the BLASTn hit identity threshold is increased from 90% to 95%, no *Sapria* chromosome shares >15% of its length with other Malpighiales taxa while shared sequences with *Tetrastigma* remain mostly unaffected (Table S3).

Across the 15 foreign chromosomes in the mtDNA of *Sapria*, sequence similarity to *Rafflesia* or *Rhizanthes* mtDNA is minimal (<10% and <3%, respectively). Exceptionally, chromosome 22 of *Sapria* shares ∼37% of its length with both *Rafflesia leonardi* (KJ154982) and over 90% with *Tetrastigma* (Table S3). A phylogenetic tree of this region shows that *Sapria* and *Rafflesia* are not sister taxa, indicating that these regions with similarity to the host plant are the result of independent HGT events from *Tetrastigma* (Figure S5).

In contrast, *Rhizanthes* mtDNA shares very few of the mitochondrial intergenic regions with other Malpighiales mtDNAs (Figure 4, Supplementary Dataset 2) and only minimal evidence of intergenic HGT was detected (Figure S4). Only one chromosome (Chr30) shares >45% of its sequence with non-Rafflesiaceae Malpighiales, and ten chromosomes do not share sequences at all (considering BLASTn identity >90% and >200 bp in length) with the mitochondrial genome database (Figure 4, Table S4). The few BLASTn matches recovered against Vitales correspond either to genes or to intergenic regions broadly shared across angiosperms (Supplementary Dataset 3, Table S4). In *Rafflesia spp.*, HGT events were detected in low levels: approximately 3-10% of the draft mtDNA correspond to Vitaceae-derived sequences, distributed among 3-8 contigs (Figure S4, Table S5-S7). Notably, most foreign content in *R. leonardi* is concentrated in a single, fully foreign 22,595 bp contig (KJ154982), approximately half of which is present in the other *Rafflesia spp.* (Table S5), and which was the result of an independent HGT event in *Sapria* (see above).

### Origin of mitochondrial genes in Rafflesiaceae

Phylogenetic analyses of individual genes revealed that, among the 30 protein-coding genes encoded by Rafflesiaceae mitochondrial genomes, only *atp1* shows a strongly-supported foreign origin in all three genera examined (Table 2, Fig S6). This foreign gene in Rafflesiaceae forms a monophyletic clade indicating that it was acquired by the common ancestor and maintained along the speciation events. Noticeably, a native copy of *atp1* is no longer present in any of the parasites. In contrast, other foreign genes co-exist with native homologs. Second copies of *atp8* and *cox3* in *R. leonardi* are placed phylogenetically as sister to, or nested within, the angiosperm family Vitaceae, with bootstrap support values ≥70%. In addition, a second copy of *sdh4* exhibits a foreign origin in *Sapria* and *R. leonardi*, as inferred from comparative analyses that show this copy embedded in a large host-derived region shared with *Tetrastigma* (Fig S6). Genes that affiliate with Malpighiales, as well as those lacking robust phylogenetic support were classified as native. Those genes showing affiliations with Vitaceae but supported by low bootstrap values, namely *ccmB*, *ccmC*, *cox2*, and *nad7* in all three species, and *nad5* (exons 4 and 5) in *Rhizanthes* and *Rafflesia* (Figure S6) might be chimeric (part native, part foreign). Expanded taxon sampling across both host and parasite lineages will be necessary to resolve the evolutionary origins of these genes.

**Table 2.**
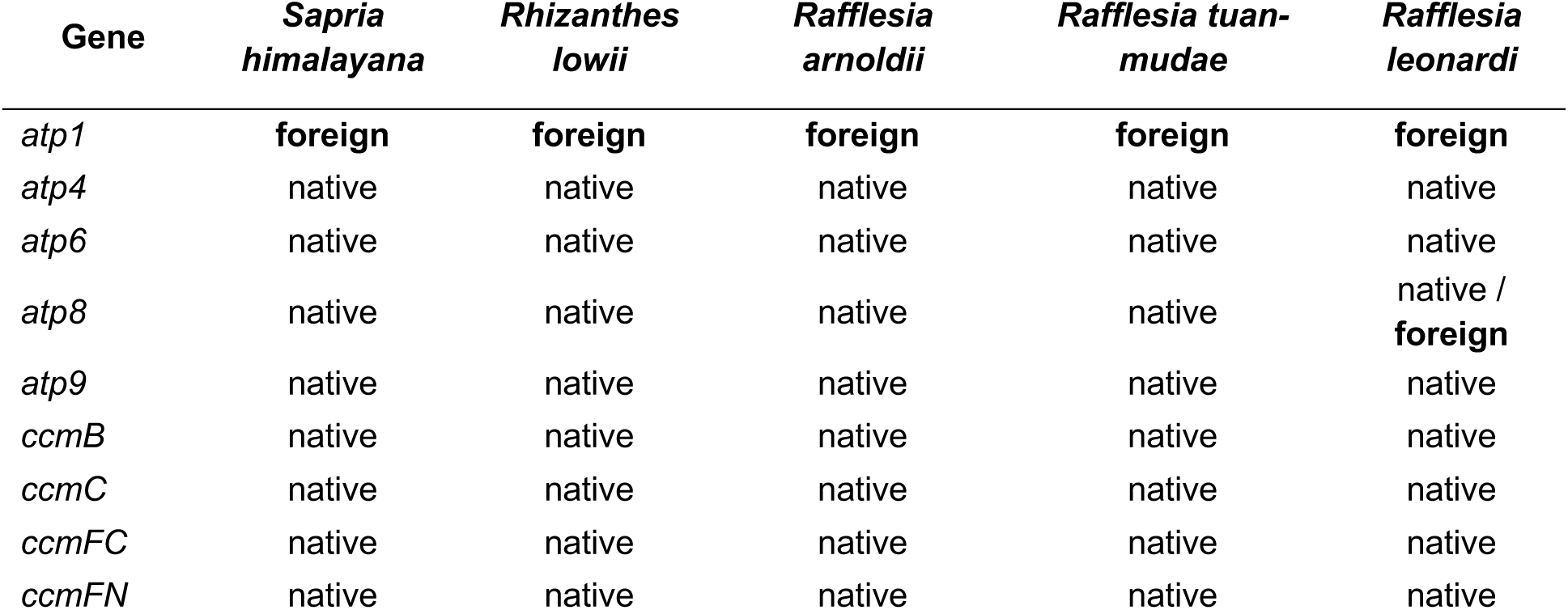
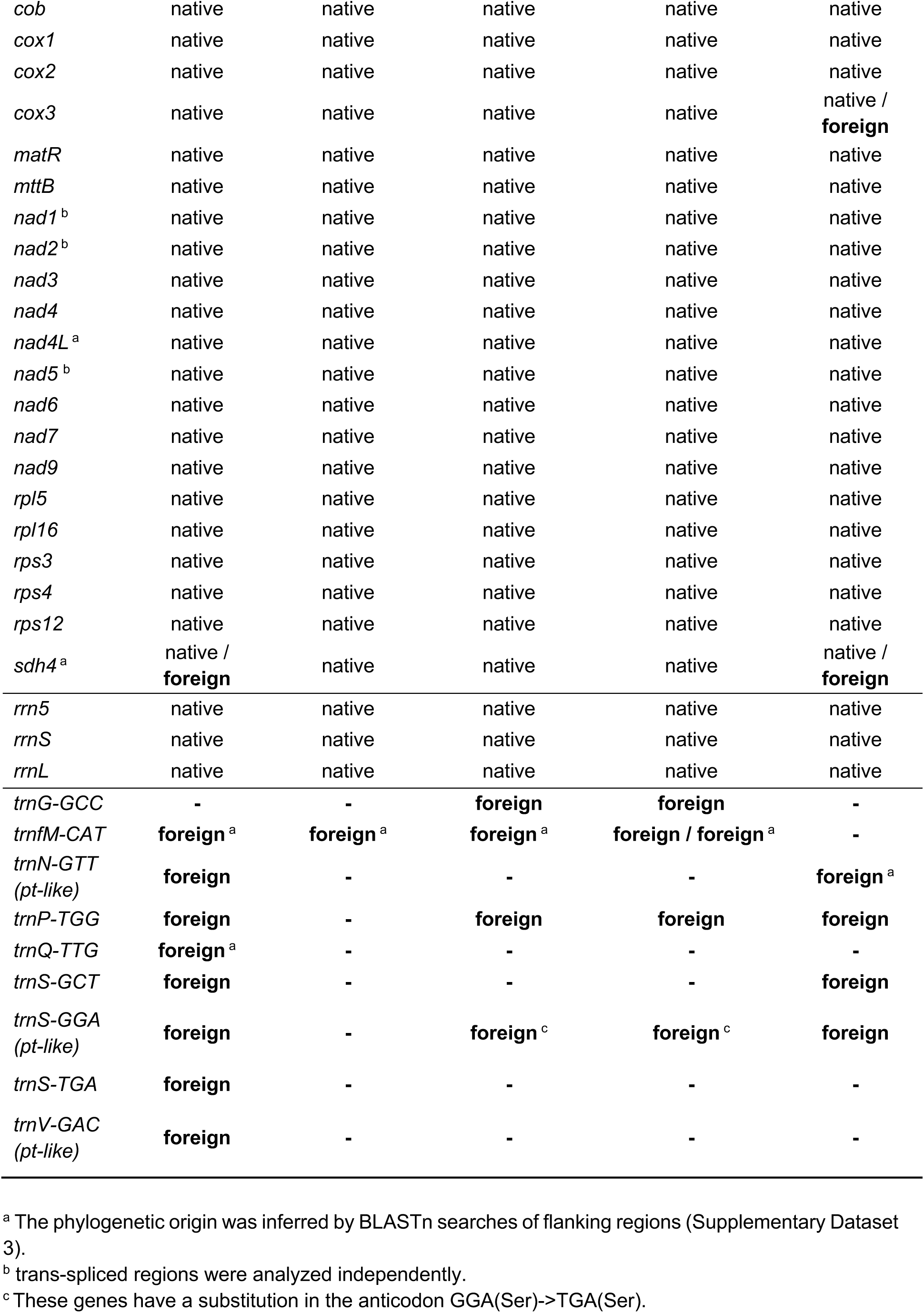
Phylogenetic origin of mitochondrial genes of Rafflesiaceae.

Noticeably, no native mitochondrial tRNA genes were identified in Rafflesiaceae mtDNAs (Figure S7, Table 2). Seven tRNAs detected in *Sapria*, six or seven in *Rafflesia spp.,* and the single *trnfM-CAT* gene identified in *Rhizanthes* show phylogenetic affiliation and/or higher sequence similarity of their flanking regions with Vitaceae mtDNAs (Figure S7, Supplementary Dataset 3). The only exception is the *trnQ* gene identified in *Sapria* which shows sequence similarity with Araliaceae mtDNAs (Supplementary Dataset 3). The three and two tRNAs of plastidic origin in *Sapria* and *Rafflesia* spp., respectively, are embedded in cpDNA-like tracts (MTPTs) that are also present in Vitaceae mitochondrial genomes. They are likely the result of intracellular cp-to-mt transfers in the hosts, followed by mt-to-mt host-to-parasite HGT, as it has been described in other angiosperm mitochondria (Gandini2017). This is likely the case of the other MTPTs identified in *Sapria* since all of them show phylogenetic affiliations with Vitaceae (Figure S8).

### Organellar DNA replication, recombination, and repair genes in Rafflesiaceae

To investigate the genetic basis of the striking differences among Rafflesiaceae mtDNAs, we performed a comparative analysis of nuclear-encoded genes involved in organellar DNA replication, recombination, and repair (DNA-RRR) across the three genera (Table S8). For this, we assembled the transcriptome of *Rhizanthes* and *Rafflesia* species, which contained 66.6-84.5% of the conserved eukaryotic BUSCOs and 45.7-58.4% of the Viridiplantae BUSCOs, either complete or fragmented (Table S9). These values are comparable with those of the complete nuclear genome of *Sapria* (84.5% and 44.6%, respectively; (Cai *et al*., 2021)) and are broadly consistent with previous assessments based on the same *Rhizanthes* transcriptomic dataset (Yu *et al*., 2026). The high proportion of missing BUSCOs is expected for holoparasitic plants as previously reported (Cai *et al*., 2021; Yu *et al*., 2025; Roulet *et al*., 2026).

Using *Arabidopsis thaliana* as a reference framework, nuclear-encoded DNA-RRR genes targeted exclusively to the plastid were found to be extensively lost in the Rafflesiaceae (Table S8), a pattern consistent with previously reported cases of complete plastid genome elimination in this family (Molina *et al*., 2014; Cai *et al*., 2021; Yu *et al*., 2026). This genomic depletion was further supported by our failure to recover any plastid-derived sequences from the *de novo* assemblies of *Rhizanthes lowii*, *Sapria*, or *Rafflesia* species. In contrast, the majority of mitochondrial- and dual-targeted DNA-RRR loci are well-retained across the family, underscoring stringent selective constraints on the maintenance of the mitochondrial genome.

Our profiling revealed distinct shared patterns of gene erosion: *ODB1* and *DJ1D* were the only exclusively mitochondrial-targeted DNA-RRR genes absent from all three genera, while four dual-targeted genes, *CRY3*, *PHR1*, *RECX*, and *UVR3*, were also uniformly missing. These shared absences point to ancestral deletion events that occurred prior to the divergence of the Rafflesiaceae. Independent interrogations of the two available nuclear genomes of *Sapria* yielded identical profiles, and were generally congruent with the inventory reported by Guo et al. (Guo *et al*., 2023). Finally, phylogenetic analyses of the DNA-RRR genes found in Rafflesiaceae confirmed their native, vertical origin and validated their orthology assignments (Figure S9). Differences among genera are limited to the dual targeted RECG1 gene, which was not detected in the *Rhizanthes* transcriptome. Additionally, there are a few absences restricted to a single species of *Rafflesia* (Table S8), which might represent false positives or species-specific losses.

## DISCUSSION

The transition to a holoparasitic lifestyle represents one of the most radical evolutionary shifts in plants. Because heterotrophic plants maintain a continuous reliance on cellular respiration and other vital mitochondrial metabolic pathways, their mitochondrial genomes (mtDNAs) are expected to experience far less evolutionary erosion than their photosynthetic-defective plastomes. However, because parasitic plant mtDNAs have historically been studied less extensively than plastomes, fully testing this hypothesis has proven challenging. Comparative mitogenomics has so far revealed a striking lack of evolutionary convergence in structure, gene content, nucleotide composition, and genome size across diverse holoparasitic lineages (Zervas *et al*., 2019; Petersen *et al*., 2020; Yu *et al*., 2025). Furthermore, while a universally accelerated nucleotide substitution rate was initially proposed as a hallmark of parasitic plant mitochondria (Bromham *et al*., 2013), broader taxonomic sampling has largely deconstructed this rule (Zervas *et al*., 2019). Instead, the most prominent unifying feature of holoparasite mitogenomes is their propensity to acquire foreign DNA via host-to-parasite horizontal gene transfer (HGT), a phenomenon that varies dramatically in magnitude across independent lineages (Petersen *et al*., 2020; Sanchez-Puerta *et al*., 2023).

The giant-flowered holoparasites of the family Rafflesiaceae provide an extraordinary, yet understudied, system to explore the outer limits of this genomic plasticity. In this study, we provide a comprehensive, comparative analysis of the mitochondrial genomes representing all three extant genera of the family: *Sapria*, *Rhizanthes*, and *Rafflesia*. Our findings expose an extraordinary level of structural differences and asymmetric host DNA integration that refines our understanding of organellar evolution under extreme parasitic constraints.

### Shared and divergent mitochondrial features across Rafflesiaceae

The mitochondrial genomes of the Rafflesiaceae represent some of the most structurally divergent organellar systems reported in angiosperms to date. The extraordinary variation in genome size, spanning from the expanded 824 kb mitogenome of *Sapria* to the streamlined 282 kb mitogenome of *Rhizanthes*, underscores a profound lack of structural constraint. Equally striking is the absolute absence of syntenic or shared mitochondrial chromosomes among these closely related genera. This extreme degree of sequence turnover strongly suggests that the mitogenome of the Rafflesiaceae common ancestor underwent severe genomic transformation. This was followed by independent, lineage-specific stabilization into highly multichromosomal architectures, resulting in 40 circular chromosomes in *Sapria* and 35 in *Rhizanthes*.

Remarkably, beneath this layer of structural divergence lies a highly conserved functional core. All three Rafflesiaceae genera maintain an identical complement of 30 protein-coding genes (excluding duplicated copies) and exhibit a highly uniform nucleotide composition, with GC contents tightly clustered between 45.3% and 46.7%. This constancy indicates that while the mechanisms governing genome physical structure and dynamics have relaxed, purifying selection on mitochondrial respiration remains uncompromised. In stark contrast to this core conservation, our comparative analysis reveals that these mtDNAs have taken highly divergent evolutionary trajectories regarding their repetitive landscapes (ranging from 7% to 36%) and their susceptibility to host-derived HGT, which accounts for an astonishing 0.8% to 60% of total genomic content.

### Asymmetric HGT dynamics and mitogenomic repeat content across Rafflesiaceae

Our comparative framework demonstrates that mitochondrial HGT within the family Rafflesiaceae exhibits highly divergent, asymmetric patterns, with minimal, intermediate, and exceptionally high levels observed in *Rhizanthes*, *Rafflesia*, and *Sapria*, respectively. This stark intra-familial variation directly accounts for the remarkable differences in mitogenome sizes across these genera, most notably driving the massive genome expansion captured in *Sapria*. Our analyses reveal that approximately 60% of the *Sapria* mitochondrial genome was acquired horizontally from its *Tetrastigma* host, an extraordinary proportion that aligns with the upper limits of host-derived integration documented in other extreme holoparasites, such as *Lophophytum* spp. (Sanchez-Puerta *et al*., 2019) and *Mitrastemon yamamotoi* (Roulet *et al*., 2026). Conversely, *Rhizanthes* exhibits virtually no intergenic HGT, despite sharing an identical endoparasitic lifestyle and host genus with *Sapria*. This profound disparity in foreign DNA retention echoes the evolutionary dynamics observed within the Balanophoraceae, where genera like *Balanophora*, *Thonningia*, and *Rhopalocnemis* display negligible HGT impacts in stark contrast to the record-setting *Lophophytum mirabile*, whose host-derived core constitutes nearly 75% of its mitochondrial genome (Roulet *et al*., 2024), expanding to over 93% when accounting for its total pan-mitogenome (Gatica-Soria *et al*., 2025).

Historically, variable levels of HGT across parasitic plants have been attributed to different degrees of nutritional dependence and the physical intimacy of the host-parasite physiological bridge (Yang *et al*., 2016). For instance, holoparasitic Orobanchaceae species exhibit substantially higher rates of host-derived nuclear HGT than their hemiparasitic relatives (Yang *et al*., 2016; Kado & Innan, 2018; Yoshida & Kee, 2021). Furthermore, the presence of an extended vegetative endophytic stage, such as the one shared by the Rafflesiaceae, has been recognized as a primary anatomical facilitator for the frequent exchange of mitochondrial DNA fragments (Sanchez-Puerta *et al*., 2023).

However, the wide spectrum of HGT retention we observed among the fully endoparasitic Rafflesiaceae forces a reconsideration of purely anatomical models, indicating that cell-biological and genomic features act as critical checkpoints to enable or limit the stable integration of foreign mtDNA. We hypothesize that a strong deletion bias in the mitochondrial DNA maintenance machinery may actively purge foreign non-coding sequences, preventing genome expansion. Another relevant aspect is the massive proliferation of short repeats, which comprise over 25% of the *Rhizanthes* mitogenome, a phenomenon also documented in *Balanophora* spp. (Yu *et al*., 2025). Mechanistically, the lineage-specific absence of the dual-targeted DNA repair gene *RECG1* in *Rhizanthes*, which is involved in suppressing recombination at short repeat clusters (Wallet *et al*., 2015), may provide a molecular explanation for the highly repetitive landscape as a result of increased replication slippage and/or microhomology-mediated repair pathways. Additionally, population genetic dynamics, structural constraints imposed during cellular divisions in the endophyte, and variation in mitochondrial fusion-fission frequencies across the three genera likely intersect to shape these distinct genomic architectures.

### Chromosome-specific concentration of host-derived DNA in *Sapria*

Notably, the host-derived sequences within the *Sapria* mitogenome are not uniformly distributed across its architecture. Instead, foreign DNA is concentrated into specific circular molecules, most of which lack known functional genes and effectively function as non-coding chromosomes. Among the 40 circular chromosomes constituting the mitogenome, ten exhibit an exceptional >90% sequence length identity to available *Tetrastigma* mtDNAs and are classified as entirely foreign. An additional five non-coding chromosomes display a substantial 71–88% host affinity. Because genomic data from the exact sympatric *Tetrastigma* host species remains unavailable, our current estimations likely represent a conservative baseline; hence, the actual proportion of foreign sequence integration could be even higher.

The structural isolation of these 15 predominantly foreign chromosomes strongly supports the hypothesis that they originated via the circularization of host mitochondrial DNA fragments, a phenomenon conceptualized in the circle-mediated HGT model (Roulet *et al*., 2024). The mechanistic predictions of this model have gained empirical support in other holoparasites. For instance, in *Mitrastemon yamamotoi*, 60% of the mitochondrial genome is horizontally acquired and similarly compartmentalized into seven entirely foreign circular chromosomes (Roulet *et al*., 2026). This structural partitioning is also reminiscent of the multi-chromosomal architectures of *Lophophytum mirabile* and *L. pyramidale*, where the massive influx of foreign genetic material led to the stabilization of independent, host-derived circular chromosomes (Roulet *et al*., 2024; Gatica-Soria *et al*., 2025). Taken together, these parallel patterns across phylogenetically distant holoparasites indicate that the sorting and maintenance of foreign DNA in circular non-coding blocks is a recurring architectural solution to massive host-to-parasite genetic transfers.

### Reduced functional impact of HGT in Rafflesiaceae mitochondrial genes

The primary evolutionary outcome of mitochondrial HGT in angiosperms is typically duplicative transfer, wherein foreign and native gene copies temporarily coexist within the recipient genome (Mower *et al*., 2010; Rice *et al*., 2013; Sanchez-Puerta, 2014; Roulet *et al*., 2020). Under relaxed selective constraints, these redundant foreign copies commonly degenerate into pseudogenes and are eventually purged from the genome (Rice *et al*., 2013). Alternatively, homologous recombination between native and foreign counterparts can trigger gene conversion, yielding stable chimeric genes through a process of partial replacement HGT (Hao *et al*., 2010; Mower *et al*., 2010). Such genomic mosaicism heavily complicates phylogenetic inference derived from mitochondrial genes, underscoring the need for heightened caution when interpreting complex organellar evolutionary histories in parasitic plant systems.

Notably, the number of intact foreign mitochondrial genes identified in our Rafflesiaceae datasets is substantially smaller than previously reported for this family (Xi *et al*., 2013; Guo *et al*., 2023). This discrepancy reflects critical differences in assembly completeness, annotation criteria, and phylogenetic stringency. For instance, several loci previously classified as functional foreign genes in *Rafflesia* and/or *Sapria*, such as *cox3*, *rpl2*, and *rps1*, were recovered in our assemblies as truncated open reading frames and are thus annotated here as non-functional foreign pseudogenes. Furthermore, our high-resolution *Sapria* assemblies failed to recover several foreign copies described by Xi et al. (Xi *et al*., 2013), including *cob*, *cox1*, and *atp9*. Other previously reported foreign candidates (*rpl5*, *rps14*, and *sdh3*) were identified only in our preliminary raw assemblies and were subsequently traced to mitochondrial insertions within the nuclear genome (NUMTs, NoteS1).

We also found no robust support for the proposed Cucurbitaceae origin of the single-copy *atp4* gene in *Rafflesia* spp. and *Sapria* (Xi *et al*., 2013). In our extended analyses, *atp4* lacked definitive phylogenetic affinity to any angiosperm lineage and was consequently categorized as native. Similarly, the single-copy *atp6* and *nad4L* genes in *Sapria*, previously inferred to be acquisitions from Fagaceae and Vitaceae, respectively (Guo *et al*., 2023), displayed no compelling evidence of foreign origin in our phylogenies. Collectively, these reassessments strongly imply that earlier workflows overestimated the functional scale of mitochondrial HGT in this family due to limitations associated with incomplete genome assemblies and/or methodological artifacts. Conversely, we do suggest the potential presence of chimeric genes (e.g., *ccmB*, *ccmC*, *cox2*, and *nad7*), which are characterized by weak bootstrap support for an affiliation with their hosts, suggesting that partial gene conversion represents the primary mechanism extending the functional footprint of HGT in these parasites.

To date, *atp1* stands out as the only core mitochondrial gene that has been fully and functionally replaced by a host-derived homolog across the Rafflesiaceae. The stable inheritance of this foreign *atp1* gene across all three examined genera demonstrates an ancestral, family-wide replacement event that occurred prior to generic divergence. In contrast, the remaining intact foreign genes identified in our study co-exist alongside their native counterparts (such as the duplicated *sdh4* in *Sapria* and *Rafflesia leonardi*). Based on functional insights from other massive HGT systems, such as *Amborella trichopoda* and *Lophophytum* spp., where co-existing native alleles remain the exclusively transcribed and functional copies (Rice *et al*., 2013; Garcia *et al*., 2021; Roulet *et al*., 2024), we hypothesize that these supplementary foreign copies are largely transcriptionally silent or evolutionary dead-ends.

### The DNA-RRR complement as a driver of mitogenomic architecture in Rafflesiaceae

A combination of genomic and life-history factors likely underlies the evolution of holoparasite organellar genomes. The loss of photosynthesis, normally the dominant metabolic process in photoautotrophs, has been proposed to shift the population-genetic regime toward relaxed purifying selection, reduced effective population size, and cascading gene loss across plastid and nuclear genomes (Cai *et al*., 2021).

While many of these genomic deletions are direct structural consequences of losing photosynthetic function, others reflect broader systemic responses to heterotrophy and altered selective pressures. Prominent among these are the losses of nuclear genes encoding organellar DNA replication, recombination, and repair (DNA-RRR) proteins, which have been increasingly linked to the anomalous architectural features observed in diverse parasitic lineages (Ceriotti *et al*., 2022; Yu *et al*., 2022; Yu *et al*., 2025; Roulet *et al*., 2026).

In the Rafflesiaceae, specific DNA-RRR gene absences may have catalyzed mitochondrial structural destabilization or facilitated plastome degradation leading to its ultimate elimination, whereas other losses may represent indirect signatures of their parasitic lifestyle. We infer that fifteen DNA-RRR genes were lost in the common ancestor of the Rafflesiaceae, exhibiting a pronounced and asymmetric bias toward plastid-targeted components. This genomic erosion strongly correlates with the putative loss of the plastid genome in this family (Molina *et al*., 2014; Cai *et al*., 2021; Yu *et al*., 2025). It further suggests that the disruption of the plastid maintenance machinery was a foundational, potentially causal step in its evolutionary decay. As documented in other non-photosynthetic plants with highly degraded plastomes (Schelkunov *et al*., 2019; Schelkunov *et al*., 2021; Ceriotti *et al*., 2022; Yu *et al*., 2025), the collapse of plastid maintenance is typically accompanied by elevated substitution rates, extreme AT bias, and genome compaction prior to complete loss. Notably, *ARP*, which encodes an apurinic/apyrimidinic endonuclease critical for plastid base excision repair(Gutman & Niyogi, 2009; Córdoba-Cañero *et al*., 2014), is the sole plastid-targeted RRR gene retained in the family. Given the putative absence of a physical plastome in these taxa, its unexpected retention strongly implies functional targeting to the mitochondria or nucleus, consistent with evidence that ARP proteins can fulfill multi-compartmental repair roles under stress (Chevigny *et al*., 2020). Furthermore, there is evidence for ARP presence and activity in the mitochondria of *Arabidopsis* and *Solanum tuberosum* (Boesch *et al*., 2010; Ferrando *et al*., 2019; Fuchs *et al*., 2020).

Notably, dual-targeted photolyases *CRY3*, *PHR1*, and *UVR3*, which mediate the repair of UV-induced DNA lesions, are entirely absent across all examined Rafflesiaceae lineages. Because these loci represent the complete photolyase repertoire in angiosperms, their total loss is remarkable. A convergent multi-gene deletion has been documented in the common ancestor of the Balanophoraceae and attributed to the “deep-shade” hypothesis, where a lifetime spent inside host tissues or underground relaxes selection on light-dependent repair functions (Yu *et al*., 2025). Given that the vegetative structures of the Rafflesiaceae develop exclusively as endophytes within underground host roots, an identical evolutionary relaxation scenario is highly plausible. This interpretation is further supported by the root-endoparasitic Mitrastemon, which also lost the photolyases CRY3 and UVR3 (Roulet *et al*., 2026).

The structural integrity of the mitochondrial genome may have been compromised by other ancestral losses of DNA repair genes. In addition to photolyases, the dual-targeted recombination surveillance factor RECX and the mitochondrially targeted organellar DNA-binding protein 1 (ODB1) are universally absent in the Rafflesiaceae. Because both proteins regulate homologous recombination and maintain structural stability in plant mitochondria (Janicka *et al*., 2012; Odahara & Sekine, 2018), their ancestral absence likely explains the profound physical reshuffling and lack of chromosomal synteny observed among the three genera. Another shared missing factor is DJ1D, which is responsible for repairing glycated DNA and RNA in plant mitochondria (Prasad *et al*., 2022). The loss of DJ1D, in concert with ODB1 and RECX, likely weakened the efficiency of homologous recombination, the primary high-fidelity DNA repair pathway in plant mitochondria (Christensen, 2013; Chevigny *et al*., 2020), contributing to the moderately elevated substitution rate in Rafflesiaceae mitogenomes.

While the shared absence of these core DNA-RRR components is robustly supported by congruent data across the five transcriptomes and two nuclear assemblies, we also identified lineage-specific variations with much weaker support. Most notably, the dual-targeted gene *RECG1* was absent from the *Rhizanthes* transcriptome. Although this sporadic absence should be verified with complete genomic data for *Rhizanthes*, it provides a molecular rationale for the idiosyncratic features of its mitogenome, where the lack of *RECG1*-mediated suppression explains the massive expansion of short repeat families.

### Remodeling of mitochondrial translation following putative plastid genome loss

The evolutionary transition to obligate holoparasitism entails the complete loss of photosynthesis, which triggers an extensive remodeling of organellar translation systems, including the systematic purging of plastid-encoded tRNAs (Wicke & Naumann, 2018; Ceriotti *et al*., 2021; Kim *et al*., 2023; Ceriotti *et al*., 2025) and associated translational components (DeTar *et al*., 2024). Relaxed purifying selection on plastid function appears to initiate a cascade of reductive evolution that destabilizes the plastid translational machinery, exerting correlated, systemic effects on mitochondrial translation. In the Rafflesiaceae, where both plastid genomes and plastid translation have been likely eliminated (Molina *et al*., 2014; Cai *et al*., 2021; Yu *et al*., 2026), these evolutionary repercussions are expected to be uniquely pronounced within their mitochondria.

Indeed, one of the most extraordinary features of the Rafflesiaceae mitogenomes is the entire absence of native mitochondrial tRNA genes. Across all examined genera, the remaining mitochondrial tRNA complement is almost exclusively host-derived: *Sapria* encodes eight tRNAs, *Rafflesia* species harbor six to seven, and *Rhizanthes* retains solely a single *trnfM* gene. This complete replacement aligns with growing evidence that plant mitochondrial tRNA repertoires are evolutionarily labile and frequently reshaped by horizontal acquisitions (Warren *et al*., 2025; Ceriotti *et al*., 2026). While many foreign mitochondrial insertions often degenerate into non-functional pseudogenes, multiple benchmark studies have demonstrated the expression, and occasionally the aminoacylation, of horizontally acquired tRNAs of plastid (Joyce & Gray, 1989; Marchfelder *et al*., 1990; Ceriotti *et al*., 2026), bacterial (Kitazaki *et al*., 2011; Knie *et al*., 2015), fungal (Sinn & Barrett, 2020; Warren *et al*., 2025), or foreign mitochondrial origin (Ceriotti *et al*., 2026). These functional precedents establish that foreign tRNAs can be successfully integrated into host organellar translation systems.

In typical angiosperms, the organellar tRNAs are aminoacylated by their corresponding nuclear-encoded, dual-targeted organellar aminoacyl-tRNA synthetases (aaRSs) (Duchêne et al., 2005). Previous surveys in non-photosynthetic lineages suggested that the retention of these organellar aaRSs strictly correlates with the remaining organellar tRNA content (DeTar *et al*., 2024; Ceriotti *et al*., 2026). The Rafflesiaceae, however, represent a radical and unprecedented endpoint of this evolutionary trajectory. In *Sapria* and *Rafflesia*, all but a single organellar aaRS (PheRS) have been entirely purged from the nuclear genome, constituting the most severe collapse of organellar aaRS genes ever documented in plants (DeTar *et al*., 2024), which perfectly mirrors the complete loss of native plastid and mitochondrial tRNAs. Consequently, mitochondrial tRNA aminoacylation in these parasites must rely almost entirely on cytoplasmic aaRSs imported into the organelle, presenting two distinct evolutionary scenarios. First, the majority of the horizontally acquired mitochondrial tRNAs may be transcriptionally silent or non-functional, leaving mitochondrial translation completely dependent on the direct import of both cytosolic tRNAs and their cognate cytoplasmic aaRSs. Alternatively, these foreign mitochondrial tRNAs may be functional and aminoacylated by the cytoplasmic aaRSs within the mitochondrial matrix. Whichever scenario is true, the Rafflesiaceae provide an unparalleled natural laboratory for testing the absolute boundaries of coordinated coevolution and translational flexibility following massive gene loss and horizontal gene acquisition.

Comparative insights from other holoparasitic lineages further highlight the idiosyncratic nature of these translational dynamics. *Lophophytum* (Balanophoraceae) is the only other angiosperm lineage known to harbor an entirely foreign mitochondrial tRNA complement (Roulet *et al*., 2024). In that system, most foreign tRNAs are transcriptionally active and correctly processed, signaling stable functional assimilation, while only a subset shows evidence of ongoing pseudogenization (Ceriotti *et al*., 2026). Across the Balanophoraceae, the incorporation of foreign tRNAs follows a direct replacement model, where foreign homologs substituted for native mitochondrial genes prior to or immediately following their loss; a process accompanied by the strict nuclear retention of their corresponding organellar aaRSs (Ceriotti *et al*., 2026). In sharp contrast, the Rafflesiaceae display a uniquely derived configuration characterized by the near-complete loss of organellar aaRS loci. This highlights a distinct, far more radical evolutionary pathway of organellar translation remodeling, where the parasite has shattered the traditional organellar (bacterial-like) infrastructure in favor of a structural reliance on cytoplasmic translational components.

## Materials and Methods

### Sample collection, DNA extraction, and sequencing

Individuals of *Sapria himalayana* (Individual 1) and *Rhizanthes lowii* were collected in Xishuangbanna, Yunnan, China, and Kapit, Sarawak, Malaysia, respectively. In both cases, individuals were found parasitizing the roots of *Tetrastigma* spp. Flower tissue was used to minimize host-derived contamination. Total genomic DNA was extracted using the CTAB protocol, and Illumina libraries with an average insert size of 350 bp were constructed and sequenced on the Illumina Novaseq 6000 platform. Clean reads were deposited in the NCBI Sequence Read Archive under accession numbers SRR37894703 and SRR37894702 (Table S10).

### Assembly of mitochondrial genomes

The sequencing data was used to assemble the mitochondrial genomes of *Sapria* (Individual 1) and *Rhizanthes* with SPAdes v3.15.2 (Bankevich et al., 2012) using the parameters --careful --only-assembler -k 31,51,77. Mitochondrial contigs were identified by BLASTn searches (Camacho *et al*., 2009) against a custom database of mitochondrial sequences from Rafflesiaceae and Vitaceae retrieved from NCBI (Table S11). The mitochondrial contigs were visualized and manually joined with Bandage v0.8.1 (Wick *et al*., 2015). Circularity of mitochondrial chromosomes was confirmed by inspection of paired-end read with CONSED v29 (Gordon & Green, 2013), specifically by identifying read pairs whose mates mapped to the opposite ends of the same mitochondrial contigs and verifying consistent coverage across junctions. In addition, an independent mitochondrial assembly was generated for a second sample of *Sapria himalayana* (Individual 2, also collected in Yunnan, China by Guo et al., 2023), as well as for individuals of *Rafflesia arnoldii* and *Rafflesia tuan-mudae*, using reads downloaded from SRA (Table S11) and processed with the same workflow described above. The DNA read depth for the mitochondrial assemblies was calculated using Bowtie2 (Langmead & Salzberg, 2012) with the parameters *--end-to-end* and *--very-sensitive*. The graphical representations of the organellar genomes and their corresponding read-depth profiles were generated in R. The final assembled chromosomes of *Sapria* individuals 1 and 2 and *Rhizanthes* have average read depths of ∼350-600x, ∼900-2750x, and ∼1000-2000x, respectively (Figure S10). In addition, we inspected PacBio reads (SRR18272677) generated by Guo et al. (Guo *et al*., 2023) for *S. himalayana* individual 2 and identified head-to-tail concatemers of each of the mitochondrial chromosomes including two or three intact chromosome units, which can lead to individual circular chromosomes.

Because publicly available mitochondrial genomes of known Rafflesiaceae hosts, i.e. *Tetrastigma* spp. (tribe Cayratieae, family Vitaceae), are limited, we expanded the reference dataset to improve the phylogenetic inference of the parasite mitochondrial sequences. Accordingly, we assembled the complete mitochondrial genomes of two *Tetrastigma spp.* (Individual 1 and Individual 2) and their plastid genomes for taxonomic purposes (Note S2; Table S10; Figure S11; Supplementary Dataset 4). The DNA read depth of both *Tetrastigma* mtDNAs was calculated and visualized as detailed above (Figure S10).

To better capture host (i.e. putative donors) diversity, additional DNAseq data were retrieved from SRA for five of the eight recognized genera within the tribe Cayratieae (Table S11). Only DNAseq datasets exceeding 500 MB were selected to ensure sufficient mitochondrial read depth. This dataset included 15 *Tetrastigma* individuals, representing ∼16% of the currently recognized species in the genus (Wen *et al*., 2018), as well as species from four other genera of the tribe. In total, mitochondrial sequence data were obtained for 19 individuals across the tribe Cayratieae (Table S11). For each individual, mitochondrial sequences were assembled with SPAdes (--careful --only-assembler -k 31,51,77). Candidate mitochondrial contigs were identified through BLASTn searches against a custom database of mitochondrial sequences from Vitaceae (Table S12) and visualized in Bandage. To reduce redundancy and produce streamlined species-level assemblies, overlapping contigs were merged in Geneious Prime v2024.0.3 using a minimum overlap of 200 bp and requiring 100% identity across the shared region.

### Mitochondrial genome characterization

The mitochondrial genomes of Rafflesiaceae and *Tetrastigma* were annotated using Geneious v.11.1.5 and through BLASTn searches (Camacho *et al*., 2009) with angiosperm mitochondrial genes as queries. The identification of tRNA genes was performed using the tRNAscan-SE algorithm (Lowe & Eddy, 1997). To detect plastid-derived mitochondrial sequences (MTPTs) longer than 100 bp, BLASTn searches were conducted against a custom angiosperm chloroplast database (Table S13A). Repeats were annotated and clusterized based on sequence similarity using the *get_repeats.sh* script developed by Gandini et al. (Gandini *et al*., 2019) specifying - perc_identity flag 90. The genomic coordinates of all annotated repeats have been deposited in the Supplementary Dataset 5. For a graphical view, repeat regions were identified with LASTZ v1.0.4 and visualized in AliTV v1.0.6 (Ankenbrand *et al*., 2017), retaining only homologous blocks longer than 400 bp and with >90% nucleotide identity.

### Collinearity analysis

Collinearity analyses were performed to assess both interspecific and intraspecific variability of the mitochondrial genome structure within Rafflesiaceae. Homologous regions were identified using BLASTn searches, and pairwise whole-genome alignments were generated with LASTZ v1.0.4 and visualized in AliTV v1.0.6 (Ankenbrand *et al*., 2017), retaining homologous blocks longer than 1 kb and exhibiting >70% nucleotide identity. Interspecific comparisons were conducted among the mitochondrial genomes of *Sapria*, *Rhizanthes*, and *Rafflesia* spp. Intraspecific variation was evaluated by comparing the two *Sapria* individuals assembled in this study.

### Phylogenetic origin of mitochondrial sequences in Rafflesiaceae

To infer the phylogenetic origin of the mitochondrial DNA in the endoparasitic Rafflesiaceae, each mitochondrial chromosome/contig was queried individually using BLASTn against a customized angiosperm mitochondrial database (Table S12). To visualize the distribution of BLASTn matches, all BLASTn hits with an e-value < 2 × 10⁻^10^ were plotted using the Sushi R package v1.20.0 (Phanstiel *et al*., 2014). Hits were arranged by taxonomic category to facilitate interpretation. When multiple overlapping matches occurred within a category, the highest-scoring hit was plotted above the others to enhance visual clarity. Plots available at Supplementary Dataset 3.

The proportion of putative foreign sequence per mitochondrial chromosome/contig was estimated from the BLASTn searches (e-value < 2 × 10⁻^10^). Hits to *Tetrastigma* mtDNA with lengths >200 bp and with >90% identity were retained when they were the best hits relative to other taxonomic groups. Redundant and overlapping matches were excluded. These criteria follow established approaches for detecting HGT events, integrating both (i) high sequence similarity and (ii) phylogenetically irregular distribution (Bock, 2010; Wickell & Li, 2020; Aubin *et al*., 2021; Mariault *et al*., 2025). Proportions of foreign matches were summarized and visualized using the ggplot2 package in R (Wickham *et al*., 2016).

### Phylogenetic origin of the genes

To assess the origin of the mitochondrial genes in Rafflesciaceae, phylogenetic analyses were conducted using a Maximum Likelihood (ML) approach. Nucleotide sequences of protein-coding genes were extracted from various angiosperms mtDNAs (Table S13B), and trans-spliced gene regions were analyzed separately. For tRNAs the analysis was extended to the flanking regions, including BLASTn hits >250 bp against a local database of angiosperm mtDNAs (Table S12) and considering only one sequence per genus. In addition, we analyzed the origin of MTPTs >400 bp in length. Alignments were prepared with MAFFT v.7.407 using -localpair -maxiterate 1000 options, and poorly aligned regions were removed with BMGE 1.12 (Supplementary Dataset 6). The best substitution model for each alignment was estimated using the ModelFinder Plus option of IQ-TREE2 v2.2.0 [Minh et al. 2020] and ML trees were inferred with RAxML v.8.2.11 (Stamatakis, 2014) under the GTR + gamma model, including 1000 rapid bootstrap pseudoreplicates. The resulting trees were visualized using FIGTREE v.1.4.4. For phylogenetic inference of genic regions, RNA editing sites previously identified in multiple angiosperms (Edera *et al*., 2018) were excluded from the alignments.

### Identification of organellar DNA-RRR genes

To identify DNA replication, recombination, and repair (DNA-RRR) genes encoding proteins targeted to organelles, we used transcriptomic data from species of *Rhizanthes, Rafflesia* and two nuclear genome assemblies from *Sapria*. For *Sapria*, the nuclear genome assemblies of *Sapria himalayana* (GCA_029783825.1 and GCA_016808135.1; (Cai *et al*., 2021; Guo *et al*., 2023) were used. *De novo* transcriptome assemblies of *Rhizanthes zippelii* (SRR13325878; (Cai *et al*., 2021)), *Rafflesia cantleyi* (SRR518525; (Xi *et al*., 2012)), *Rafflesia tuan-mudae* (SRR13325879; (Cai *et al*., 2021)), and *Rafflesia speciosa* ((Molina *et al*., 2025); SRR24479220) were generated from publicly available RNA-seq datasets retrieved from NCBI using Trinity v2.15.0 (Grabherr et al., 2011), using default parameters and specifying a non–strand-specific library type and --min_contig_length 100. Open reading frames (ORFs) were predicted from assembled transcripts using TransDecoder specifying a minimum length of 80 amino acids. Transcriptome assembly completeness was assessed with BUSCO v6.0 (Tegenfeldt *et al*., 2025) run in genome mode, using the following datasets: eukaryota_odb12 (129 genes), viridiplantae_odb12 (822 genes), embryophyta_odb12 (2,026 genes), and eudicotyledons_odb12 (2805 genes).

We focused on a curated set of 39 DNA-RRR genes whose products are known to be targeted to mitochondria and/or plastids (Gualberto & Newton, 2017; Ceriotti *et al*., 2022; Schatz *et al*., 2025). Sequences from *Arabidopsis thaliana* were used as queries in tBLASTx and BLASTp searches against the assembled transcriptomes and predicted protein datasets of *Rhizanthes* and *Rafflesia*, as well as against the genomic datasets of *Sapria*, using an e-value threshold of 1 × 10⁻³. When an RRR gene was detected in one genus but not in the others (*Rafflesia*, *Sapria* or *Rhizanthes*), a second search was performed using the sequence identified in the detected species as query. Identified ORFs were verified if they correspond to homologs of the Arabidopsis gene used as query by making reciprocal BLAST searches. A third round of searches was subsequently performed using HMMER (hmmsearch) against the *Rafflesia*, *Sapria*, and *Rhizanthes* datasets, employing domain profiles inferred from InterPro annotations together with Hidden Markov Models obtained from Pfam and PANTHER. Phylogenetic analyses of all identified genes were performed using RAxML v8.2.11, as described above, to verify orthology assignments.

## Data Availability statement

The data underlying this article are available in Figshare, at https://figshare.com/account/mycontent/projects/278234 and *in* the GenBank Nucleotide Database at https://www.ncbi.nlm.nih.gov/nucleotide/, *and can be accessed with* accession numbers PZ290500-PZ290534, PZ290541-PZ290580, and PZ584705-PZ584712.

## Supplementary information

**Note S1.** Mitochondrial genome assemblies of two *Sapria himalayana* individuals.

**Note S2.** Assembly of *Tetrastigma sp.* mitochondrial and plastid genomes.

**Figure S1. Collinearity among mitochondrial sequences from two individuals of *Sapria himalayana*.** Gray blocks represent mitochondrial chromosomes, with block width proportional to sequence length. Internal colored lines denote homologous regions (>1 kb, >70% identity), with color intensity reflecting sequence identity (70– 100%). Alignments were generated with LASTZ v1.0.4 and visualized using AliTV v1.0.6. Both individuals were collected in Yunnan, China.

**Figure S2. Read mapping profiles revealing structural misassemblies in previously published *Sapria himalayana* mitogenomic contigs.** (A–D) Sequencing coverage (read depth) across two mitochondrial contigs originally assembled by Guo et al. (2023) (GenBank accessions OQ719965 [Chr1] and OQ719978 [Chr14]). Raw paired-end reads from two *Sapria* individuals (Individual 1 and Individual 2) were mapped to these reference sequences. Coverage profiles are shown for OQ719965 using reads from Individual 2 (A) and Individual 1 (C), and for OQ719978 using reads from Individual 2 (B) and Individual 1 (D). Read depth was calculated using Bowtie2 v2.4.4 with the parameters --end-to-end --very-sensitive -- no-contain --no-discordant --no-mixed. Below each coordinate axis, homologous regions are arbitrarily colored based on BLASTn alignments against the assembled *Sapria* mitogenome (Individual 1), selected mitochondrial coding genes (*sdh3*, *rpl5*, *rps14*), and the plastome of *Tetrastigma* sp (Individual 1).

**Figure S3. Mitochondrial genomes of Rafflesiaceae species.** (A) Complete mitochondrial genome (mtDNA) of *Rhizanthes lowii*. The mtDNA is 282,577 bp in length and is partitioned into 35 circular chromosomes (Chr) of varying sizes. (B) Draft mitochondrial genome of *Rafflesia tuan-mudae*. (C) Draft mitochondrial genome of *Rafflesia arnoldii*. (D) Draft mitochondrial genome of *Rafflesia leonardi*. Concentric tracks depict genomic and structural features..

**Figure S4. Distribution of host-derived mitochondrial DNA across Rafflesiaceae mitogenomes.** Linear representations of mitochondrial genomes from *Sapria himalayana* (two individuals), *Rhizanthes lowii*, and three *Rafflesia* species, with lengths scaled proportionally to their size. Each rectangle represents a mitochondrial chromosome or contig. Contigs are colored based on the proportion of BLASTn hits (>200 bp and 90% identity) to the mitochondrial data from host genus *Tetrastigma* (Tables S3-S7). White segments correspond to mitochondrial DNA of native or unknown origin. Pie charts summarize the total proportion of foreign DNA detected in each mitogenome. This proportion was calculated by retaining BLASTn hits to *Tetrastigma* mitochondrial DNA longer than 200 bp and with >90% sequence identity, considering only the best hit per region relative to other taxonomic groups. Redundant and overlapping matches were excluded.

**Figure S5. Phylogenetic and sequence similarity evidence of convergent mitochondrial horizontal gene transfer in Rafflesiaceae.** (Top) Maximum-likelihood (ML) phylogenetic reconstructions of two intergenic regions. Branch support values are based on 1,000 rapid bootstrap replicates. Rafflesiaceae taxa are highlighted in magenta. (Bottom) Sequence similarity profile generated with the R package Sushi, showing BLASTn alignments against multiple taxonomic groups. A *Rafflesia leonardi* mitochondrial contig (GenBank accession KJ154982) was used as the reference sequence. The color gradient represents percentage nucleotide identity (pident), ranging from 60% to 100%. Black rectangles and solid arrows indicate the genomic regions analyzed in the phylogenetic reconstructions shown above. Nearby coding sequences are annotated along the coordinate axis.

**Figure S6. Phylogenetic analyses of mitochondrial genes of Rafflesiaceae.** Maximum likelihood analyses were performed with RAxML. ML bootstrap values >50% are shown. The scale bar corresponds to substitutions per site. For species for which only a subset of mitochondrial genes is available, corresponding NCBI accession numbers are provided. Sequences from *Rafflesia cantleyi*, *R. tuan-mudae*, and *Sapria himalayana* with sequence identifiers starting with RC, RT, and SH, respectively, were extracted from Xi *et al*. 2013 supplementary information.

**Figure S7. Phylogenetic analyses of mitochondrial tRNA and rRNA genes of Rafflesiaceae.** Maximum likelihood analyses were performed with RAxML. ML bootstrap values >50% are shown. The scale bar corresponds to substitutions per site.

**Figure S8. Phylogenetic analyses of mitochondrial plastid-derived sequences (MTPTs) >400 bp of *Sapria himalayana*.** Maximum likelihood analyses were performed with RAxML. ML bootstrap values >50% are shown. The scale bar corresponds to substitutions per site.

**Figure S9. Phylogenetic analyses of organellar repair, recombination, and replication (RRR) genes.** Maximum likelihood analyses were performed with RAxML. ML bootstrap values >50% are shown. The scale bar corresponds to substitutions per site.

**Figure S10. DNA read depth of the mitochondrial genomes calculated with Bowtie2 v.2.4.4 (parameters: --end-to-end --very-sensitive --no-contain --no-discordant --no-mixed).** (A) *Sapria himalayana* (individual 1), (B) *Sapria himalayana* (individual 2), (C) *Rhizanthes lowii*, (D) *Tetrastigma* sp. (individual 1), and (E) *Tetrastigma* sp. (individual 2).

**Figure S11. Maximum-likelihood (ML) phylogenetic reconstruction of plastid regions from *Tetrastigma* (Vitaceae).** The phylogeny includes 38 *Tetrastigma* species, including the two individuals assembled in this study (*Tetrastigma* individual 1 and *Tetrastigma* individual 2). Four plastomes from *Cayratia* (Vitaceae) were used as outgroups. The alignment comprises approximately 37 kb of plastid DNA containing 43 plastid genes. Maximum-likelihood analyses were performed with RAxML. Branch support values were estimated from 1,000 rapid bootstrap replicates, and bootstrap values >50% are shown. The scale bar represents substitutions per site.

**Table S1.** Sequence comparison between Rafflesiaceae mtDNAs (BLASTn, e-value<0.0001). (A) *Sapria himalayana* individual 1 versus individual 2 mitochondrial chromosomes. (B) Query coverage (%) of the full length mitochondrial genome assemblies among Rafflesiaceae.

**Table S2**. Repeat landscape characterization of mitochondrial genomes in Rafflesiaceae. A) Summary of repeat content across mitochondrial genomes of Rafflesiaceae. B) Clustering analysis of short repeats in *Rhizanthes lowii* mitochondrial genome.

**Table S3.** Comparative BLASTn coverage of *Sapria himalayana* individual 1 mitochondrial chromosomes across taxonomic categories.

**Table S4.** Comparative BLASTn coverage of *Rhizanthes* sp. mitochondrial chromosomes across taxonomic categories.

**Table S5.** Comparative BLASTn coverage of *Rafflesia leonardi* mitochondrial chromosomes across taxonomic categories.

**Table S6.** Comparative BLASTn coverage of *Rafflesia tuan-mudae* mitochondrial chromosomes across different taxonomic categories.

**Table S7.** Comparative BLASTn coverage of *Raflessia arnoldii* mitochondrial chromosomes across different taxonomic categories.

**Table S8.** Organellar DNA-RRR genes in Rafflesiaceae.

**Table S9.** BUSCO analysis of the transcriptome assembly of *Rhizanthes zipelli* and *Rafflesia species*.

**Table S10.** Sample collection and sequencing information of species analyzed in this study.

**Table S11.** Sequencing datasets from Rafflesiaceae and Vitaceae used to generate de novo mitochondrial genome assemblies and/or genome annotations.

**Table S12.** Accessions of the local angiosperm mitochondrial database.

**Table S13.** GenBank accession numbers of (A) the plastid genomes used in MTPTs identification and (B) the mitochondrial genes included in phylogenetic analyses.

## Supplementary Datasets

Supplementary Datasets available at Figshare: https://figshare.com/account/mycontent/projects/278234

**Supplementary Dataset 1:** Fasta sequences of the mtDNA of *Sapria himalayana* individual 2.

**Supplementary Dataset 2:** Fasta sequences of the draft mitochondrial genomes assembled in this study (Table S11).

**Supplementary Dataset 3:** Plots of the BLASTn hits found when using each Rafflesiaceae mitochondrial genome assembly against an angiosperm genome mitochondrial database (Table S12).

**Supplementary Dataset 4:** Fasta sequence of the cpDNA of *Tetrastigma* sp. individual 1.

**Supplementary Dataset 5:** List of repeats identified in the mitochondrial assemblies of Rafflesiaceae, using the *get_repeats.sh* script developed by Gandini et al. (2019) specifying -perc_identity flag 90.

**Supplementary Dataset 6:** Nucleotide alignments used for the phylogenetic trees.

