## Supplementary material for "Extensive mitogenome divergence across the Rafflesiaceae in size and impact of horizontal gene transfer": Supplementary Information.pdf

**Note S1.** Mitochondrial genome assemblies of two *Sapria himalayana* individuals

The mitochondrial chromosomes 1 and 14 of *Sapria himalayana* assembled as linear molecules by Guo et al. (Guo *et al.*, 2023) and deposited in NCBI databases under accession numbers OQ719965 and OQ719978, respectively, were not recovered in either of the mtDNA assemblies of *Sapria himalayana* of this study, even when using the same dataset as Guo *et al.* (2023) for individual 2.

The mitochondrial genome of *Sapria himalayana* (individual 2 in this study) was previously assembled by Guo *et al.* (2023) as a multichromosomal genome totaling 659,631 bp (160 kb shorter than our assembly using the same dataset) and consisting of 21 circular and 8 linear contigs. The published assembly is entirely contained within our final mitochondrial genome assemblies of *Sapria* individuals 1 and 2, with the exception of chromosomes 1 and 14, which are absent from both of our mitochondrial assemblies. Chromosome 1 is 41,730 bp in length, it is linear and 99% identical to the plastid genome of *Tetrastigma* spp., it shows no evidence of structural rearrangements compared to the cpDNA, and it contains polymorphisms that disrupt several plastid ORFs. This chromosome is evenly covered by sequencing reads from individual 2 but with much lower read depth than the other mitochondrial chromosomes (~200× vs. ~900-2750x shown in Fig S10). Similarly, chromosome 14, which includes foreign copies of *rpl5*, *rps14* and *sdh3* genes, also showed a much lower read depth (~200x) than other mitochondrial chromosomes when mapping with DNAseq reads from individual 2, except for those regions that are shared with other mitochondrial chromosomes (Figure S2). Furthermore, the absence of reads from individual 1 mapping to chromosomes 1 and 14 (with the sole exception of MTPTs and regions already assembled in other mitochondrial chromosomes) indicates that they are not present in the mtDNA of individual 1 (Figure S4). Overall, we conclude that the sequences contained in chromosomes 1 and 14 likely represent mitochondrial- and plastid-derived sequences inserted in the nuclear genome of *S. himalayana* individual 2.

**Note S2.** Assembly of *Tetrastigma sp.* mitochondrial and plastid genomes

Leaves from *Tetrastigma* sp. individual 1 were collected in Yunnan, China. DNA was extracted using the CTAB method and sequenced using Illumina Novaseq 6000 platform (Table S1). The mitochondrial and plastid genomes were assembled using GetOrganelle (Jin *et al.*, 2020). The DNA read depth was ~10x, except for peaks corresponding to o mitochondrial plastid-derived sequences (MTPTs; Figure S10D). The mitochondrial genome assembled into eight circular molecules totaling 680,471 bp (Table S10).

For *Tetrastigma* sp. individual 2, Illumina paired-end reads (151 bp in length) were downloaded from SRA (Table S11) and assembled de novo using SPAdes (Bankevich *et al.*, 2012), with expected coverage and coverage cutoff set to automatic and k-mer lengths of 55, 77, and 99. SPAdes assembled 1,576 contigs longer than 1 kb. Putative mitochondrial and plastidic contigs were identified using BANDAGE v.0.8.1 (Wick *et al.*, 2015) by running BLAST against *Liriodendron tulipifera* mitochondrial CDSs and *Tetrastigma planicaule* cpDNA (NCBI accessions NC_021152 and NC_057118, respectively), applying alignment length and identity thresholds of 100 bp and 95% (Camacho *et al.*, 2009). We identified 36 contigs spanning 287,719 bp with a mean depth of 560.6x, all of which were connected through other contigs in a single structure comprising 131 contigs totaling 826,402 bp with a mean depth of 330.6x. Nine contigs exhibited >98% query cover against the cpDNA, spanning 128,958 bp. The expected single-copy and inverted repeat regions showed sequencing depths of 844.7x and 1,627.8x, respectively. These contigs were manually joined and edited using Illumina paired-end read information in CONSED v.29.0 (Gordon & Green, 2013), resulting in the final 160,500 bp circular cpDNA (Supplementary Dataset 4). Similarly, the mtDNA was obtained by assembling and editing the remaining 122 contigs in CONSED, forming a single 711,396 bp circular chromosome, with relatively uniform coverage of ~500x, except for pronounced coverage peaks corresponding to MTPTs (Figure S10E).

To assess the phylogenetic placement of each individual of *Tetrastigma* analyzed, we conducted a phylogenetic analysis based on a plastid region of approximately 37,000 bp comprising 43 plastid genes. The dataset included the two *Tetrastigma* individuals and 34 species of *Tetrastigma* plus species of the genus *Caryratia* as outgroups. Sequence alignments were generated using MAFFT v7.407 with the -localpair and -maxiterate 1000 options, and poorly aligned regions were removed with BMGE v1.12. Maximum-likelihood trees were inferred using RAxML v8.2.11 (Stamatakis, 2014) under the GTR+gamma model, with 1,000 rapid bootstrap replicates. The resulting phylogeny indicates that the two individuals analyzed here belong to different *Tetrastigma* species; however, their precise species identification could not be confidently resolved (Figure S11).
