## Supplementary material for "Extensive mitogenome divergence across the Rafflesiaceae in size and impact of horizontal gene transfer": SupplFigures.pdf

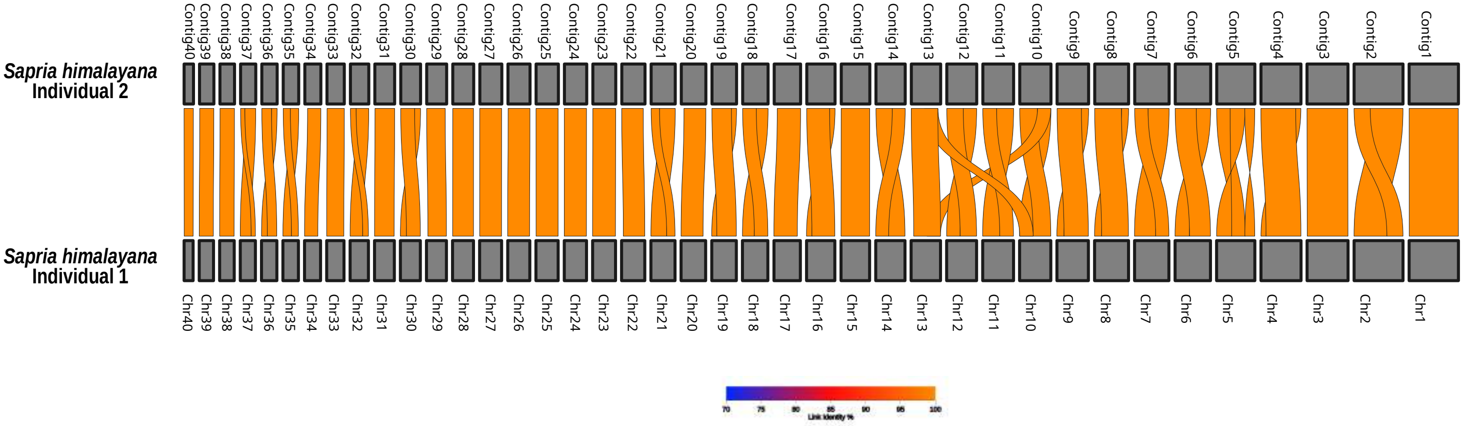


**Figure S1. Collinearity among mitochondrial sequences from two individuals of *Sapria himalayana*.** Gray blocks represent mitochondrial chromosomes, with block width proportional to sequence length. Internal colored lines denote homologous regions (>1 kb, >70% identity), with color intensity reflecting sequence identity (70–100%). Alignments were generated with LASTZ v1.0.4 and visualized using AliTV v1.0.6. Both individuals were collected in Yunnan, China.

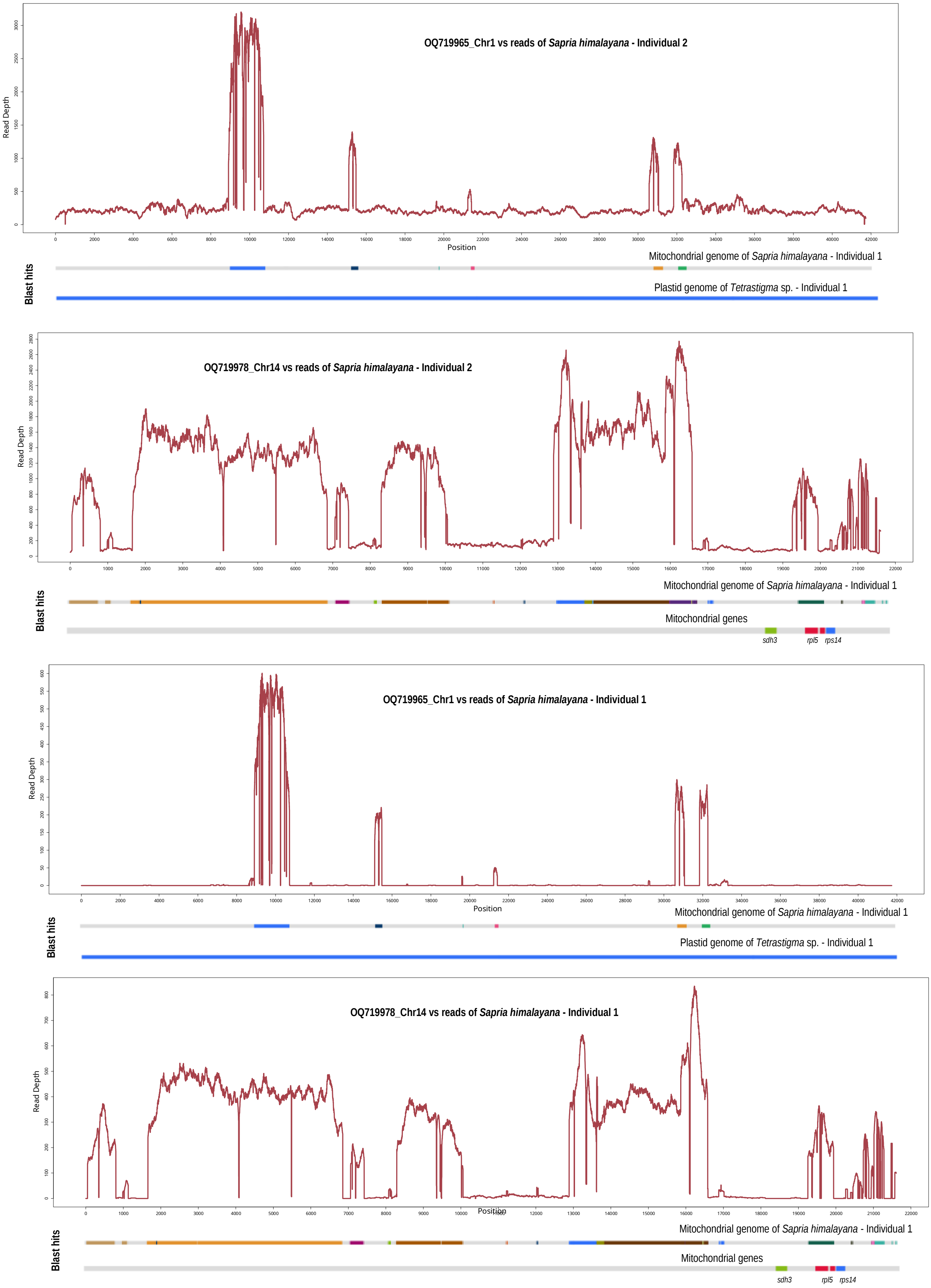


**Figure S2. Read mapping profiles revealing structural misassemblies in previously published *Sapria himalayana* mitogenomic contigs.** (A–D) Sequencing coverage (read depth) across two mitochondrial contigs originally assembled by Guo et al. (2023) (GenBank accessions OQ719965 [Chr1] and OQ719978 [Chr14]). Raw paired-end reads from two *Sapria* individuals (Individual 1 and Individual 2) were mapped to these reference sequences. Coverage profiles are shown for OQ719965 using reads from Individual 2 (A) and Individual 1 (C), and for OQ719978 using reads from Individual 2 (B) and Individual 1 (D). Read depth was calculated using Bowtie2 v2.4.4 with the parameters --end-to-end --very-sensitive --no-contain --no-discordant --no- mixed. Below each coordinate axis, homologous regions are arbitrarily colored based on BLASTn alignments against the assembled *Sapria* mitogenome (Individual 1), selected mitochondrial coding genes (*sdh3, rpl5, rps14*), and the plastome of *Tetrastigma* sp (Individual 1).

**Figure S3. Mitochondrial genomes of Rafflesiaceae species.** (A) Complete mitochondrial genome (mtDNA) of *Rhizanthes lowii*. The mtDNA is 282,577 bp in length and is partitioned into 35 circular chromosomes (Chr) of varying sizes. (B) Draft mitochondrial genome of *Rafflesia tuan-mudae*. (C) Draft mitochondrial genome of *Rafflesia arnoldii*. (D) Draft mitochondrial genome of *Rafflesia leonardi*. Concentric tracks depict genomic and structural features, including intact protein- coding genes, ribosomal RNAs (rRNAs), and transfer RNAs (tRNAs); pseudogenes >100 bp; repetitive elements >1,000 bp; and plastid-derived mitochondrial sequences (MTPTs) >100 bp.

**A)** *Rhizanthes lowii*

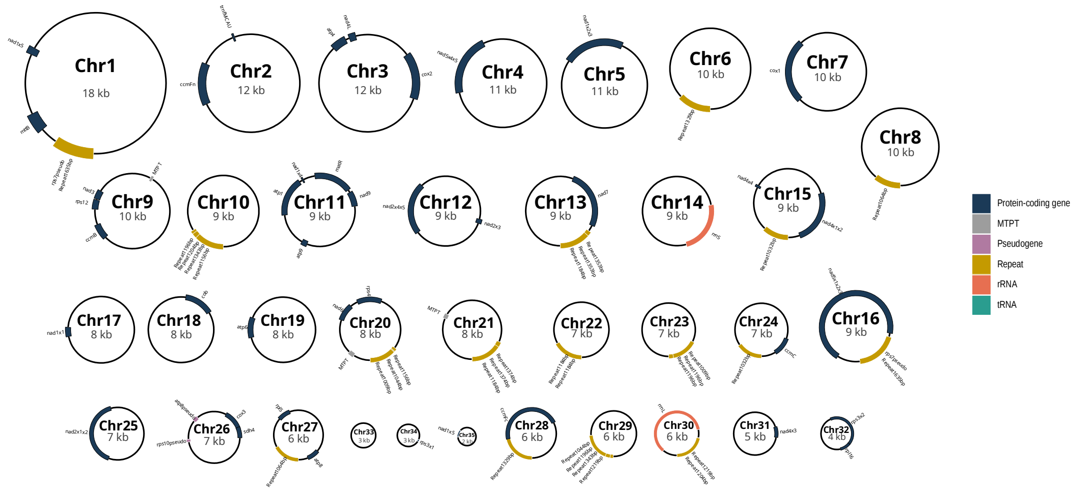

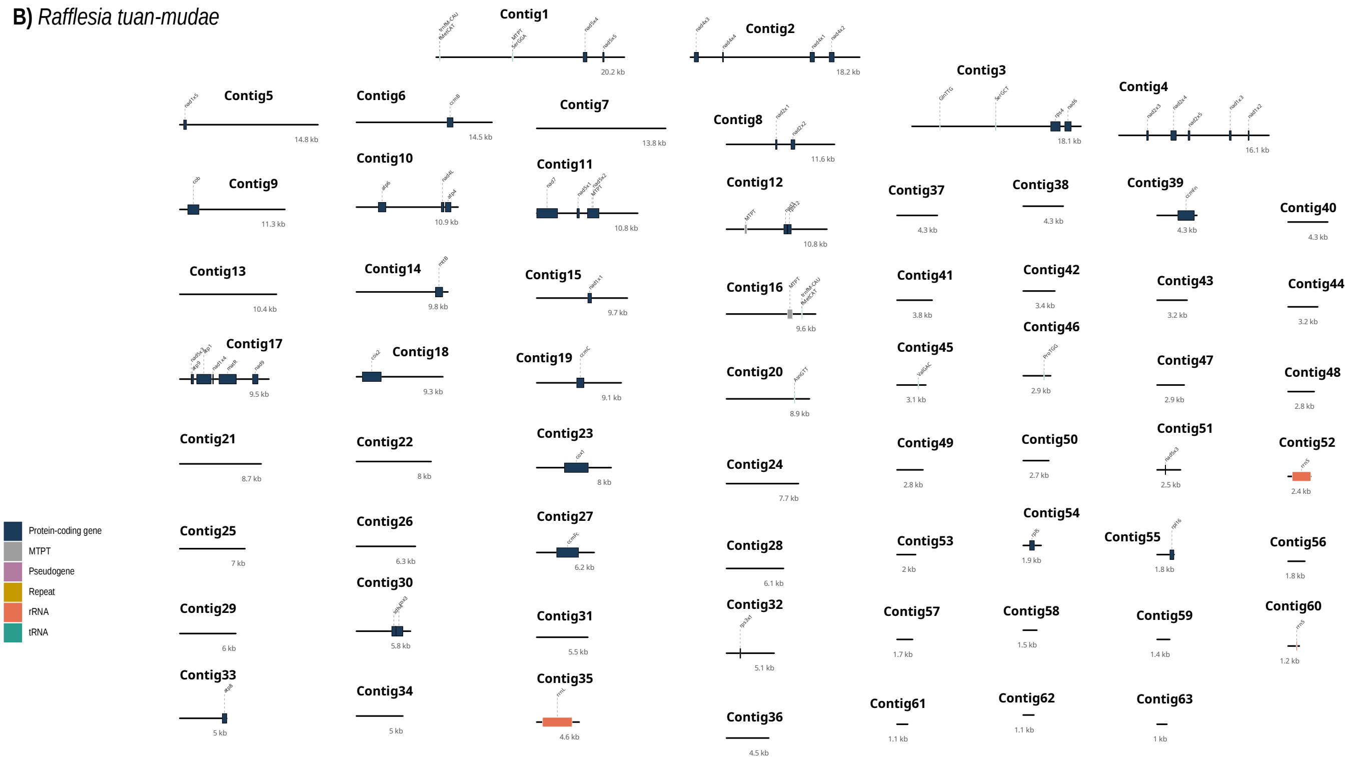

B) Rafflesia tuan-mudae
Contig1
20.2 kb
Contig2
18.2 kb
Contig3
18.1 kb
Contig4
16.1 kb
Contig5
14.8 kb
Contig6
14.5 kb
Contig7
13.8 kb
Contig8
11.6 kb
Contig9
11.3 kb
Contig10
10.9 kb
Contig11
10.8 kb
Contig12
10.8 kb
Contig13
10.4 kb
Contig14
9.8 kb
Contig15
9.7 kb
Contig16
9.6 kb
Contig17
9.5 kb
Contig18
9.3 kb
Contig19
9.1 kb
Contig20
8.9 kb
Contig21
8.7 kb
Contig22
8 kb
Contig23
8 kb
Contig24
7.7 kb
Contig25
7 kb
Contig26
6.3 kb
Contig27
6.2 kb
Contig28
6.1 kb
Contig29
6 kb
Contig30
5.8 kb
Contig31
5.5 kb
Contig32
5.1 kb
Contig33
5 kb
Contig34
5 kb
Contig35
4.6 kb
Contig36
4.5 kb
Contig37
4.3 kb
Contig38
4.3 kb
Contig39
4.3 kb
Contig40
4.3 kb
Contig41
3.8 kb
Contig42
3.4 kb
Contig43
3.2 kb
Contig44
3.2 kb
Contig45
3.1 kb
Contig46
2.9 kb
Contig47
2.9 kb
Contig48
2.8 kb
Contig49
2.8 kb
Contig50
2.7 kb
Contig51
2.5 kb
Contig52
2.4 kb
Contig53
2 kb
Contig54
1.9 kb
Contig55
1.8 kb
Contig56
1.8 kb
Contig57
1.7 kb
Contig58
1.5 kb
Contig59
1.4 kb
Contig60
1.2 kb
Contig61
1.1 kb
Contig62
1.1 kb
Contig63
1 kb
Protein-coding gene
MTPT
Pseudogene
Repeat
rRNA
tRNA

C) *Rafflesia arnoldii*

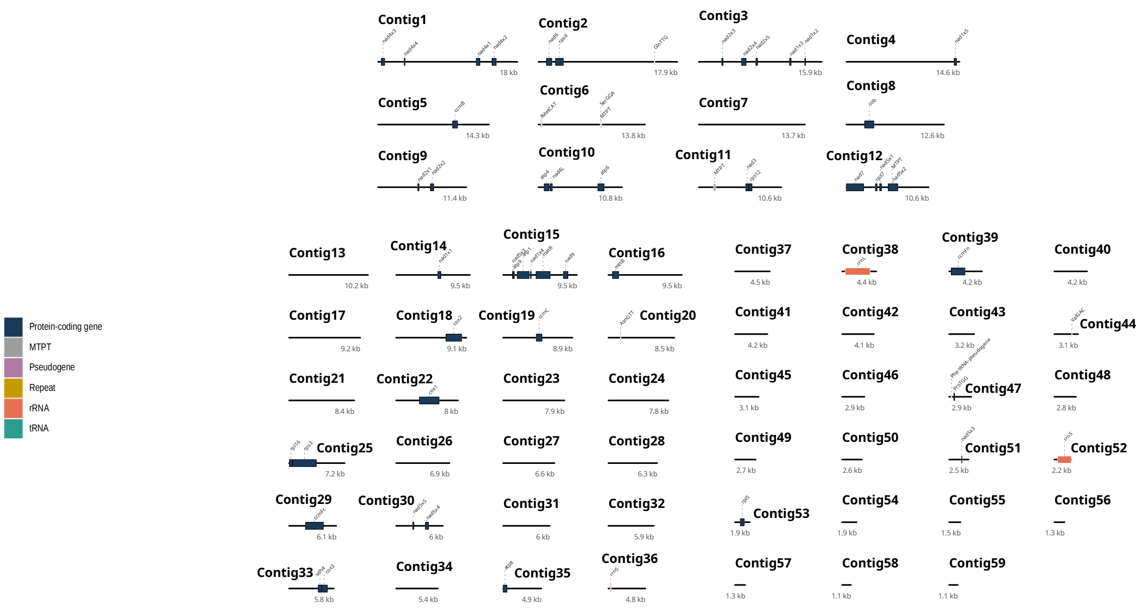

**D)** *Rafflesia leonardi*

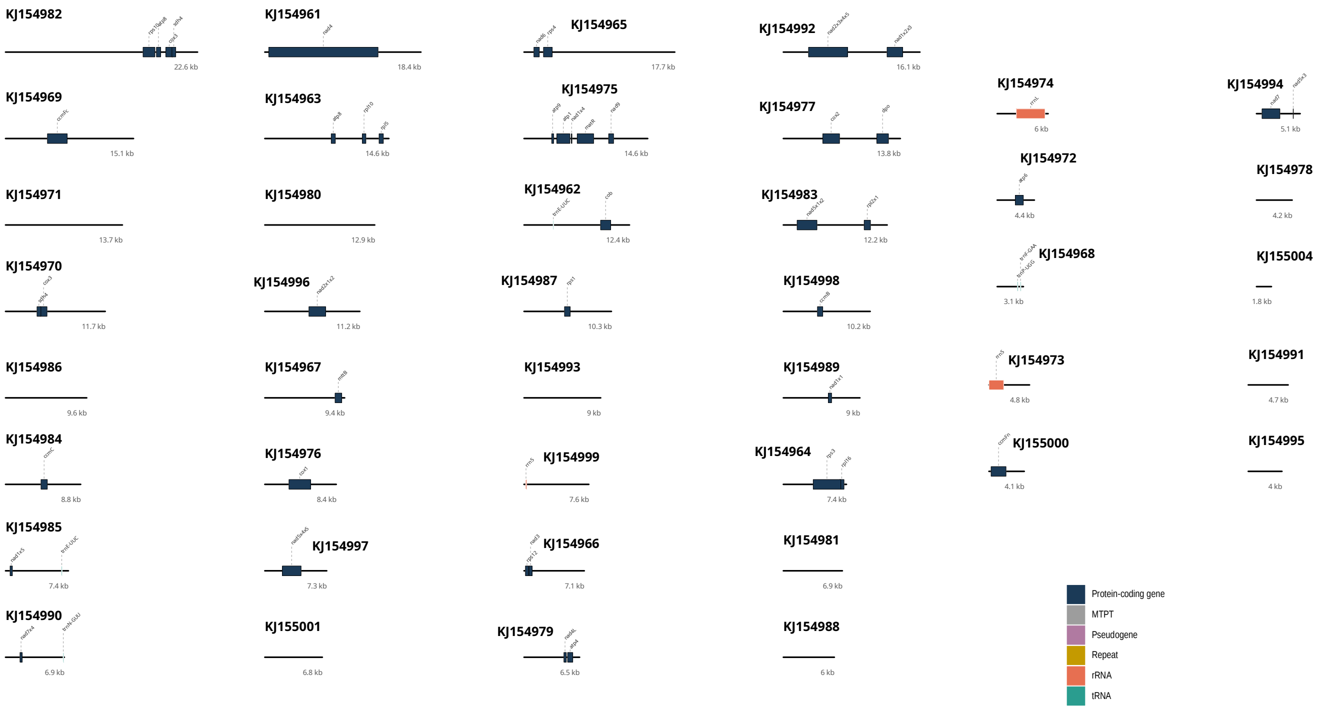

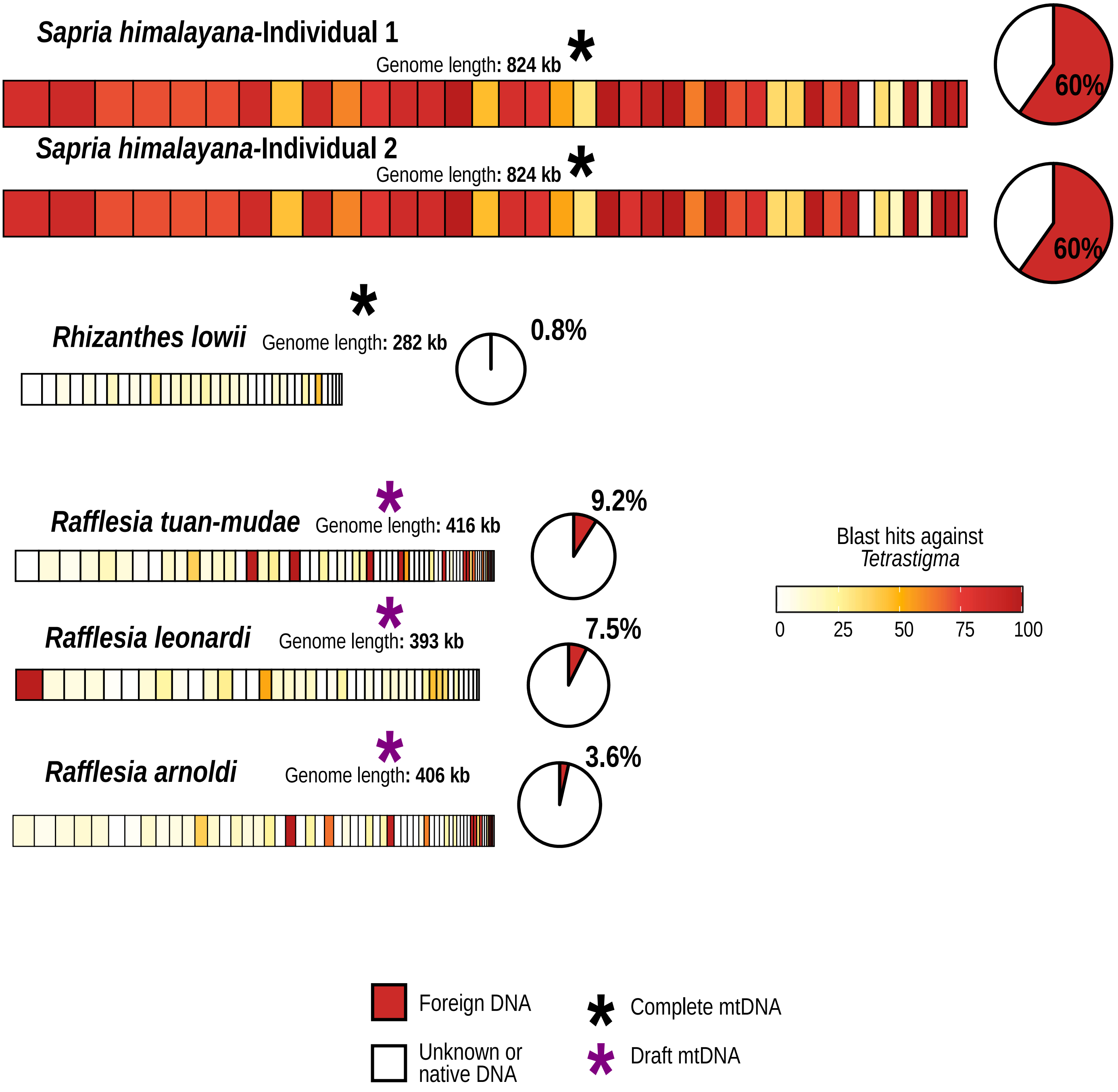


**Figure S4. Distribution of host-derived mitochondrial DNA across Rafflesiaceae mitogenomes.** Linear representations of mitochondrial genomes from *Sapria himalayana* (two individuals), *Rhizanthes lowi*i, and three *Rafflesia* species, with lengths scaled proportionally to their size. Each rectangle represents a mitochondrial chromosome or contig. Contigs are colored based on the proportion of BLASTn hits (>200 bp and 90% identity) to the mitochondrial data from host genus *Tetrastigma* (Tables S3-S7). White segments correspond to mitochondrial DNA of native or unknown origin. Pie charts summarize the total proportion of foreign DNA detected in each mitogenome. This proportion was calculated by retaining BLASTn hits to *Tetrastigma* mitochondrial DNA longer than 200 bp and with >90% sequence identity, considering only the best hit per region relative to other taxonomic groups. Redundant and overlapping matches were excluded.

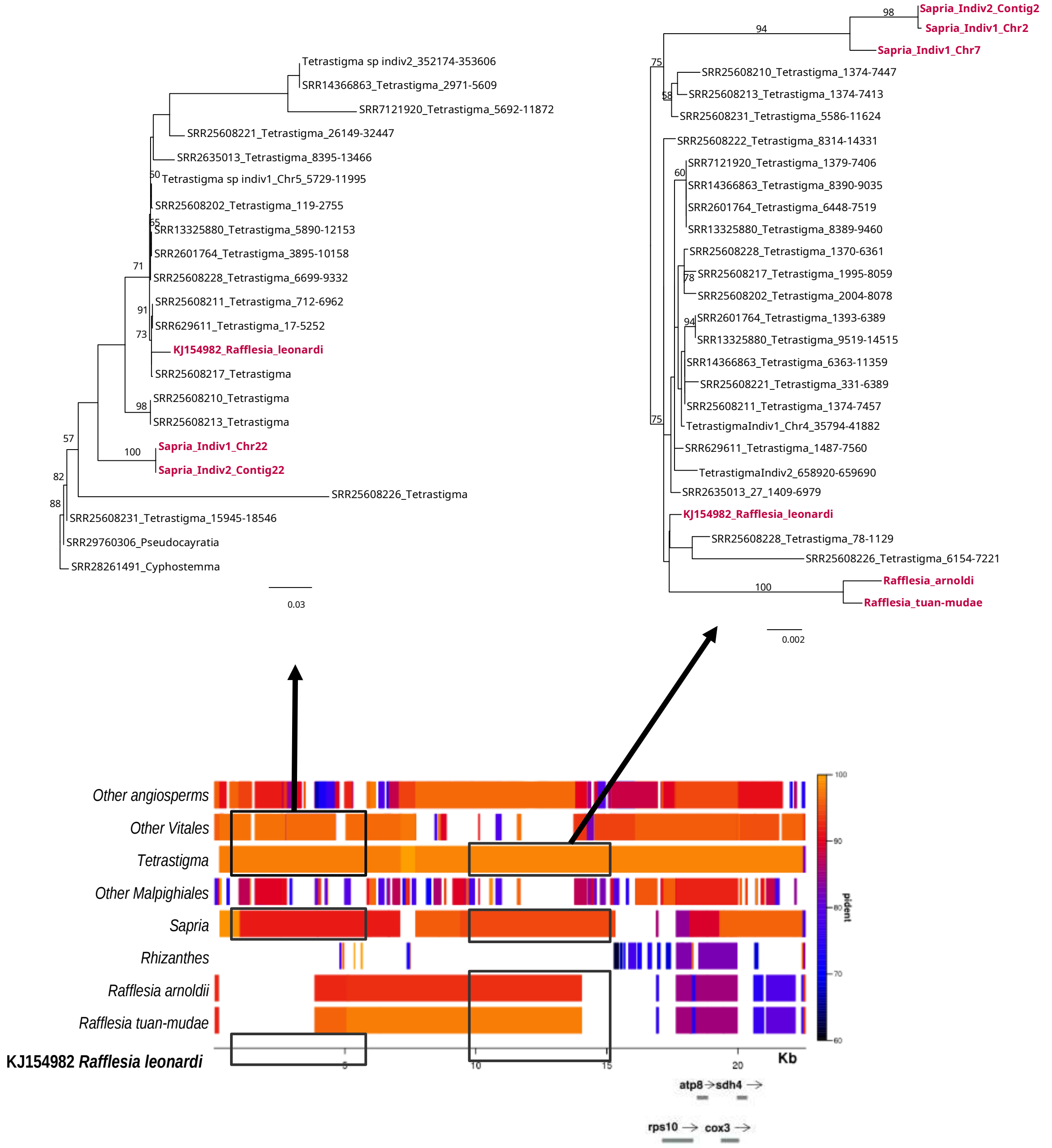


**Figure S5. Phylogenetic and sequence similarity evidence of convergent mitochondrial horizontal gene transfer in Rafflesiaceae**. (Top) Maximum-likelihood (ML) phylogenetic reconstructions of two intergenic regions. Branch support values are based on 1,000 rapid bootstrap replicates. Rafflesiaceae taxa are highlighted in magenta. (Bottom) Sequence similarity profile generated with the R package Sushi, showing BLASTn alignments against multiple taxonomic groups. A *Rafflesia leonardi* mitochondrial contig (GenBank accession KJ154982) was used as the reference sequence. The color gradient represents percentage nucleotide identity (pident), ranging from 60% to 100%. Black rectangles and solid arrows indicate the genomic regions analyzed in the phylogenetic reconstructions shown above. Nearby coding sequences are annotated along the coordinate axis.

**Figure S6. Phylogenetic analyses of mitochondrial genes of Rafflesiaceae.** Maximum likelihood analyses were performed with RAxML. ML bootstrap values >50% are shown. The scale bar corresponds to substitutions per site. For species for which only a subset of mitochondrial genes is available, corresponding NCBI accession numbers are provided. Sequences from *Rafflesia cantleyi*, *R. tuan-mudae*, and *Sapria himalayana* with sequence identifiers starting with RC, RT, and SH, respectively, were extracted from Xi *et al.* 2013 supplementary information.

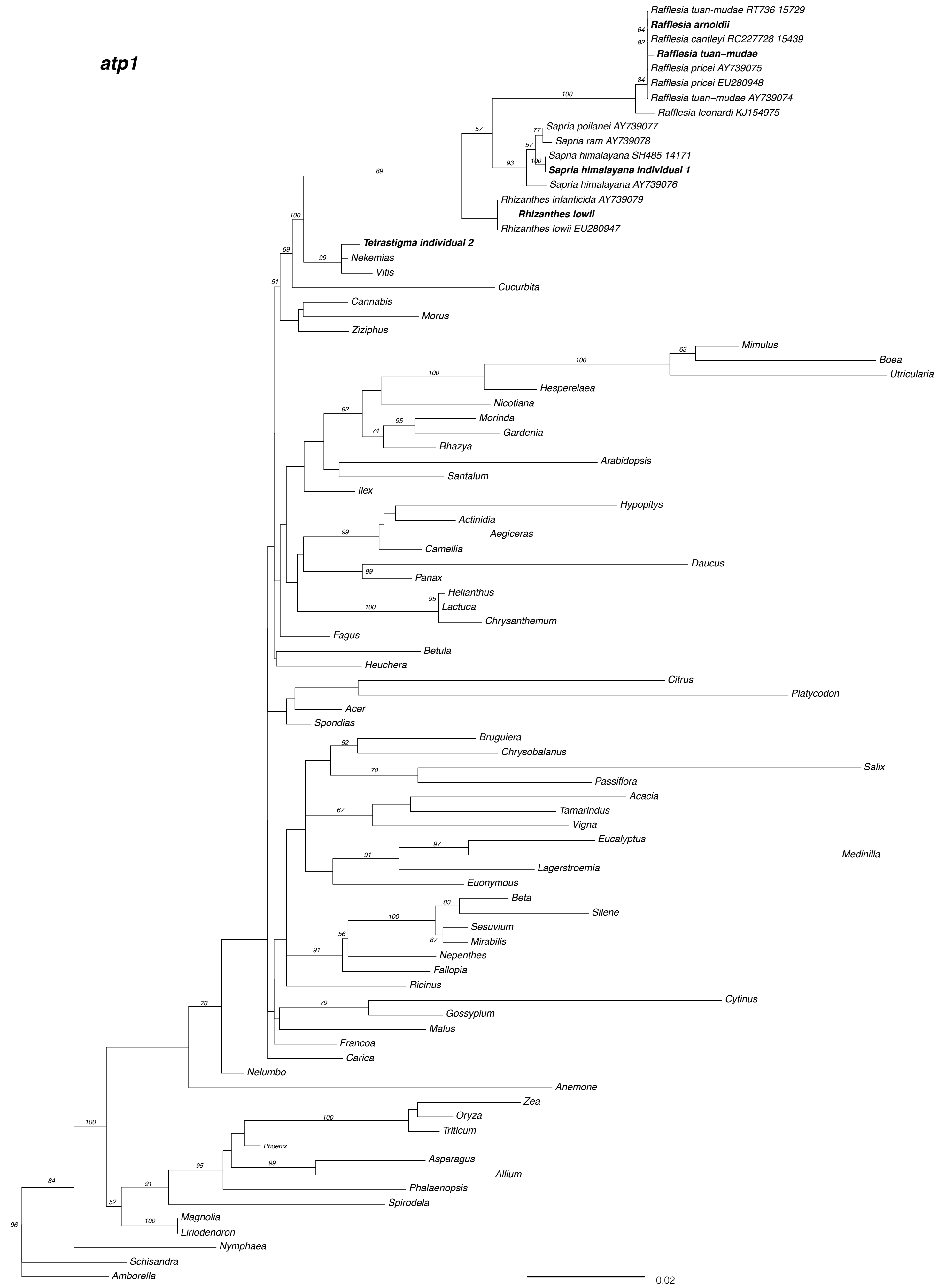

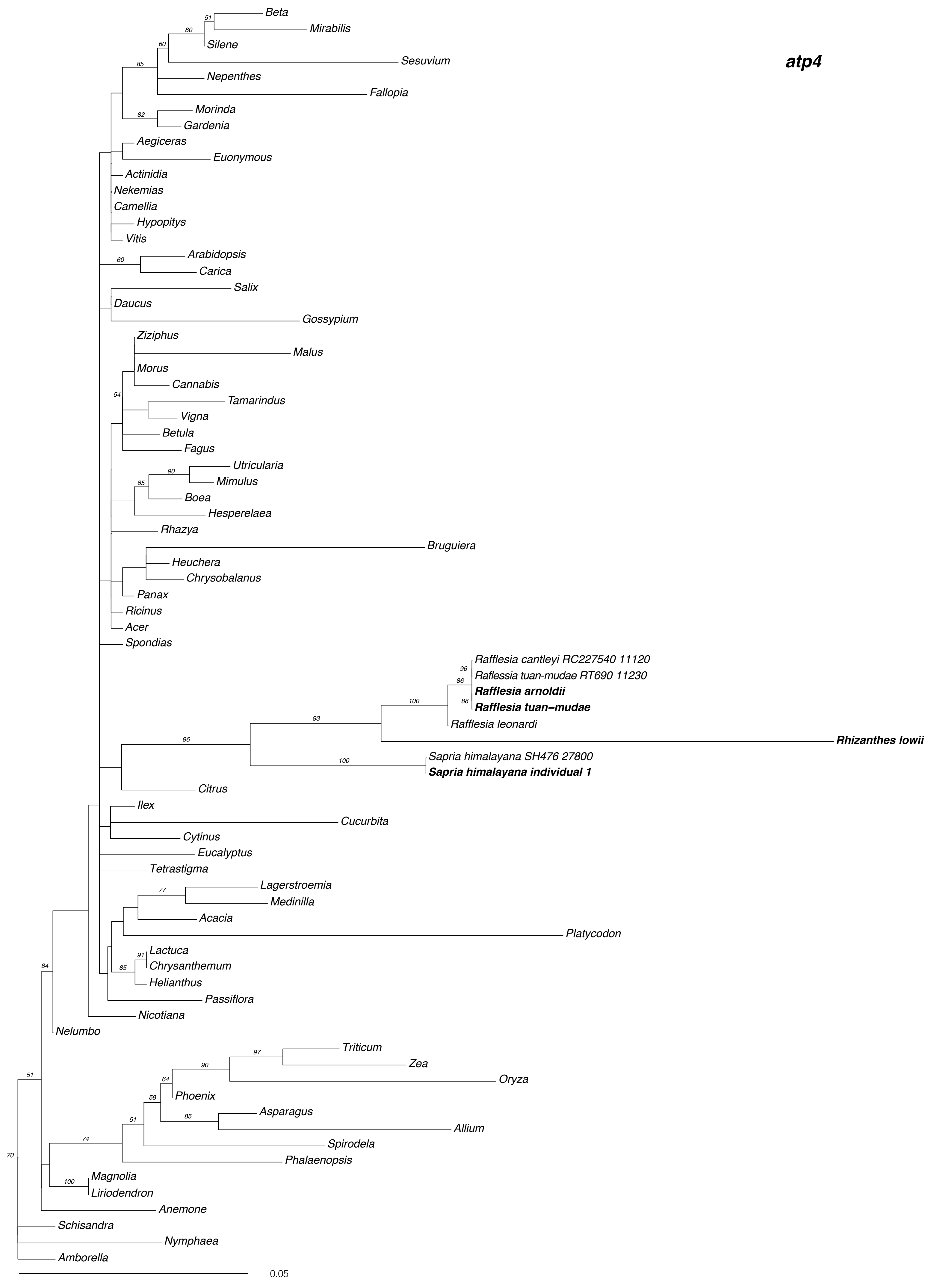
atp4
Beta
Mirabilis
Silene
Sesuvium
Nepenthes
Fallopia
Morinda
Gardenia
Aegiceras
Euonymous
Actinidia
Nekemias
Camellia
Hypopitys
Vitis
Arabidopsis
Carica
Salix
Daucus
Gossypium
Ziziphus
Malus
Morus
Cannabis
Tamarindus
Vigna
Betula
Fagus
Utricularia
Mimulus
Boea
Hesperelaea
Rhazya
Bruguiera
Heuchera
Chrysobalanus
Panax
Ricinus
Acer
Spondias
Rafflesia cantleyi RC227540 11120
Raflessia tuan-mudae RT690 11230
Rafflesia arnoldii
Rafflesia tuan–mudae
Rafflesia leonardi
Rhizanthes lowii
Sapria himalayana SH476 27800
Sapria himalayana individual 1
Citrus
Ilex
Cucurbita
Cytinus
Eucalyptus
Tetrastigma
Lagerstroemia
Medinilla
Acacia
Platycodon
Lactuca
Chrysanthemum
Helianthus
Passiflora
Nicotiana
Nelumbo
Triticum
Zea
Oryza
Phoenix
Asparagus
Allium
Spirodela
Phalaenopsis
Magnolia
Liriodendron
Anemone
Schisandra
Nymphaea
Amborella
0.05

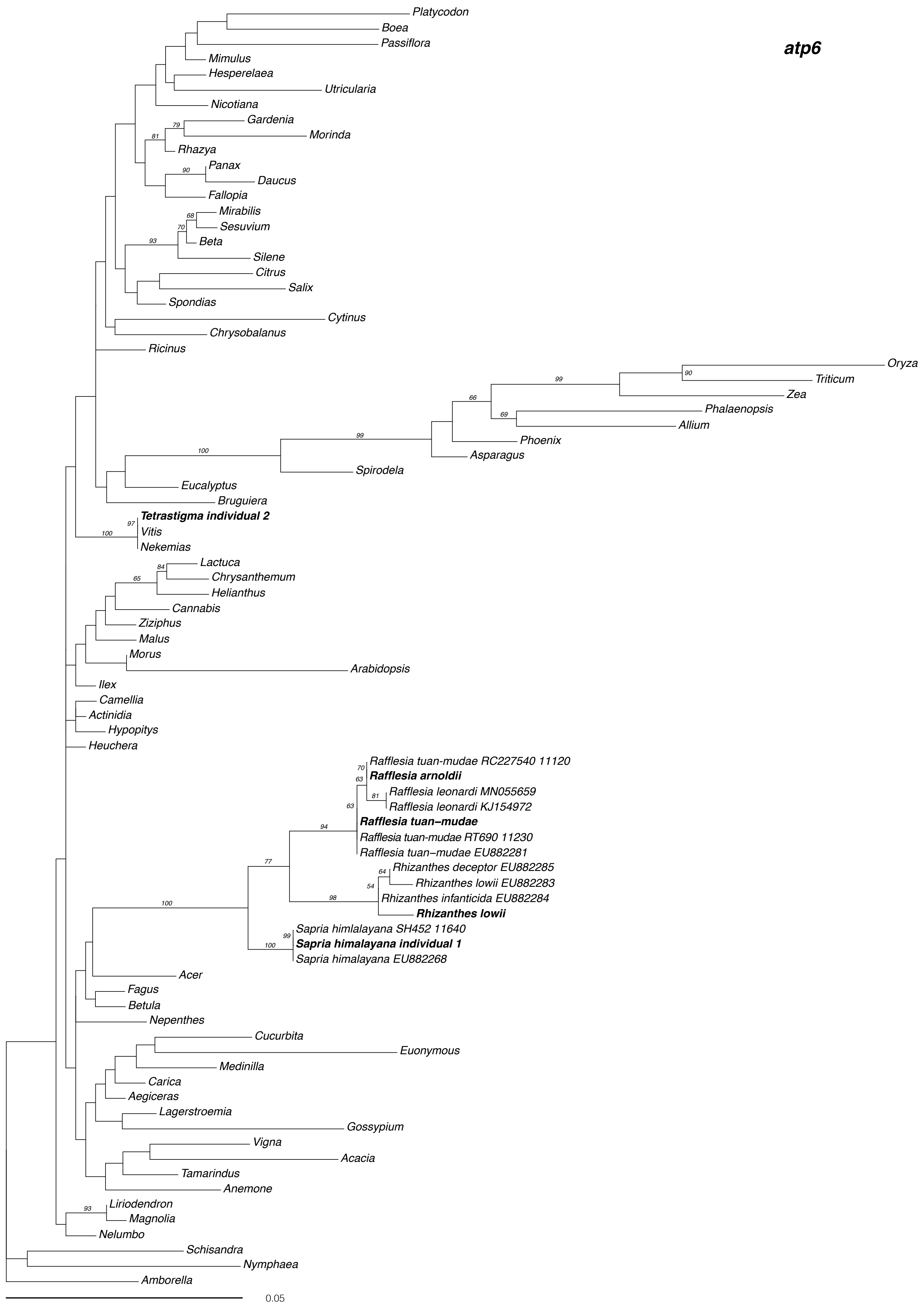
atp6
Platycodon
Boea
Passiflora
Mimulus
Hesperelaea
Utricularia
Nicotiana
Gardenia
Morinda
Rhazya
Panax
Daucus
Fallopia
Mirabilis
Sesuvium
Beta
Silene
Citrus
Salix
Spondias
Cytinus
Chrysobalanus
Ricinus
Oryza
Triticum
Zea
Phalaenopsis
Allium
Phoenix
Asparagus
Spirodela
Eucalyptus
Bruguiera
Tetrastigma individual 2
Vitis
Nekemias
Lactuca
Chrysanthemum
Helianthus
Cannabis
Ziziphus
Malus
Morus
Arabidopsis
Ilex
Camellia
Actinidia
Hypopitys
Heuchera
Rafflesia tuan-mudae RC227540 11120
Rafflesia arnoldii
Rafflesia leonardi MN055659
Rafflesia leonardi KJ154972
Rafflesia tuan–mudae
Rafflesia tuan-mudae RT690 11230
Rafflesia tuan–mudae EU882281
Rhizanthes deceptor EU882285
Rhizanthes lowii EU882283
Rhizanthes infanticida EU882284
Rhizanthes lowii
Sapria himlalayana SH452 11640
Sapria himalayana individual 1
Sapria himalayana EU882268
Acer
Fagus
Betula
Nepenthes
Cucurbita
Euonymous
Medinilla
Carica
Aegiceras
Lagerstroemia
Gossypium
Vigna
Acacia
Tamarindus
Anemone
Liriodendron
Magnolia
Nelumbo
Schisandra
Nymphaea
Amborella
0.05

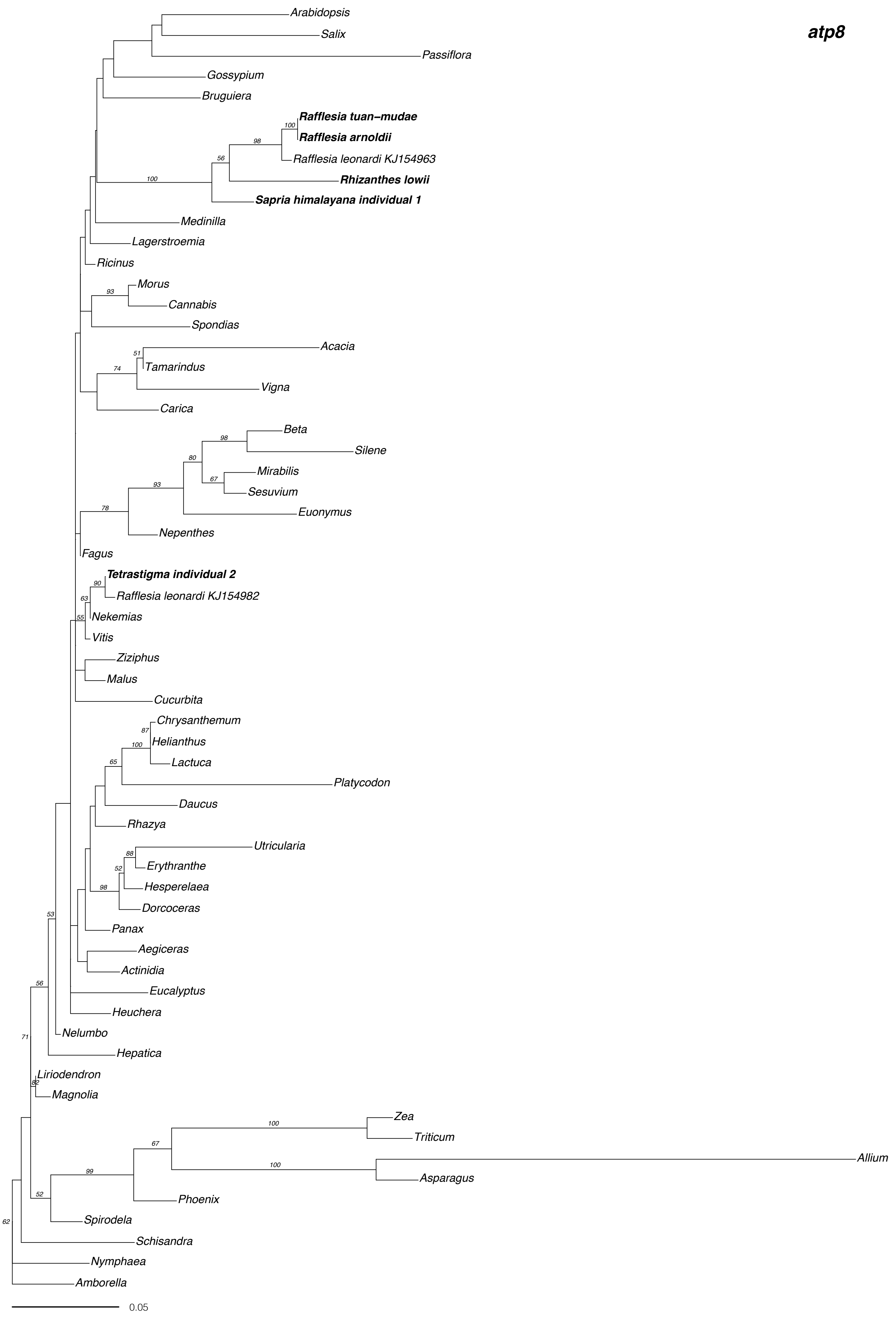
atp8
Arabidopsis
Salix
Passiflora
Gossypium
Bruguiera
Rafflesia tuan-mudae
Rafflesia arnoldii
Rafflesia leonardi KJ154963
Rhizanthes lowii
Sapria himalayana individual 1
Medinilla
Lagerstroemia
Ricinus
Morus
Cannabis
Spondias
Acacia
Tamarindus
Vigna
Carica
Beta
Silene
Mirabilis
Sesuvium
Euonymus
Nepenthes
Fagus
Tetrastigma individual 2
Rafflesia leonardi KJ154982
Nekemias
Vitis
Ziziphus
Malus
Cucurbita
Chrysanthemum
Helianthus
Lactuca
Platycodon
Daucus
Rhazya
Utricularia
Erythranthe
Hesperelaea
Dorcoceras
Panax
Aegiceras
Actinidia
Eucalyptus
Heuchera
Nelumbo
Hepatica
Liriodendron
Magnolia
Zea
Triticum
Allium
Asparagus
Phoenix
Spirodela
Schisandra
Nymphaea
Amborella
0.05

### atp9

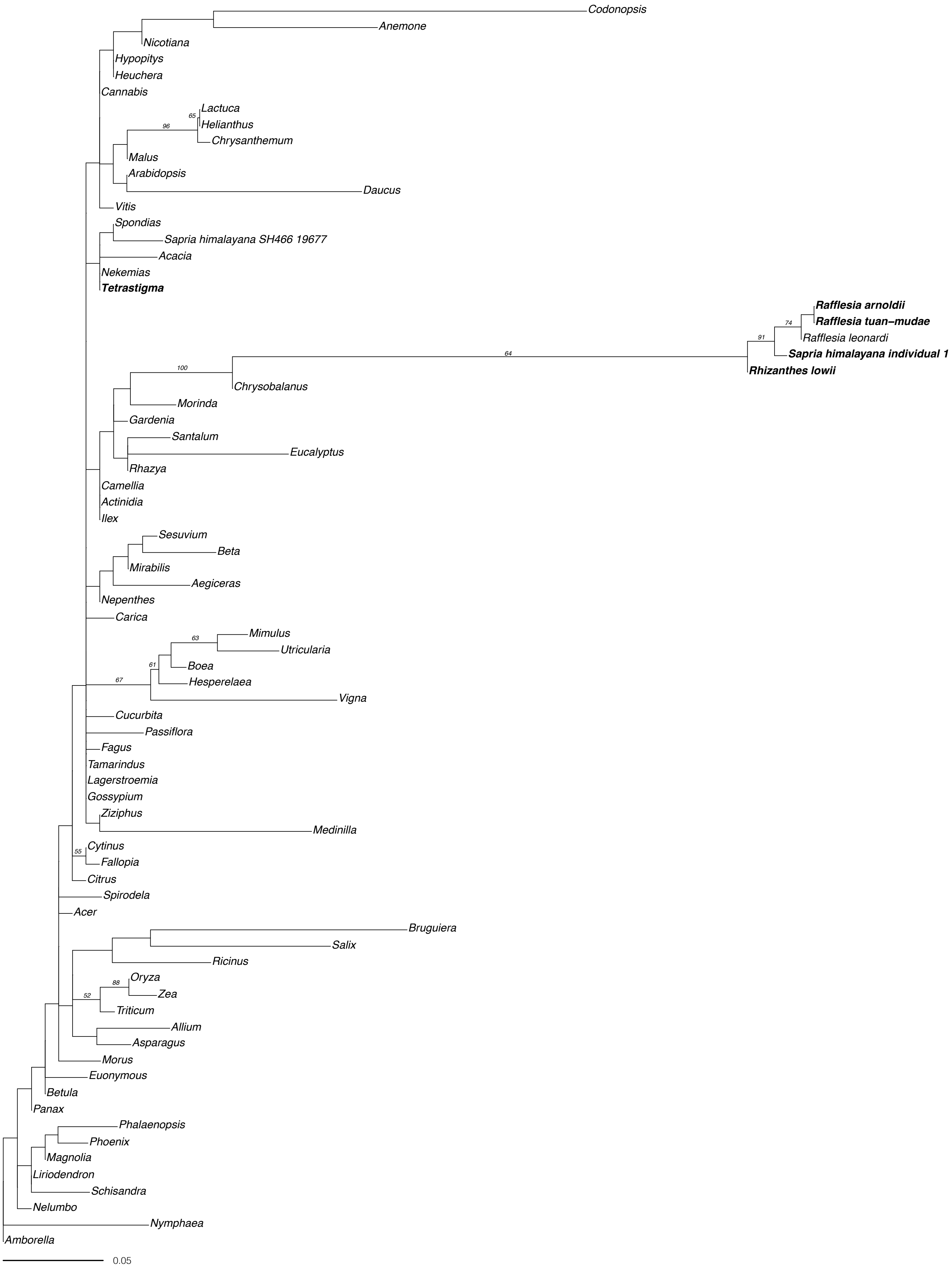
Codonopsis
Anemone
Nicotiana
Hypopitys
Heuchera
Cannabis
65
Lactuca
96
Helianthus
Chrysanthemum
Malus
Arabidopsis
Daucus
Vitis
Spondias
Sapria himalayana SH466 19677
Acacia
Nekemias
Tetrastigma
Rafflesia arnoldii
74
Rafflesia tuan-mudae
91
Rafflesia leonardi
64
Sapria himalayana individual 1
100
Rhizanthes lowii
Chrysobalanus
Morinda
Gardenia
Santalum
Eucalyptus
Rhazya
Camellia
Actinidia
Ilex
Sesuvium
Beta
Mirabilis
Aegiceras
Nepenthes
Carica
Mimulus
63
Utricularia
61
Boea
67
Hesperelaea
Vigna
Cucurbita
Passiflora
Fagus
Tamarindus
Lagerstroemia
Gossypium
Ziziphus
Medinilla
55
Cytinus
Fallopia
Citrus
Spirodela
Acer
Bruguiera
Salix
Ricinus
88
Oryza
52
Zea
Triticum
Allium
Asparagus
Morus
Euonymous
Betula
Panax
Phalaenopsis
Phoenix
Magnolia
Liriodendron
Schisandra
Nelumbo
Nymphaea
Amborella
0.05

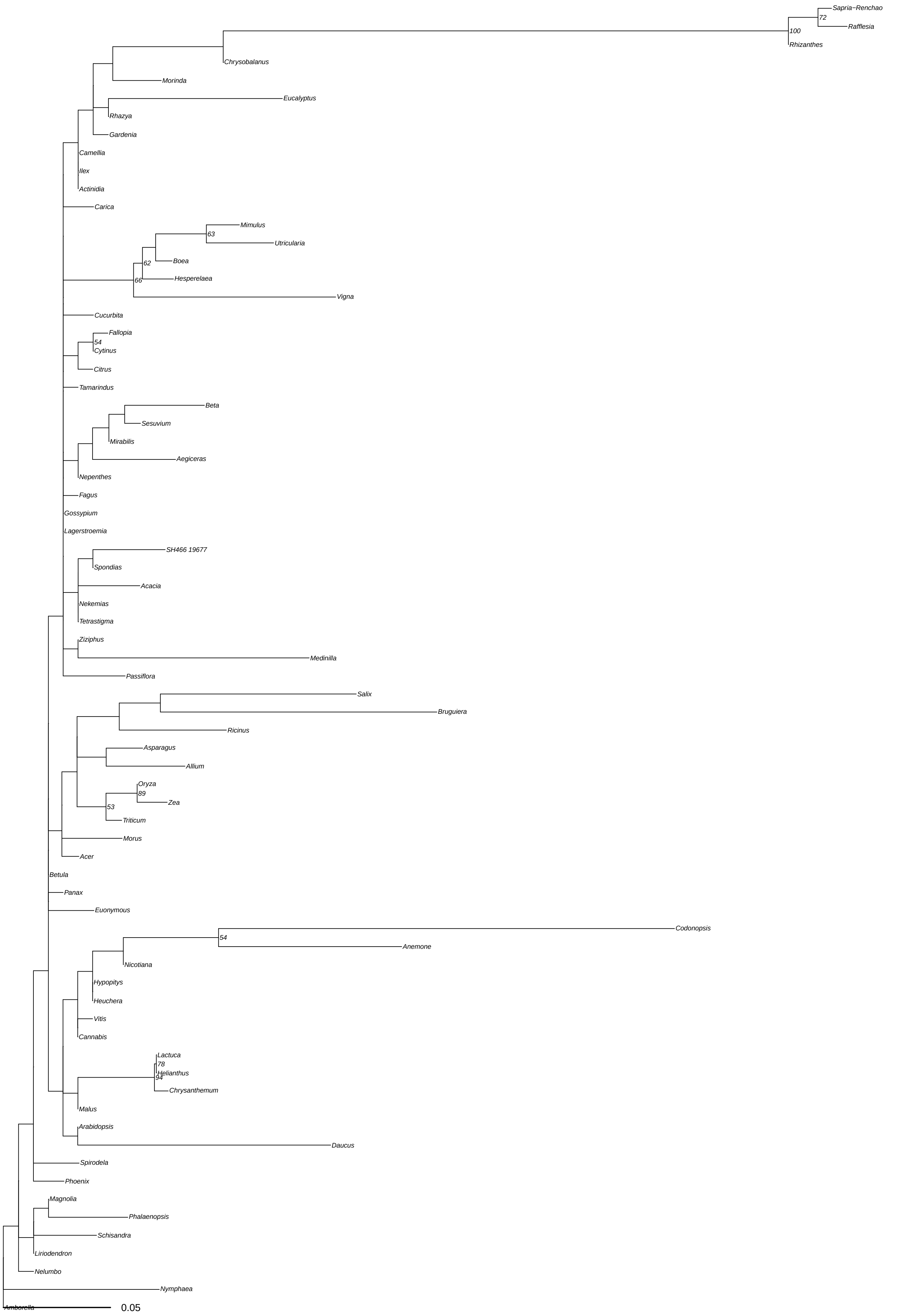
Sapria−Renchao
72
Rafflesia
100
Rhizanthes
Chrysobalanus
Morinda
Eucalyptus
Rhazya
Gardenia
Camellia
Ilex
Actinidia
Carica
Mimulus
63
Utricularia
Boea
62
Hesperelaea
66
Vigna
Cucurbita
Fallopia
54
Cytinus
Citrus
Tamarindus
Beta
Sesuvium
Mirabilis
Aegiceras
Nepenthes
Fagus
Gossypium
Lagerstroemia
SH466 19677
Spondias
Acacia
Nekemias
Tetrastigma
Ziziphus
Medinilla
Passiflora
Salix
Bruguiera
Ricinus
Asparagus
Allium
Oryza
89
Zea
53
Triticum
Morus
Acer
Betula
Panax
Euonymous
Codonopsis
54
Anemone
Nicotiana
Hypopitys
Heuchera
Vitis
Cannabis
Lactuca
78
Helianthus
94
Chrysanthemum
Malus
Arabidopsis
Daucus
Spirodela
Phoenix
Magnolia
Phalaenopsis
Schisandra
Liriodendron
Nelumbo
Nymphaea
Amborella
0.05

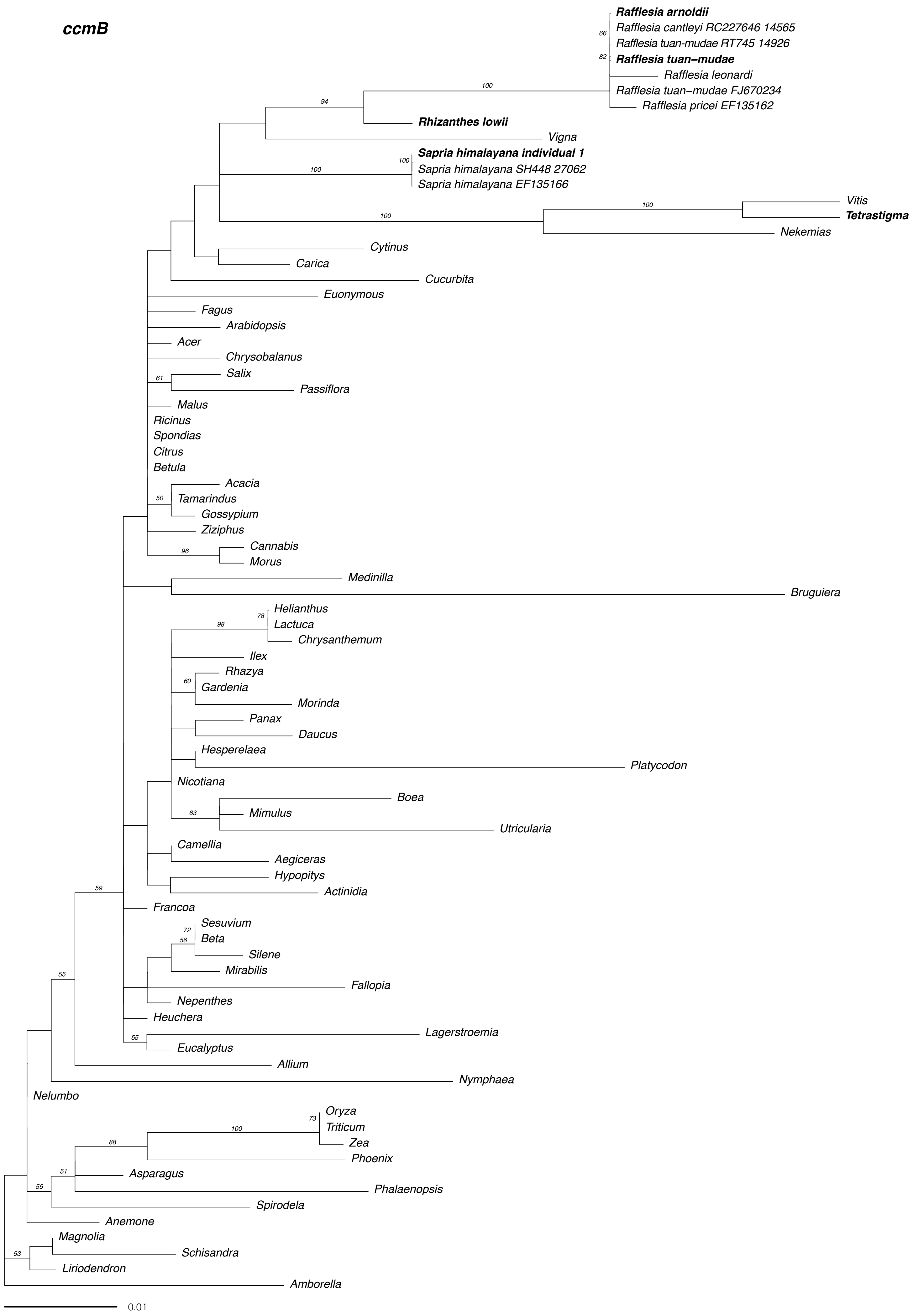
ccmB
Rafflesia arnoldii
Rafflesia cantleyi RC227646 14565
Rafflesia tuan-mudae RT745 14926
Rafflesia tuan–mudae
Rafflesia leonardi
Rafflesia tuan–mudae FJ670234
Rafflesia pricei EF135162
Rhizanthes lowii
Vigna
Sapria himalayana individual 1
Sapria himalayana SH448 27062
Sapria himalayana EF135166
Vitis
Tetrastigma
Nekemias
Cytinus
Carica
Cucurbita
Euonymous
Fagus
Arabidopsis
Acer
Chrysobalanus
Salix
Passiflora
Malus
Ricinus
Spondias
Citrus
Betula
Acacia
Tamarindus
Gossypium
Ziziphus
Cannabis
Morus
Medinilla
Bruguiera
Helianthus
Lactuca
Chrysanthemum
Ilex
Rhazya
Gardenia
Morinda
Panax
Daucus
Hesperelaea
Platycodon
Nicotiana
Boea
Mimulus
Utricularia
Camellia
Aegiceras
Hypopitys
Actinidia
Francoa
Sesuvium
Beta
Silene
Mirabilis
Fallopia
Nepenthes
Heuchera
Lagerstroemia
Eucalyptus
Allium
Nymphaea
Nelumbo
Oryza
Triticum
Zea
Phoenix
Asparagus
Phalaenopsis
Spirodela
Anemone
Magnolia
Schisandra
Liriodendron
Amborella
0.01

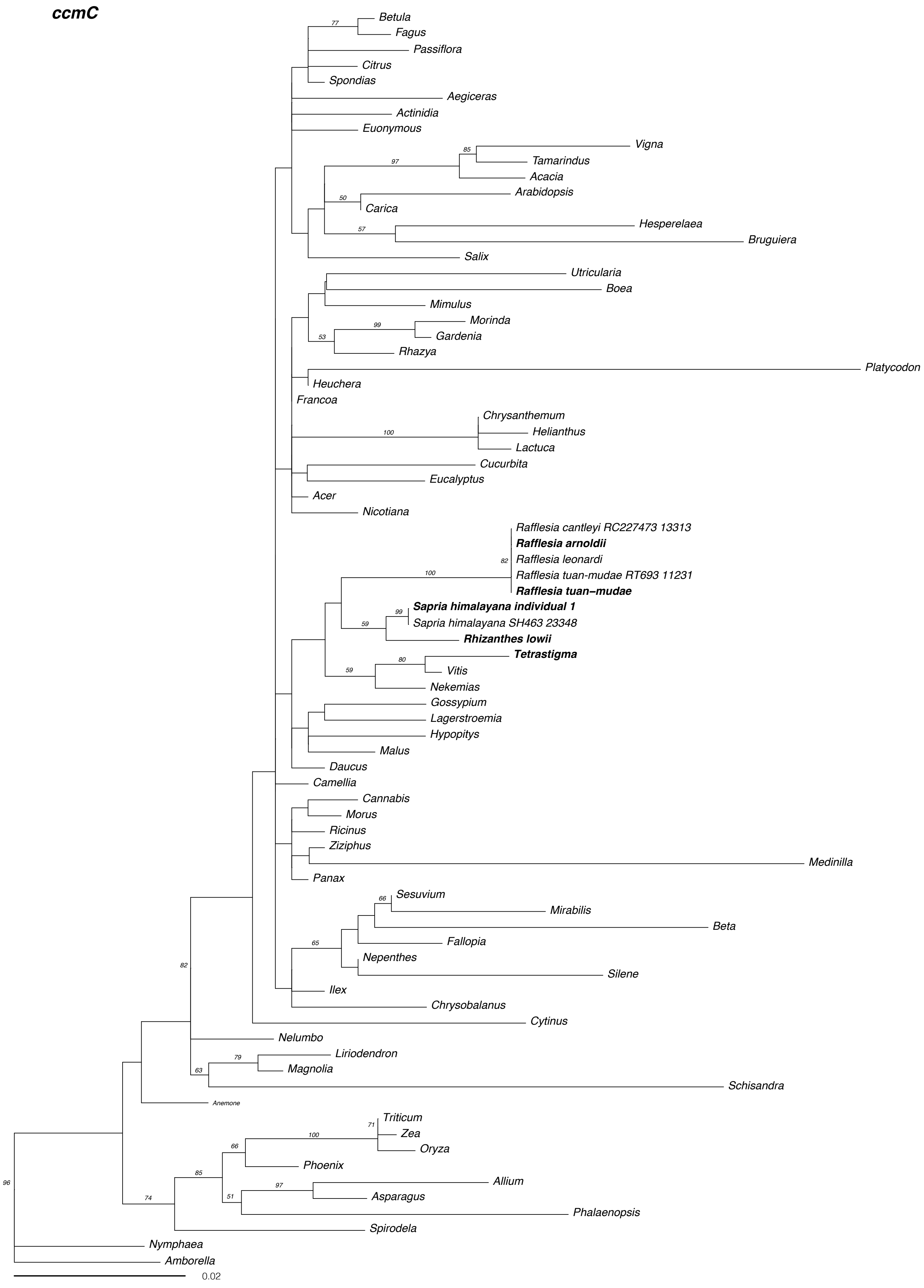
ccmC
Betula
Fagus
Passiflora
Citrus
Spondias
Aegiceras
Actinidia
Euonymous
Vigna
Tamarindus
Acacia
Arabidopsis
Carica
Hesperelaea
Bruguiera
Salix
Utricularia
Boea
Mimulus
Morinda
Gardenia
Rhazya
Platycodon
Heuchera
Francoa
Chrysanthemum
Helianthus
Lactuca
Cucurbita
Eucalyptus
Acer
Nicotiana
Rafflesia cantleyi RC227473 13313
Rafflesia arnoldii
Rafflesia leonardi
Rafflesia tuan-mudae RT693 11231
Rafflesia tuan–mudae
Sapria himalayana individual 1
Sapria himalayana SH463 23348
Rhizanthes lowii
Tetrastigma
Vitis
Nekemias
Gossypium
Lagerstroemia
Hypopitys
Malus
Daucus
Camellia
Cannabis
Morus
Ricinus
Ziziphus
Medinilla
Panax
Sesuvium
Mirabilis
Beta
Fallopia
Nepenthes
Silene
Ilex
Chrysobalanus
Cytinus
Nelumbo
Liriodendron
Magnolia
Schisandra
Anemone
Triticum
Zea
Oryza
Phoenix
Allium
Asparagus
Phalaenopsis
Spirodela
Nymphaea
Amborella
0.02

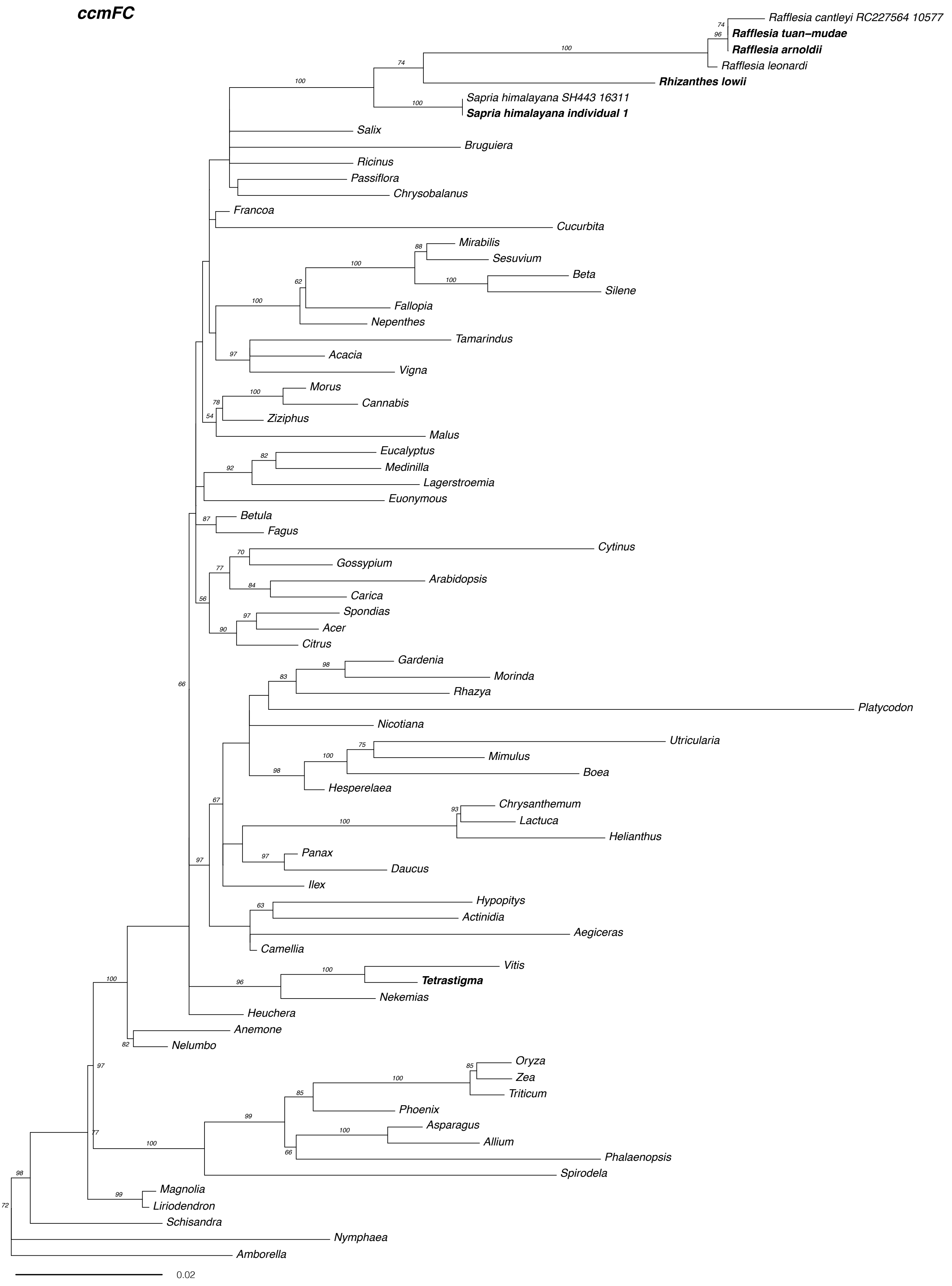
ccmFC
Rafflesia cantleyi RC227564 10577
Rafflesia tuan–mudae
Rafflesia arnoldii
Rafflesia leonardi
Rhizanthes lowii
Sapria himalayana SH443 16311
Sapria himalayana individual 1
Salix
Bruguiera
Ricinus
Passiflora
Chrysobalanus
Francoa
Cucurbita
Mirabilis
Sesuvium
Beta
Silene
Fallopia
Nepenthes
Tamarindus
Acacia
Vigna
Morus
Cannabis
Ziziphus
Malus
Eucalyptus
Medinilla
Lagerstroemia
Euonymous
Betula
Fagus
Cytinus
Gossypium
Arabidopsis
Carica
Spondias
Acer
Citrus
Gardenia
Morinda
Rhazya
Platycodon
Nicotiana
Utricularia
Mimulus
Boea
Hesperelaea
Chrysanthemum
Lactuca
Helianthus
Panax
Daucus
Ilex
Hypopitys
Actinidia
Aegiceras
Camellia
Vitis
Tetrastigma
Nekemias
Heuchera
Anemone
Nelumbo
Oryza
Zea
Triticum
Phoenix
Asparagus
Allium
Phalaenopsis
Spirodela
Magnolia
Liriodendron
Schisandra
Nymphaea
Amborella
0.02

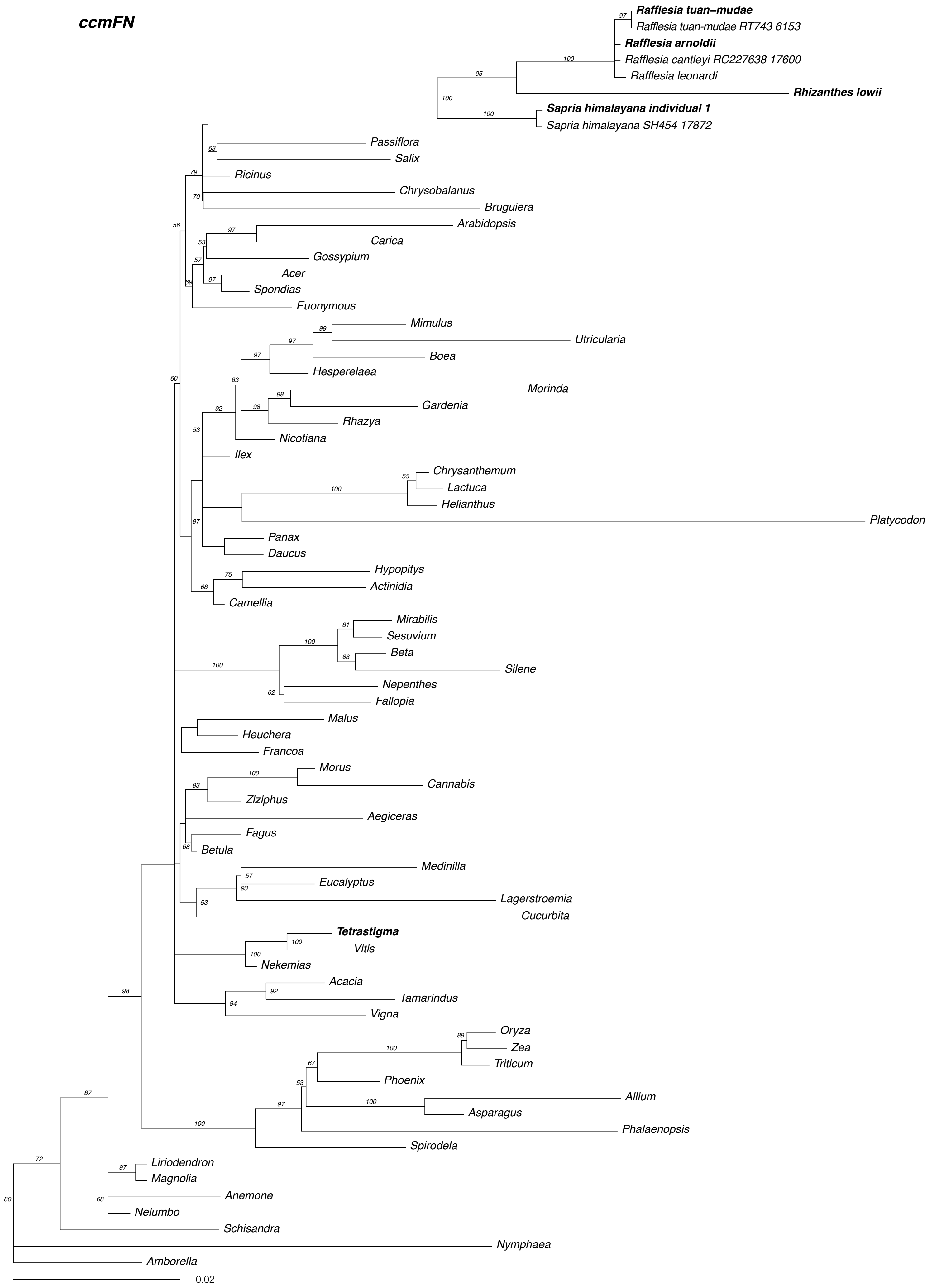
ccmFN
Rafflesia tuan–mudae
Rafflesia tuan-mudae RT743 6153
Rafflesia arnoldii
Rafflesia cantleyi RC227638 17600
Rafflesia leonardi
Rhizanthes lowii
Sapria himalayana individual 1
Sapria himalayana SH454 17872
Passiflora
Salix
Ricinus
Chrysobalanus
Bruguiera
Arabidopsis
Carica
Gossypium
Acer
Spondias
Euonymous
Mimulus
Utricularia
Boea
Hesperelaea
Morinda
Gardenia
Rhazya
Nicotiana
Ilex
Chrysanthemum
Lactuca
Helianthus
Platycodon
Panax
Daucus
Hypopitys
Actinidia
Camellia
Mirabilis
Sesuvium
Beta
Silene
Nepenthes
Fallopia
Malus
Heuchera
Francoa
Morus
Cannabis
Ziziphus
Aegiceras
Fagus
Betula
Medinilla
Eucalyptus
Lagerstroemia
Cucurbita
Tetrastigma
Vitis
Nekemias
Acacia
Tamarindus
Vigna
Oryza
Zea
Triticum
Phoenix
Allium
Asparagus
Phalaenopsis
Spirodela
Liriodendron
Magnolia
Anemone
Nelumbo
Schisandra
Nymphaea
Amborella
0.02

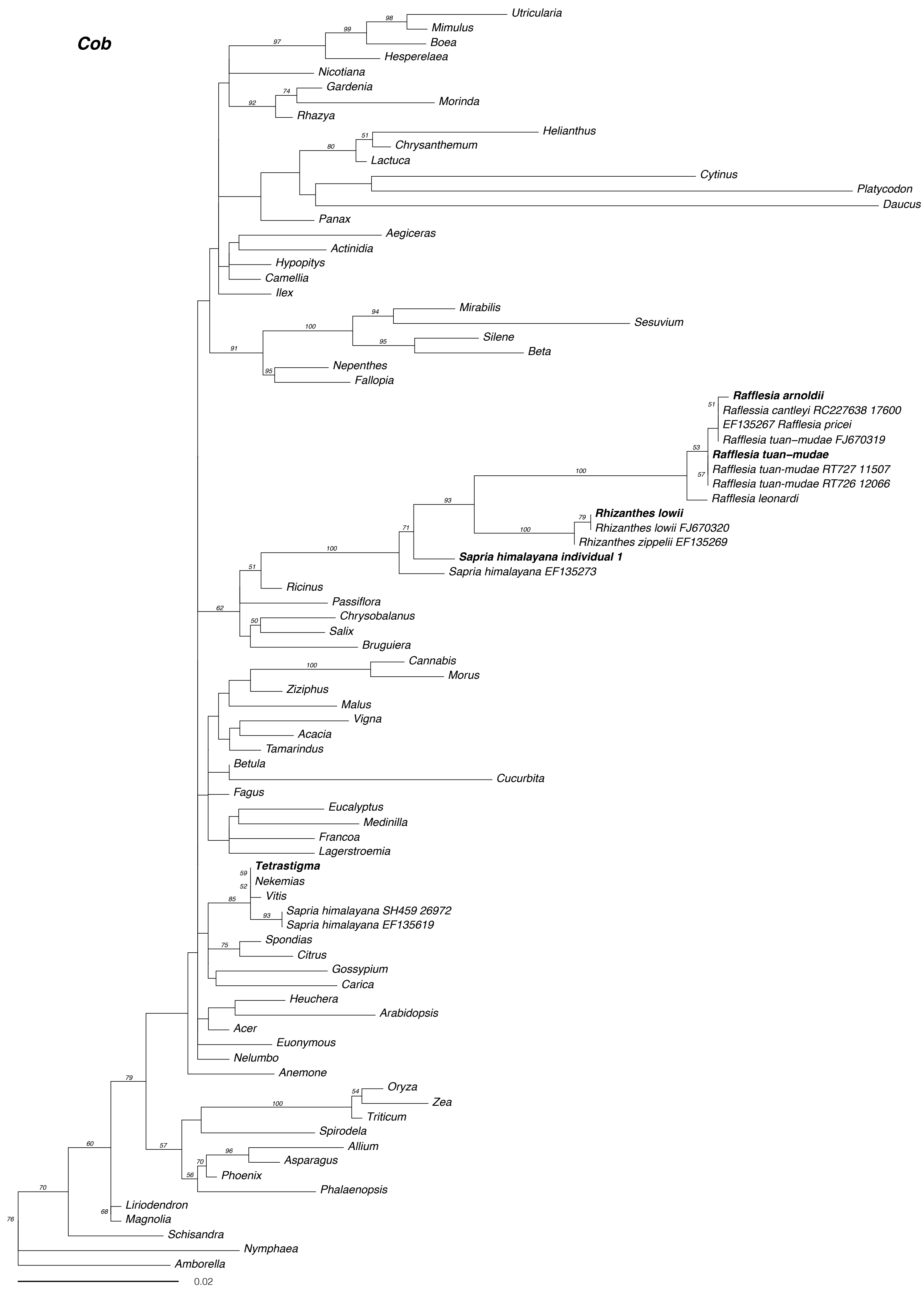
Cob
Utricularia
Mimulus
Boea
Hesperelaea
Nicotiana
Gardenia
Morinda
Rhazya
Helianthus
Chrysanthemum
Lactuca
Cytinus
Platycodon
Daucus
Panax
Aegiceras
Actinidia
Hypopitys
Camellia
Ilex
Mirabilis
Sesuvium
Silene
Beta
Nepenthes
Fallopia
Rafflesia arnoldii
Raflessia cantleyi RC227638 17600
EF135267 Rafflesia pricei
Rafflesia tuan–mudae FJ670319
Rafflesia tuan–mudae
Rafflesia tuan–mudae RT727 11507
Rafflesia tuan–mudae RT726 12066
Rafflesia leonardi
Rhizanthes lowii
Rhizanthes lowii FJ670320
Rhizanthes zippelii EF135269
Sapria himalayana individual 1
Sapria himalayana EF135273
Ricinus
Passiflora
Chrysobalanus
Salix
Bruguiera
Cannabis
Morus
Ziziphus
Malus
Vigna
Acacia
Tamarindus
Betula
Cucurbita
Fagus
Eucalyptus
Medinilla
Francoa
Lagerstroemia
Tetrastigma
Nekemias
Vitis
Sapria himalayana SH459 26972
Sapria himalayana EF135619
Spondias
Citrus
Gossypium
Carica
Heuchera
Arabidopsis
Acer
Euonymous
Nelumbo
Anemone
Oryza
Zea
Triticum
Spirodela
Allium
Asparagus
Phoenix
Phalaenopsis
Liriodendron
Magnolia
Schisandra
Nymphaea
Amborella
0.02

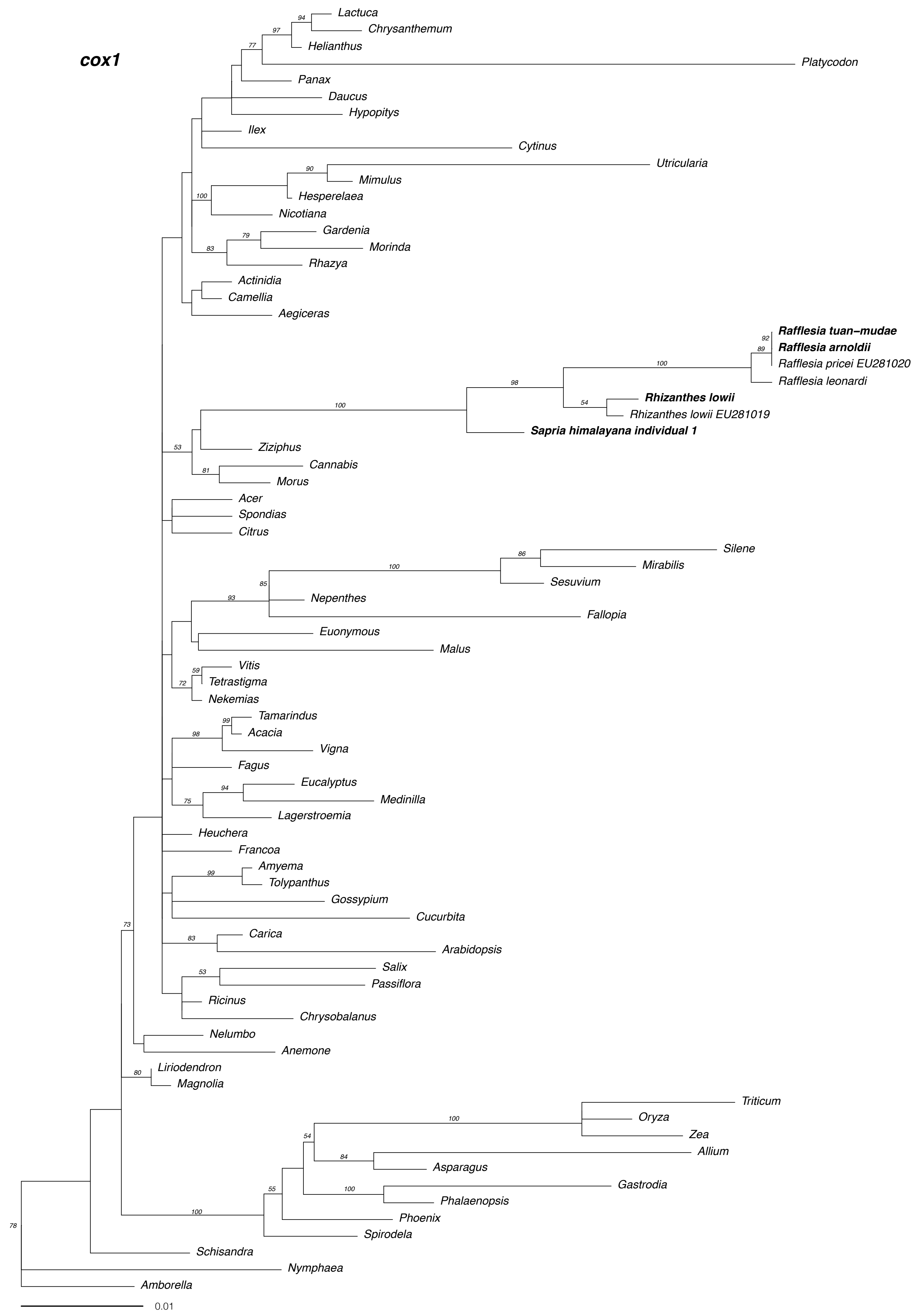

cox1
Lactuca
Chrysanthemum
Helianthus
Platycodon
Panax
Daucus
Hypopitys
Ilex
Cytinus
Utricularia
Mimulus
Hesperelaea
Nicotiana
Gardenia
Morinda
Rhazya
Actinidia
Camellia
Aegiceras
Rafflesia tuan-mudae
Rafflesia arnoldii
Rafflesia pricei EU281020
Rafflesia leonardi
Rhizanthes lowii
Rhizanthes lowii EU281019
Sapria himalayana individual 1
Ziziphus
Cannabis
Morus
Acer
Spondias
Citrus
Silene
Mirabilis
Sesuvium
Nepenthes
Fallopia
Euonymous
Malus
Vitis
Tetrastigma
Nekemias
Tamarindus
Acacia
Vigna
Fagus
Eucalyptus
Medinilla
Lagerstroemia
Heuchera
Francoa
Amyema
Tolypanthus
Gossypium
Cucurbita
Carica
Arabidopsis
Salix
Passiflora
Ricinus
Chrysobalanus
Nelumbo
Anemone
Liriodendron
Magnolia
Triticum
Oryza
Zea
Allium
Asparagus
Gastrodia
Phalaenopsis
Phoenix
Spirodela
Schisandra
Nymphaea
Amborella
0.01

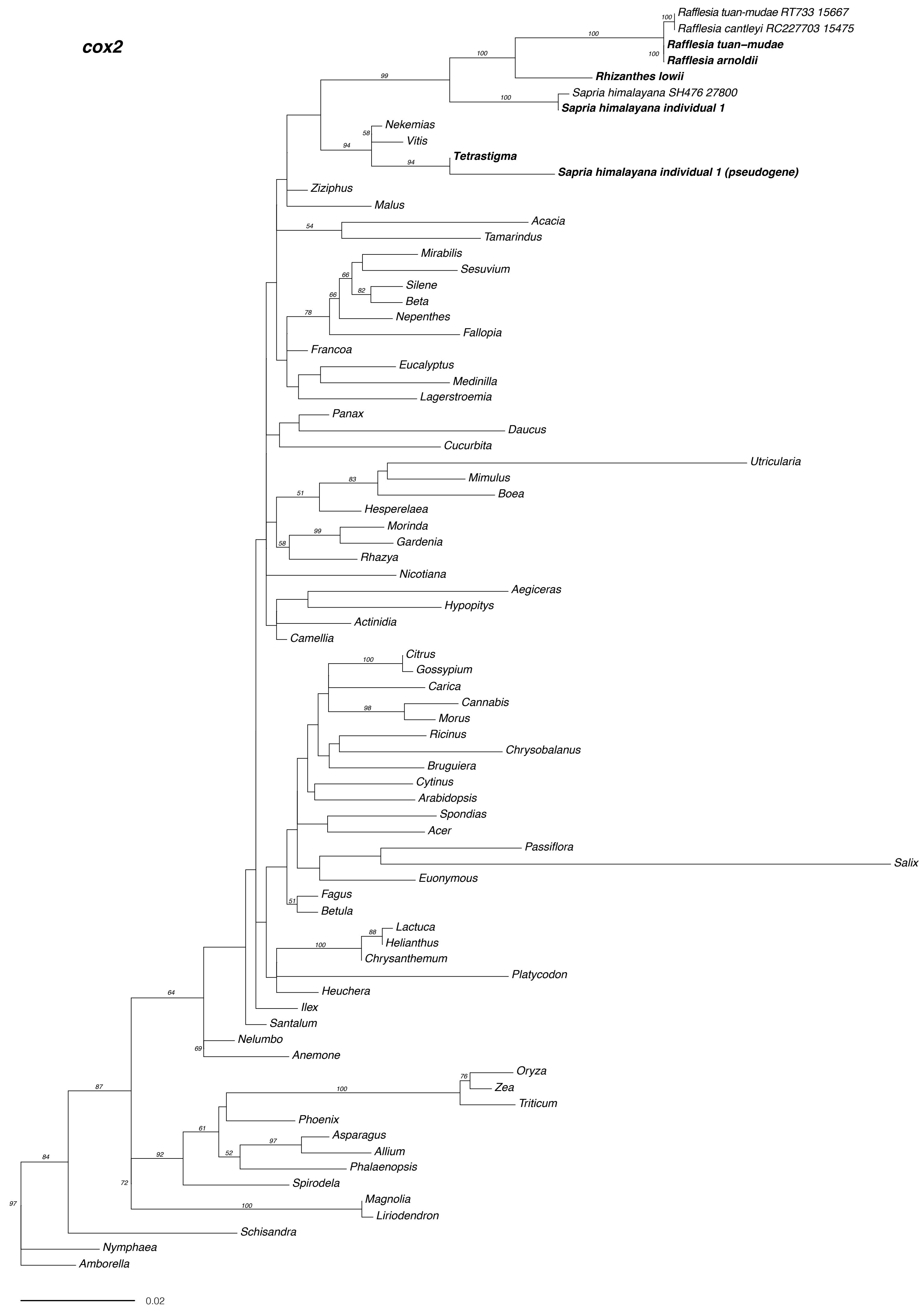

cox2
Rafflesia tuan-mudae RT733 15667
Rafflesia cantleyi RC227703 15475
Rafflesia tuan-mudae
Rafflesia arnoldii
Rhizanthes lowii
Sapria himalayana SH476 27800
Sapria himalayana individual 1
Nekemias
Vitis
Tetrastigma
Sapria himalayana individual 1 (pseudogene)
Ziziphus
Malus
Acacia
Tamarindus
Mirabilis
Sesuvium
Silene
Beta
Nepenthes
Fallopia
Francoa
Eucalyptus
Medinilla
Lagerstroemia
Panax
Daucus
Cucurbita
Utricularia
Mimulus
Boea
Hesperelaea
Morinda
Gardenia
Rhazya
Nicotiana
Aegiceras
Hypopitys
Actinidia
Camellia
Citrus
Gossypium
Carica
Cannabis
Morus
Ricinus
Chrysobalanus
Bruguiera
Cytinus
Arabidopsis
Spondias
Acer
Passiflora
Salix
Euonymous
Fagus
Betula
Lactuca
Helianthus
Chrysanthemum
Platycodon
Heuchera
Ilex
Santalum
Nelumbo
Anemone
Oryza
Zea
Triticum
Phoenix
Asparagus
Allium
Phalaenopsis
Spirodela
Magnolia
Liriodendron
Schisandra
Nymphaea
Amborella
0.02

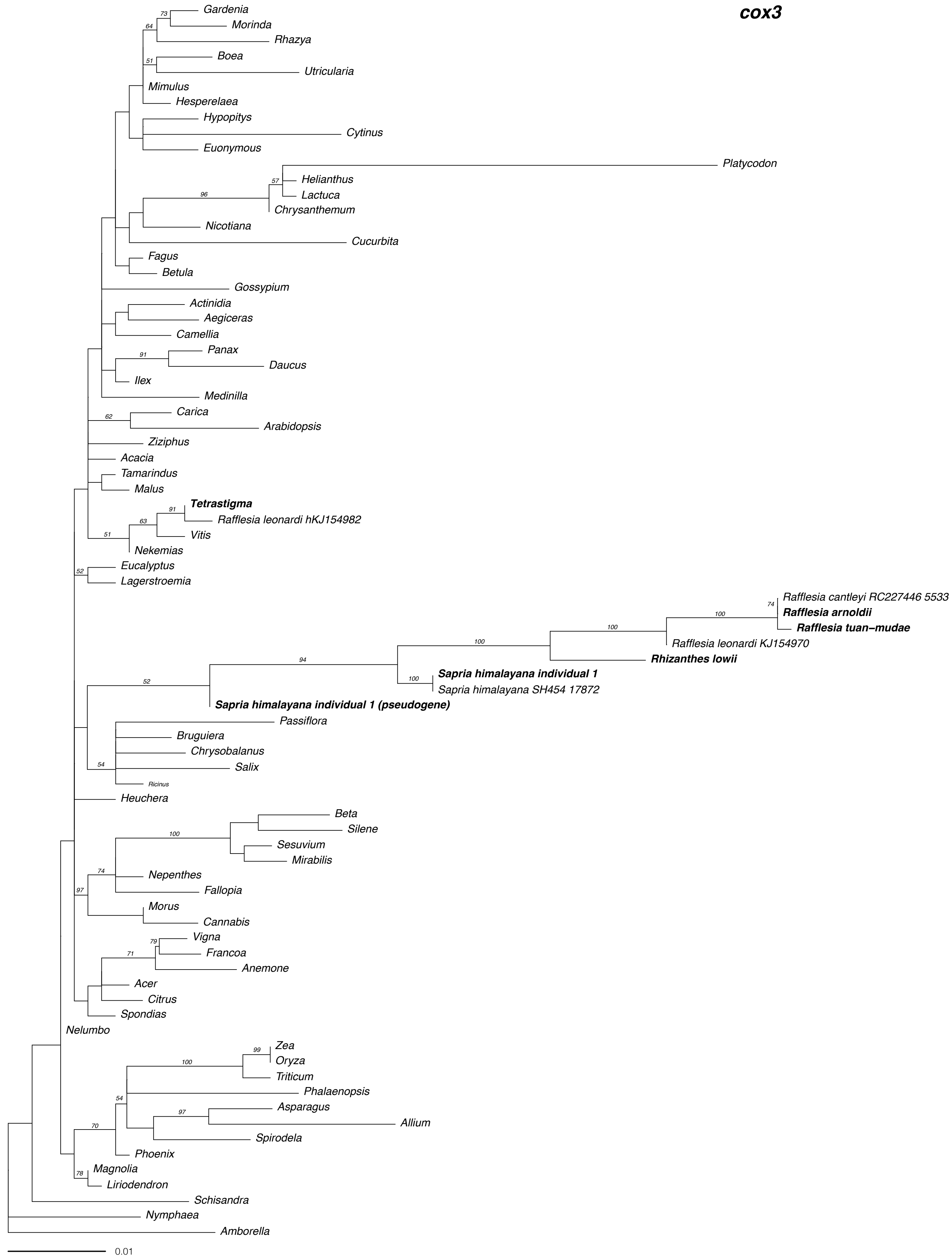

cox3
Gardenia
Morinda
Rhazya
Boea
Utricularia
Mimulus
Hesperelaea
Hypopitys
Cytinus
Euonymous
Platycodon
Helianthus
Lactuca
Chrysanthemum
Nicotiana
Cucurbita
Fagus
Betula
Gossypium
Actinidia
Aegiceras
Camellia
Panax
Daucus
Ilex
Medinilla
Carica
Arabidopsis
Ziziphus
Acacia
Tamarindus
Malus
Tetrastigma
Rafflesia leonardi hKJ154982
Vitis
Nekemias
Eucalyptus
Lagerstroemia
Rafflesia cantleyi RC227446 5533
Rafflesia arnoldii
Rafflesia tuan-mudae
Rafflesia leonardi KJ154970
Rhizanthes lowii
Sapria himalayana individual 1
Sapria himalayana SH454 17872
Sapria himalayana individual 1 (pseudogene)
Passiflora
Bruguiera
Chrysobalanus
Salix
Ricinus
Heuchera
Beta
Silene
Sesuvium
Mirabilis
Nepenthes
Fallopia
Morus
Cannabis
Vigna
Francoa
Anemone
Acer
Citrus
Spondias
Nelumbo
Zea
Oryza
Triticum
Phalaenopsis
Asparagus
Allium
Spirodela
Phoenix
Magnolia
Liriodendron
Schisandra
Nymphaea
Amborella
0.01

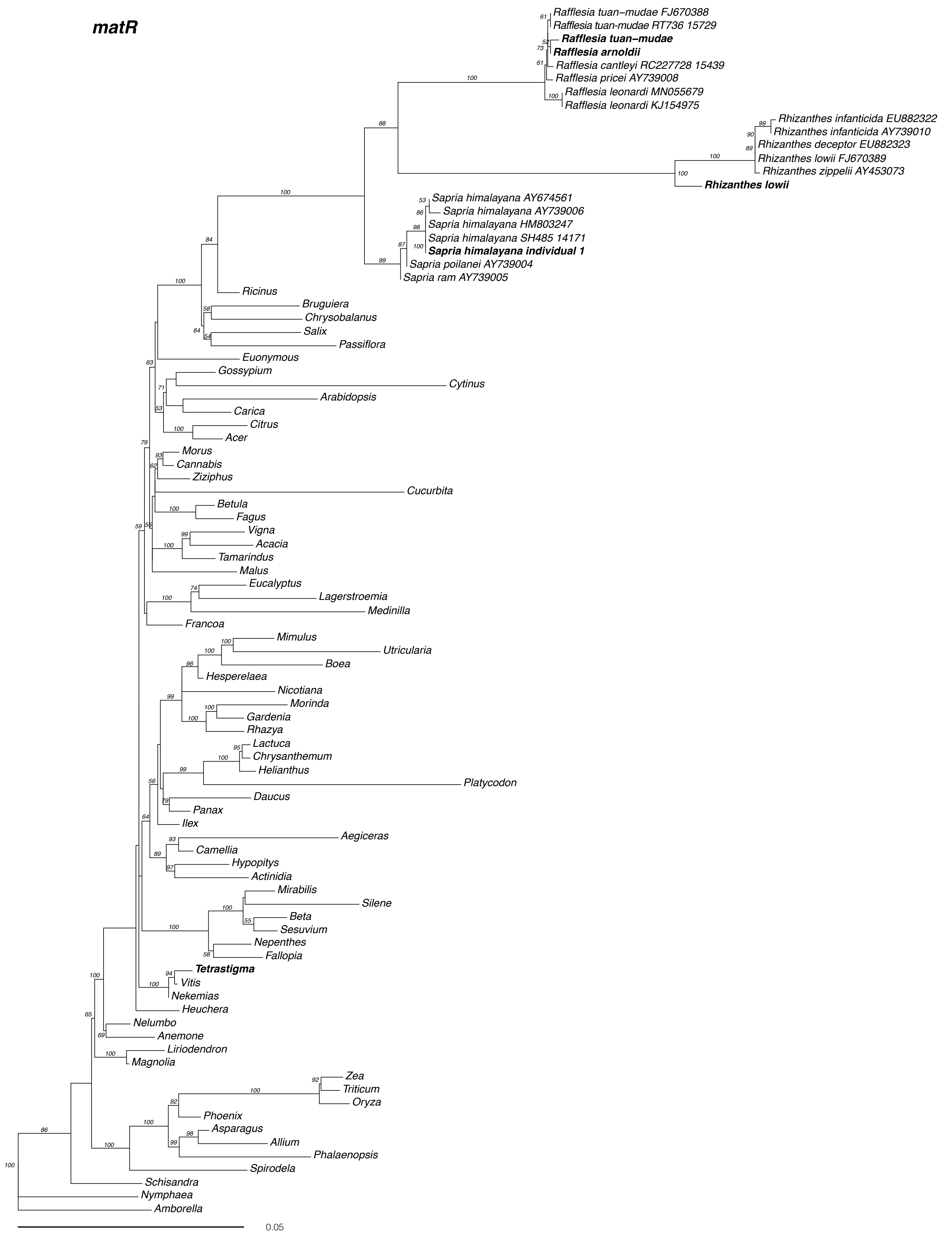
matR
Rafflesia tuan-mudae FJ670388
Rafflesia tuan-mudae RT736 15729
Rafflesia tuan-mudae
Rafflesia arnoldii
Rafflesia cantleyi RC227728 15439
Rafflesia pricei AY739008
Rafflesia leonardi MN055679
Rafflesia leonardi KJ154975
Rhizanthes infanticida EU882322
Rhizanthes infanticida AY739010
Rhizanthes deceptor EU882323
Rhizanthes lowii FJ670389
Rhizanthes zippelii AY453073
Rhizanthes lowii
Sapria himalayana AY674561
Sapria himalayana AY739006
Sapria himalayana HM803247
Sapria himalayana SH485 14171
Sapria himalayana individual 1
Sapria poilanei AY739004
Sapria ram AY739005
Ricinus
Bruguiera
Chrysobalanus
Salix
Passiflora
Euonymous
Gossypium
Cytinus
Arabidopsis
Carica
Citrus
Acer
Morus
Cannabis
Ziziphus
Cucurbita
Betula
Fagus
Vigna
Acacia
Tamarindus
Malus
Eucalyptus
Lagerstroemia
Medinilla
Francoa
Mimulus
Utricularia
Boea
Hesperelaea
Nicotiana
Morinda
Gardenia
Rhazya
Lactuca
Chrysanthemum
Helianthus
Platycodon
Daucus
Panax
Ilex
Aegiceras
Camellia
Hypopitys
Actinidia
Mirabilis
Silene
Beta
Sesuvium
Nepenthes
Fallopia
Tetrastigma
Vitis
Nekemias
Heuchera
Nelumbo
Anemone
Liriodendron
Magnolia
Zea
Triticum
Oryza
Phoenix
Asparagus
Allium
Phalaenopsis
Spirodela
Schisandra
Nymphaea
Amborella
0.05

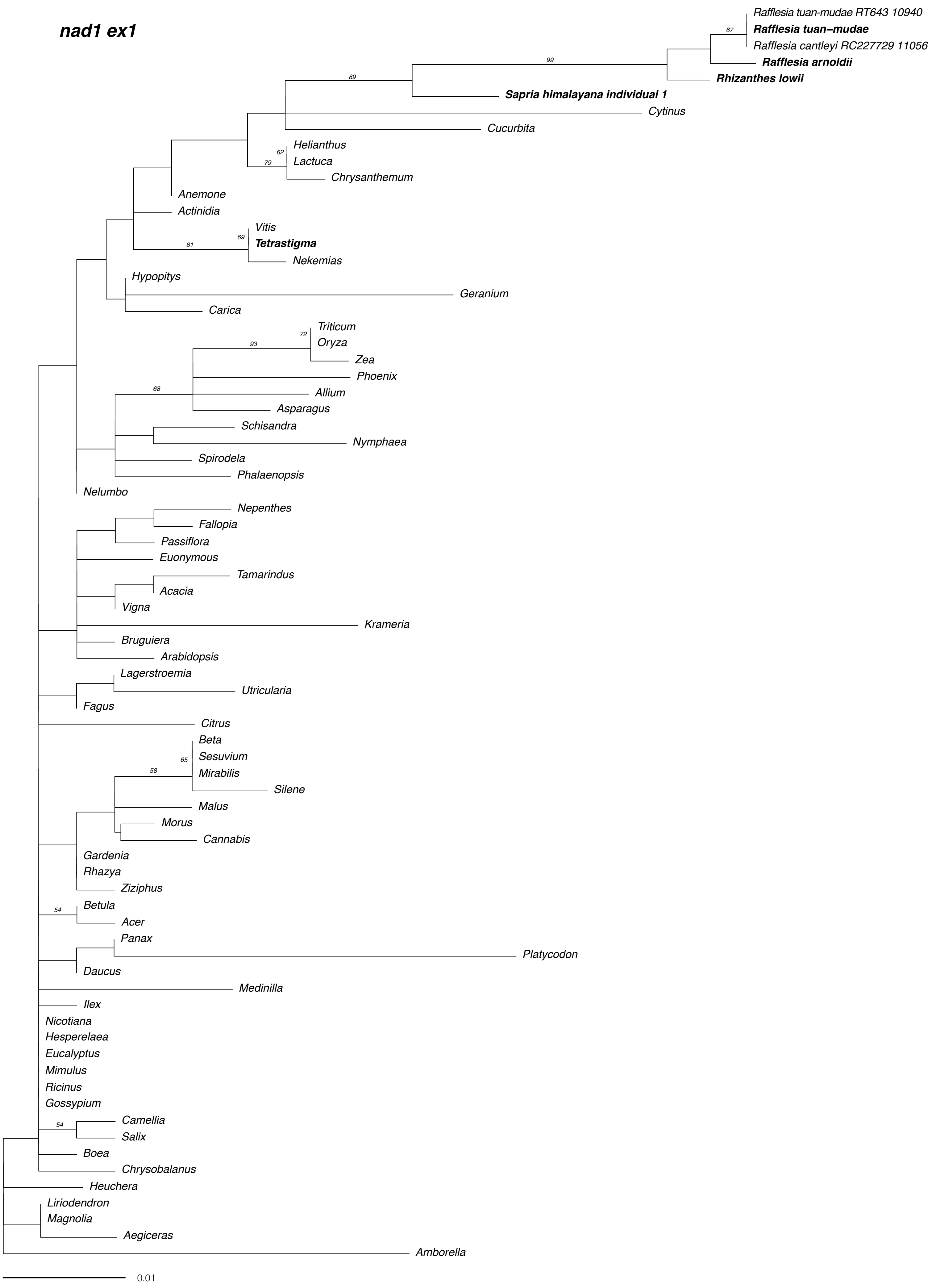

nad1 ex1
Rafflesia tuan-mudae RT643 10940
Rafflesia tuan–mudae
Rafflesia cantleyi RC227729 11056
Rafflesia arnoldii
Rhizanthes lowii
Sapria himalayana individual 1
Cytinus
Cucurbita
Helianthus
Lactuca
Chrysanthemum
Anemone
Actinidia
Vitis
Tetrastigma
Nekemias
Hypopitys
Geranium
Carica
Triticum
Oryza
Zea
Phoenix
Allium
Asparagus
Schisandra
Nymphaea
Spirodela
Phalaenopsis
Nelumbo
Nepenthes
Fallopia
Passiflora
Euonymous
Tamarindus
Acacia
Vigna
Krameria
Bruguiera
Arabidopsis
Lagerstroemia
Utricularia
Fagus
Citrus
Beta
Sesuvium
Mirabilis
Silene
Malus
Morus
Cannabis
Gardenia
Rhazya
Ziziphus
Betula
Acer
Panax
Platycodon
Daucus
Medinilla
Ilex
Nicotiana
Hesperelaea
Eucalyptus
Mimulus
Ricinus
Gossypium
Camellia
Salix
Boea
Chrysobalanus
Heuchera
Liriodendron
Magnolia
Aegiceras
Amborella
0.01

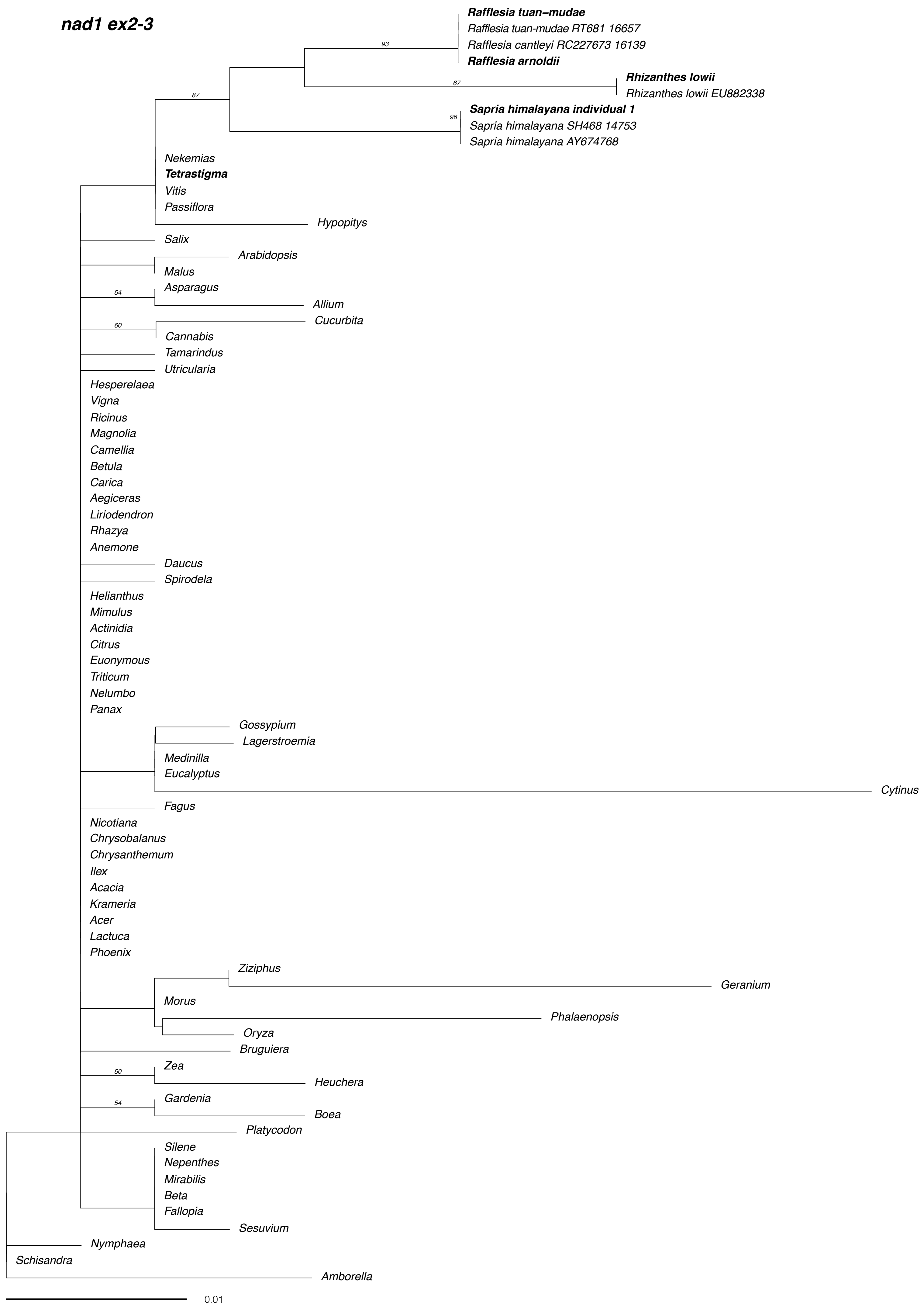
nad1 ex2-3
Rafflesia tuan–mudae
Rafflesia tuan-mudae RT681 16657
Rafflesia cantleyi RC227673 16139
Rafflesia arnoldii
93
Rhizanthes lowii
67
Rhizanthes lowii EU882338
87
Sapria himalayana individual 1
96
Sapria himalayana SH468 14753
Sapria himalayana AY674768
Nekemias
Tetrastigma
Vitis
Passiflora
Hypopitys
Salix
Arabidopsis
Malus
Asparagus
54
Allium
Cucurbita
60
Cannabis
Tamarindus
Utricularia
Hesperelaea
Vigna
Ricinus
Magnolia
Camellia
Betula
Carica
Aegiceras
Liriodendron
Rhazya
Anemone
Daucus
Spirodela
Helianthus
Mimulus
Actinidia
Citrus
Euonymous
Triticum
Nelumbo
Panax
Gossypium
Lagerstroemia
Medinilla
Eucalyptus
Cytinus
Fagus
Nicotiana
Chrysobalanus
Chrysanthemum
Ilex
Acacia
Krameria
Acer
Lactuca
Phoenix
Ziziphus
Geranium
Morus
Phalaenopsis
Oryza
Bruguiera
Zea
50
Heuchera
Gardenia
54
Boea
Platycodon
Silene
Nepenthes
Mirabilis
Beta
Fallopia
Sesuvium
Nymphaea
Schisandra
Amborella
0.01

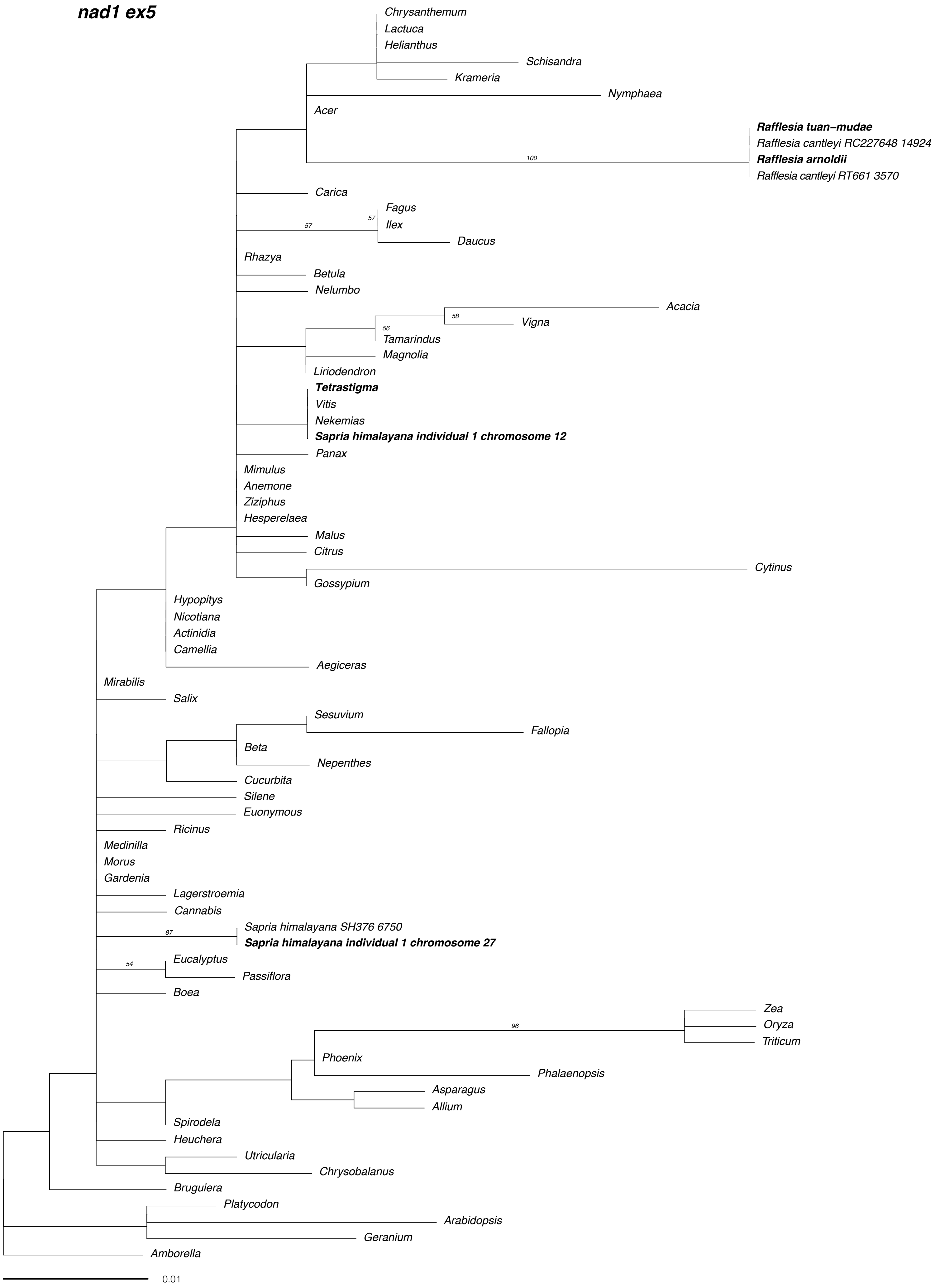
nad1 ex5
Chrysanthemum
Lactuca
Helianthus
Schisandra
Krameria
Nymphaea
Acer
Rafflesia tuan−mudae
Rafflesia cantleyi RC227648 14924
100
Rafflesia arnoldii
Rafflesia cantleyi RT661 3570
Carica
Fagus
57
57
Ilex
Daucus
Rhazya
Betula
Nelumbo
Acacia
58
Vigna
56
Tamarindus
Magnolia
Liriodendron
Tetrastigma
Vitis
Nekemias
Sapria himalayana individual 1 chromosome 12
Panax
Mimulus
Anemone
Ziziphus
Hesperelaea
Malus
Citrus
Cytinus
Gossypium
Hypopitys
Nicotiana
Actinidia
Camellia
Aegiceras
Mirabilis
Salix
Sesuvium
Fallopia
Beta
Nepenthes
Cucurbita
Silene
Euonymous
Ricinus
Medinilla
Morus
Gardenia
Lagerstroemia
Cannabis
Sapria himalayana SH376 6750
87
Sapria himalayana individual 1 chromosome 27
Eucalyptus
54
Passiflora
Boea
Zea
Oryza
96
Triticum
Phoenix
Phalaenopsis
Asparagus
Allium
Spirodela
Heuchera
Utricularia
Chrysobalanus
Bruguiera
Platycodon
Arabidopsis
Geranium
Amborella
0.01

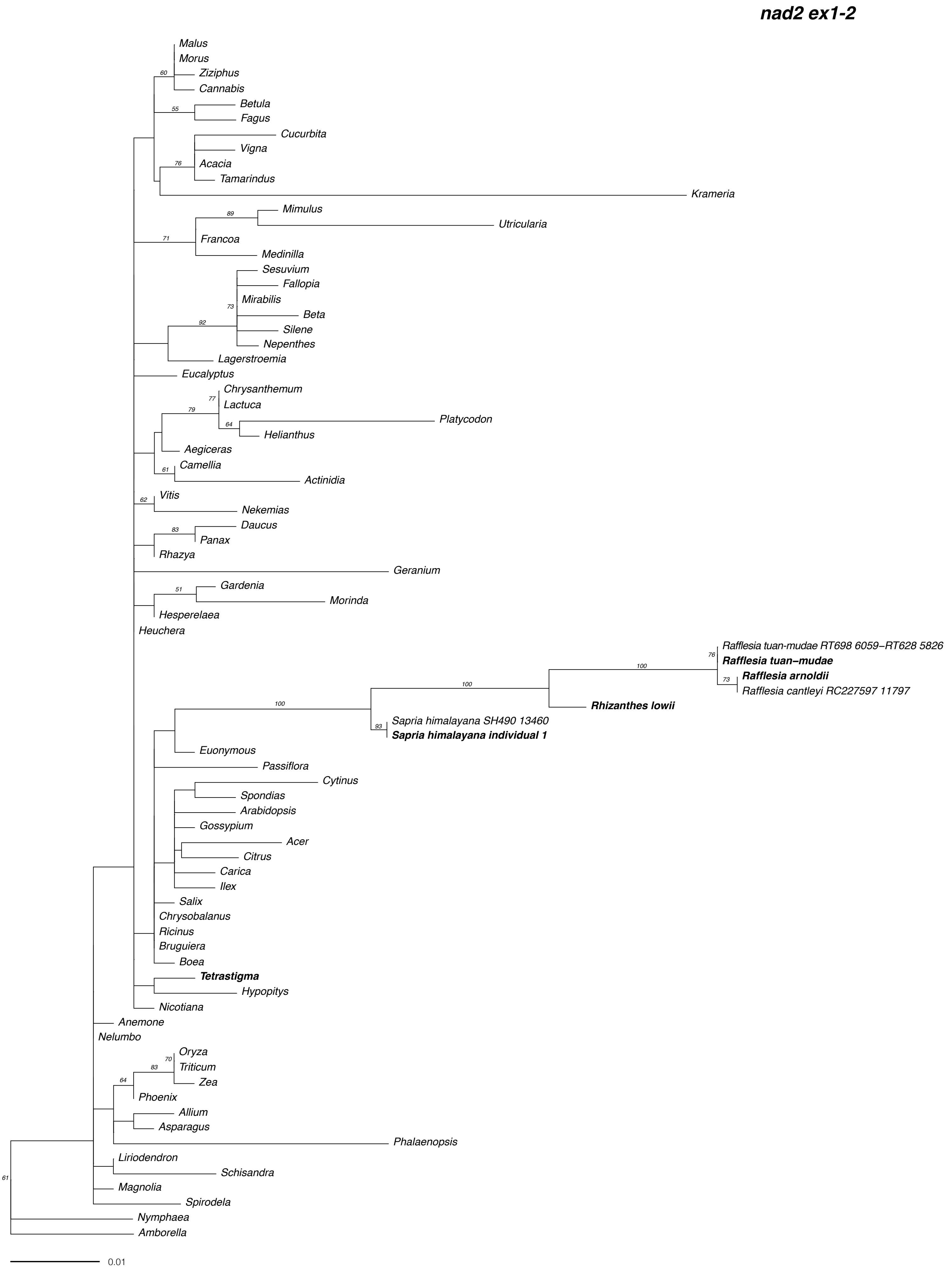
nad2 ex1-2
Malus
Morus
Ziziphus
Cannabis
60
Betula
Fagus
55
Cucurbita
Vigna
Acacia
Tamarindus
76
Krameria
Mimulus
Utricularia
89
Francoa
71
Medinilla
Sesuvium
Fallopia
Mirabilis
73
Beta
Silene
Nepenthes
92
Lagerstroemia
Eucalyptus
Chrysanthemum
77
Lactuca
79
Platycodon
64
Helianthus
Aegiceras
Camellia
61
Actinidia
Vitis
62
Nekemias
Daucus
83
Panax
Rhazya
Geranium
Gardenia
51
Morinda
Hesperelaea
Heuchera
Rafflesia tuan-mudae RT698 6059–RT628 5826
76
Rafflesia tuan–mudae
100
Rafflesia arnoldii
73
Rafflesia cantleyi RC227597 11797
100
Rhizanthes lowii
100
Sapria himalayana SH490 13460
93
Sapria himalayana individual 1
Euonymous
Passiflora
Cytinus
Spondias
Arabidopsis
Gossypium
Acer
Citrus
Carica
Ilex
Salix
Chrysobalanus
Ricinus
Bruguiera
Boea
Tetrastigma
Hypopitys
Nicotiana
Anemone
Nelumbo
Oryza
70
Triticum
83
Zea
64
Phoenix
Allium
Asparagus
Phalaenopsis
Liriodendron
Schisandra
Magnolia
Spirodela
61
Nymphaea
Amborella
0.01

**nad2 ex3-4-5**

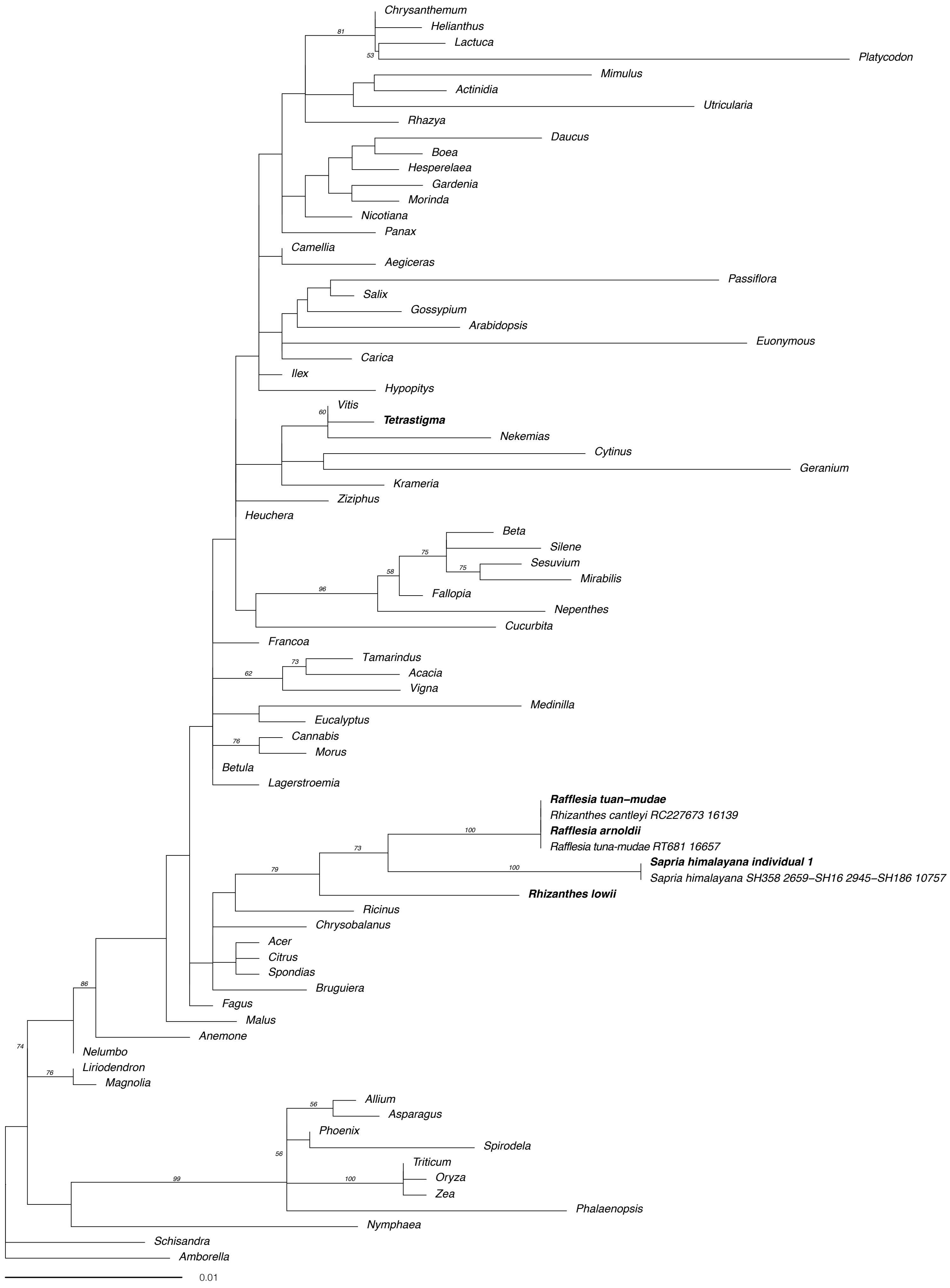
Chrysanthemum
Helianthus
Lactuca
Platycodon
81
53
Mimulus
Actinidia
Utricularia
Rhazya
Daucus
Boea
Hesperelaea
Gardenia
Morinda
Nicotiana
Panax
Camellia
Aegiceras
Passiflora
Salix
Gossypium
Arabidopsis
Euonymous
Carica
Ilex
Hypopitys
Vitis
60
Tetrastigma
Nekemias
Cytinus
Geranium
Krameria
Ziziphus
Heuchera
Beta
Silene
75
Sesuvium
75
Mirabilis
58
96
Fallopia
Nepenthes
Cucurbita
Francoa
Tamarindus
73
Acacia
62
Vigna
Medinilla
Eucalyptus
Cannabis
76
Morus
Betula
Lagerstroemia
Rafflesia tuan-mudae
Rhizanthes cantleyi RC227673 16139
Rafflesia arnoldii
100
Rafflesia tuna-mudae RT681 16657
73
Sapria himalayana individual 1
100
Sapria himalayana SH358 2659–SH16 2945–SH186 10757
79
Rhizanthes lowii
Ricinus
Chrysobalanus
Acer
Citrus
Spondias
Bruguiera
Fagus
Malus
86
Anemone
74
Nelumbo
Liriodendron
76
Magnolia
Allium
56
Asparagus
Phoenix
Spirodela
56
Triticum
99
100
Oryza
Zea
Phalaenopsis
Nymphaea
Schisandra
Amborella
0.01

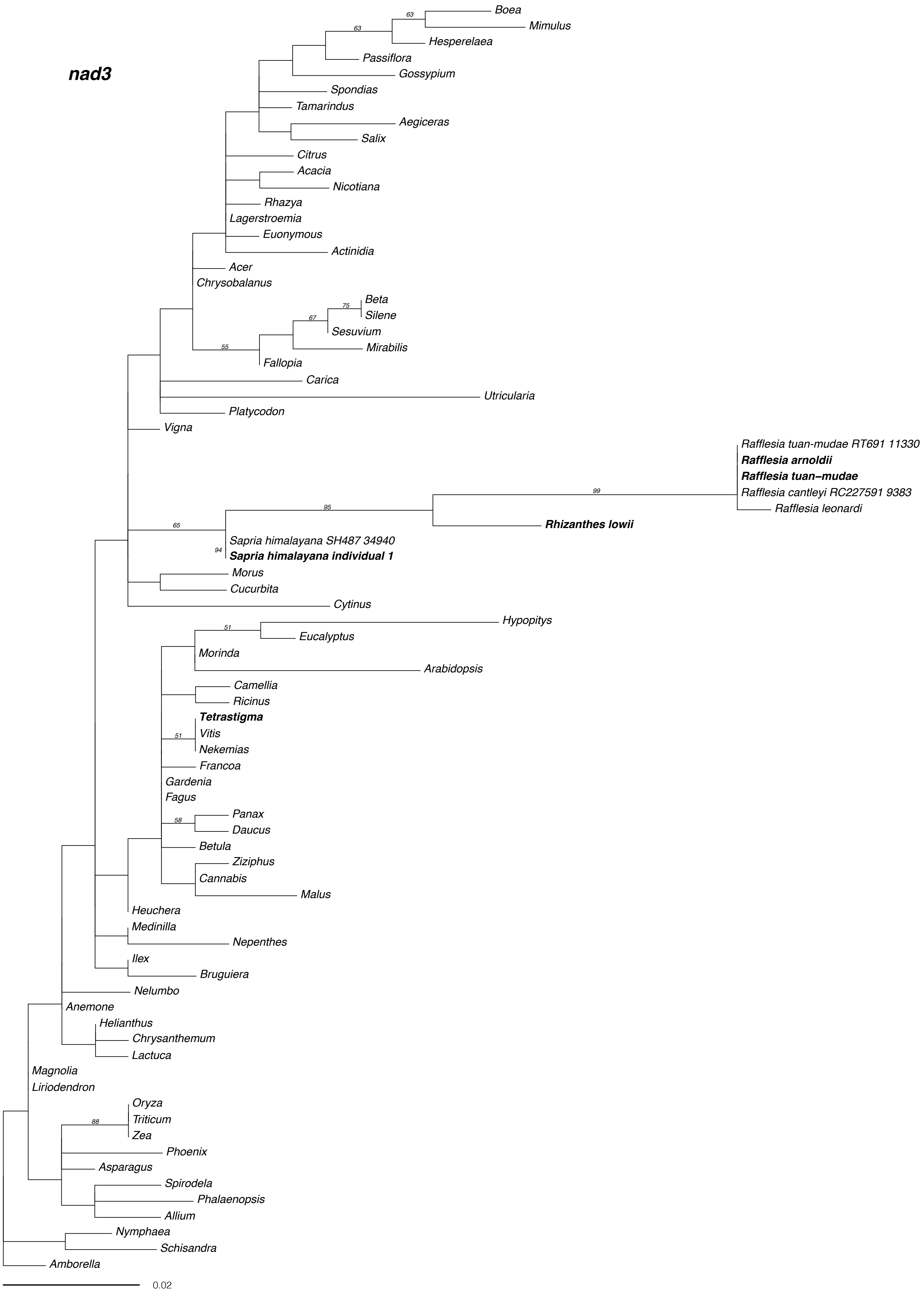
nad3
Boea
Mimulus
Hesperelaea
Passiflora
Gossypium
Spondias
Tamarindus
Aegiceras
Salix
Citrus
Acacia
Nicotiana
Rhazya
Lagerstroemia
Euonymous
Actinidia
Acer
Chrysobalanus
Beta
Silene
Sesuvium
Mirabilis
Fallopia
Carica
Utricularia
Platycodon
Vigna
Rafflesia tuan-mudae RT691 11330
Rafflesia arnoldii
Rafflesia tuan–mudae
Rafflesia cantleyi RC227591 9383
Rafflesia leonardi
Rhizanthes lowii
Sapria himalayana SH487 34940
Sapria himalayana individual 1
Morus
Cucurbita
Cytinus
Hypopitys
Eucalyptus
Morinda
Arabidopsis
Camellia
Ricinus
Tetrastigma
Vitis
Nekemias
Francoa
Gardenia
Fagus
Panax
Daucus
Betula
Ziziphus
Cannabis
Malus
Heuchera
Medinilla
Nepenthes
Ilex
Bruguiera
Nelumbo
Anemone
Helianthus
Chrysanthemum
Lactuca
Magnolia
Liriodendron
Oryza
Triticum
Zea
Phoenix
Asparagus
Spirodela
Phalaenopsis
Allium
Nymphaea
Schisandra
Amborella
0.02

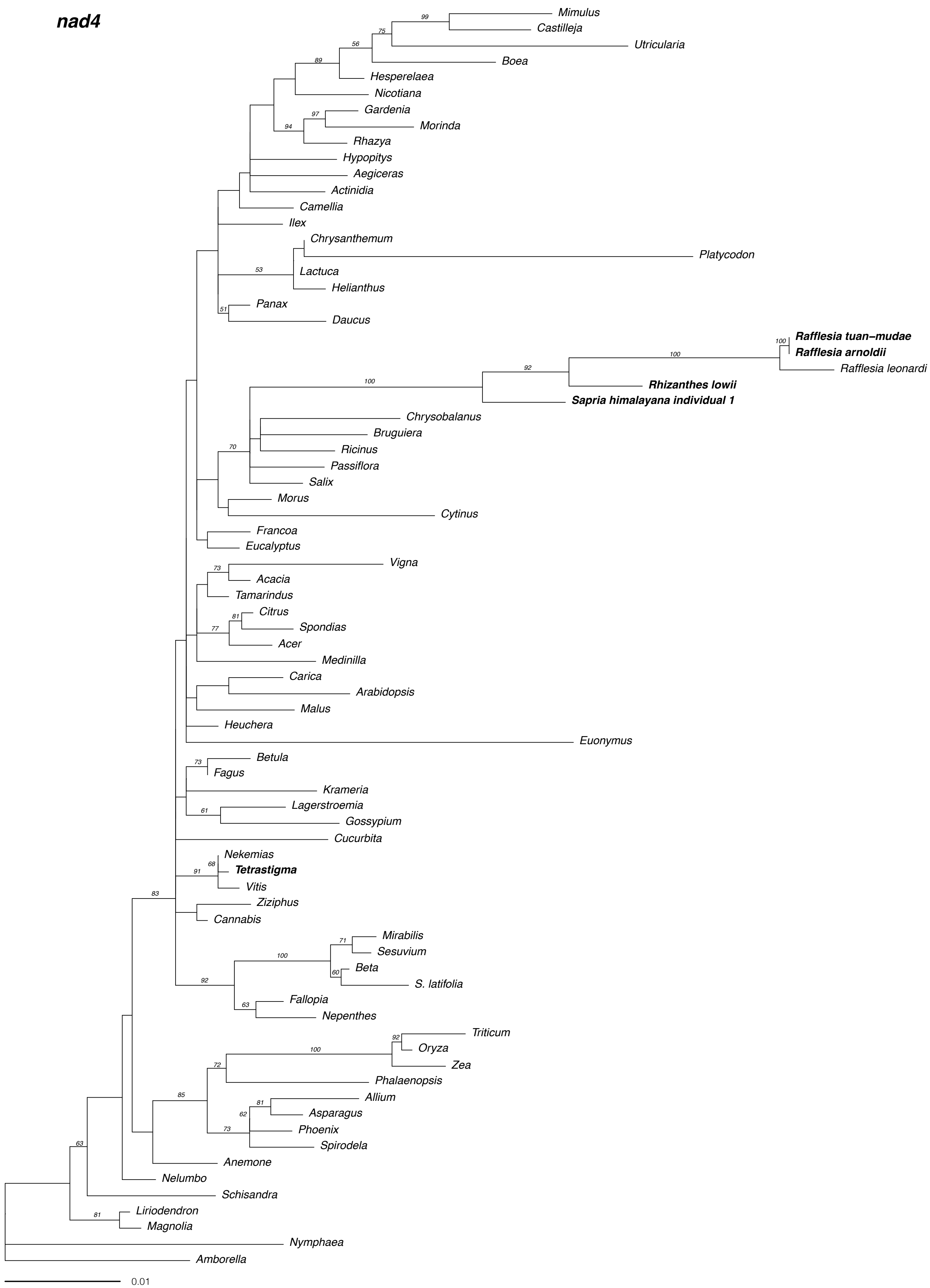

nad4
Mimulus
Castilleja
Utricularia
Boea
Hesperelaea
Nicotiana
Gardenia
Morinda
Rhazya
Hypopitys
Aegiceras
Actinidia
Camellia
Ilex
Chrysanthemum
Platycodon
Lactuca
Helianthus
Panax
Daucus
Rafflesia tuan-mudae
Rafflesia arnoldii
Rafflesia leonardi
Rhizanthes lowii
Sapria himalayana individual 1
Chrysobalanus
Bruguiera
Ricinus
Passiflora
Salix
Morus
Cytinus
Francoa
Eucalyptus
Vigna
Acacia
Tamarindus
Citrus
Spondias
Acer
Medinilla
Carica
Arabidopsis
Malus
Heuchera
Euonymus
Betula
Fagus
Krameria
Lagerstroemia
Gossypium
Cucurbita
Nekemias
Tetrastigma
Vitis
Ziziphus
Cannabis
Mirabilis
Sesuvium
Beta
S. latifolia
Fallopia
Nepenthes
Triticum
Oryza
Zea
Phalaenopsis
Allium
Asparagus
Phoenix
Spirodela
Anemone
Nelumbo
Schisandra
Liriodendron
Magnolia
Nymphaea
Amborella
0.01

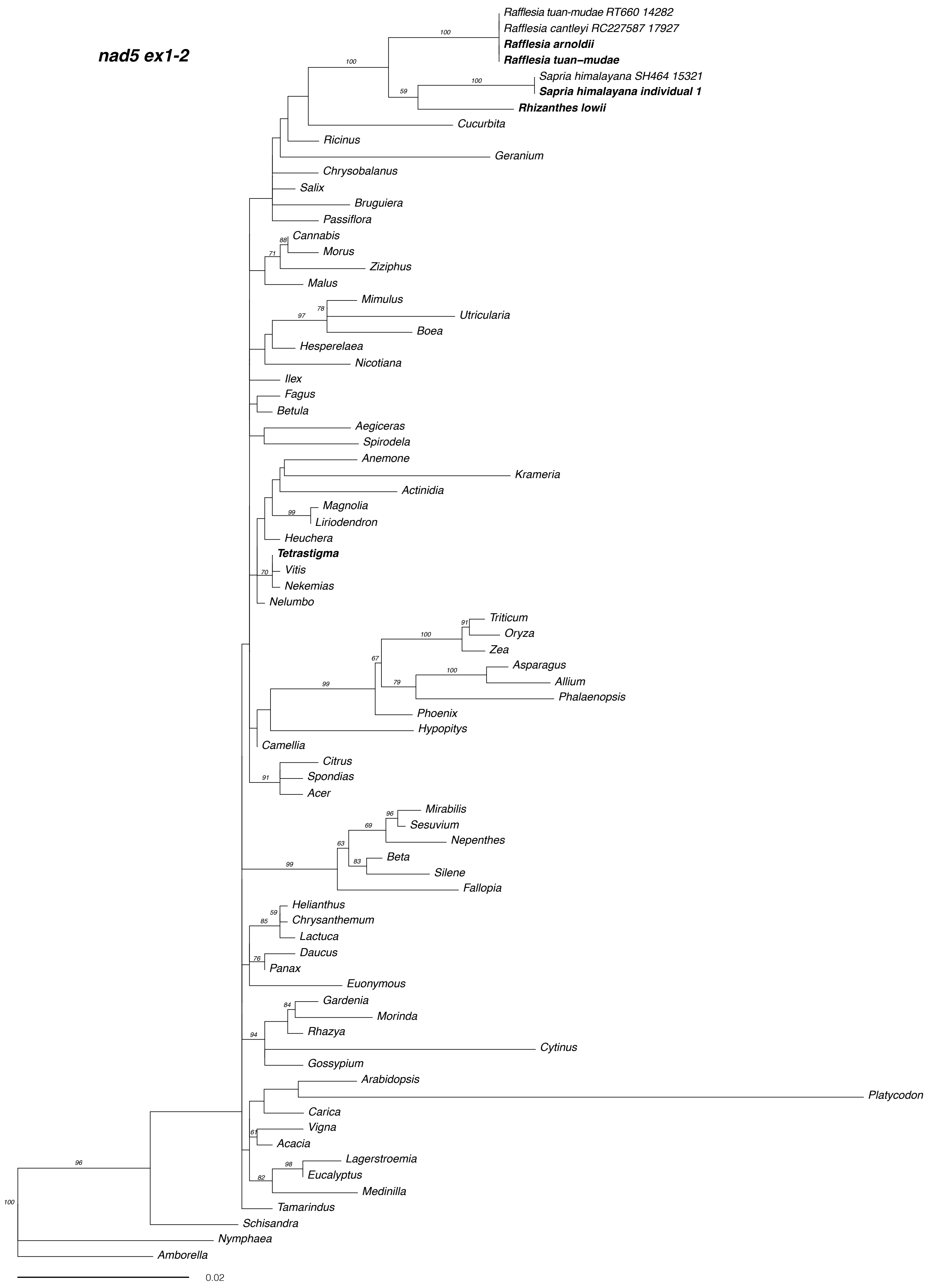
nad5 ex1-2
Rafflesia tuan-mudae RT660 14282
Rafflesia cantleyi RC227587 17927
Rafflesia arnoldii
Rafflesia tuan–mudae
Sapria himalayana SH464 15321
Sapria himalayana individual 1
Rhizanthes lowii
Cucurbita
Ricinus
Geranium
Chrysobalanus
Salix
Bruguiera
Passiflora
Cannabis
Morus
Ziziphus
Malus
Mimulus
Utricularia
Boea
Hesperelaea
Nicotiana
Ilex
Fagus
Betula
Aegiceras
Spirodela
Anemone
Krameria
Actinidia
Magnolia
Liriodendron
Heuchera
Tetrastigma
Vitis
Nekemias
Nelumbo
Triticum
Oryza
Zea
Asparagus
Allium
Phalaenopsis
Phoenix
Hypopitys
Camellia
Citrus
Spondias
Acer
Mirabilis
Sesuvium
Nepenthes
Beta
Silene
Fallopia
Helianthus
Chrysanthemum
Lactuca
Daucus
Panax
Euonymous
Gardenia
Morinda
Rhazya
Cytinus
Gossypium
Arabidopsis
Platycodon
Carica
Vigna
Acacia
Lagerstroemia
Eucalyptus
Medinilla
Tamarindus
Schisandra
Nymphaea
Amborella
0.02

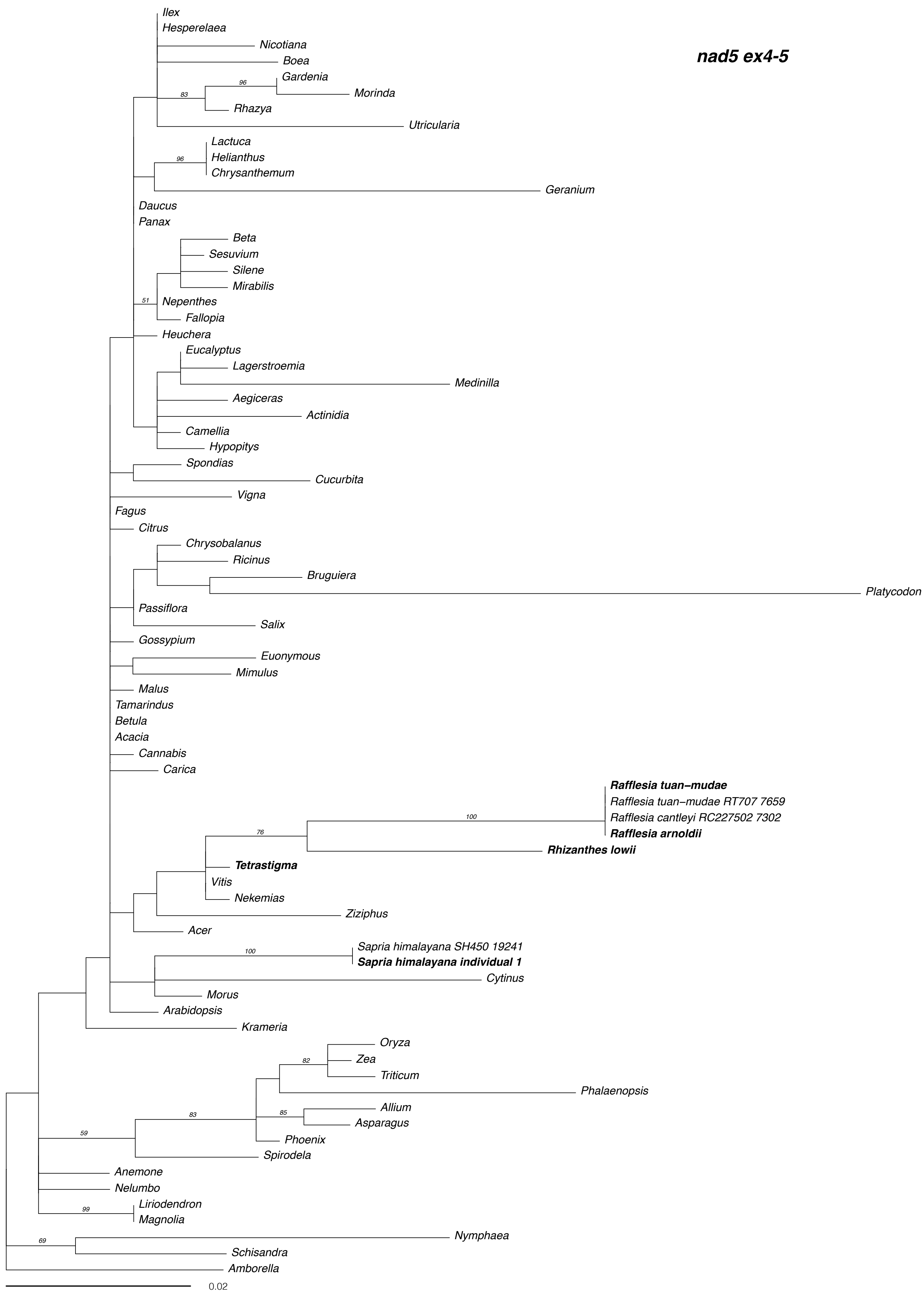
nad5 ex4-5
Ilex
Hesperelaea
Nicotiana
Boea
Gardenia
Morinda
Rhazya
Utricularia
Lactuca
Helianthus
Chrysanthemum
Geranium
Daucus
Panax
Beta
Sesuvium
Silene
Mirabilis
Nepenthes
Fallopia
Heuchera
Eucalyptus
Lagerstroemia
Medinilla
Aegiceras
Actinidia
Camellia
Hypopitys
Spondias
Cucurbita
Vigna
Fagus
Citrus
Chrysobalanus
Ricinus
Bruguiera
Platycodon
Passiflora
Salix
Gossypium
Euonymous
Mimulus
Malus
Tamarindus
Betula
Acacia
Cannabis
Carica
Rafflesia tuan-mudae
Rafflesia tuan-mudae RT707 7659
Rafflesia cantleyi RC227502 7302
Rafflesia arnoldii
Rhizanthes lowii
Tetrastigma
Vitis
Nekemias
Ziziphus
Acer
Sapria himalayana SH450 19241
Sapria himalayana individual 1
Cytinus
Morus
Arabidopsis
Krameria
Oryza
Zea
Triticum
Phalaenopsis
Allium
Asparagus
Phoenix
Spirodela
Anemone
Nelumbo
Liriodendron
Magnolia
Nymphaea
Schisandra
Amborella
0.02

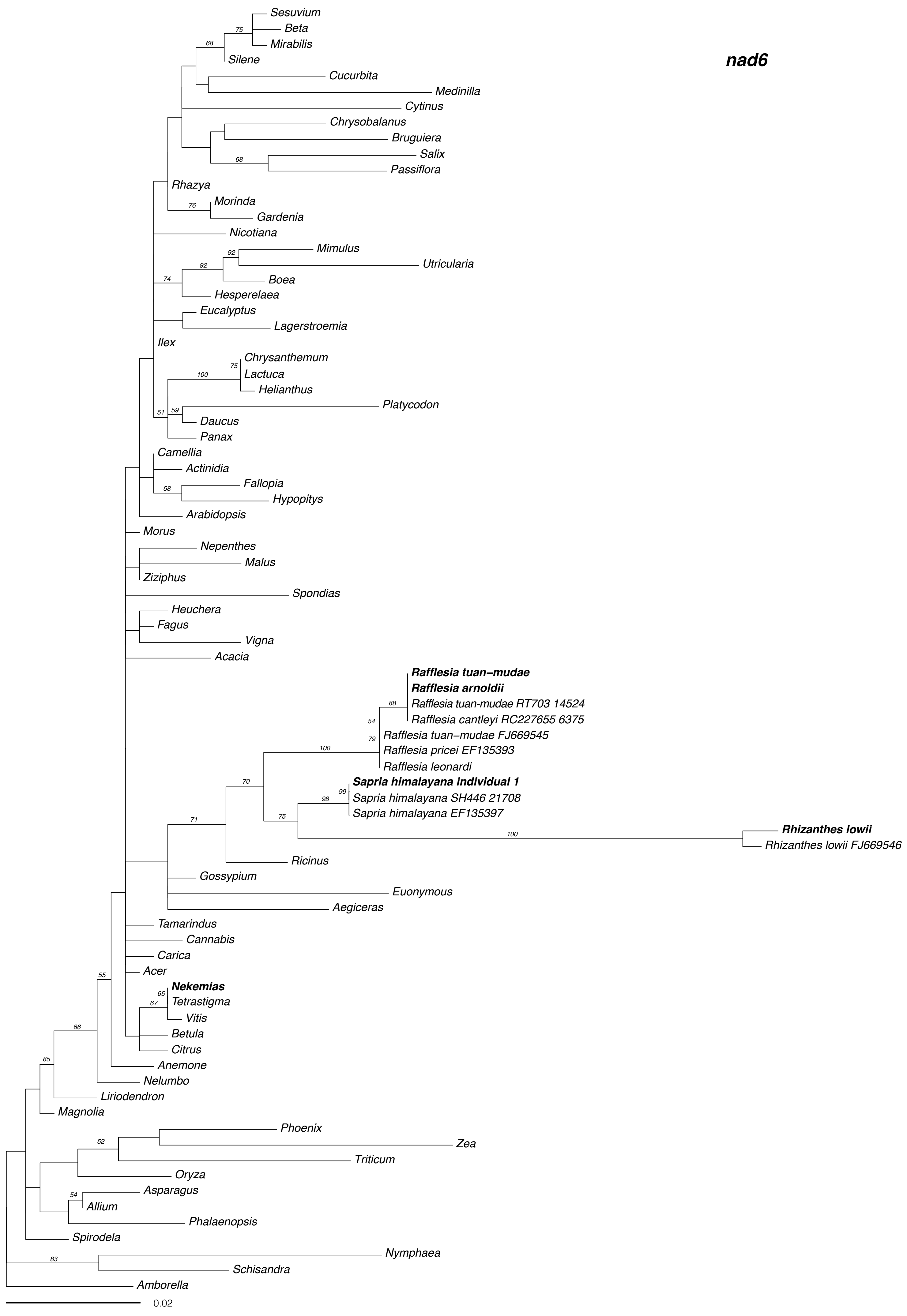

nad6
Sesuvium
Beta
Mirabilis
Silene
Cucurbita
Medinilla
Cytinus
Chrysobalanus
Bruguiera
Salix
Passiflora
Rhazya
Morinda
Gardenia
Nicotiana
Mimulus
Utricularia
Boea
Hesperelaea
Eucalyptus
Lagerstroemia
Ilex
Chrysanthemum
Lactuca
Helianthus
Platycodon
Daucus
Panax
Camellia
Actinidia
Fallopia
Hypopitys
Arabidopsis
Morus
Nepenthes
Malus
Ziziphus
Spondias
Heuchera
Fagus
Vigna
Acacia
Rafflesia tuan-mudae
Rafflesia arnoldii
Rafflesia tuan-mudae RT703 14524
Rafflesia cantleyi RC227655 6375
Rafflesia tuan-mudae FJ669545
Rafflesia pricei EF135393
Rafflesia leonardi
Sapria himalayana individual 1
Sapria himalayana SH446 21708
Sapria himalayana EF135397
Rhizanthes lowii
Rhizanthes lowii FJ669546
Ricinus
Gossypium
Euonymous
Aegiceras
Tamarindus
Cannabis
Carica
Acer
Nekemias
Tetrastigma
Vitis
Betula
Citrus
Anemone
Nelumbo
Liriodendron
Magnolia
Phoenix
Zea
Triticum
Oryza
Asparagus
Allium
Phalaenopsis
Spirodela
Nymphaea
Schisandra
Amborella
0.02

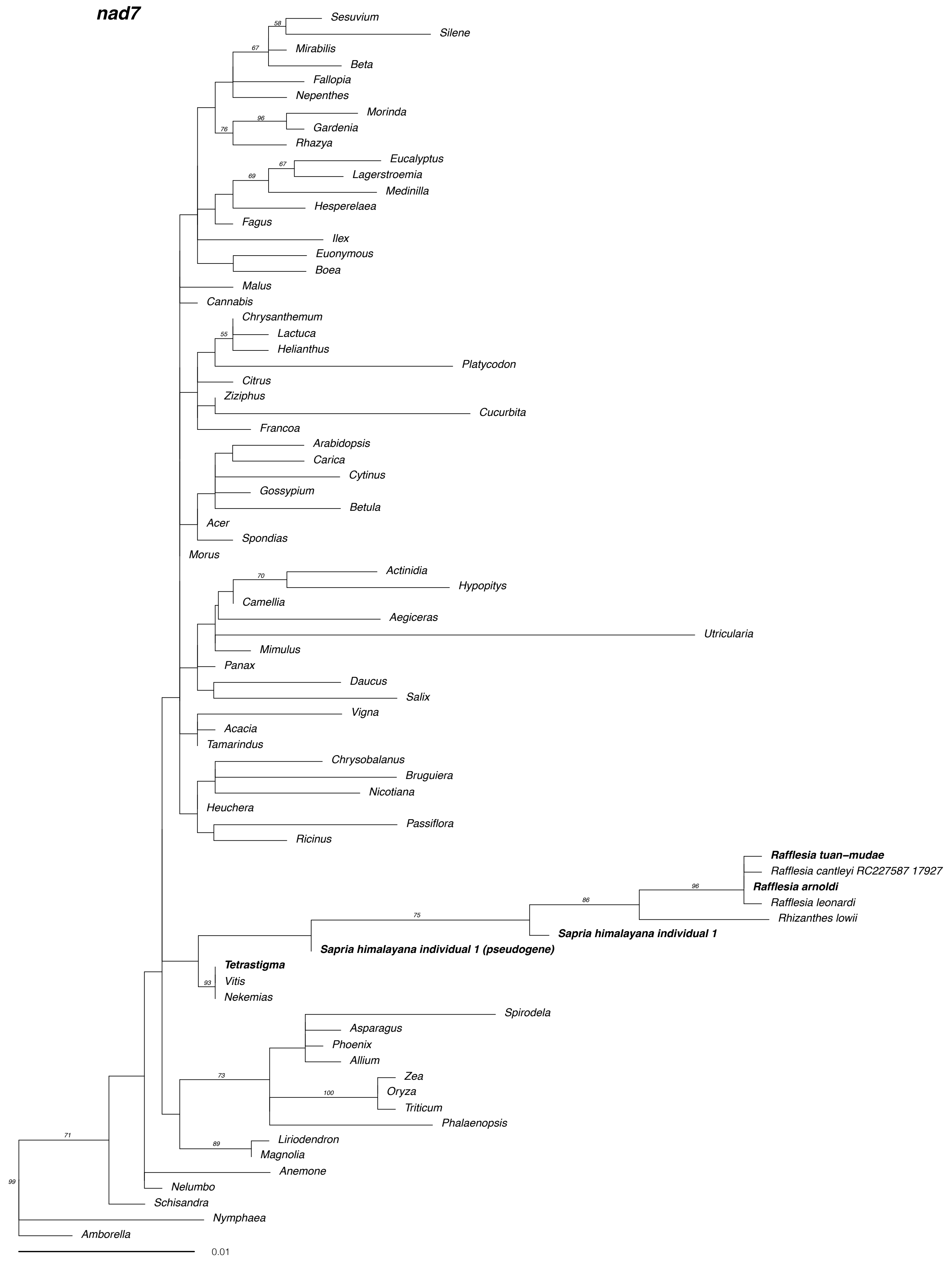
nad7
58
Sesuvium
Silene
67
Mirabilis
Beta
Fallopia
Nepenthes
96
Morinda
76
Gardenia
Rhazya
67
Eucalyptus
69
Lagerstroemia
Medinilla
Hesperelaea
Fagus
Ilex
Euonymous
Boea
Malus
Cannabis
Chrysanthemum
55
Lactuca
Helianthus
Platycodon
Citrus
Ziziphus
Cucurbita
Francoa
Arabidopsis
Carica
Cytinus
Gossypium
Betula
Acer
Spondias
Morus
70
Actinidia
Hypopitys
Camellia
Aegiceras
Utricularia
Mimulus
Panax
Daucus
Salix
Vigna
Acacia
Tamarindus
Chrysobalanus
Bruguiera
Nicotiana
Heuchera
Passiflora
Ricinus
Rafflesia tuan-mudae
Rafflesia cantleyi RC227587 17927
96
Rafflesia arnoldi
86
Rafflesia leonardi
75
Rhizanthes lowii
Sapria himalayana individual 1
Sapria himalayana individual 1 (pseudogene)
Tetrastigma
93
Vitis
Nekemias
Spirodela
Asparagus
Phoenix
Allium
Zea
73
100
Oryza
Triticum
Phalaenopsis
89
Liriodendron
Magnolia
71
Anemone
99
Nelumbo
Schisandra
Nymphaea
Amborella
0.01

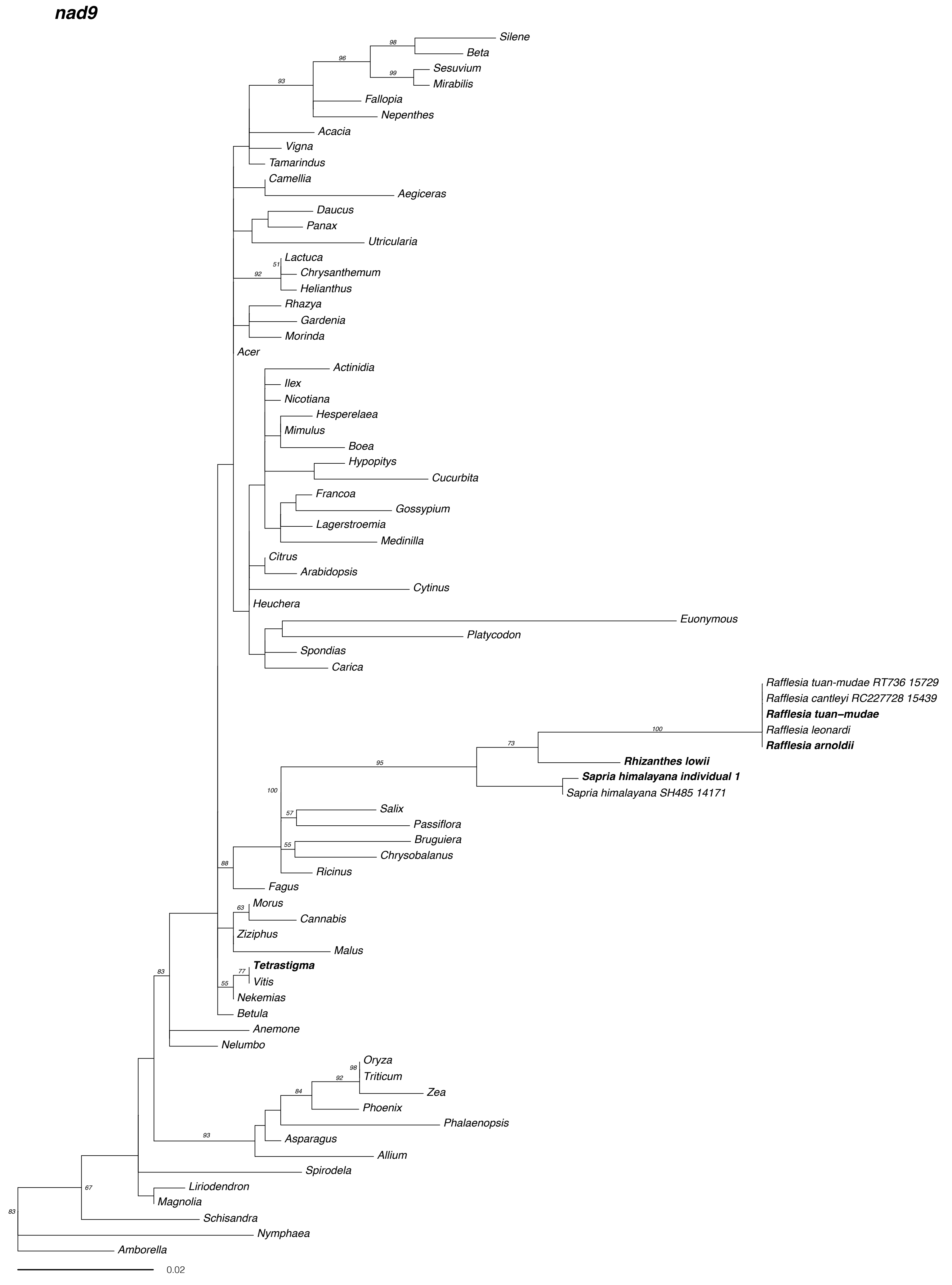
nad9
Silene
Beta
Sesuvium
Mirabilis
Fallopia
Nepenthes
Acacia
Vigna
Tamarindus
Camellia
Aegiceras
Daucus
Panax
Utricularia
Lactuca
Chrysanthemum
Helianthus
Rhazya
Gardenia
Morinda
Acer
Actinidia
Ilex
Nicotiana
Hesperelaea
Mimulus
Boea
Hypopitys
Cucurbita
Francoa
Gossypium
Lagerstroemia
Medinilla
Citrus
Arabidopsis
Cytinus
Heuchera
Euonymous
Platycodon
Spondias
Carica
Rafflesia tuan-mudae RT736 15729
Rafflesia cantleyi RC227728 15439
Rafflesia tuan–mudae
Rafflesia leonardi
Rafflesia arnoldii
Rhizanthes lowii
Sapria himalayana individual 1
Sapria himalayana SH485 14171
Salix
Passiflora
Bruguiera
Chrysobalanus
Ricinus
Fagus
Morus
Cannabis
Ziziphus
Malus
Tetrastigma
Vitis
Nekemias
Betula
Anemone
Nelumbo
Oryza
Triticum
Zea
Phoenix
Phalaenopsis
Asparagus
Allium
Spirodela
Liriodendron
Magnolia
Schisandra
Nymphaea
Amborella
0.02

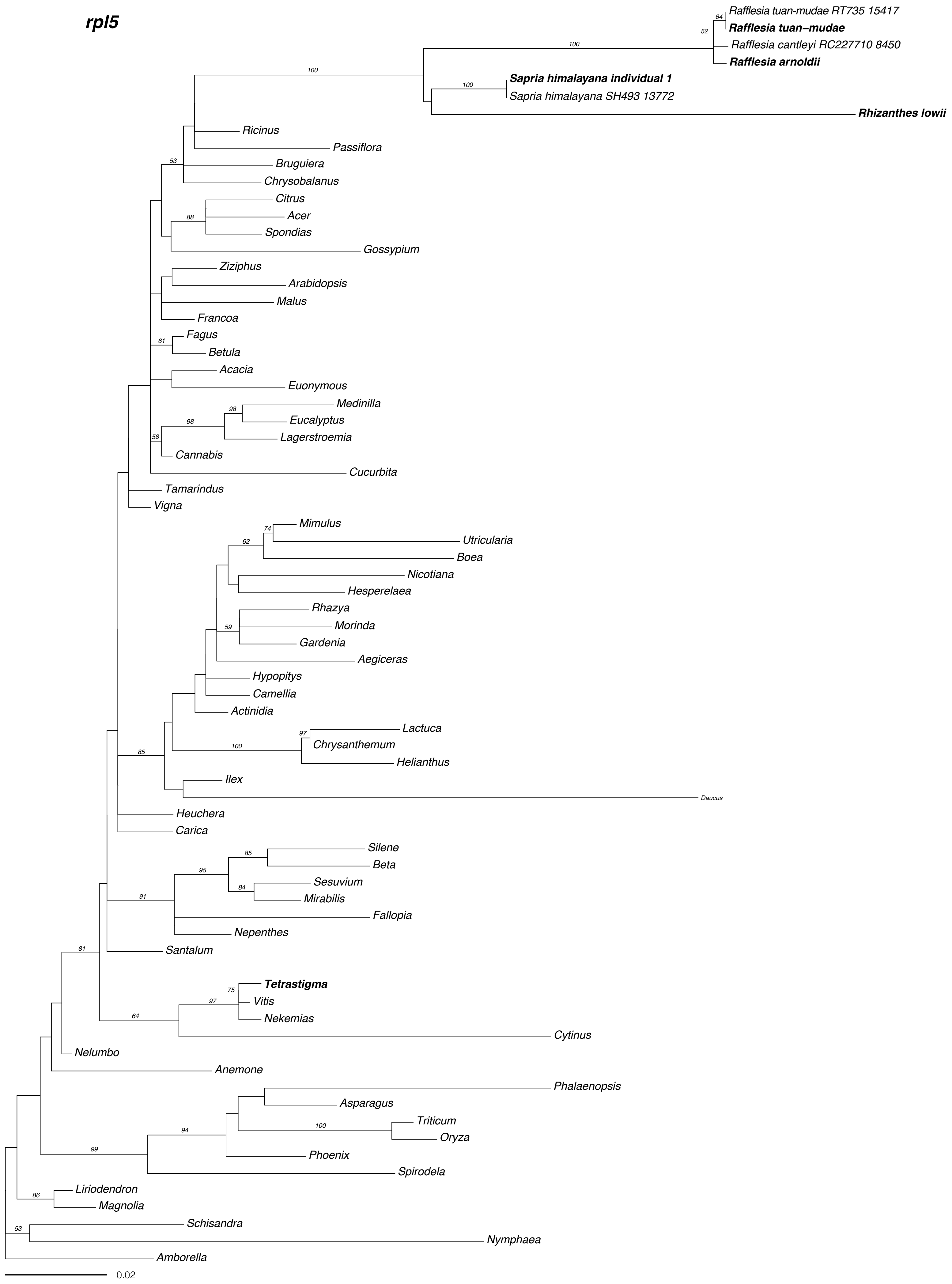
rpl5
Rafflesia tuan-mudae RT735 15417
Rafflesia tuan–mudae
Rafflesia cantleyi RC227710 8450
Rafflesia arnoldii
Sapria himalayana individual 1
Sapria himalayana SH493 13772
Rhizanthes lowii
Ricinus
Passiflora
Bruguiera
Chrysobalanus
Citrus
Acer
Spondias
Gossypium
Ziziphus
Arabidopsis
Malus
Francoa
Fagus
Betula
Acacia
Euonymous
Medinilla
Eucalyptus
Lagerstroemia
Cannabis
Cucurbita
Tamarindus
Vigna
Mimulus
Utricularia
Boea
Nicotiana
Hesperelaea
Rhazya
Morinda
Gardenia
Aegiceras
Hypopitys
Camellia
Actinidia
Lactuca
Chrysanthemum
Helianthus
Ilex
Daucus
Heuchera
Carica
Silene
Beta
Sesuvium
Mirabilis
Fallopia
Nepenthes
Santalum
Tetrastigma
Vitis
Nekemias
Cytinus
Nelumbo
Anemone
Phalaenopsis
Asparagus
Triticum
Oryza
Phoenix
Spirodela
Liriodendron
Magnolia
Schisandra
Nymphaea
Amborella
0.02

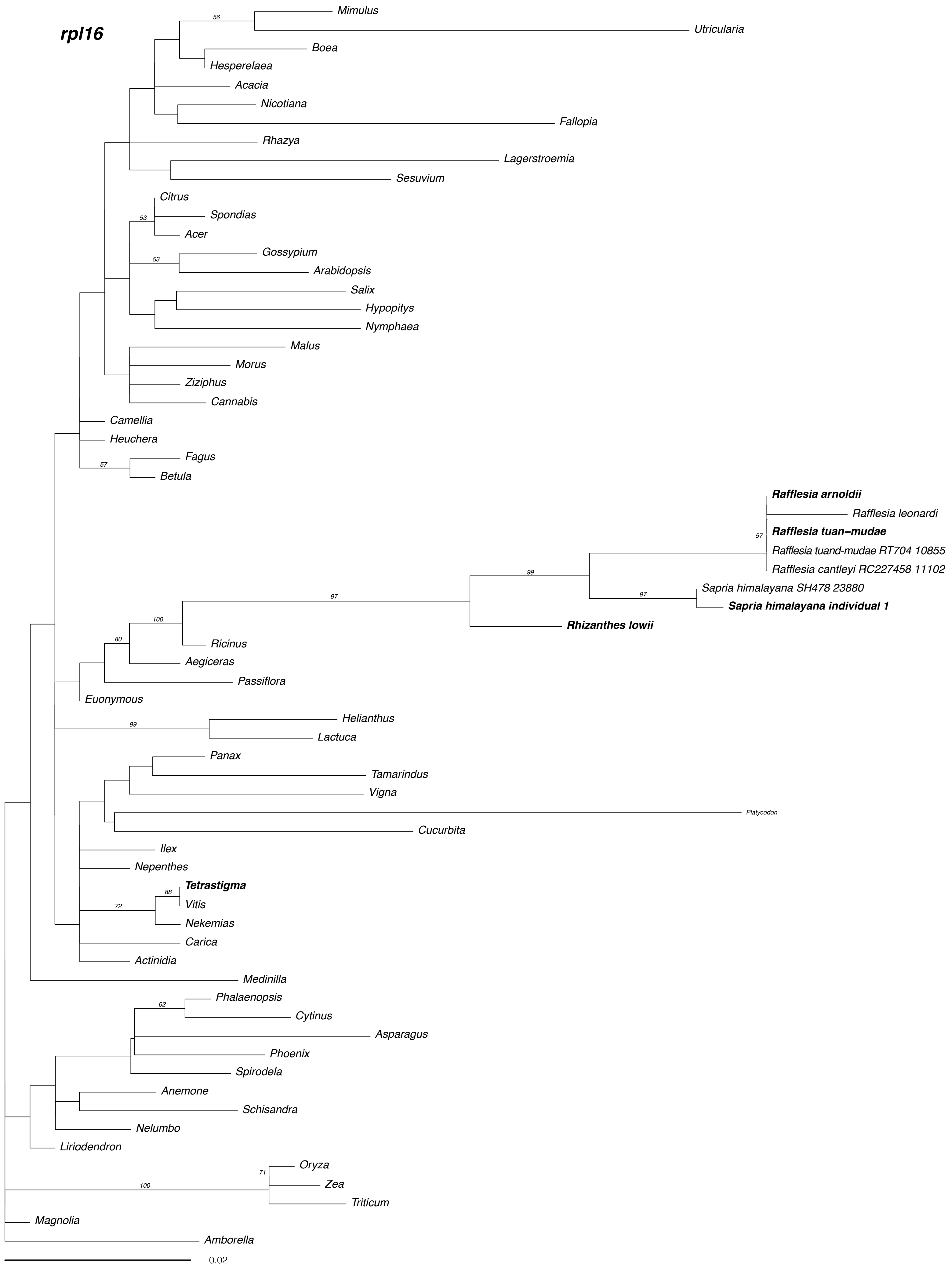
rpl16
Mimulus
Utricularia
Boea
Hesperelaea
Acacia
Nicotiana
Fallopia
Rhazya
Lagerstroemia
Sesuvium
Citrus
Spondias
Acer
Gossypium
Arabidopsis
Salix
Hypopitys
Nymphaea
Malus
Morus
Ziziphus
Cannabis
Camellia
Heuchera
Fagus
Betula
Rafflesia arnoldii
Rafflesia leonardi
Rafflesia tuan–mudae
Rafflesia tuand-mudae RT704 10855
Rafflesia cantleyi RC227458 11102
Sapria himalayana SH478 23880
Sapria himalayana individual 1
Rhizanthes lowii
Ricinus
Aegiceras
Passiflora
Euonymous
Helianthus
Lactuca
Panax
Tamarindus
Vigna
Platycodon
Cucurbita
Ilex
Nepenthes
Tetrastigma
Vitis
Nekemias
Carica
Actinidia
Medinilla
Phalaenopsis
Cytinus
Asparagus
Phoenix
Spirodela
Anemone
Schisandra
Nelumbo
Liriodendron
Oryza
Zea
Triticum
Magnolia
Amborella
0.02

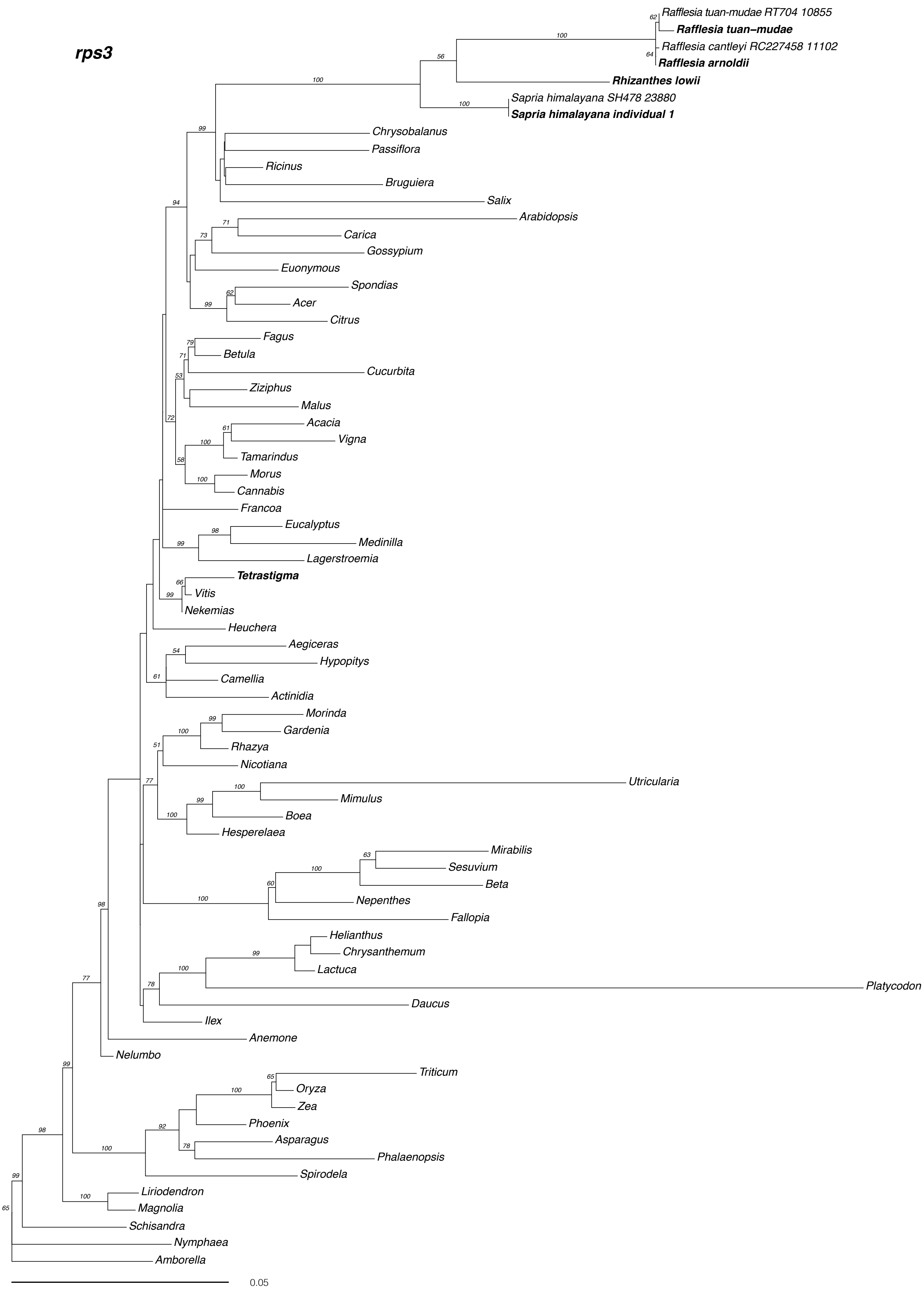

rps3
Rafflesia tuan-mudae RT704 10855
Rafflesia tuan–mudae
Rafflesia cantleyi RC227458 11102
Rafflesia arnoldii
Rhizanthes lowii
Sapria himalayana SH478 23880
Sapria himalayana individual 1
Chrysobalanus
Passiflora
Ricinus
Bruguiera
Salix
Arabidopsis
Carica
Gossypium
Euonymous
Spondias
Acer
Citrus
Fagus
Betula
Cucurbita
Ziziphus
Malus
Acacia
Vigna
Tamarindus
Morus
Cannabis
Francoa
Eucalyptus
Medinilla
Lagerstroemia
Tetrastigma
Vitis
Nekemias
Heuchera
Aegiceras
Hypopitys
Camellia
Actinidia
Morinda
Gardenia
Rhazya
Nicotiana
Utricularia
Mimulus
Boea
Hesperelaea
Mirabilis
Sesuvium
Beta
Nepenthes
Fallopia
Helianthus
Chrysanthemum
Lactuca
Platycodon
Daucus
Ilex
Anemone
Nelumbo
Triticum
Oryza
Zea
Phoenix
Asparagus
Phalaenopsis
Spirodela
Liriodendron
Magnolia
Schisandra
Nymphaea
Amborella
0.05

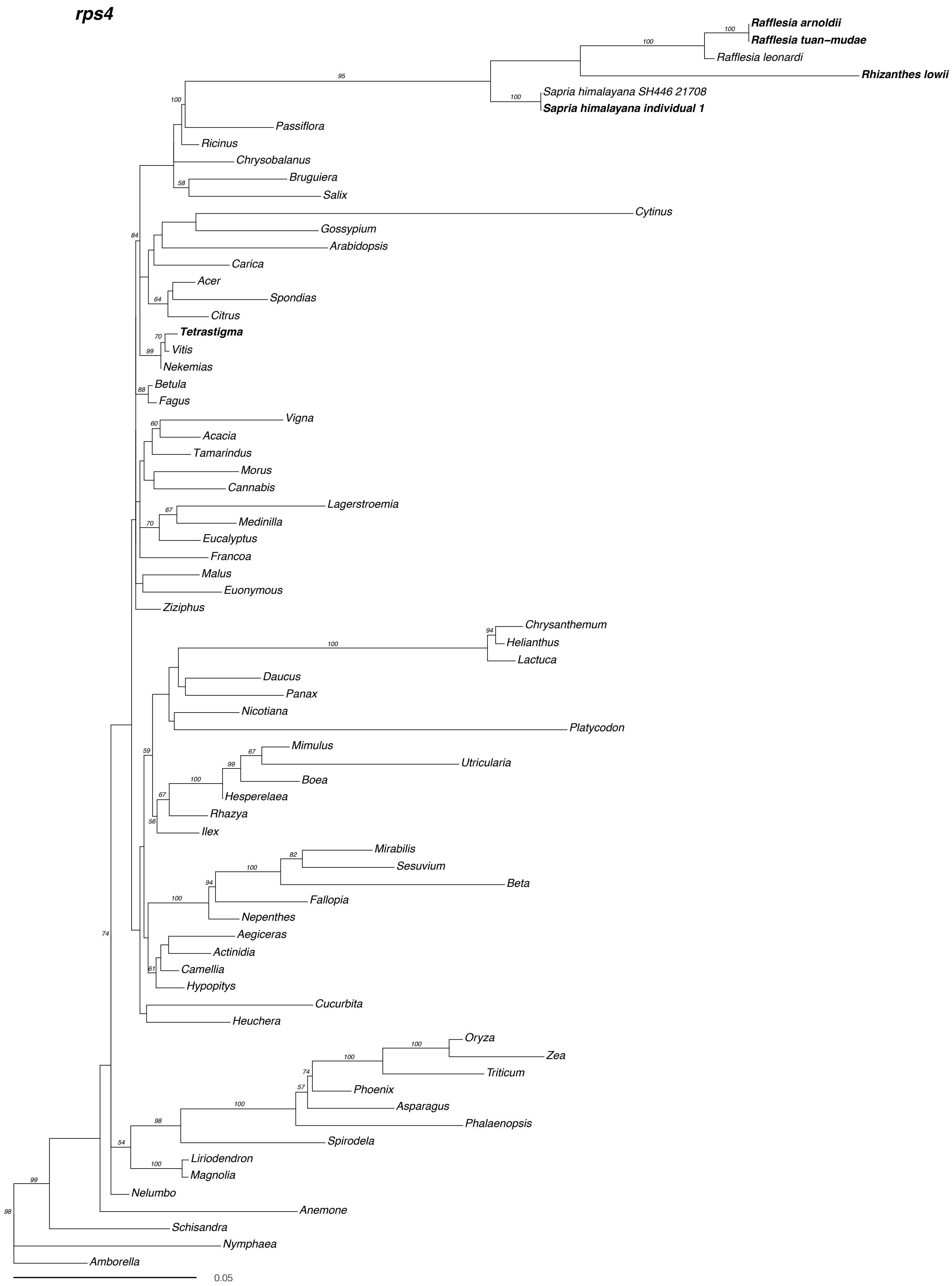

rps4
Rafflesia arnoldii
Rafflesia tuan-mudae
Rafflesia leonardi
Rhizanthes lowii
Sapria himalayana SH446 21708
Sapria himalayana individual 1
Passiflora
Ricinus
Chrysobalanus
Bruguiera
Salix
Cytinus
Gossypium
Arabidopsis
Carica
Acer
Spondias
Citrus
Tetrastigma
Vitis
Nekemias
Betula
Fagus
Vigna
Acacia
Tamarindus
Morus
Cannabis
Lagerstroemia
Medinilla
Eucalyptus
Francoa
Malus
Euonymous
Ziziphus
Chrysanthemum
Helianthus
Lactuca
Daucus
Panax
Nicotiana
Platycodon
Mimulus
Utricularia
Boea
Hesperelaea
Rhazya
Ilex
Mirabilis
Sesuvium
Beta
Fallopia
Nepenthes
Aegiceras
Actinidia
Camellia
Hypopitys
Cucurbita
Heuchera
Oryza
Zea
Triticum
Phoenix
Asparagus
Phalaenopsis
Spirodela
Liriodendron
Magnolia
Nelumbo
Anemone
Schisandra
Nymphaea
Amborella
0.05

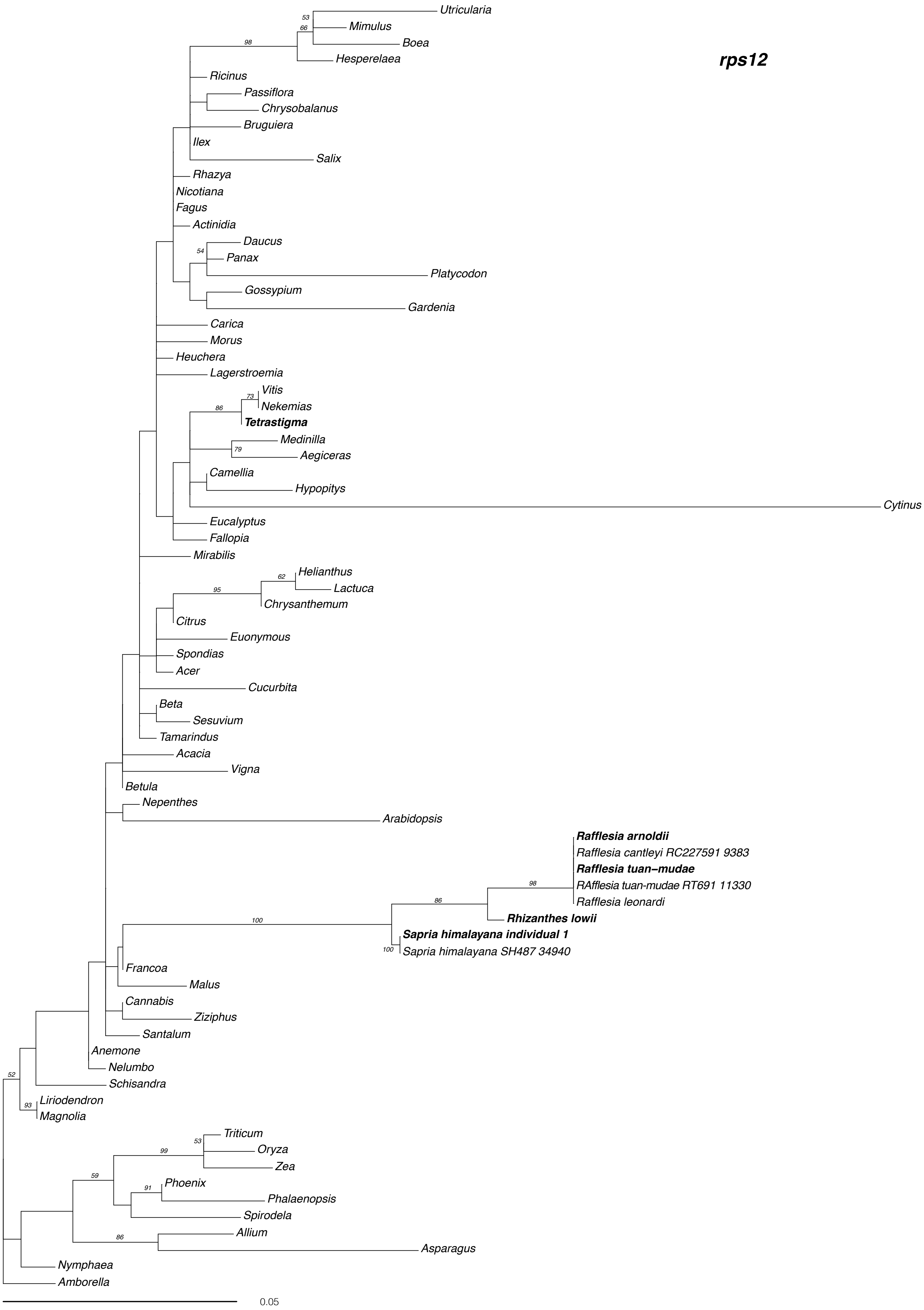
rps12
Utricularia
Mimulus
Boea
Hesperelaea
Ricinus
Passiflora
Chrysobalanus
Bruguiera
Ilex
Salix
Rhazya
Nicotiana
Fagus
Actinidia
Daucus
Panax
Platycodon
Gossypium
Gardenia
Carica
Morus
Heuchera
Lagerstroemia
Vitis
Nekemias
Tetrastigma
Medinilla
Aegiceras
Camellia
Hypopitys
Cytinus
Eucalyptus
Fallopia
Mirabilis
Helianthus
Lactuca
Chrysanthemum
Citrus
Euonymous
Spondias
Acer
Cucurbita
Beta
Sesuvium
Tamarindus
Acacia
Vigna
Betula
Nepenthes
Arabidopsis
Rafflesia arnoldii
Rafflesia cantleyi RC227591 9383
Rafflesia tuan-mudae
RAfflesia tuan-mudae RT691 11330
Rafflesia leonardi
Rhizanthes lowii
Sapria himalayana individual 1
Sapria himalayana SH487 34940
Francoa
Malus
Cannabis
Ziziphus
Santalum
Anemone
Nelumbo
Schisandra
Liriodendron
Magnolia
Triticum
Oryza
Zea
Phoenix
Phalaenopsis
Spirodela
Allium
Asparagus
Nymphaea
Amborella
0.05

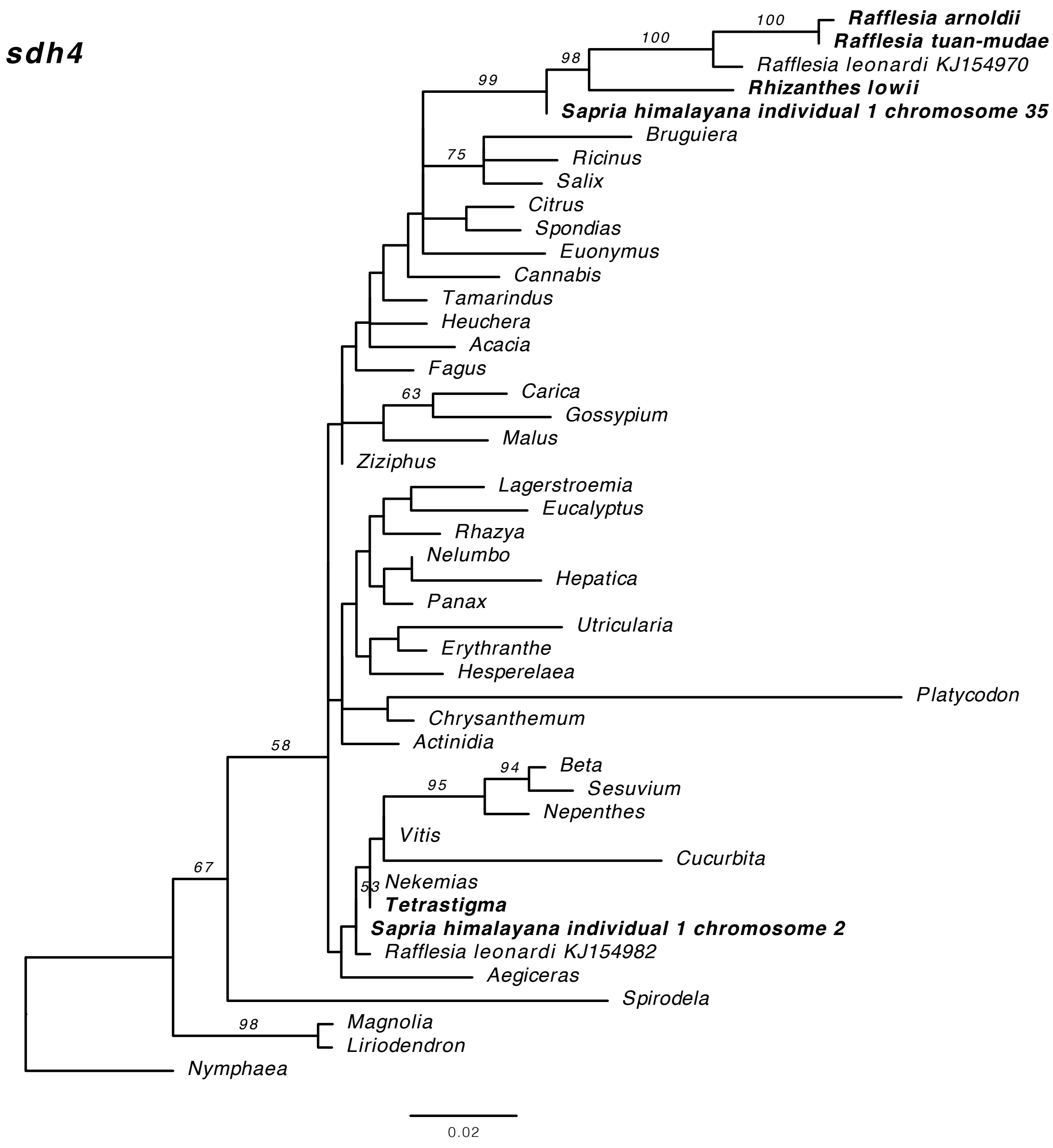


BLASTn hits against *Sapria himalayana* individual 1 Chromosome 2

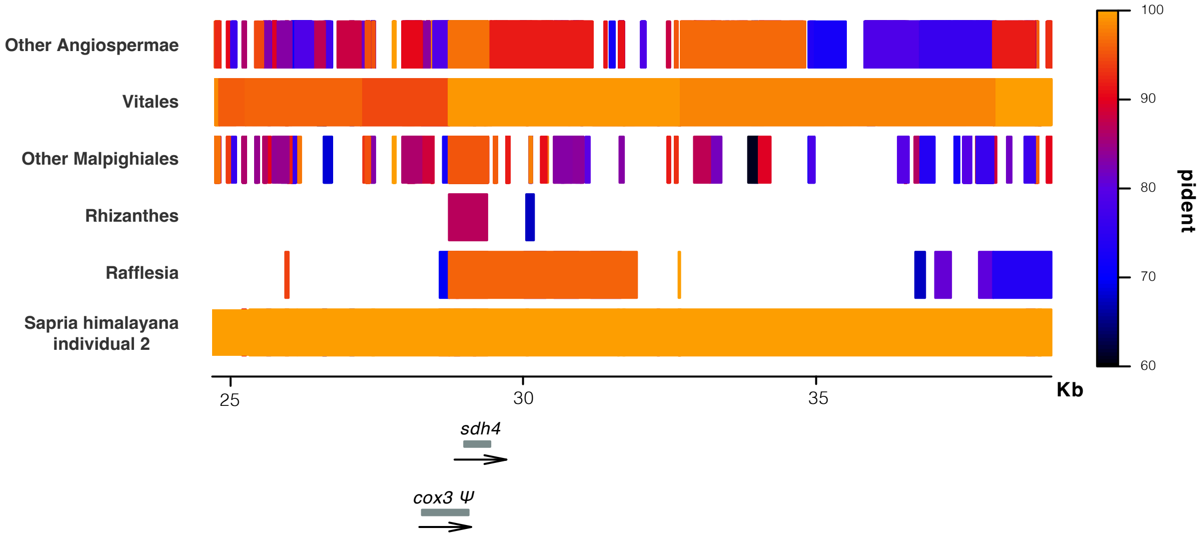

**Figure S7. Phylogenetic analyses of mitochondrial tRNA and rRNA genes of Rafflesiaceae**. Maximum likelihood analyses were performed with RAxML. ML bootstrap values >50% are shown. The scale bar corresponds to substitutions per site.

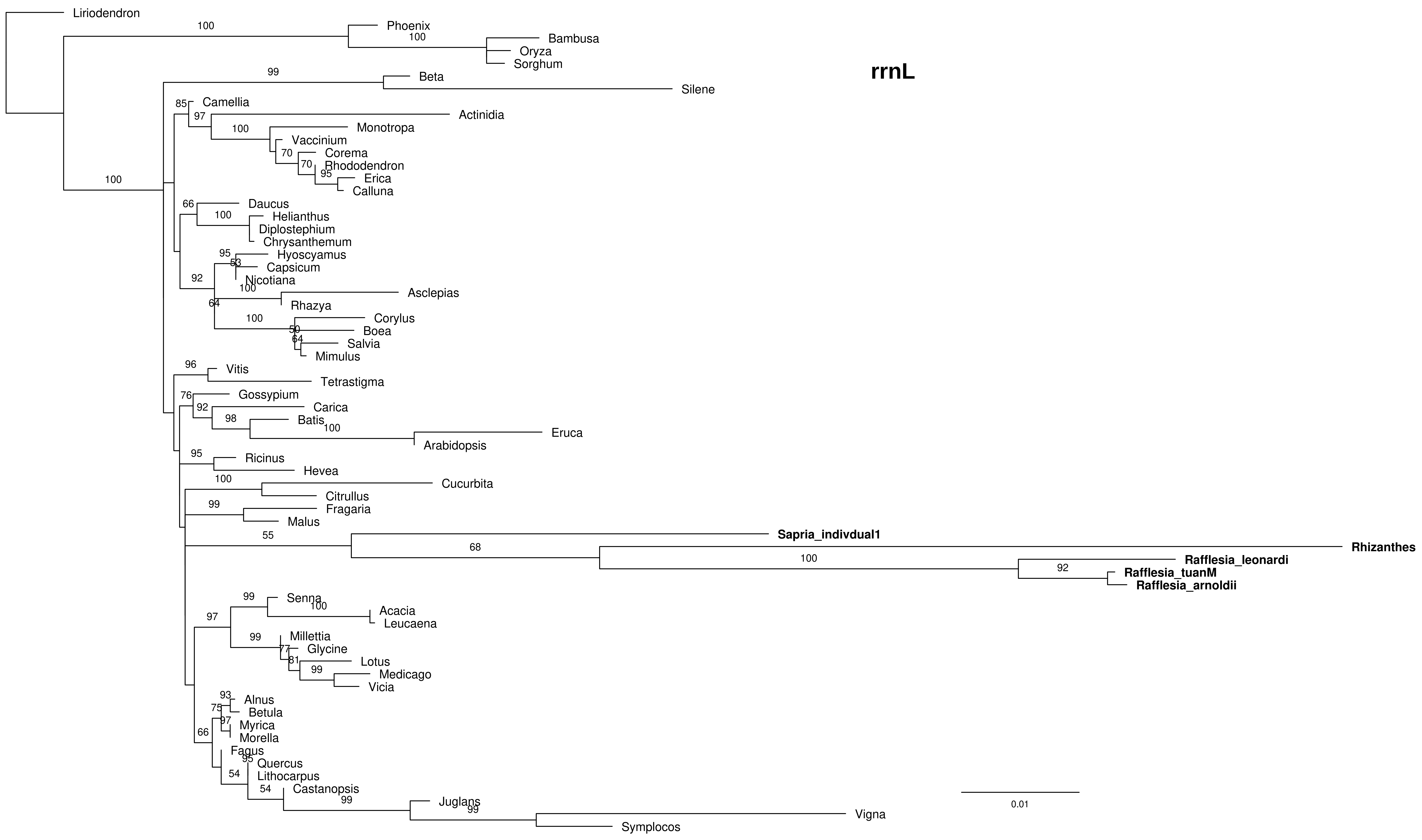

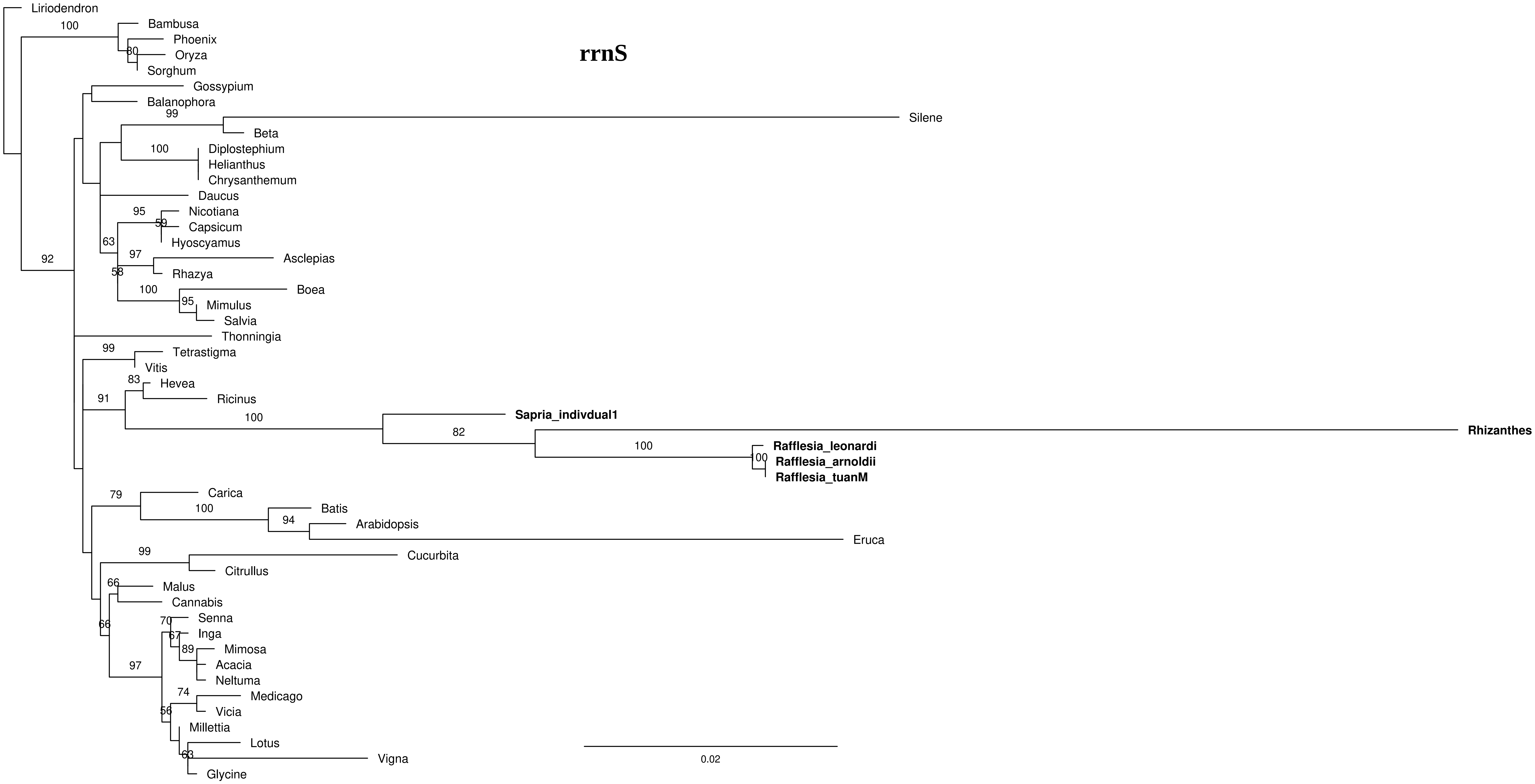
rrnS
Liriodendron
Bambusa
Phoenix
Oryza
Sorghum
Gossypium
Balanophora
Silene
Beta
Diplostephium
Helianthus
Chrysanthemum
Daucus
Nicotiana
Capsicum
Hyoscyamus
Asclepias
Rhazya
Boea
Mimulus
Salvia
Thonningia
Tetrastigma
Vitis
Hevea
Ricinus
Sapria_indivdual1
Rhizanthes
Rafflesia_leonardi
Rafflesia_arnoldii
Rafflesia_tuanM
Carica
Batis
Arabidopsis
Eruca
Cucurbita
Citrullus
Malus
Cannabis
Senna
Inga
Mimosa
Acacia
Neltuma
Medicago
Vicia
Millettia
Lotus
Vigna
Glycine
0.02

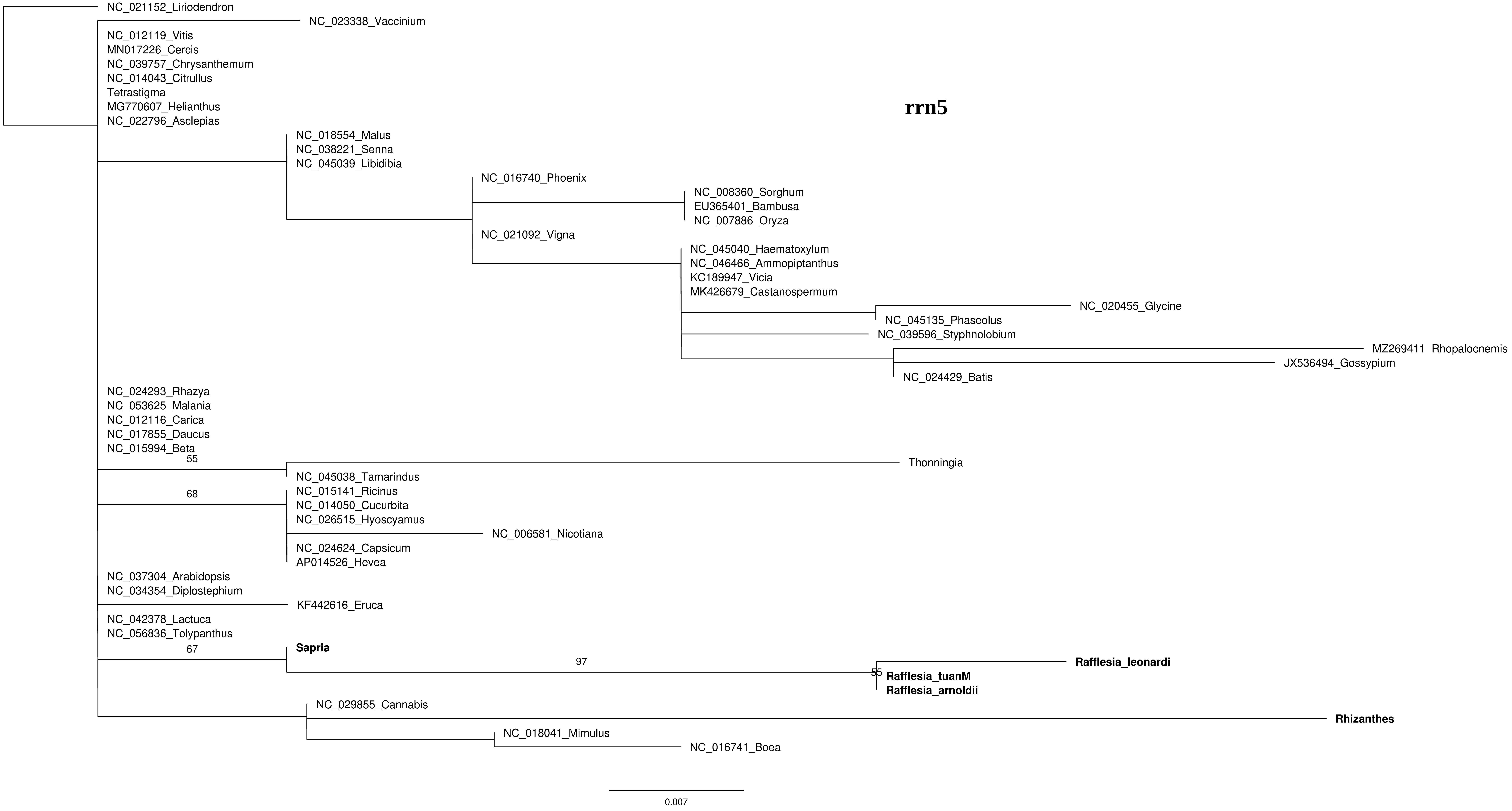
rrn5
NC_021152_Liriodendron
NC_023338_Vaccinium
NC_012119_Vitis
MN017226_Cercis
NC_039757_Chrysanthemum
NC_014043_Citrullus
Tetrastigma
MG770607_Helianthus
NC_022796_Asclepias
NC_018554_Malus
NC_038221_Senna
NC_045039_Libidibia
NC_016740_Phoenix
NC_008360_Sorghum
EU365401_Bambusa
NC_007886_Oryza
NC_021092_Vigna
NC_045040_Haematoxylum
NC_046466_Ammopiptanthus
KC189947_Vicia
MK426679_Castanospermum
NC_020455_Glycine
NC_045135_Phaseolus
NC_039596_Styphnolobium
MZ269411_Rhopalocnemis
JX536494_Gossypium
NC_024429_Batis
NC_024293_Rhazya
NC_053625_Malania
NC_012116_Carica
NC_017855_Daucus
NC_015994_Beta
55
Thonningia
NC_045038_Tamarindus
68
NC_015141_Ricinus
NC_014050_Cucurbita
NC_026515_Hyoscyamus
NC_006581_Nicotiana
NC_024624_Capsicum
AP014526_Hevea
NC_037304_Arabidopsis
NC_034354_Diplostephium
KF442616_Eruca
NC_042378_Lactuca
NC_056836_Tolypanthus
67
Sapria
97
Rafflesia_leonardi
55
Rafflesia_tuanM
Rafflesia_arnoldii
NC_029855_Cannabis
Rhizanthes
NC_018041_Mimulus
NC_016741_Boea
0.007

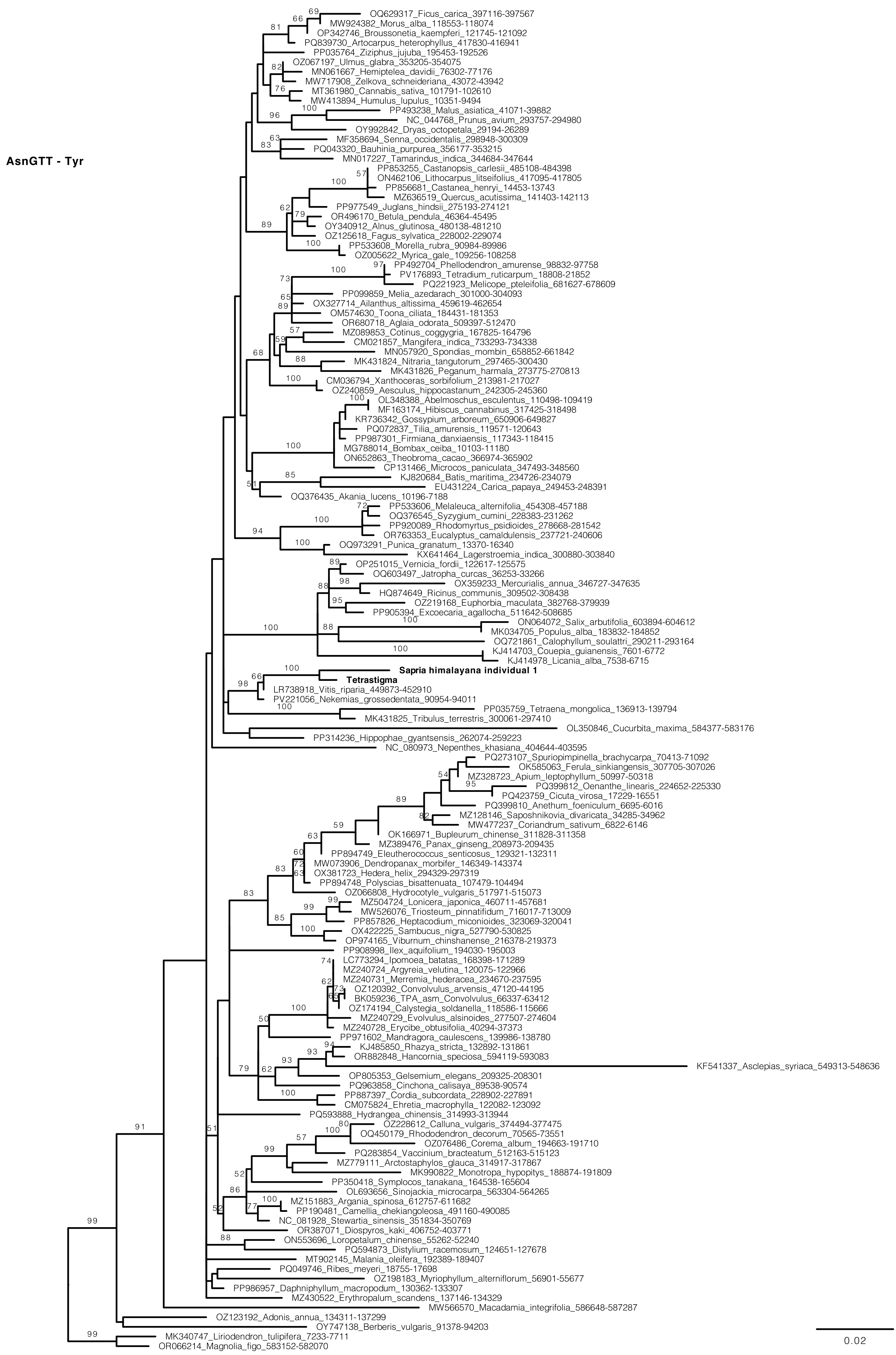
AsnGTT - Tyr
OQ629317_Ficus_carica_397116-397567
MW924382_Morus_alba_118553-118074
OP342746_Broussonetia_kaempferi_121745-121092
PQ839730_Artocarpus_heterophyllus_417830-416941
PP035764_Ziziphus_jujuba_195453-192526
OZ067197_Ulmus_glabra_353205-354075
MN061667_Hemiptelea_davidii_76302-77176
MW717908_Zelkova_schneideriana_43072-43942
MT361980_Cannabis_sativa_101791-102610
MW413894_Humulus_lupulus_10351-9494
PP493238_Malus_asiatica_41071-39882
NC_044768_Prunus_avium_293757-294980
OY992842_Dryas_octopetala_29194-26289
MF358694_Senna_occidentalis_298948-300309
PQ043320_Bauhinia_purpurea_356177-353215
MN017227_Tamarindus_indica_344684-347644
PP853255_Castanopsis_carlesii_485108-484398
ON462106_Lithocarpus_litseifolius_417095-417805
PP856681_Castanea_henryi_14453-13743
MZ636519_Quercus_acutissima_141403-142113
PP977549_Juglans_hindsii_275193-274121
OR496170_Betula_pendula_46364-45495
OY340912_Alnus_glutinosa_480138-481210
OZ125618_Fagus_sylvatica_228002-229074
PP533608_Morella_rubra_90984-89986
OZ005622_Myrica_gale_109256-108258
PP492704_Phellodendron_amurense_98832-97758
PV176893_Tetradium_ruticarpum_18808-21852
PQ221923_Melicope_pteleifolia_681627-678609
PP099859_Melia_azedarach_301000-304093
OX327714_Ailanthus_altissima_459619-462654
OM574630_Toona_ciliata_184431-181353
OR680718_Aglaia_odorata_509397-512470
MZ089853_Cotinus_coggygria_167825-164796
CM021857_Mangifera_indica_733293-734338
MN057920_Spondias_mombin_658852-661842
MK431824_Nitraria_tangutorum_297465-300430
MK431826_Peganum_harmala_273775-270813
CM036794_Xanthoceras_sorbifolium_213981-217027
OZ240859_Aesculus_hippocastanum_242305-245360
OL348388_Abelmoschus_esculentus_110498-109419
MF163174_Hibiscus_cannabinus_317425-318498
KR736342_Gossypium_arboreum_650906-649827
PQ072837_Tilia_amurensis_119571-120643
PP987301_Firmiana_danxiaensis_117343-118415
MG788014_Bombax_ceiba_10103-11180
ON652863_Theobroma_cacao_366974-365902
CP131466_Microcos_paniculata_347493-348560
KJ820684_Batis_maritima_234726-234079
EU431224_Carica_papaya_249453-248391
OQ376435_Akania_lucens_10196-7188
PP533606_Melaleuca_alternifolia_454308-457188
OQ376545_Syzygium_cumini_228383-231262
PP920089_Rhodomyrtus_psidioides_278668-281542
OR763353_Eucalyptus_camaldulensis_237721-240606
OQ973291_Punica_granatum_13370-16340
KX641464_Lagerstroemia_indica_300880-303840
OP251015_Vernicia_fordii_122617-125575
OQ603497_Jatropha_curcas_36253-33266
OX359233_Mercurialis_annua_346727-347635
HQ874649_Ricinus_communis_309502-308438
OZ219168_Euphorbia_maculata_382768-379939
PP905394_Excoecaria_agallocha_511642-508685
ON064072_Salix_arbutifolia_603894-604612
MK034705_Populus_alba_183832-184852
OQ721861_Calophyllum_soulattri_290211-293164
KJ414703_Couepia_guianensis_7601-6772
KJ414978_Licania_alba_7538-6715
Sapria himalayana individual 1
Tetrastigma
LR738918_Vitis_riparia_449873-452910
PV221056_Nekemias_grossedentata_90954-94011
PP035759_Tetraena_mongolica_136913-139794
MK431825_Tribulus_terrestris_300061-297410
OL350846_Cucurbita_maxima_584377-583176
PP314236_Hippophae_gyantsensis_262074-259223
NC_080973_Nepenthes_khasiana_404644-403595
PQ273107_Spuriopimpinella_brachycarpa_70413-71092
OK585063_Ferula_sinkiangensis_307705-307026
MZ328723_Apium_leptophyllum_50997-50318
PQ399812_Oenanthe_linearis_224652-225330
PQ423759_Cicuta_virosa_17229-16551
PQ399810_Anethum_foeniculum_6695-6016
MZ128146_Saposhnikovia_divaricata_34285-34962
MW477237_Coriandrum_sativum_6822-6146
OK166971_Bupleurum_chinense_311828-311358
MZ389476_Panax_ginseng_208973-209435
PP894749_Eleutherococcus_senticosus_129321-132311
MW073906_Dendropanax_morbifer_146349-143374
OX381723_Hedera_helix_294329-297319
PP894748_Polyscias_bisattenuata_107479-104494
OZ066808_Hydrocotyle_vulgaris_517971-515073
MZ504724_Lonicera_japonica_460711-457681
MW526076_Triosteum_pinnatifidum_716017-713009
PP857826_Heptacodium_miconioides_323069-320041
OX422225_Sambucus_nigra_527790-530825
OP974165_Viburnum_chinshanense_216378-219373
PP908998_Ilex_aquifolium_194030-195003
LC773294_Ipomoea_batatas_168398-171289
MZ240724_Argyreia_velutina_120075-122966
MZ240731_Merremia_hederacea_234670-237595
OZ120392_Convolvulus_arvensis_47120-44195
BK059236_TPA_asm_Convolvulus_66337-63412
OZ174194_Calystegia_soldanella_118586-115666
MZ240729_Evolvulus_alsinoides_277507-274604
MZ240728_Erycibe_obtusifolia_40294-37373
PP971602_Mandragora_caulescens_139986-138780
KJ485850_Rhazya_stricta_132892-131861
OR882848_Hancornia_speciosa_594119-593083
KF541337_Asclepias_syriaca_549313-548636
OP805353_Gelsemium_elegans_209325-208301
PQ963858_Cinchona_calisaya_89538-90574
PP887397_Cordia_subcordata_228902-227891
CM075824_Ehretia_macrophylla_122082-123092
PQ593888_Hydrangea_chinensis_314993-313944
OZ228612_Calluna_vulgaris_374494-377475
OQ450179_Rhododendron_decorum_70565-73551
OZ076486_Corema_album_194663-191710
PQ283854_Vaccinium_bracteatum_512163-515123
MZ779111_Arctostaphylos_glauca_314917-317867
MK990822_Monotropa_hypopitys_188874-191809
PP350418_Symplocos_tanakana_164538-165604
OL693656_Sinojackia_microcarpa_563304-564265
MZ151883_Argania_spinosa_612757-611682
PP190481_Camellia_chekiangoleosa_491160-490085
NC_081928_Stewartia_sinensis_351834-350769
OR387071_Diospyros_kaki_406752-403771
ON553696_Loropetalum_chinense_55262-52240
PQ594873_Distylium_racemosum_124651-127678
MT902145_Malania_oleifera_192389-189407
PQ049746_Ribes_meyeri_18755-17698
OZ198183_Myriophyllum_alterniflorum_56901-55677
PP986957_Daphniphyllum_macropodum_130362-133307
MZ430522_Erythropalum_scandens_137146-134329
MW566570_Macadamia_integrifolia_586648-587287
OZ123192_Adonis_annua_134311-137299
OY747138_Berberis_vulgaris_91378-94203
MK340747_Liriodendron_tulipifera_7233-7711
OR066214_Magnolia_figo_583152-582070
0.02

**GluTTC**

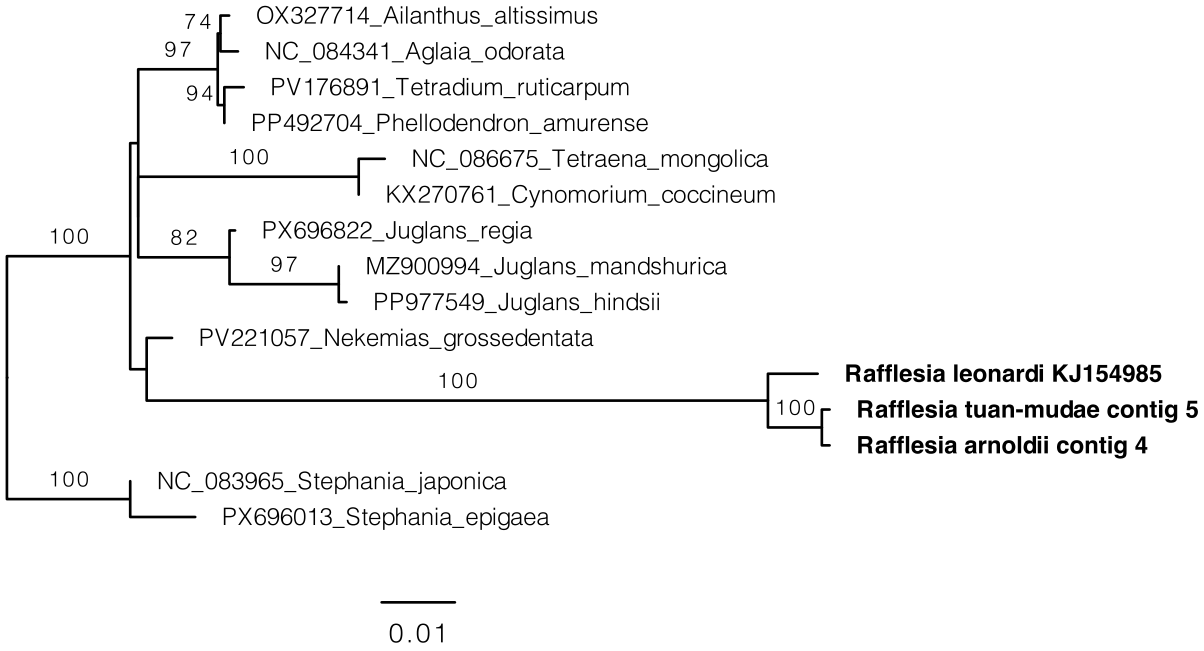
74
OX327714_Ailanthus_altissimus
97
NC_084341_Aglaia_odorata
94
PV176891_Tetradium_ruticarpum
PP492704_Phellodendron_amurense
100
NC_086675_Tetraena_mongolica
KX270761_Cynomorium_coccineum
100
82
PX696822_Juglans_regia
97
MZ900994_Juglans_mandshurica
PP977549_Juglans_hindsii
PV221057_Nekemias_grossedentata
100
Rafflesia leonardi KJ154985
100
Rafflesia tuan-mudae contig 5
Rafflesia arnoldii contig 4
100
NC_083965_Stephania_japonica
PX696013_Stephania_epigaea
0.01

**GluTTC**

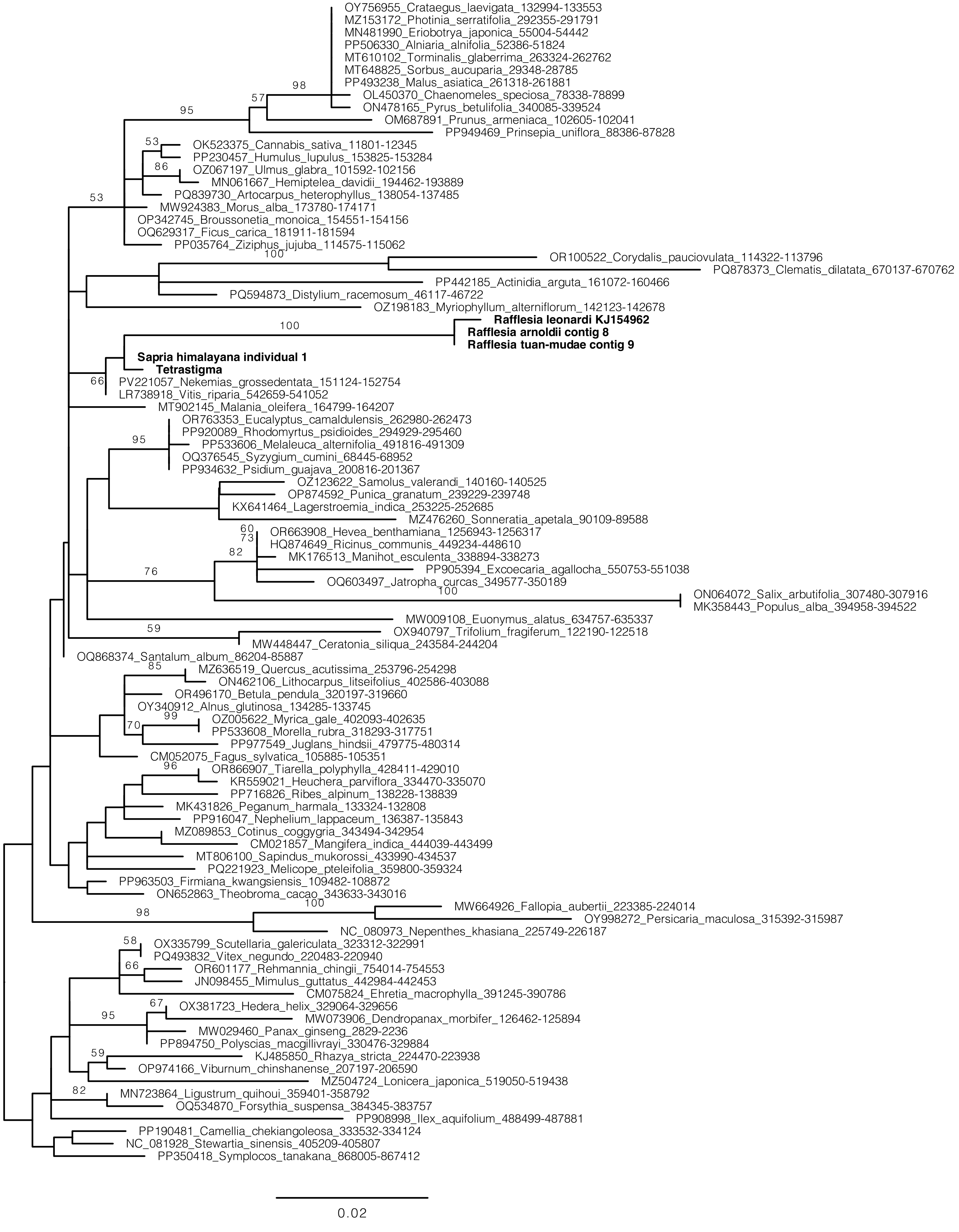

OY756955_Crataegus_laevigata_132994-133553
MZ153172_Photinia_serratifolia_292355-291791
MN481990_Eriobotrya_japonica_55004-54442
PP506330_Alniaria_alnifolia_52386-51824
MT610102_Torminalis_glaberrima_263324-262762
MT648825_Sorbus_aucuparia_29348-28785
PP493238_Malus_asiatica_261318-261881
OL450370_Chaenomeles_speciosa_78338-78899
ON478165_Pyrus_betulifolia_340085-339524
OM687891_Prunus_armeniaca_102605-102041
PP949469_Prinsepia_uniflora_88386-87828
OK523375_Cannabis_sativa_11801-12345
PP230457_Humulus_lupulus_153825-153284
OZ067197_Ulmus_glabra_101592-102156
MN061667_Hemiptelea_davidii_194462-193889
PQ839730_Artocarpus_heterophyllus_138054-137485
MW924383_Morus_alba_173780-174171
OP342745_Broussonetia_monoica_154551-154156
OQ629317_Ficus_carica_181911-181594
PP035764_Ziziphus_jujuba_114575-115062
OR100522_Corydalis_pauciovulata_114322-113796
PQ878373_Clematis_dilatata_670137-670762
PP442185_Actinidia_arguta_161072-160466
PQ594873_Distylium_racemosum_46117-46722
OZ198183_Myriophyllum_alterniflorum_142123-142678
Rafflesia leonardi KJ154962
Rafflesia arnoldii contig 8
Rafflesia tuan-mudae contig 9
Sapria himalayana individual 1
Tetrastigma
PV221057_Nekemias_grossedentata_151124-152754
LR738918_Vitis_riparia_542659-541052
MT902145_Malania_oleifera_164799-164207
OR763353_Eucalyptus_camaldulensis_262980-262473
PP920089_Rhodomyrtus_psidioides_294929-295460
PP533606_Melaleuca_alternifolia_491816-491309
OQ376545_Syzygium_cumini_68445-68952
PP934632_Psidium_guajava_200816-201367
OZ123622_Samolus_valerandi_140160-140525
OP874592_Punica_granatum_239229-239748
KX641464_Lagerstroemia_indica_253225-252685
MZ476260_Sonneratia_apetala_90109-89588
OR663908_Hevea_benthamiana_1256943-1256317
HQ874649_Ricinus_communis_449234-448610
MK176513_Manihot_esculenta_338894-338273
PP905394_Excoecaria_agallocha_550753-551038
OQ603497_Jatropha_curcas_349577-350189
ON064072_Salix_arbutifolia_307480-307916
MK358443_Populus_alba_394958-394522
MW009108_Euonymus_alatus_634757-635337
OX940797_Trifolium_fragiferum_122190-122518
MW448447_Ceratonia_siliqua_243584-244204
OQ868374_Santalum_album_86204-85887
MZ636519_Quercus_acutissima_253796-254298
ON462106_Lithocarpus_litseifolius_402586-403088
OR496170_Betula_pendula_320197-319660
OY340912_Alnus_glutinosa_134285-133745
OZ005622_Myrica_gale_402093-402635
PP533608_Morella_rubra_318293-317751
PP977549_Juglans_hindsii_479775-480314
CM052075_Fagus_sylvatica_105885-105351
OR866907_Tiarella_polyphylla_428411-429010
KR559021_Heuchera_parviflora_334470-335070
PP716826_Ribes_alpinum_138228-138839
MK431826_Peganum_harmala_133324-132808
PP916047_Nephelium_lappaceum_136387-135843
MZ089853_Cotinus_coggygria_343494-342954
CM021857_Mangifera_indica_444039-443499
MT806100_Sapindus_mukorossi_433990-434537
PQ221923_Melicope_pteleifolia_359800-359324
PP963503_Firmiana_kwangsiensis_109482-108872
ON652863_Theobroma_cacao_343633-343016
MW664926_Fallopia_aubertii_223385-224014
OY998272_Persicaria_maculosa_315392-315987
NC_080973_Nepenthes_khasiana_225749-226187
OX335799_Scutellaria_galericulata_323312-322991
PQ493832_Vitex_negundo_220483-220940
OR601177_Rehmannia_chingii_754014-754553
JN098455_Mimulus_guttatus_442984-442453
CM075824_Ehretia_macrophylla_391245-390786
OX381723_Hedera_helix_329064-329656
MW073906_Dendropanax_morbifer_126462-125894
MW029460_Panax_ginseng_2829-2236
PP894750_Polyscias_macgillivrayi_330476-329884
KJ485850_Rhazya_stricta_224470-223938
OP974166_Viburnum_chinshanense_207197-206590
MZ504724_Lonicera_japonica_519050-519438
MN723864_Ligustrum_quihoui_359401-358792
OQ534870_Forsythia_suspensa_384345-383757
PP908998_Ilex_aquifolium_488499-487881
PP190481_Camellia_chekiangoleosa_333532-334124
NC_081928_Stewartia_sinensis_405209-405807
PP350418_Symplocos_tanakana_868005-867412
0.02

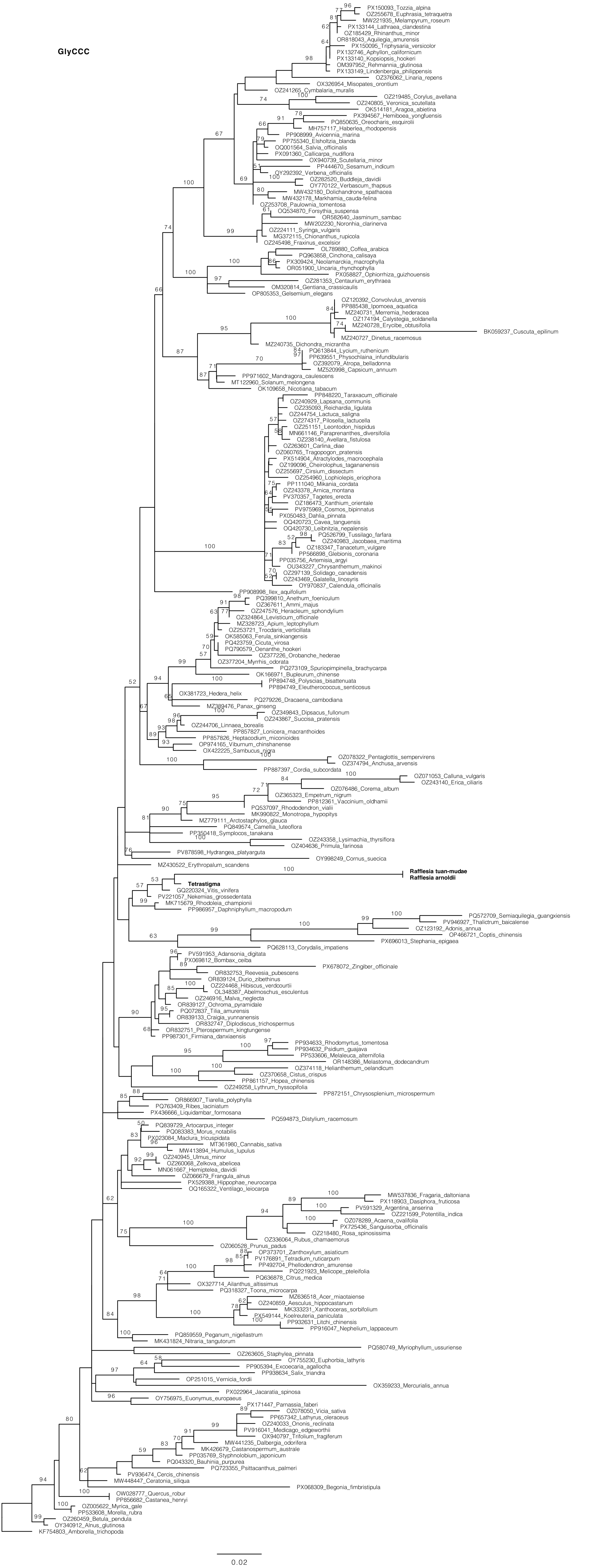
GlyCCC
PX150093_Tozzia_alpina
OZ255678_Euphrasia_tetraquetra
MW221935_Melampyrum_roseum
PX133144_Lathraea_clandestina
OZ185429_Rhinanthus_minor
OR818043_Aquilegia_amurensis
PX150095_Triphysaria_versicolor
PX132746_Aphyllon_californicum
PX133140_Kopsiopsis_hookeri
OM397952_Rehmannia_glutinosa
PX133149_Lindenbergia_philippensis
OZ376062_Linaria_repens
OX326954_Misopates_orontium
OZ241265_Cymbalaria_muralis
OZ219485_Corylus_avellana
OZ240805_Veronica_scutellata
OK514181_Aragoa_abietina
PX394567_Hemiboea_yongfuensis
PQ850635_Oreocharis_esquirolii
MH757117_Haberlea_rhodopensis
PP908999_Avicennia_marina
PP755340_Elsholtzia_blanda
OQ001564_Salvia_officinalis
PX091360_Callicarpa_nudiflora
OX940739_Scutellaria_minor
PP444670_Sesamum_indicum
OY292392_Verbena_officinalis
OZ282520_Buddleja_davidii
OY770122_Verbascum_thapsus
MW432180_Dolichandrone_spathacea
MW432178_Markhamia_cauda-felina
OZ253708_Paulownia_tomentosa
OQ534870_Forsythia_suspensa
OR582640_Jasminum_sambac
MW202230_Noronhia_clarinerva
OZ224111_Syringa_vulgaris
MG372115_Chionanthus_rupicola
OZ245498_Fraxinus_excelsior
OL789880_Coffea_arabica
PQ963858_Cinchona_calisaya
PX309424_Neolamarckia_macrophylla
OR051900_Uncaria_rhynchophylla
PX058827_Ophiorrhiza_guizhouensis
OZ281353_Centaurium_erythraea
OM320814_Gentiana_crassicaulis
OP805353_Gelsemium_elegans
OZ120392_Convolvulus_arvensis
PP885438_Ipomoea_aquatica
MZ240731_Merremia_hederacea
OZ174194_Calystegia_soldanella
MZ240728_Erycibe_obtusifolia
BK059237_Cuscuta_epilinum
MZ240727_Dinetus_racemosus
MZ240735_Dichondra_micrantha
PQ613844_Lycium_ruthenicum
PP639551_Physochlaina_infundibularis
OZ392079_Atropa_belladonna
MZ520998_Capsicum_annuum
PP971602_Mandragora_caulescens
MT122960_Solanum_melongena
OK109658_Nicotiana_tabacum
PP848220_Taraxacum_officinale
OZ240929_Lapsana_communis
OZ235093_Reichardia_ligulata
OZ244754_Lactuca_saligna
OZ274317_Pilosella_lactucella
OZ251151_Leontodon_hispidus
MN661146_Paraprenanthes_diversifolia
OZ238140_Avellara_fistulosa
OZ263601_Carlina_diae
OZ060765_Tragopogon_pratensis
PX514904_Atractylodes_macrocephala
OZ199096_Cheirolophus_tagananensis
OZ255697_Cirsium_dissectum
OZ254960_Lophiolepis_eriophora
PP111040_Mikania_cordata
OZ243378_Arnica_montana
PV370357_Tagetes_erecta
OZ186473_Xanthium_orientale
PV975969_Cosmos_bipinnatus
PX050483_Dahlia_pinnata
OQ420723_Cavea_tanguensis
OQ420730_Leibnitzia_nepalensis
PQ526799_Tussilago_farfara
OZ240983_Jacobaea_maritima
OZ183347_Tanacetum_vulgare
PP566898_Glebionis_coronaria
PP035756_Artemisia_argyi
OU343227_Chrysanthemum_makinoi
OZ297139_Solidago_canadensis
OZ243469_Galatella_linosyris
OY970837_Calendula_officinalis
PP908998_Ilex_aquifolium
PQ399810_Anethum_foeniculum
OZ367611_Ammi_majus
OZ247576_Heracleum_sphondylium
OZ324864_Levisticum_officinale
MZ328723_Apium_leptophyllum
OZ253721_Trocdaris_verticillata
OK585063_Ferula_sinkiangensis
PQ423759_Cicuta_virosa
PQ790579_Oenanthe_hookeri
OZ377226_Orobanche_hederae
OZ377204_Myrrhis_odorata
PQ273109_Spuriopimpinella_brachycarpa
OK166971_Bupleurum_chinense
PP894748_Polyscias_bisattenuata
PP894749_Eleutherococcus_senticosus
OX381723_Hedera_helix
PQ279226_Dracaena_cambodiana
MZ389476_Panax_ginseng
OZ349843_Dipsacus_fullonum
OZ243867_Succisa_pratensis
OZ244706_Linnaea_borealis
PP857827_Lonicera_macranthoides
PP857826_Heptacodium_miconioides
OP974165_Viburnum_chinshanense
OX422225_Sambucus_nigra
OZ078322_Pentaglottis_sempervirens
OZ374794_Anchusa_arvensis
PP887397_Cordia_subcordata
OZ071053_Calluna_vulgaris
OZ243140_Erica_ciliaris
OZ076486_Corema_album
OZ365323_Empetrum_nigrum
PP812361_Vaccinium_oldhamii
PQ537097_Rhododendron_vialii
MK990822_Monotropa_hypopitys
MZ779111_Arctostaphylos_glauca
PQ849574_Camellia_luteoflora
PP350418_Symplocos_tanakana
OZ243358_Lysimachia_thyrsiflora
OZ404636_Primula_farinosa
PV878598_Hydrangea_platyarguta
OY998249_Cornus_suecica
MZ430522_Erythropalum_scandens
Rafflesia tuan-mudae
Rafflesia arnoldii
Tetrastigma
GQ220324_Vitis_vinifera
PV221057_Nekemias_grossedentata
MK715679_Rhodoleia_championii
PP986957_Daphniphyllum_macropodum
PQ572709_Semiaquilegia_guangxiensis
PV946927_Thalictrum_baicalense
OZ123192_Adonis_annua
OP466721_Coptis_chinensis
PX696013_Stephania_epigaea
PQ628113_Corydalis_impatiens
PV591953_Adansonia_digitata
PX069812_Bombax_ceiba
OR832753_Reevesia_pubescens
PX678072_Zingiber_officinale
OR839124_Durio_zibethinus
OZ224468_Hibiscus_verdcourtii
OL348387_Abelmoschus_esculentus
OZ246916_Malva_neglecta
OR839127_Ochroma_pyramidale
PQ072837_Tilia_amurensis
OR839133_Craigia_yunnanensis
OR832747_Diplodiscus_trichospermus
OR832751_Pterospermum_kingtungense
PP987301_Firmiana_danxiaensis
PP934633_Rhodomyrtus_tomentosa
PP934632_Psidium_guajava
PP533606_Melaleuca_alternifolia
OR148386_Melastoma_dodecandrum
OZ374118_Helianthemum_oelandicum
OZ370658_Cistus_crispus
PP861157_Hopea_chinensis
OZ249258_Lythrum_hyssopifolia
PP872151_Chrysosplenium_microspermum
OR866907_Tiarella_polyphylla
PQ763409_Ribes_laciniatum
PX436666_Liquidambar_formosana
PQ594873_Distylium_racemosum
PQ839729_Artocarpus_integer
PQ083383_Morus_notabilis
PX023084_Maclura_tricuspidata
MT361980_Cannabis_sativa
MW413894_Humulus_lupulus
OZ240945_Ulmus_minor
OZ260068_Zelkova_abelicea
MN061667_Hemiptelea_davidii
OZ066679_Frangula_alnus
PX529388_Hippophae_neurocarpa
OQ165322_Ventilago_leiocarpa
MW537836_Fragaria_daltoniana
PX118903_Dasiphora_fruticosa
PV591329_Argentina_anserina
OZ221599_Potentilla_indica
OZ078289_Acaena_ovalifolia
PX725436_Sanguisorba_officinalis
OZ218480_Rosa_spinosissima
OZ336064_Rubus_chamaemorus
OZ060528_Prunus_padus
OP373701_Zanthoxylum_asiaticum
PV176891_Tetradium_ruticarpum
PP492704_Phellodendron_amurense
PQ221923_Melicope_pteleifolia
PQ636878_Citrus_medica
OX327714_Ailanthus_altissimus
PQ318327_Toona_microcarpa
MZ636518_Acer_miaotaiense
OZ240859_Aesculus_hippocastanum
MK333231_Xanthoceras_sorbifolium
PX549144_Koelreuteria_paniculata
PP932631_Litchi_chinensis
PP916047_Nephelium_lappaceum
PQ859559_Peganum_nigellastrum
MK431824_Nitraria_tangutorum
PQ580749_Myriophyllum_ussuriense
OZ263605_Staphylea_pinnata
OY755230_Euphorbia_lathyris
PP905394_Excoecaria_agallocha
PP938634_Salix_triandra
OP251015_Vernicia_fordii
OX359233_Mercurialis_annua
PX022964_Jacaratia_spinosa
OY756975_Euonymus_europaeus
PX171447_Parnassia_faberi
OZ078050_Vicia_sativa
PP657342_Lathyrus_oleraceus
OZ240033_Ononis_reclinata
PV916041_Medicago_edgeworthii
OX940797_Trifolium_fragiferum
MW441235_Dalbergia_odorifera
MK426679_Castanospermum_australe
PP035769_Styphnolobium_japonicum
PQ043320_Bauhinia_purpurea
PQ723355_Psittacanthus_palmeri
PV936474_Cercis_chinensis
MW448447_Ceratonia_siliqua
PX068309_Begonia_fimbristipula
OW028777_Quercus_robur
PP856682_Castanea_henryi
OZ005622_Myrica_gale
PP533608_Morella_rubra
OZ260459_Betula_pendula
OY340912_Alnus_glutinosa
KF754803_Amborella_trichopoda
0.02

ProTGG-Phe-SerGCT

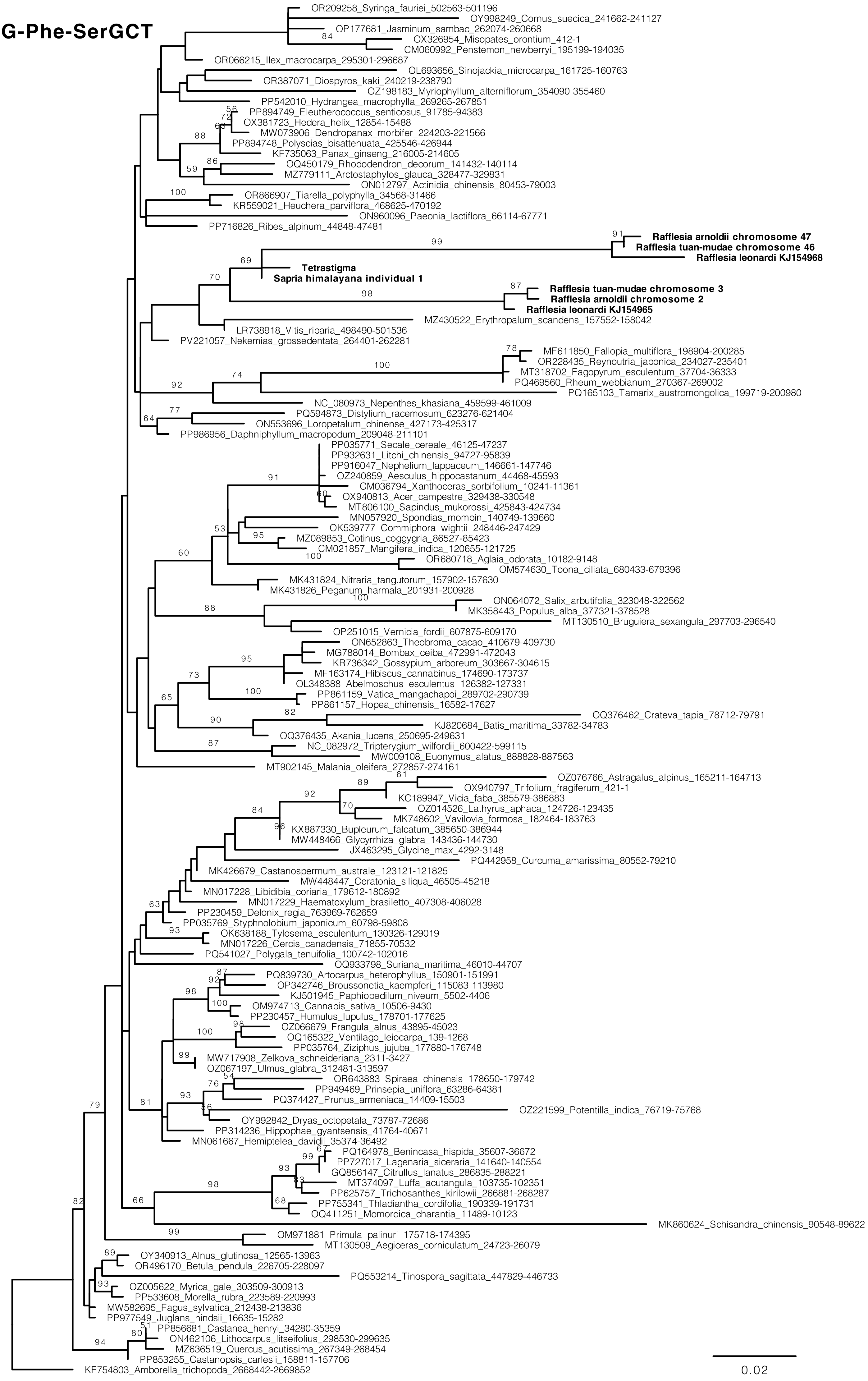

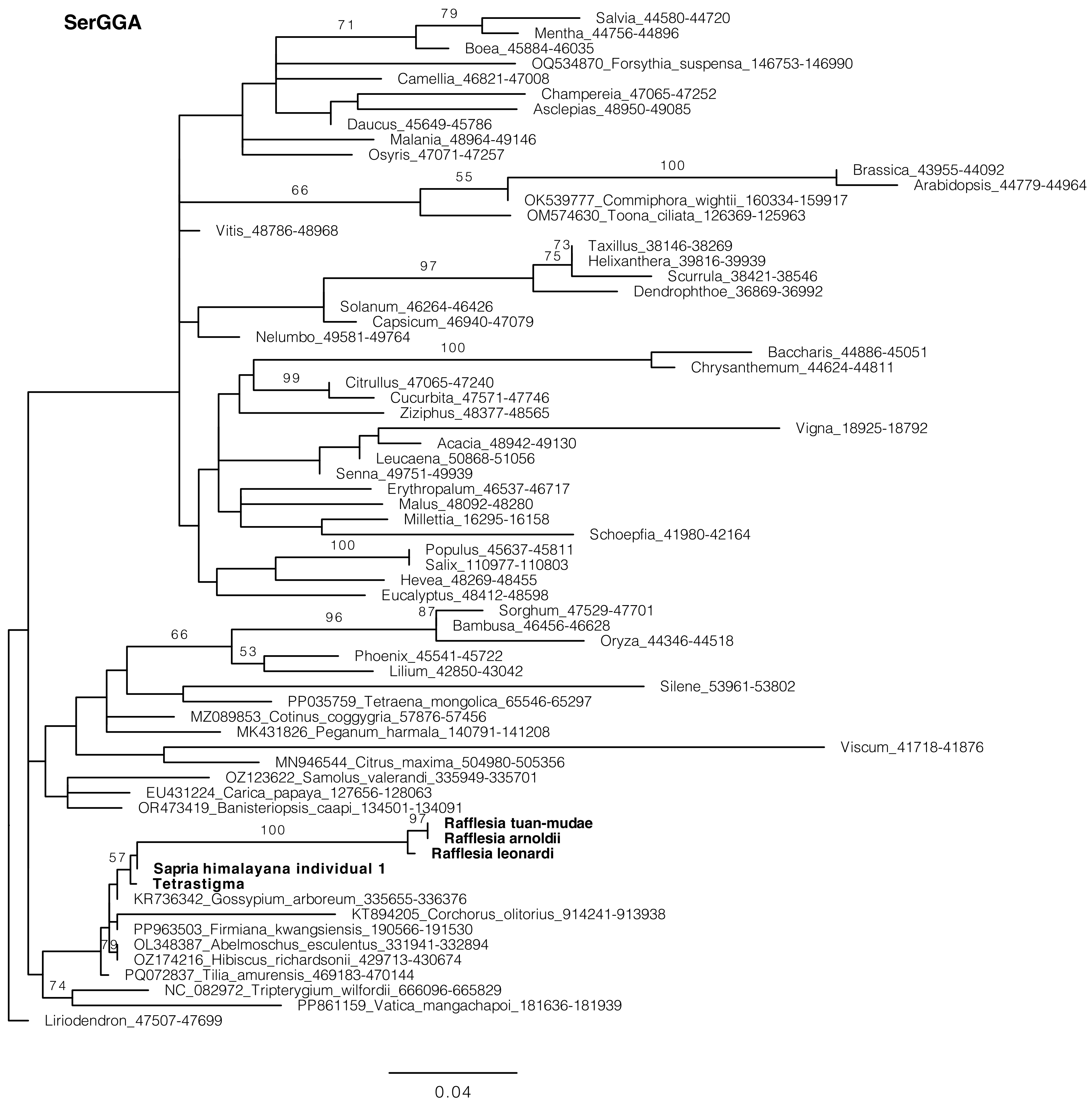
SerGGA
71
79
Salvia_44580-44720
Mentha_44756-44896
Boea_45884-46035
OQ534870_Forsythia_suspensa_146753-146990
Camellia_46821-47008
Champereia_47065-47252
Asclepias_48950-49085
Daucus_45649-45786
Malania_48964-49146
Osyris_47071-47257
100
Brassica_43955-44092
Arabidopsis_44779-44964
66
55
OK539777_Commiphora_wightii_160334-159917
OM574630_Toona_ciliata_126369-125963
Vitis_48786-48968
73
Taxillus_38146-38269
75
Helixanthera_39816-39939
97
Scurrula_38421-38546
Dendrophthoe_36869-36992
Solanum_46264-46426
Capsicum_46940-47079
Nelumbo_49581-49764
100
Baccharis_44886-45051
Chrysanthemum_44624-44811
99
Citrullus_47065-47240
Cucurbita_47571-47746
Ziziphus_48377-48565
Vigna_18925-18792
Acacia_48942-49130
Leucaena_50868-51056
Senna_49751-49939
Erythropalum_46537-46717
Malus_48092-48280
Millettia_16295-16158
Schoepfia_41980-42164
100
Populus_45637-45811
Salix_110977-110803
Hevea_48269-48455
Eucalyptus_48412-48598
87
Sorghum_47529-47701
96
Bambusa_46456-46628
66
Oryza_44346-44518
53
Phoenix_45541-45722
Lilium_42850-43042
Silene_53961-53802
PP035759_Tetraena_mongolica_65546-65297
MZ089853_Cotinus_coggygria_57876-57456
MK431826_Peganum_harmala_140791-141208
Viscum_41718-41876
MN946544_Citrus_maxima_504980-505356
OZ123622_Samolus_valerandi_335949-335701
EU431224_Carica_papaya_127656-128063
OR473419_Banisteriopsis_caapi_134501-134091
97
Rafflesia tuan-mudae
100
Rafflesia arnoldii
Rafflesia leonardi
57
Sapria himalayana individual 1
Tetrastigma
KR736342_Gossypium_arboreum_335655-336376
KT894205_Corchorus_olitorius_914241-913938
PP963503_Firmiana_kwangsiensis_190566-191530
79
OL348387_Abelmoschus_esculentus_331941-332894
OZ174216_Hibiscus_richardsonii_429713-430674
PQ072837_Tilia_amurensis_469183-470144
74
NC_082972_Tripterygium_wilfordii_666096-665829
PP861159_Vatica_mangachapoi_181636-181939
Liriodendron_47507-47699
0.04

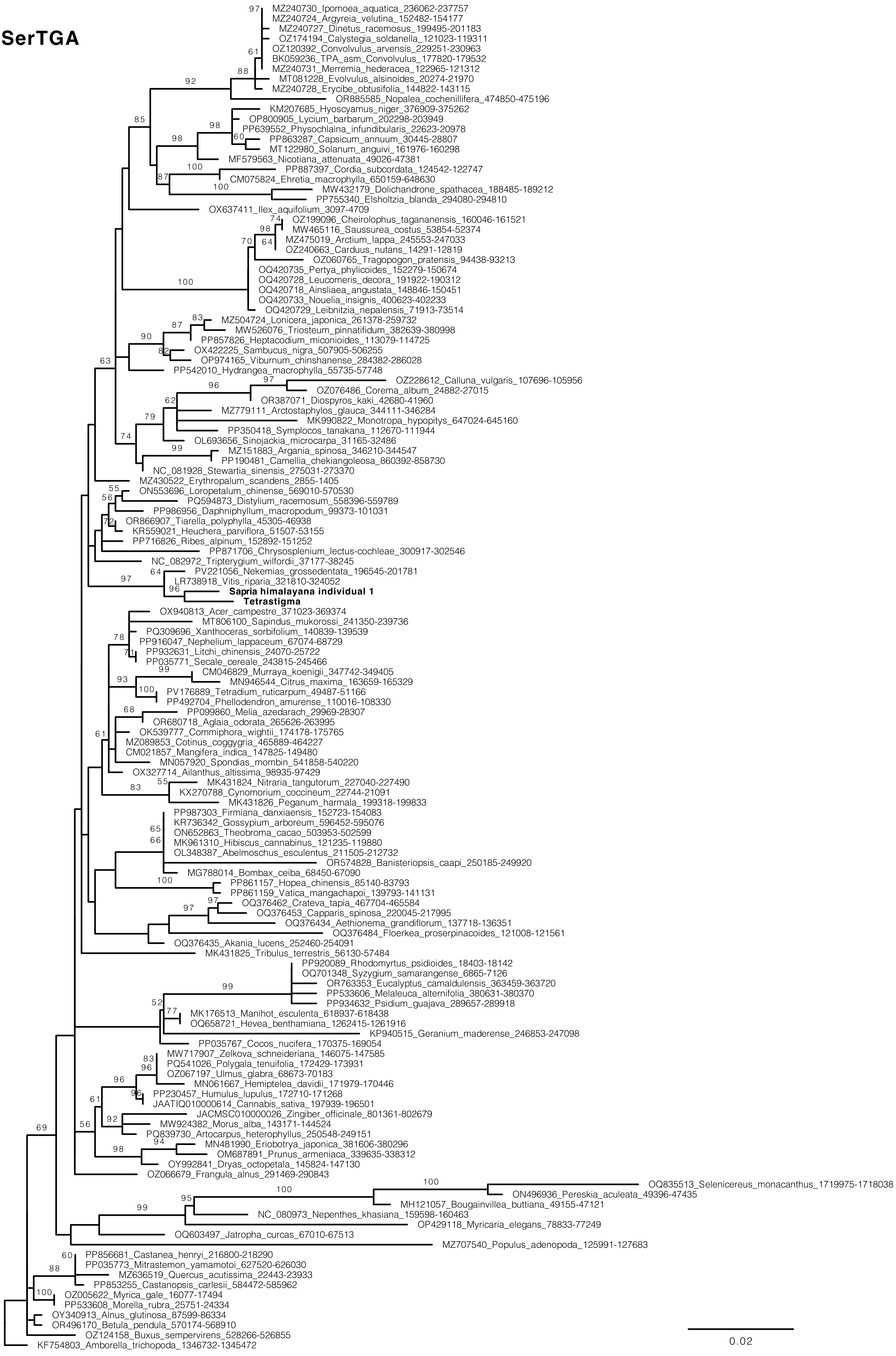
SerTGA
MZ240730_Ipomoea_aquatica_236062-237757
MZ240724_Argyreia_velutina_152482-154177
MZ240727_Dinetus_racemosus_199495-201183
OZ174194_Calystegia_soldanella_121023-119311
OZ120392_Convolvulus_arvensis_229251-230963
BK059236_TPA_asm_Convolvulus_177820-179532
MZ240731_Merremia_hederacea_122965-121312
MT081228_Evolvulus_alsinoides_20274-21970
MZ240728_Erycibe_obtusifolia_144822-143115
OR885585_Nopalea_cochenillifera_474850-475196
KM207685_Hyoscyamus_niger_376909-375262
OP800905_Lycium_barbarum_202298-203949
PP639552_Physochlaina_infundibularis_22623-20978
PP863287_Capsicum_annuum_30445-28807
MT122980_Solanum_anguivi_161976-160298
MF579563_Nicotiana_attenuata_49026-47381
PP887397_Cordia_subcordata_124542-122747
CM075824_Ehretia_macrophylla_650159-648630
MW432179_Dolichandrone_spathacea_188485-189212
PP755340_Elsholtzia_blanda_294080-294810
OX637411_Ilex_aquifolium_3097-4709
OZ199096_Cheirolophus_tagananensis_160046-161521
MW465116_Saussurea_costus_53854-52374
MZ475019_Arctium_lappa_245553-247033
OZ240663_Carduus_nutans_14291-12819
OZ060765_Tragopogon_pratensis_94438-93213
OQ420735_Pertya_phylicoides_152279-150674
OQ420728_Leucomeris_decora_191922-190312
OQ420718_Ainsliaea_angustata_148846-150451
OQ420733_Nouelia_insignis_400623-402233
OQ420729_Leibnitzia_nepalensis_71913-73514
MZ504724_Lonicera_japonica_261378-259732
MW526076_Triosteum_pinnatifidum_382639-380998
PP857826_Heptacodium_miconioides_113079-114725
OX422225_Sambucus_nigra_507905-506255
OP974165_Viburnum_chinshanense_284382-286028
PP542010_Hydrangea_macrophylla_55735-57748
OZ228612_Calluna_vulgaris_107696-105956
OZ076486_Corema_album_24882-27015
OR387071_Diospyros_kaki_42680-41960
MZ779111_Arctostaphylos_glauca_344111-346284
MK990822_Monotropa_hypopitys_647024-645160
PP350418_Symplocos_tanakana_112670-111944
OL693656_Sinojackia_microcarpa_31165-32486
MZ151883_Argania_spinosa_346210-344547
PP190481_Camellia_chekiangoleosa_860392-858730
NC_081928_Stewartia_sinensis_275031-273370
MZ430522_Erythropalum_scandens_2855-1405
ON553696_Loropetalum_chinense_569010-570530
PQ594873_Distylium_racemosum_558396-559789
PP986956_Daphniphyllum_macropodum_99373-101031
OR866907_Tiarella_polyphylla_45305-46938
KR559021_Heuchera_parviflora_51507-53155
PP716826_Ribes_alpinum_152892-151252
PP871706_Chrysosplenium_lectus-cochleae_300917-302546
NC_082972_Tripterygium_wilfordii_37177-38245
PV221056_Nekemias_grossedentata_196545-201781
LR738918_Vitis_riparia_321810-324052
Sapria himalayana individual 1
Tetrastigma
OX940813_Acer_campestre_371023-369374
MT806100_Sapindus_mukorossi_241350-239736
PQ309696_Xanthoceras_sorbifolium_140839-139539
PP916047_Nephelium_lappaceum_67074-68729
PP932631_Litchi_chinensis_24070-25722
PP035771_Secale_cereale_243815-245466
CM046829_Murraya_koenigii_347742-349405
MN946544_Citrus_maxima_163659-165329
PV176889_Tetradium_ruticarpum_49487-51166
PP492704_Phellodendron_amurense_110016-108330
PP099860_Melia_azedarach_29969-28307
OR680718_Aglaia_odorata_265626-263995
OK539777_Commiphora_wightii_174178-175765
MZ089853_Cotinus_coggygria_465889-464227
CM021857_Mangifera_indica_147825-149480
MN057920_Spondias_mombin_541858-540220
OX327714_Ailanthus_altissima_98935-97429
MK431824_Nitraria_tangutorum_227040-227490
KX270788_Cynomorium_coccineum_22744-21091
MK431826_Peganum_harmala_199318-199833
PP987303_Firmiana_danxiaensis_152723-154083
KR736342_Gossypium_arboreum_596452-595076
ON652863_Theobroma_cacao_503953-502599
MK961310_Hibiscus_cannabinus_121235-119880
OL348387_Abelmoschus_esculentus_211505-212732
OR574828_Banisteriopsis_caapi_250185-249920
MG788014_Bombax_ceiba_68450-67090
PP861157_Hopea_chinensis_85140-83793
PP861159_Vatica_mangachapoi_139793-141131
OQ376462_Crateva_tapia_467704-465584
OQ376453_Capparis_spinosa_220045-217995
OQ376434_Aethionema_grandiflorum_137718-136351
OQ376484_Floerkea_proserpinacoides_121008-121561
OQ376435_Akania_lucens_252460-254091
MK431825_Tribulus_terrestris_56130-57484
PP920089_Rhodomyrtus_psidioides_18403-18142
OQ701348_Syzygium_samarangense_6865-7126
OR763353_Eucalyptus_camaldulensis_363459-363720
PP533606_Melaleuca_alternifolia_380631-380370
PP934632_Psidium_guajava_289657-289918
MK176513_Manihot_esculenta_618937-618438
OQ658721_Hevea_benthamiana_1262415-1261916
KP940515_Geranium_maderense_246853-247098
PP035767_Cocos_nucifera_170375-169054
MW717907_Zelkova_schneideriana_146075-147585
PQ541026_Polygala_tenuifolia_172429-173931
OZ067197_Ulmus_glabra_68673-70183
MN061667_Hemiptelea_davidii_171979-170446
PP230457_Humulus_lupulus_172710-171268
JAATIQ010000614_Cannabis_sativa_197939-196501
JACMSC010000026_Zingiber_officinale_801361-802679
MW924382_Morus_alba_143171-144524
PQ839730_Artocarpus_heterophyllus_250548-249151
MN481990_Eriobotrya_japonica_381606-380296
OM687891_Prunus_armeniaca_339635-338312
OY992841_Dryas_octopetala_145824-147130
OZ066679_Frangula_alnus_291469-290843
OQ835513_Selenicereus_monacanthus_1719975-1718038
ON496936_Pereskia_aculeata_49396-47435
MH121057_Bougainvillea_buttiana_49155-47121
NC_080973_Nepenthes_khasiana_159598-160463
OP429118_Myricaria_elegans_78833-77249
OQ603497_Jatropha_curcas_67010-67513
MZ707540_Populus_adenopoda_125991-127683
PP856681_Castanea_henryi_216800-218290
PP035773_Mitrastemon_yamamotoi_627520-626030
MZ636519_Quercus_acutissima_22443-23933
PP853255_Castanopsis_carlesii_584472-585962
OZ005622_Myrica_gale_16077-17494
PP533608_Morella_rubra_25751-24334
OY340913_Alnus_glutinosa_87599-86334
OR496170_Betula_pendula_570174-568910
OZ124158_Buxus_sempervirens_528266-526855
KF754803_Amborella_trichopoda_1346732-1345472
0.02

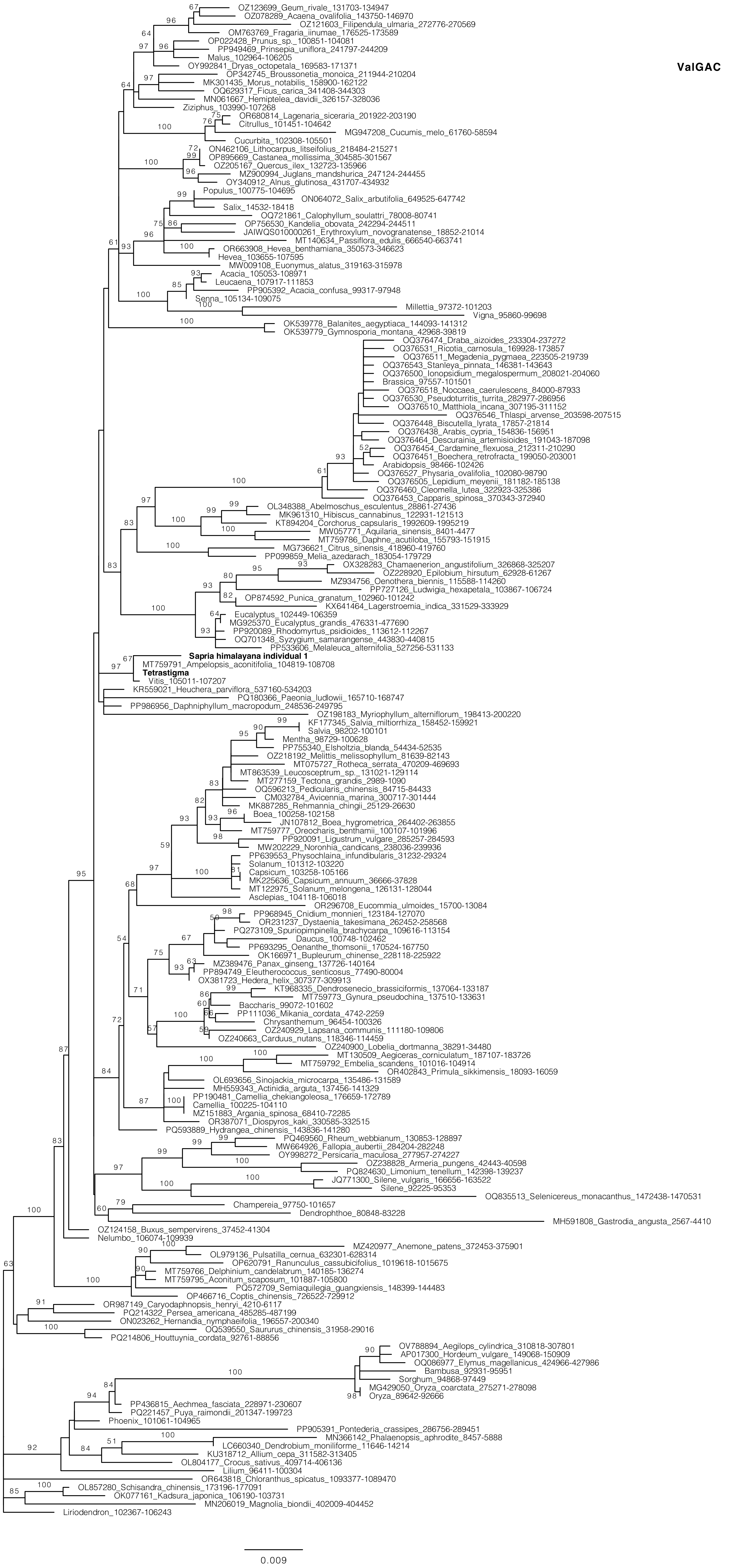

ValGAC
OZ123699_Geum_rivale_131703-134947
OZ078289_Acaena_ovalifolia_143750-146970
OZ121603_Filipendula_ulmaria_272776-270569
OM763769_Fragaria_iinumae_176525-173589
OP022428_Prunus_sp._100851-104081
PP949469_Prinsepia_uniflora_241797-244209
Malus_102964-106205
OY992841_Dryas_octopetala_169583-171371
OP342745_Broussonetia_monoica_211944-210204
MK301435_Morus_notabilis_158900-162122
OQ629317_Ficus_carica_341408-344303
MN061667_Hemiptelea_davidii_326157-328036
Ziziphus_103990-107268
OR680814_Lagenaria_siceraria_201922-203190
Citrullus_101451-104642
MG947208_Cucumis_melo_61760-58594
Cucurbita_102308-105501
ON462106_Lithocarpus_litseifolius_218484-215271
OP895669_Castanea_mollissima_304585-301567
OZ205167_Quercus_ilex_132723-135966
MZ900994_Juglans_mandshurica_247124-244455
OY340912_Alnus_glutinosa_431707-434932
Populus_100775-104695
ON064072_Salix_arbutifolia_649525-647742
Salix_14532-18418
OQ721861_Calophyllum_soulattri_78008-80741
OP756530_Kandelia_obovata_242294-244511
JAIWQS010000261_Erythroxylum_novogranatense_18852-21014
MT140634_Passiflora_edulis_666540-663741
OR663908_Hevea_benthamiana_350573-346623
Hevea_103655-107595
MW009108_Euonymus_alatus_319163-315978
Acacia_105053-108971
Leucaena_107917-111853
PP905392_Acacia_confusa_99317-97948
Senna_105134-109075
Millettia_97372-101203
Vigna_95860-99698
OK539778_Balanites_aegyptiaca_144093-141312
OK539779_Gymnosporia_montana_42968-39819
OQ376474_Draba_aizoides_233304-237272
OQ376531_Ricotia_carnosula_169928-173857
OQ376511_Megadenia_pygmaea_223505-219739
OQ376543_Stanleya_pinnata_146381-143643
OQ376500_Ionopsidium_megalospermum_208021-204060
Brassica_97557-101501
OQ376518_Noccaea_caerulescens_84000-87933
OQ376530_Pseudoturritis_turrita_282977-286956
OQ376510_Matthiola_incana_307195-311152
OQ376546_Thlaspi_arvense_203598-207515
OQ376448_Biscutella_lyrata_17857-21814
OQ376438_Arabis_cypria_154836-156951
OQ376464_Descurainia_artemisioides_191043-187098
OQ376454_Cardamine_flexuosa_212311-210290
OQ376451_Boechera_retrofracta_199050-203001
Arabidopsis_98466-102426
OQ376527_Physaria_ovalifolia_102080-98790
OQ376505_Lepidium_meyenii_181182-185138
OQ376460_Cleomella_lutea_322923-325386
OQ376453_Capparis_spinosa_370343-372940
OL348388_Abelmoschus_esculentus_28861-27436
MK961310_Hibiscus_cannabinus_122931-121513
KT894204_Corchorus_capsularis_1992609-1995219
MW057771_Aquilaria_sinensis_8401-4477
MT759786_Daphne_acutiloba_155793-151915
MG736621_Citrus_sinensis_418960-419760
PP099859_Melia_azedarach_183054-179729
OX328283_Chamaenerion_angustifolium_326868-325207
OZ228920_Epilobium_hirsutum_62928-61267
MZ934756_Oenothera_biennis_115588-114260
PP727126_Ludwigia_hexapetala_103867-106724
OP874592_Punica_granatum_102960-101242
KX641464_Lagerstroemia_indica_331529-333929
Eucalyptus_102449-106359
MG925370_Eucalyptus_grandis_476331-477690
PP920089_Rhodomyrtus_psidioides_113612-112267
OQ701348_Syzygium_samarangense_443830-440815
PP533606_Melaleuca_alternifolia_527256-531133
Sapria himalayana individual 1
MT759791_Ampelopsis_aconitifolia_104819-108708
Tetrastigma
Vitis_105011-107207
KR559021_Heuchera_parviflora_537160-534203
PQ180366_Paeonia_ludlowii_165710-168747
PP986956_Daphniphyllum_macropodum_248536-249795
OZ198183_Myriophyllum_alterniflorum_198413-200220
KF177345_Salvia_miltiorrhiza_158452-159921
Salvia_98202-100101
Mentha_98729-100628
PP755340_Elsholtzia_blanda_54434-52535
OZ218192_Melittis_melissophyllum_81639-82143
MT075727_Rotheca_serrata_470209-469693
MT863539_Leucosceptrum_sp._131021-129114
MT277159_Tectona_grandis_2989-1090
OQ596213_Pedicularis_chinensis_84715-84433
CM032784_Avicennia_marina_300717-301444
MK887285_Rehmannia_chingii_25129-26630
Boea_100258-102158
JN107812_Boea_hygrometrica_264402-263855
MT759777_Oreocharis_benthamii_100107-101996
PP920091_Ligustrum_vulgare_285257-284593
MW202229_Noronhia_candicans_238036-239936
PP639553_Physochlaina_infundibularis_31232-29324
Solanum_101312-103220
Capsicum_103258-105166
MK225636_Capsicum_annuum_36666-37828
MT122975_Solanum_melongena_126131-128044
Asclepias_104118-106018
OR296708_Eucommia_ulmoides_15700-13084
PP968945_Cnidium_monnieri_123184-127070
OR231237_Dystaenia_takesimana_262452-258568
PQ273109_Spuriopimpinella_brachycarpa_109616-113154
Daucus_100748-102462
PP693295_Oenanthe_thomsonii_170524-167750
OK166971_Bupleurum_chinense_228118-225922
MZ389476_Panax_ginseng_137726-140164
PP894749_Eleutherococcus_senticosus_77490-80004
OX381723_Hedera_helix_307377-309913
KT968335_Dendrosenecio_brassiciformis_137064-133187
MT759773_Gynura_pseudochina_137510-133631
Baccharis_99072-101602
PP111036_Mikania_cordata_4742-2259
Chrysanthemum_96454-100326
OZ240929_Lapsana_communis_111180-109806
OZ240663_Carduus_nutans_118346-114459
OZ240900_Lobelia_dortmanna_38291-34480
MT130509_Aegiceras_corniculatum_187107-183726
MT759792_Embelia_scandens_101016-104914
OR402843_Primula_sikkimensis_18093-16059
OL693656_Sinojackia_microcarpa_135486-131589
MH559343_Actinidia_arguta_137456-141329
PP190481_Camellia_chekiangoleosa_176659-172789
Camellia_100225-104110
MZ151883_Argania_spinosa_68410-72285
OR387071_Diospyros_kaki_330585-332515
PQ593889_Hydrangea_chinensis_143836-141280
PQ469560_Rheum_webbianum_130853-128897
MW664926_Fallopia_aubertii_284204-282248
OY998272_Persicaria_maculosa_277957-274227
OZ238828_Armeria_pungens_42443-40598
PQ824630_Limonium_tenellum_142398-139237
JQ771300_Silene_vulgaris_166656-163522
Silene_92225-95353
OQ835513_Selenicereus_monacanthus_1472438-1470531
Champereia_97750-101657
Dendrophthoe_80848-83228
MH591808_Gastrodia_angusta_2567-4410
OZ124158_Buxus_sempervirens_37452-41304
Nelumbo_106074-109939
MZ420977_Anemone_patens_372453-375901
OL979136_Pulsatilla_cernua_632301-628314
OP620791_Ranunculus_cassubicifolius_1019618-1015675
MT759766_Delphinium_candelabrum_140185-136274
MT759795_Aconitum_scaposum_101887-105800
PQ572709_Semiaquilegia_guangxiensis_148399-144483
OP466716_Coptis_chinensis_726522-729912
OR987149_Caryodaphnopsis_henryi_4210-6117
PQ214322_Persea_americana_485285-487199
ON023262_Hernandia_nymphaeifolia_196557-200340
OQ539550_Saururus_chinensis_31958-29016
PQ214806_Houttuynia_cordata_92761-88856
OV788894_Aegilops_cylindrica_310818-307801
AP017300_Hordeum_vulgare_149068-150909
OQ086977_Elymus_magellanicus_424966-427986
Bambusa_92931-95951
Sorghum_94868-97449
MG429050_Oryza_coarctata_275271-278098
Oryza_89642-92666
PP436815_Aechmea_fasciata_228971-230607
PQ221457_Puya_raimondii_201347-199723
Phoenix_101061-104965
PP905391_Pontederia_crassipes_286756-289451
MN366142_Phalaenopsis_aphrodite_8457-5888
LC660340_Dendrobium_moniliforme_11646-14214
KU318712_Allium_cepa_311582-313405
OL804177_Crocus_sativus_409714-406136
Lilium_96411-100304
OR643818_Chloranthus_spicatus_1093377-1089470
OL857280_Schisandra_chinensis_173196-177091
OK077161_Kadsura_japonica_106190-103731
MN206019_Magnolia_biondii_402009-404452
Liriodendron_102367-106243
0.009

**Figure S8. Phylogenetic analyses of mitochondrial plastid-derived sequences (MTPTs) >400 bp of *Sapria himalayana*.** Maximum likelihood analyses were performed with RAxML. ML bootstrap values >50% are shown. The scale bar corresponds to substitutions per site.

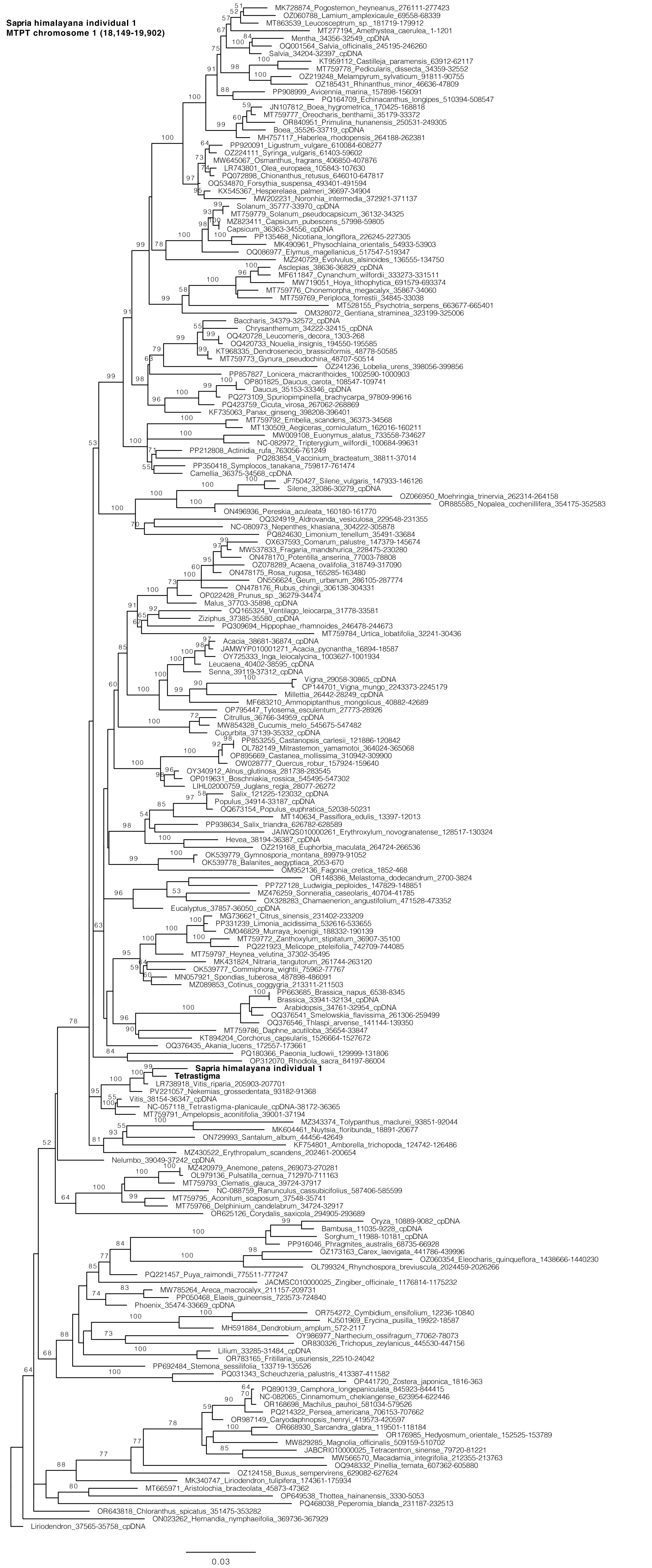

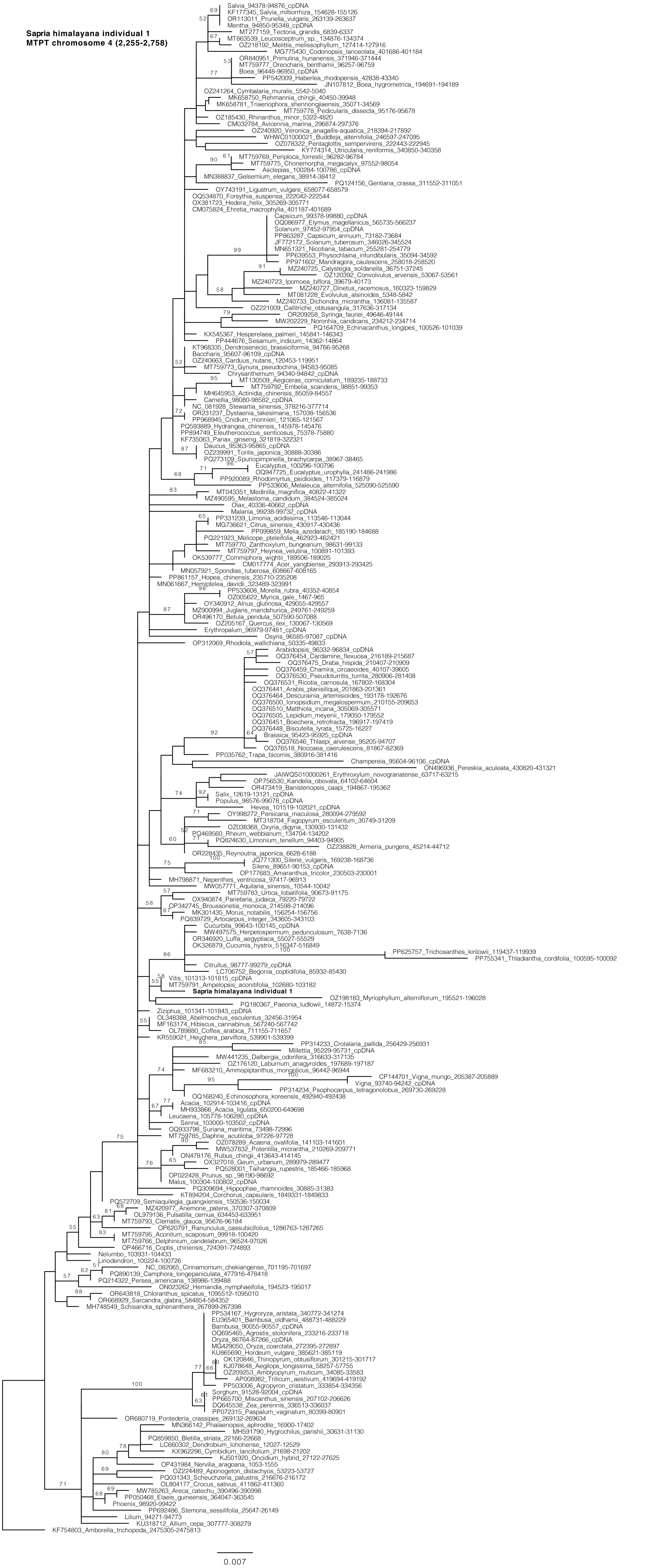

Sapria himalayana individual 1
MTPT chromosome 4 (2,255-2,758)
Salvia_94378-94876_cpDNA
KF177345_Salvia_miltiorrhiza_154628-155126
OR113011_Prunella_vulgaris_263139-263637
Mentha_94850-95348_cpDNA
MT277159_Tectona_grandis_6839-6337
MT863539_Leucosceptrum_sp._134876-134374
OZ218192_Melittis_melissophyllum_127414-127916
MG775430_Codonopsis_lanceolata_401686-401184
OR840951_Primulina_hunanensis_371946-371444
MT759777_Oreocharis_benthamii_96257-96759
Boea_96448-96950_cpDNA
PP542009_Haberlea_rhodopensis_42838-43340
JN107812_Boea_hygrometrica_194691-194189
OZ241264_Cymbalaria_muralis_5542-5040
MK658750_Rehmannia_chingii_40450-39948
MK658781_Triaenophora_shennongjiaensis_35071-34569
MT759778_Pedicularis_dissecta_95176-95678
OZ185430_Rhinanthus_minor_5322-4820
CM032784_Avicennia_marina_296874-297376
OZ240920_Veronica_anagallis-aquatica_218394-217892
WHWC01000021_Buddleja_alternifolia_246597-247095
OZ078322_Pentaglottis_sempervirens_222443-222945
KY774314_Utricularia_reniformis_340850-340358
MT759769_Periploca_forrestii_96282-96784
MT759775_Chonemorpha_megacalyx_97552-98054
Asclepias_100284-100786_cpDNA
MN388837_Gelsemium_elegans_38914-38412
PQ124156_Gentiana_crassa_311552-311051
OY743191_Ligustrum_vulgare_658077-658579
OQ534870_Forsythia_suspensa_222042-222544
OX381723_Hedera_helix_305269-305771
CM075824_Ehretia_macrophylla_401187-401689
Capsicum_99378-99880_cpDNA
OQ086977_Elymus_magellanicus_565735-566237
Solanum_97452-97954_cpDNA
PP863287_Capsicum_annuum_73182-73684
JF772172_Solanum_tuberosum_346026-345524
MN651321_Nicotiana_tabacum_255281-254779
PP639553_Physochlaina_infundibularis_35094-34592
PP971602_Mandragora_caulescens_258018-258520
MZ240725_Calystegia_soldanella_36751-37245
OZ120392_Convolvulus_arvensis_53067-53561
MZ240723_Ipomoea_biflora_39679-40173
MZ240727_Dinetus_racemosus_160323-159829
MT081228_Evolvulus_alsinoides_5348-5842
MZ240733_Dichondra_micrantha_136081-135587
OZ221009_Callitriche_obtusangula_317636-317134
OR209258_Syringa_fauriei_49646-49144
MW202229_Noronhia_candicans_234212-234714
PQ164709_Echinacanthus_longipes_100526-101039
KX545367_Hesperelaea_palmeri_145841-146343
PP444676_Sesamum_indicum_14362-14864
KT968335_Dendrosenecio_brassiciformis_94766-95268
Baccharis_95607-96109_cpDNA
OZ240663_Carduus_nutans_120453-119951
MT759773_Gynura_pseudochina_94583-95085
Chrysanthemum_94340-94842_cpDNA
MT130509_Aegiceras_corniculatum_189235-188733
MT759792_Embelia_scandens_98851-99353
MH645953_Actinidia_chinensis_85059-84557
Camellia_98080-98582_cpDNA
NC_081928_Stewartia_sinensis_378216-377714
OR231237_Dystaenia_takesimana_157038-156536
PP968945_Cnidium_monnieri_121065-121567
PQ593889_Hydrangea_chinensis_145978-145476
PP894749_Eleutherococcus_senticosus_75378-75880
KF735063_Panax_ginseng_321819-322321
Daucus_95363-95865_cpDNA
OZ239991_Torilis_japonica_30888-30386
PQ273109_Spuriopimpinella_brachycarpa_38967-38465
Eucalyptus_100296-100796
OQ947725_Eucalyptus_urophylla_241486-241986
PP920089_Rhodomyrtus_psidioides_117379-116879
PP533606_Melaleuca_alternifolia_525090-525590
MT043351_Medinilla_magnifica_40822-41322
MZ490595_Melastoma_candidum_384524-385024
Olax_40336-40662_cpDNA
Malania_99238-99732_cpDNA
PP331239_Limonia_acidissima_113546-113044
MG736621_Citrus_sinensis_430917-430436
PP099859_Melia_azedarach_185190-184688
PQ221923_Melicope_pteleifolia_462923-462421
MT759770_Zanthoxylum_bungeanum_98631-99133
MT759797_Heynea_velutina_100891-101393
OK539777_Commiphora_wightii_189506-189025
CM017774_Acer_yangbiense_293913-293425
MN057921_Spondias_tuberosa_608667-608165
PP861157_Hopea_chinensis_235710-235208
MN061667_Hemiptelea_davidii_323489-323991
PP533608_Morella_rubra_40352-40854
OZ005622_Myrica_gale_1467-965
OY340912_Alnus_glutinosa_429055-429557
MZ900994_Juglans_mandshurica_249761-249259
OR496170_Betula_pendula_507590-507088
OZ205167_Quercus_ilex_130067-130569
Erythropalum_96979-97481_cpDNA
Osyris_96585-97087_cpDNA
OP312069_Rhodiola_wallichiana_50335-49833
Arabidopsis_96332-96834_cpDNA
OQ376454_Cardamine_flexuosa_216189-215687
OQ376475_Draba_hispida_210407-210909
OQ376459_Chamira_circaeoides_40107-39605
OQ376530_Pseudoturritis_turrita_280906-281408
OQ376531_Ricotia_carnosula_167802-168304
OQ376441_Arabis_planisiliqua_201863-201361
OQ376464_Descurainia_artemisioides_193178-192676
OQ376500_Ionopsidium_megalospermum_210155-209653
OQ376510_Matthiola_incana_305069-305571
OQ376505_Lepidium_meyenii_179050-179552
OQ376451_Boechera_retrofracta_196917-197419
OQ376448_Biscutella_lyrata_15725-16227
Brassica_95423-95925_cpDNA
OQ376546_Thlaspi_arvense_95205-94707
OQ376518_Noccaea_caerulescens_81867-82369
PP035762_Trapa_bicornis_380916-381416
Champereia_95604-96106_cpDNA
ON496936_Pereskia_aculeata_430820-431321
JAIWQS010000261_Erythroxylum_novogranatense_63717-63215
OP756530_Kandelia_obovata_64102-64604
OR473419_Banisteriopsis_caapi_194867-195362
Salix_12619-13121_cpDNA
Populus_98576-99078_cpDNA
Hevea_101519-102021_cpDNA
OY998272_Persicaria_maculosa_280094-279592
MT318704_Fagopyrum_esculentum_30749-31209
OZ038368_Oxyria_digyna_130930-131432
PQ469560_Rheum_webbianum_134704-134202
PQ824630_Limonium_tenellum_94403-94905
OZ238828_Armeria_pungens_45214-44712
OR228435_Reynoutria_japonica_6628-6188
JQ771300_Silene_vulgaris_169238-168736
Silene_89651-90153_cpDNA
OP177683_Amaranthus_tricolor_230503-230001
MH798871_Nepenthes_ventricosa_97417-96913
MW057771_Aquilaria_sinensis_10544-10042
MT759783_Urtica_lobatifolia_90673-91175
OX940874_Parietaria_judaica_79220-79722
OP342745_Broussonetia_monoica_214598-214096
MK301435_Morus_notabilis_156254-156756
PQ839729_Artocarpus_integer_343605-343103
Cucurbita_99643-100145_cpDNA
MW497575_Herpetospermum_pedunculosum_7638-7136
OR346920_Luffa_aegyptiaca_55027-55529
OK326879_Cucumis_hystrix_516347-516849
PP625757_Trichosanthes_kirilowii_119437-119939
PP755341_Thladiantha_cordifolia_100595-100092
Citrullus_98777-99279_cpDNA
LC706752_Begonia_coptidifolia_85932-85430
Vitis_101313-101815_cpDNA
MT759791_Ampelopsis_aconitifolia_102680-103182
Sapria himalayana individual 1
OZ198183_Myriophyllum_alterniflorum_195521-196028
PQ180367_Paeonia_ludlowii_14872-15374
Ziziphus_101341-101843_cpDNA
OL348388_Abelmoschus_esculentus_32456-31954
MF163174_Hibiscus_cannabinus_567240-567742
OL789880_Coffea_arabica_711155-711657
KR559021_Heuchera_parviflora_539901-539399
PP314233_Crotalaria_pallida_256429-256931
Millettia_95229-95731_cpDNA
MW441235_Dalbergia_odorifera_316633-317135
OZ176120_Laburnum_anagyroides_197689-197187
MF683210_Ammopiptanthus_mongolicus_96442-96944
CP144701_Vigna_mungo_205387-205889
Vigna_93740-94242_cpDNA
PP314234_Psophocarpus_tetragonolobus_269730-269228
OQ168240_Echinosophora_koreensis_492940-492438
Acacia_102914-103416_cpDNA
MH933866_Acacia_ligulata_650200-649698
Leucaena_105778-106280_cpDNA
Senna_103000-103502_cpDNA
OQ933798_Suriana_maritima_73498-72996
MT759785_Daphne_acutiloba_97226-97728
OZ078289_Acaena_ovalifolia_141103-141601
MW537832_Potentilla_micrantha_210269-209771
ON478176_Rubus_chingii_413643-414145
OX327018_Geum_urbanum_289979-289477
PQ528001_Taihangia_rupestris_185466-185968
OP022428_Prunus_sp._98190-98692
Malus_100304-100802_cpDNA
PQ309694_Hippophae_rhamnoides_30885-31383
KT894204_Corchorus_capsularis_1849331-1849833
PQ572709_Semiaquilegia_guangxiensis_150536-150034
MZ420977_Anemone_patens_370307-370809
OL979136_Pulsatilla_cernua_634453-633951
MT759793_Clematis_glauca_95676-96184
OP620791_Ranunculus_cassubicifolius_1286763-1287265
MT759795_Aconitum_scaposum_99918-100420
MT759766_Delphinium_candelabrum_96524-97026
OP466716_Coptis_chinensis_724391-724893
Nelumbo_103931-104433
Liriodendron_100224-100726
NC_082065_Cinnamomum_chekiangense_701195-701697
PQ890139_Camphora_longepaniculata_477916-478418
PQ214322_Persea_americana_138986-139488
ON023262_Hernandia_nymphaeifolia_194523-195017
OR643818_Chloranthus_spicatus_1095512-1095010
OR668929_Sarcandra_glabra_584854-584352
MH748549_Schisandra_sphenanthera_267899-267398
PP534167_Hygroryza_aristata_340772-341274
EU365401_Bambusa_oldhamii_488731-488229
Bambusa_90055-90557_cpDNA
OQ695465_Agrostis_stolonifera_233216-233718
Oryza_86764-87266_cpDNA
MG429050_Oryza_coarctata_272395-272897
KU865690_Hordeum_vulgare_385621-385119
OK120846_Thinopyrum_obtusiflorum_301215-301717
KJ078648_Aegilops_longissima_58257-57755
OZ209253_Amblyopyrum_muticum_34085-33583
AP008982_Triticum_aestivum_419694-419192
PP503006_Agropyron_cristatum_333854-334356
Sorghum_91528-92004_cpDNA
PP665700_Miscanthus_sinensis_207102-206626
DQ645538_Zea_perennis_336513-336037
PP072315_Paspalum_vaginatum_80399-80901
OR680719_Pontederia_crassipes_269132-269634
MN366142_Phalaenopsis_aphrodite_16900-17402
MH591790_Hygrochilus_parishii_30631-31130
PQ859850_Bletilla_striata_22166-22668
LC660302_Dendrobium_lohohense_12027-12529
KX962296_Cymbidium_lancifolium_21698-21202
KJ501920_Oncidium_hybrid_27122-27625
OP431984_Nervilia_aragoana_1053-1555
OZ224489_Aponogeton_distachyos_53223-53727
PQ031343_Scheuchzeria_palustris_216676-216172
OL804177_Crocus_sativus_411862-411360
MW785263_Areca_catechu_390496-390998
PP050468_Elaeis_guineensis_364047-363545
Phoenix_98920-99422
PP692486_Stemona_sessilifolia_25647-26149
Lilium_94271-94773
KU318712_Allium_cepa_307777-308279
KF754803_Amborella_trichopoda_2475305-2475813
0.007

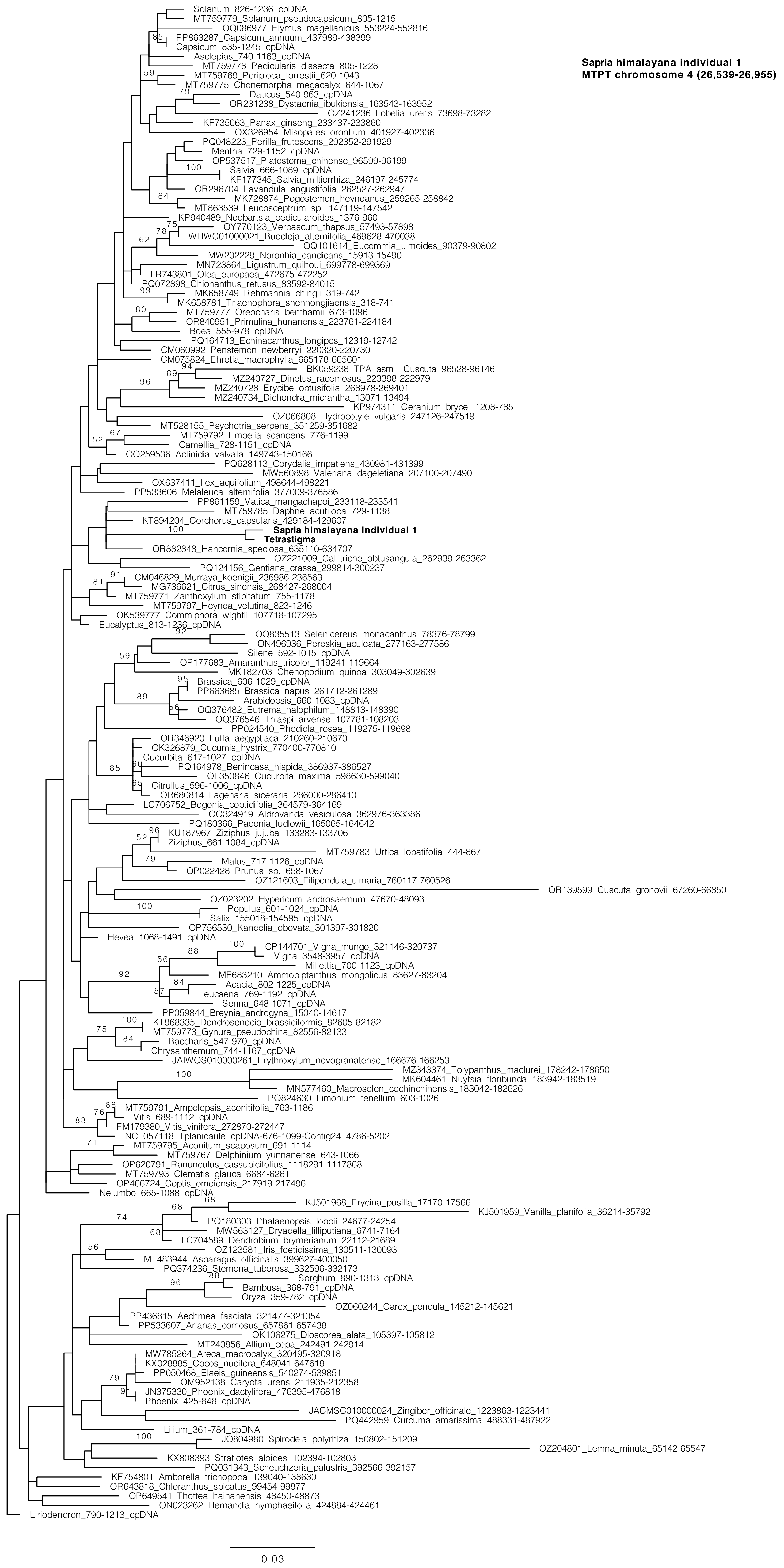

Sapria himalayana individual 1
MTPT chromosome 4 (26,539-26,955)
Solanum_826-1236_cpDNA
MT759779_Solanum_pseudocapsicum_805-1215
OQ086977_Elymus_magellanicus_553224-552816
PP863287_Capsicum_annuum_437989-438399
Capsicum_835-1245_cpDNA
Asclepias_740-1163_cpDNA
MT759778_Pedicularis_dissecta_805-1228
MT759769_Periploca_forrestii_620-1043
MT759775_Chonemorpha_megacalyx_644-1067
Daucus_540-963_cpDNA
OR231238_Dystaenia_ibukiensis_163543-163952
OZ241236_Lobelia_urens_73698-73282
KF735063_Panax_ginseng_233437-233860
OX326954_Misopates_orontium_401927-402336
PQ048223_Perilla_frutescens_292352-291929
Mentha_729-1152_cpDNA
OP537517_Platostoma_chinense_96599-96199
Salvia_666-1089_cpDNA
KF177345_Salvia_miltiorrhiza_246197-245774
OR296704_Lavandula_angustifolia_262527-262947
MK728874_Pogostemon_heyneanus_259265-258842
MT863539_Leucosceptrum_sp._147119-147542
KP940489_Neobartsia_pedicularoides_1376-960
OY770123_Verbascum_thapsus_57493-57898
WHWC01000021_Buddleja_alternifolia_469628-470038
OQ101614_Eucommia_ulmoides_90379-90802
MW202229_Noronhia_candicans_15913-15490
MN723864_Ligustrum_quihoui_699778-699369
LR743801_Olea_europaea_472675-472252
PQ072898_Chionanthus_retusus_83592-84015
MK658749_Rehmannia_chingii_319-742
MK658781_Triaenophora_shennongjiaensis_318-741
MT759777_Oreocharis_benthamii_673-1096
OR840951_Primulina_hunanensis_223761-224184
Boea_555-978_cpDNA
PQ164713_Echinacanthus_longipes_12319-12742
CM060992_Penstemon_newberryi_220320-220730
CM075824_Ehretia_macrophylla_665178-665601
BK059238_TPA_asm__Cuscuta_96528-96146
MZ240727_Dinetus_racemosus_223398-222979
MZ240728_Erycibe_obtusifolia_268978-269401
MZ240734_Dichondra_micrantha_13071-13494
KP974311_Geranium_brycei_1208-785
OZ066808_Hydrocotyle_vulgaris_247126-247519
MT528155_Psychotria_serpens_351259-351682
MT759792_Embelia_scandens_776-1199
Camellia_728-1151_cpDNA
OQ259536_Actinidia_valvata_149743-150166
PQ628113_Corydalis_impatiens_430981-431399
MW560898_Valeriana_dageletiana_207100-207490
OX637411_Ilex_aquifolium_498644-498221
PP533606_Melaleuca_alternifolia_377009-376586
PP861159_Vatica_mangachapoi_233118-233541
MT759785_Daphne_acutiloba_729-1138
KT894204_Corchorus_capsularis_429184-429607
Sapria himalayana individual 1
Tetrastigma
OR882848_Hancornia_speciosa_635110-634707
OZ221009_Callitriche_obtusangula_262939-263362
PQ124156_Gentiana_crassa_299814-300237
CM046829_Murraya_koenigii_236986-236563
MG736621_Citrus_sinensis_268427-268004
MT759771_Zanthoxylum_stipitatum_755-1178
MT759797_Heynea_velutina_823-1246
OK539777_Commiphora_wightii_107718-107295
Eucalyptus_813-1236_cpDNA
OQ835513_Selenicereus_monacanthus_78376-78799
ON496936_Pereskia_aculeata_277163-277586
Silene_592-1015_cpDNA
OP177683_Amaranthus_tricolor_119241-119664
MK182703_Chenopodium_quinoa_303049-302639
Brassica_606-1029_cpDNA
PP663685_Brassica_napus_261712-261289
Arabidopsis_660-1083_cpDNA
OQ376482_Eutrema_halophilum_148813-148390
OQ376546_Thlaspi_arvense_107781-108203
PP024540_Rhodiola_rosea_119275-119698
OR346920_Luffa_aegyptiaca_210260-210670
OK326879_Cucumis_hystrix_770400-770810
Cucurbita_617-1027_cpDNA
PQ164978_Benincasa_hispida_386937-386527
OL350846_Cucurbita_maxima_598630-599040
Citrullus_596-1006_cpDNA
OR680814_Lagenaria_siceraria_286000-286410
LC706752_Begonia_coptidifolia_364579-364169
OQ324919_Aldrovanda_vesiculosa_362976-363386
PQ180366_Paeonia_ludlowii_165065-164642
KU187967_Ziziphus_jujuba_133283-133706
Ziziphus_661-1084_cpDNA
MT759783_Urtica_lobatifolia_444-867
Malus_717-1126_cpDNA
OP022428_Prunus_sp._658-1067
OZ121603_Filipendula_ulmaria_760117-760526
OR139599_Cuscuta_gronovii_67260-66850
OZ023202_Hypericum_androsaemum_47670-48093
Populus_601-1024_cpDNA
Salix_155018-154595_cpDNA
OP756530_Kandelia_obovata_301397-301820
Hevea_1068-1491_cpDNA
CP144701_Vigna_mungo_321146-320737
Vigna_3548-3957_cpDNA
Millettia_700-1123_cpDNA
MF683210_Ammopiptanthus_mongolicus_83627-83204
Acacia_802-1225_cpDNA
Leucaena_769-1192_cpDNA
Senna_648-1071_cpDNA
PP059844_Breynia_androgyna_15040-14617
KT968335_Dendrosenecio_brassiciformis_82605-82182
MT759773_Gynura_pseudochina_82556-82133
Baccharis_547-970_cpDNA
Chrysanthemum_744-1167_cpDNA
JAIWQS010000261_Erythroxylum_novogranatense_166676-166253
MZ343374_Tolypanthus_maclurei_178242-178650
MK604461_Nuytsia_floribunda_183942-183519
MN577460_Macrosolen_cochinchinensis_183042-182626
PQ824630_Limonium_tenellum_603-1026
MT759791_Ampelopsis_aconitifolia_763-1186
Vitis_689-1112_cpDNA
FM179380_Vitis_vinifera_272870-272447
NC_057118_Tplanicaule_cpDNA-676-1099-Contig24_4786-5202
MT759795_Aconitum_scaposum_691-1114
MT759767_Delphinium_yunnanense_643-1066
OP620791_Ranunculus_cassubicifolius_1118291-1117868
MT759793_Clematis_glauca_6684-6261
OP466724_Coptis_omeiensis_217919-217496
Nelumbo_665-1088_cpDNA
KJ501968_Erycina_pusilla_17170-17566
KJ501959_Vanilla_planifolia_36214-35792
PQ180303_Phalaenopsis_lobbii_24677-24254
MW563127_Dryadella_lilliputiana_6741-7164
LC704589_Dendrobium_brymerianum_22112-21689
OZ123581_Iris_foetidissima_130511-130093
MT483944_Asparagus_officinalis_399627-400050
PQ374236_Stemona_tuberosa_332596-332173
Sorghum_890-1313_cpDNA
Bambusa_368-791_cpDNA
Oryza_359-782_cpDNA
OZ060244_Carex_pendula_145212-145621
PP436815_Aechmea_fasciata_321477-321054
PP533607_Ananas_comosus_657861-657438
OK106275_Dioscorea_alata_105397-105812
MT240856_Allium_cepa_242491-242914
MW785264_Areca_macrocalyx_320495-320918
KX028885_Cocos_nucifera_648041-647618
PP050468_Elaeis_guineensis_540274-539851
OM952138_Caryota_urens_211935-212358
JN375330_Phoenix_dactylifera_476395-476818
Phoenix_425-848_cpDNA
JACMSC010000024_Zingiber_officinale_1223863-1223441
PQ442959_Curcuma_amarissima_488331-487922
Lilium_361-784_cpDNA
JQ804980_Spirodela_polyrhiza_150802-151209
OZ204801_Lemna_minuta_65142-65547
KX808393_Stratiotes_aloides_102394-102803
PQ031343_Scheuchzeria_palustris_392566-392157
KF754801_Amborella_trichopoda_139040-138630
OR643818_Chloranthus_spicatus_99454-99877
OP649541_Thottea_hainanensis_48450-48873
ON023262_Hernandia_nymphaeifolia_424884-424461
Liriodendron_790-1213_cpDNA
0.03

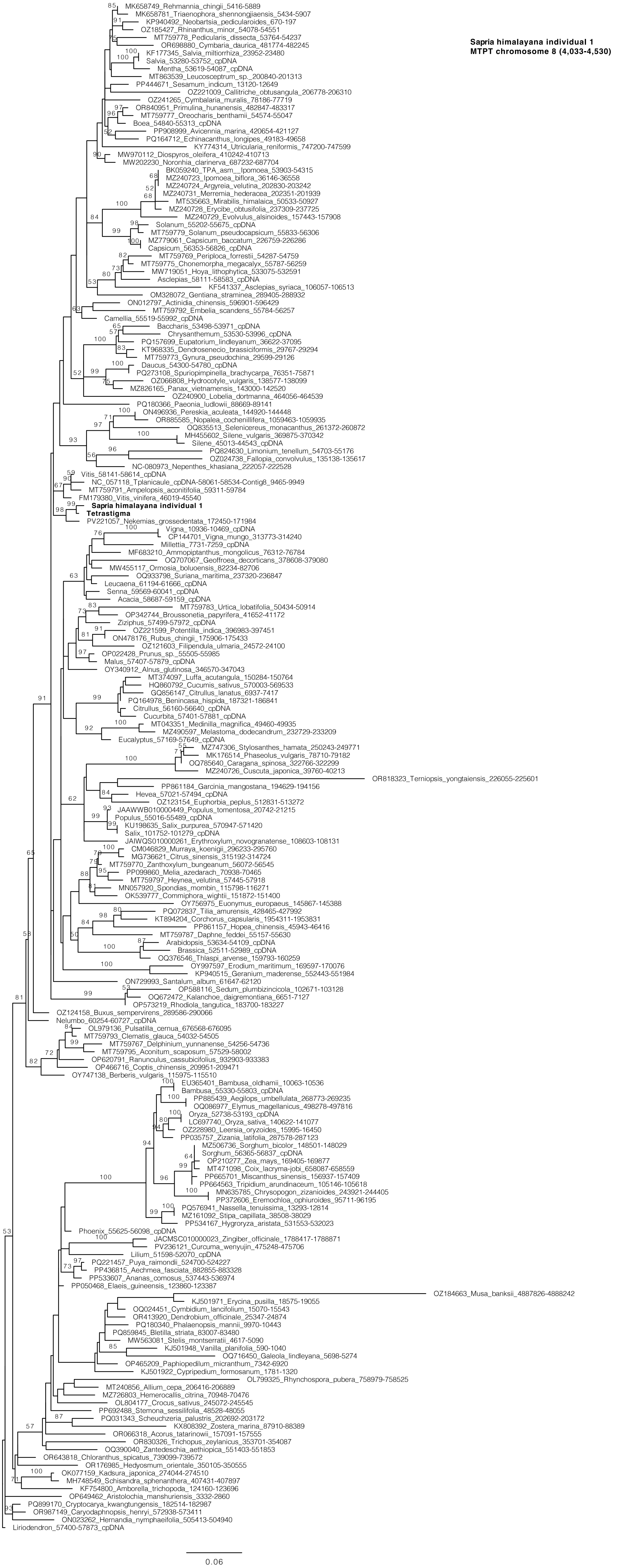
Sapria himalayana individual 1
MTPT chromosome 8 (4,033-4,530)
MK658749_Rehmannia_chingii_5416-5889
MK658781_Triaenophora_shennongjiaensis_5434-5907
KP940492_Neobartsia_pedicularoides_670-197
OZ185427_Rhinanthus_minor_54078-54551
MT759778_Pedicularis_dissecta_53764-54237
OR698880_Cymbaria_daurica_481774-482245
KF177345_Salvia_miltiorrhiza_23952-23480
Salvia_53280-53752_cpDNA
Mentha_53619-54087_cpDNA
MT863539_Leucosceptrum_sp._200840-201313
PP444671_Sesamum_indicum_13120-12649
OZ221009_Callitriche_obtusangula_206778-206310
OZ241265_Cymbalaria_muralis_78186-77719
OR840951_Primulina_hunanensis_482847-483317
MT759777_Oreocharis_benthamii_54574-55047
Boea_54840-55313_cpDNA
PP908999_Avicennia_marina_420654-421127
PQ164712_Echinacanthus_longipes_49183-49658
KY774314_Utricularia_reniformis_747200-747599
MW970112_Diospyros_oleifera_410242-410713
MW202230_Noronhia_clarinerva_687232-687704
BK059240_TPA_asm__Ipomoea_53903-54315
MZ240723_Ipomoea_biflora_36146-36558
MZ240724_Argyreia_velutina_202830-203242
MZ240731_Merremia_hederacea_202351-201939
MT535663_Mirabilis_himalaica_50533-50927
MZ240728_Erycibe_obtusifolia_237309-237725
MZ240729_Evolvulus_alsinoides_157443-157908
Solanum_55202-55675_cpDNA
MT759779_Solanum_pseudocapsicum_55833-56306
MZ779061_Capsicum_baccatum_226759-226286
Capsicum_56353-56826_cpDNA
MT759769_Periploca_forrestii_54287-54759
MT759775_Chonemorpha_megacalyx_55787-56259
MW719051_Hoya_lithophytica_533075-532591
Asclepias_58111-58583_cpDNA
KF541337_Asclepias_syriaca_106057-106513
OM328072_Gentiana_straminea_289405-288932
ON012797_Actinidia_chinensis_596901-596429
MT759792_Embelia_scandens_55784-56257
Camellia_55519-55992_cpDNA
Baccharis_53498-53971_cpDNA
Chrysanthemum_53530-53996_cpDNA
PQ157699_Eupatorium_lindleyanum_36622-37095
KT968335_Dendrosenecio_brassiciformis_29767-29294
MT759773_Gynura_pseudochina_29599-29126
Daucus_54300-54780_cpDNA
PQ273108_Spuriopimpinella_brachycarpa_76351-75871
OZ066808_Hydrocotyle_vulgaris_138577-138099
MZ826165_Panax_vietnamensis_143000-142520
OZ240900_Lobelia_dortmanna_464056-464539
PQ180366_Paeonia_ludlowii_88669-89141
ON496936_Pereskia_aculeata_144920-144448
OR885585_Nopalea_cochenillifera_1059463-1059935
OQ835513_Selenicereus_monacanthus_261372-260872
MH455602_Silene_vulgaris_369875-370342
Silene_45013-44543_cpDNA
PQ824630_Limonium_tenellum_54703-55176
OZ024738_Fallopia_convolvulus_135138-135617
NC-080973_Nepenthes_khasiana_222057-222528
Vitis_58141-58614_cpDNA
NC_057118_Tplanicaule_cpDNA-58061-58534-Contig8_9465-9949
MT759791_Ampelopsis_aconitifolia_59311-59784
FM179380_Vitis_vinifera_46019-45540
Sapria himalayana individual 1
Tetrastigma
PV221057_Nekemias_grossedentata_172450-171984
Vigna_10936-10469_cpDNA
CP144701_Vigna_mungo_313773-314240
Millettia_7731-7259_cpDNA
MF683210_Ammopiptanthus_mongolicus_76312-76784
OQ707067_Geoffroea_decorticans_378608-379080
MW455117_Ormosia_boluoensis_82234-82706
OQ933798_Suriana_maritima_237320-236847
Leucaena_61194-61666_cpDNA
Senna_59569-60041_cpDNA
Acacia_58687-59159_cpDNA
MT759783_Urtica_lobatifolia_50434-50914
OP342744_Broussonetia_papyrifera_41652-41172
Ziziphus_57499-57972_cpDNA
OZ221599_Potentilla_indica_396983-397451
ON478176_Rubus_chingii_175906-175433
OZ121603_Filipendula_ulmaria_24572-24100
OP022428_Prunus_sp._55505-55985
Malus_57407-57879_cpDNA
OY340912_Alnus_glutinosa_346570-347043
MT374097_Luffa_acutangula_150284-150764
HQ860792_Cucumis_sativus_570003-569533
GQ856147_Citrullus_lanatus_6937-7417
PQ164978_Benincasa_hispida_187321-186841
Citrullus_56160-56640_cpDNA
Cucurbita_57401-57881_cpDNA
MT043351_Medinilla_magnifica_49460-49935
MZ490597_Melastoma_dodecandrum_232729-233209
Eucalyptus_57169-57649_cpDNA
MZ747306_Stylosanthes_hamata_250243-249771
MK176514_Phaseolus_vulgaris_78710-79182
OQ785640_Caragana_spinosa_322766-322299
MZ240726_Cuscuta_japonica_39760-40213
OR818323_Terniopsis_yongtaiensis_226055-225601
PP861184_Garcinia_mangostana_194629-194156
Hevea_57021-57494_cpDNA
OZ123154_Euphorbia_peplus_512831-513272
JAAWWB010000449_Populus_tomentosa_20742-21215
Populus_55016-55489_cpDNA
KU198635_Salix_purpurea_570947-571420
Salix_101752-101279_cpDNA
JAIWQS010000261_Erythroxylum_novogranatense_108603-108131
CM046829_Murraya_koenigii_296233-295760
MG736621_Citrus_sinensis_315192-314724
MT759770_Zanthoxylum_bungeanum_56072-56545
PP099860_Melia_azedarach_70938-70465
MT759797_Heynea_velutina_57445-57918
MN057920_Spondias_mombin_115798-116271
OK539777_Commiphora_wightii_151872-151400
OY756975_Euonymus_europaeus_145867-145388
PQ072837_Tilia_amurensis_428465-427992
KT894204_Corchorus_capsularis_1954311-1953831
PP861157_Hopea_chinensis_45943-46416
MT759787_Daphne_feddei_55157-55630
Arabidopsis_53634-54109_cpDNA
Brassica_52511-52989_cpDNA
OQ376546_Thlaspi_arvense_159793-160259
OY997597_Erodium_maritimum_169597-170076
KP940515_Geranium_maderense_552443-551984
ON729993_Santalum_album_61647-62120
OP588116_Sedum_plumbizincicola_102671-103128
OQ672472_Kalanchoe_daigremontiana_6651-7127
OP573219_Rhodiola_tangutica_183700-183227
OZ124158_Buxus_sempervirens_289586-290066
Nelumbo_60254-60727_cpDNA
OL979136_Pulsatilla_cernua_676568-676095
MT759793_Clematis_glauca_54032-54505
MT759767_Delphinium_yunnanense_54256-54736
MT759795_Aconitum_scaposum_57529-58002
OP620791_Ranunculus_cassubicifolius_932903-933383
OP466716_Coptis_chinensis_209951-209471
OY747138_Berberis_vulgaris_115975-115510
EU365401_Bambusa_oldhamii_10063-10536
Bambusa_55330-55803_cpDNA
PP885439_Aegilops_umbellulata_268773-269235
OQ086977_Elymus_magellanicus_498278-497816
Oryza_52738-53193_cpDNA
LC697740_Oryza_sativa_140622-141077
OZ228980_Leersia_oryzoides_15995-16450
PP035757_Zizania_latifolia_287578-287123
MZ506736_Sorghum_bicolor_148501-148029
Sorghum_56365-56837_cpDNA
OP210277_Zea_mays_169405-169877
MT471098_Coix_lacryma-jobi_658087-658559
PP665701_Miscanthus_sinensis_156937-157409
PP664563_Tripidium_arundinaceum_105146-105618
MN635785_Chrysopogon_zizanioides_243921-244405
PP372606_Eremochloa_ophiuroides_95711-96195
PQ576941_Nassella_tenuissima_13293-12814
MZ161092_Stipa_capillata_38508-38029
PP534167_Hygroryza_aristata_531553-532023
Phoenix_55625-56098_cpDNA
JACMSC010000023_Zingiber_officinale_1788417-1788871
PV236121_Curcuma_wenyujin_475248-475706
Lilium_51598-52070_cpDNA
PQ221457_Puya_raimondii_524700-524227
PP436815_Aechmea_fasciata_882855-883328
PP533607_Ananas_comosus_537443-536974
PP050468_Elaeis_guineensis_123860-123387
OZ184663_Musa_banksii_4887826-4888242
KJ501971_Erycina_pusilla_18575-19055
OQ024451_Cymbidium_lancifolium_15070-15543
OR413920_Dendrobium_officinale_25347-24874
PQ180340_Phalaenopsis_mannii_9970-10443
PQ859845_Bletilla_striata_83007-83480
MW563081_Stelis_montserratii_4617-5090
KJ501948_Vanilla_planifolia_590-1040
OQ716450_Galeola_lindleyana_5698-5274
OP465209_Paphiopedilum_micranthum_7342-6920
KJ501922_Cypripedium_formosanum_1781-1320
OL799325_Rhynchospora_pubera_758979-758525
MT240856_Allium_cepa_206416-206889
MZ726803_Hemerocallis_citrina_70948-70476
OL804177_Crocus_sativus_245072-245545
PP692488_Stemona_sessilifolia_48528-48055
PQ031343_Scheuchzeria_palustris_202692-203172
KX808392_Zostera_marina_87910-88389
OR066318_Acorus_tatarinowii_157091-157555
OR830326_Trichopus_zeylanicus_353701-354087
OQ390040_Zantedeschia_aethiopica_551403-551853
OR643818_Chloranthus_spicatus_739099-739572
OR176985_Hedyosmum_orientale_350105-350555
OK077159_Kadsura_japonica_274044-274510
MH748549_Schisandra_sphenanthera_407431-407897
KF754800_Amborella_trichopoda_124160-123696
OP649462_Aristolochia_manshuriensis_3332-2860
PQ899170_Cryptocarya_kwangtungensis_182514-182987
OR987149_Caryodaphnopsis_henryi_572938-573411
ON023262_Hernandia_nymphaeifolia_505413-504940
Liriodendron_57400-57873_cpDNA
0.06

**Sapria himalayana individual 1**
**MTPT chromosome 14 (6,184-8,547)**

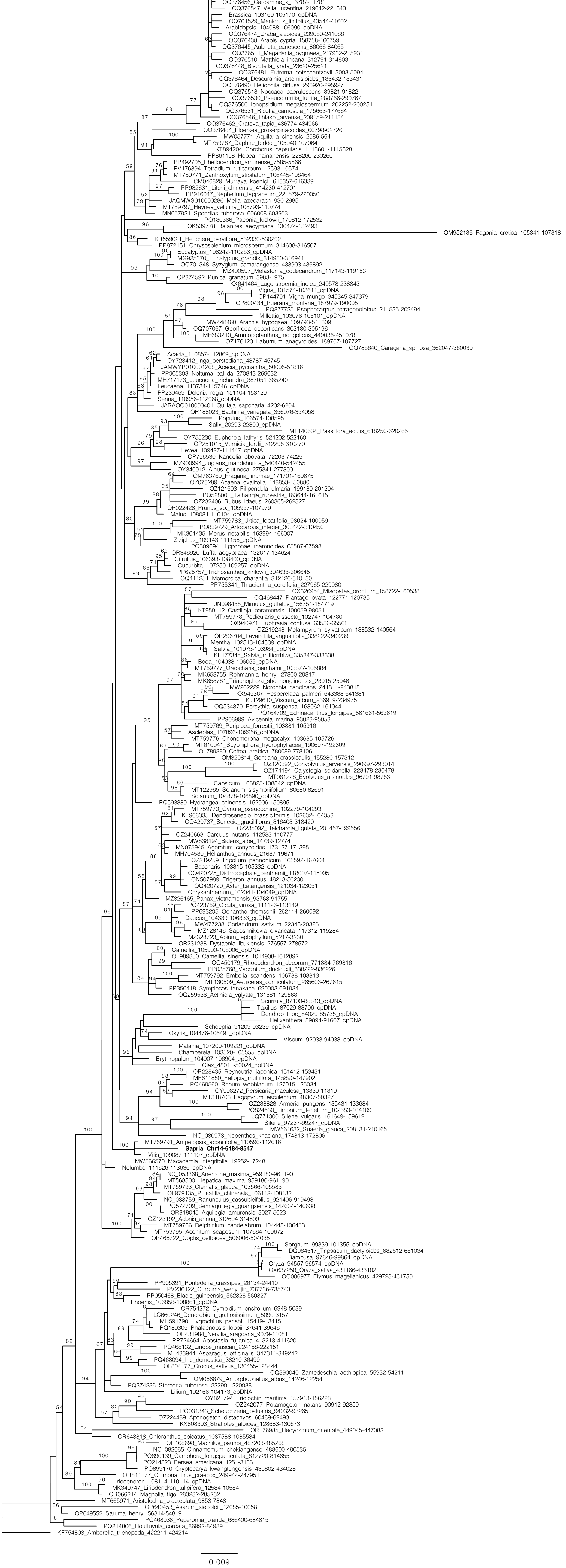

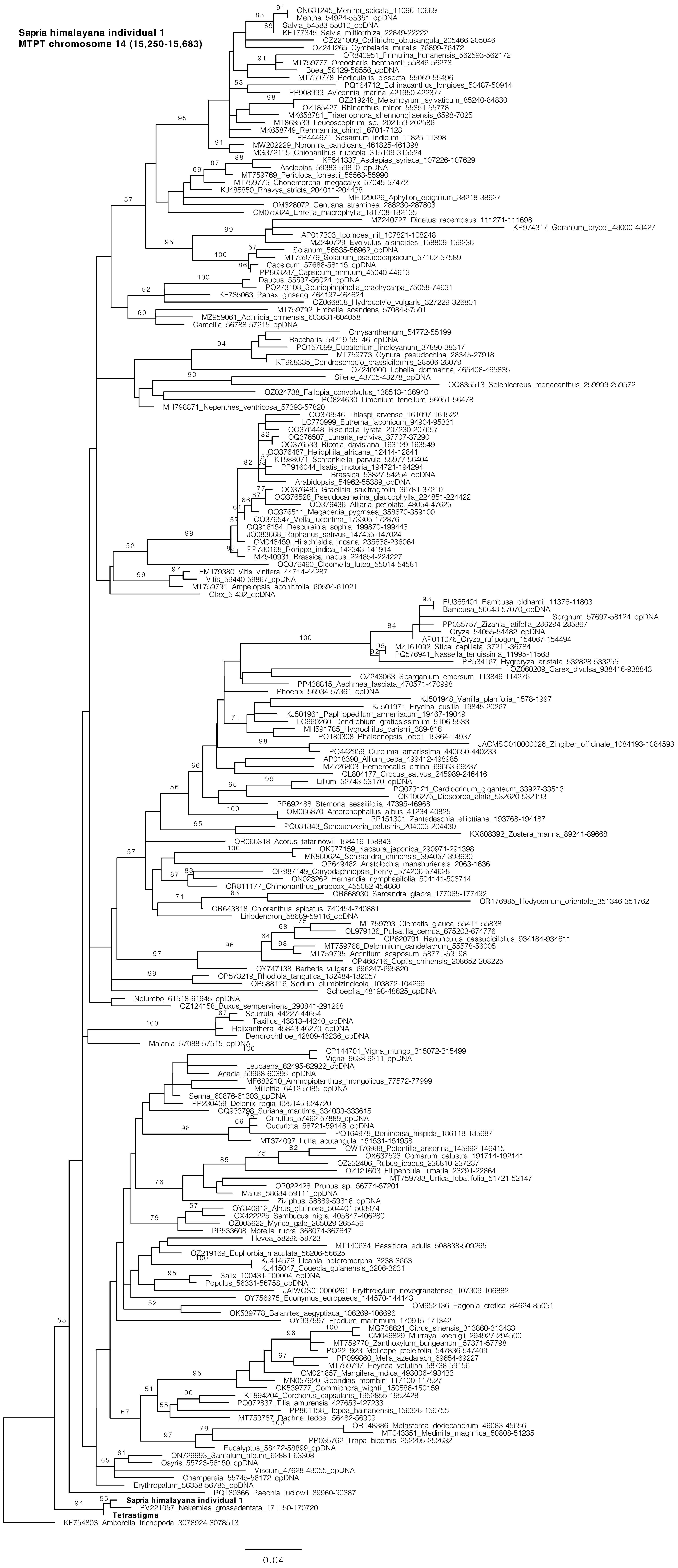

Sapria himalayana individual 1
MTPT chromosome 14 (15,250-15,683)
ON631245_Mentha_spicata_11096-10669
Mentha_54924-55351_cpDNA
Salvia_54583-55010_cpDNA
KF177345_Salvia_miltiorrhiza_22649-22222
OZ221009_Callitriche_obtusangula_205466-205046
OZ241265_Cymbalaria_muralis_76899-76472
OR840951_Primulina_hunanensis_562593-562172
MT759777_Oreocharis_benthamii_55846-56273
Boea_56129-56556_cpDNA
MT759778_Pedicularis_dissecta_55069-55496
PQ164712_Echinacanthus_longipes_50487-50914
PP908999_Avicennia_marina_421950-422377
OZ219248_Melampyrum_sylvaticum_85240-84830
OZ185427_Rhinanthus_minor_55351-55778
MK658781_Triaenophora_shennongjiaensis_6598-7025
MT863539_Leucosceptrum_sp._202159-202586
MK658749_Rehmannia_chingii_6701-7128
PP444671_Sesamum_indicum_11825-11398
MW202229_Noronhia_candicans_461825-461398
MG372115_Chionanthus_rupicola_315109-315524
KF541337_Asclepias_syriaca_107226-107629
Asclepias_59383-59810_cpDNA
MT759769_Periploca_forrestii_55563-55990
MT759775_Chonemorpha_megacalyx_57045-57472
KJ485850_Rhazya_stricta_204011-204438
MH129026_Aphyllon_epigalium_38218-38627
OM328072_Gentiana_straminea_288230-287803
CM075824_Ehretia_macrophylla_181708-182135
MZ240727_Dinetus_racemosus_111271-111698
KP974317_Geranium_brycei_48000-48427
AP017303_Ipomoea_nil_107821-108248
MZ240729_Evolvulus_alsinoides_158809-159236
Solanum_56535-56962_cpDNA
MT759779_Solanum_pseudocapsicum_57162-57589
Capsicum_57688-58115_cpDNA
PP863287_Capsicum_annuum_45040-44613
Daucus_55597-56024_cpDNA
PQ273108_Spuriopimpinella_brachycarpa_75058-74631
KF735063_Panax_ginseng_464197-464624
OZ066808_Hydrocotyle_vulgaris_327229-326801
MT759792_Embelia_scandens_57084-57501
MZ959061_Actinidia_chinensis_603631-604058
Camellia_56788-57215_cpDNA
Chrysanthemum_54772-55199
Baccharis_54719-55146_cpDNA
PQ157699_Eupatorium_lindleyanum_37890-38317
MT759773_Gynura_pseudochina_28345-27918
KT968335_Dendrosenecio_brassiciformis_28506-28079
OZ240900_Lobelia_dortmanna_465408-465835
Silene_43705-43278_cpDNA
OQ835513_Selenicereus_monacanthus_259999-259572
OZ024738_Fallopia_convolvulus_136513-136940
PQ824630_Limonium_tenellum_56051-56478
MH798871_Nepenthes_ventricosa_57393-57820
OQ376546_Thlaspi_arvense_161097-161522
LC770999_Eutrema_japonicum_94904-95331
OQ376448_Biscutella_lyrata_207230-207657
OQ376507_Lunaria_rediviva_37707-37290
OQ376533_Ricotia_davisiana_163129-163549
OQ376487_Heliophila_africana_12414-12841
KT988071_Schrenkiella_parvula_55977-56404
PP916044_Isatis_tinctoria_194721-194294
Brassica_53827-54254_cpDNA
Arabidopsis_54962-55389_cpDNA
OQ376485_Graellsia_saxifragifolia_36781-37210
OQ376528_Pseudocamelina_glaucophylla_224851-224422
OQ376436_Alliaria_petiolata_48054-47625
OQ376511_Megadenia_pygmaea_358670-359100
OQ376547_Vella_lucentina_173305-172876
OQ916154_Descurainia_sophia_199870-199443
JQ083668_Raphanus_sativus_147455-147024
CM048459_Hirschfeldia_incana_235636-236064
PP780168_Rorippa_indica_142343-141914
MZ540931_Brassica_napus_224654-224227
OQ376460_Cleomella_lutea_55014-54581
FM179380_Vitis_vinifera_44714-44287
Vitis_59440-59867_cpDNA
MT759791_Ampelopsis_aconitifolia_60594-61021
Olax_5-432_cpDNA
EU365401_Bambusa_oldhamii_11376-11803
Bambusa_56643-57070_cpDNA
Sorghum_57697-58124_cpDNA
PP035757_Zizania_latifolia_286294-285867
Oryza_54055-54482_cpDNA
AP011076_Oryza_rufipogon_154067-154494
MZ161092_Stipa_capillata_37211-36784
PQ576941_Nassella_tenuissima_11995-11568
PP534167_Hygroryza_aristata_532828-533255
OZ060209_Carex_divulsa_938416-938843
OZ243063_Sparganium_emersum_113849-114276
PP436815_Aechmea_fasciata_470571-470998
Phoenix_56934-57361_cpDNA
KJ501948_Vanilla_planifolia_1578-1997
KJ501971_Erycina_pusilla_19845-20267
KJ501961_Paphiopedilum_armeniacum_19467-19049
LC660260_Dendrobium_gratiosissimum_5106-5533
MH591785_Hygrochilus_parishii_389-816
PQ180308_Phalaenopsis_lobbii_15364-14937
JACMSC010000026_Zingiber_officinale_1084193-1084593
PQ442959_Curcuma_amarissima_440650-440233
AP018390_Allium_cepa_499412-498985
MZ726803_Hemerocallis_citrina_69663-69237
OL804177_Crocus_sativus_245989-246416
Lilium_52743-53170_cpDNA
PQ073121_Cardiocrinum_giganteum_33927-33513
OK106275_Dioscorea_alata_532620-532193
PP692488_Stemona_sessilifolia_47395-46968
OM066870_Amorphophallus_albus_41234-40825
PP151301_Zantedeschia_elliottiana_193768-194187
PQ031343_Scheuchzeria_palustris_204003-204430
KX808392_Zostera_marina_89241-89668
OR066318_Acorus_tatarinowii_158416-158843
OK077159_Kadsura_japonica_290971-291398
MK860624_Schisandra_chinensis_394057-393630
OP649462_Aristolochia_manshuriensis_2063-1636
OR987149_Caryodaphnopsis_henryi_574206-574628
ON023262_Hernandia_nymphaeifolia_504141-503714
OR811177_Chimonanthus_praecox_455082-454660
OR668930_Sarcandra_glabra_177065-177492
OR176985_Hedyosmum_orientale_351346-351762
OR643818_Chloranthus_spicatus_740454-740881
Liriodendron_58689-59116_cpDNA
MT759793_Clematis_glauca_55411-55838
OL979136_Pulsatilla_cernua_675203-674776
OP620791_Ranunculus_cassubicifolius_934184-934611
MT759766_Delphinium_candelabrum_55578-56005
MT759795_Aconitum_scaposum_58771-59198
OP466716_Coptis_chinensis_208652-208225
OY747138_Berberis_vulgaris_696247-695820
OP573219_Rhodiola_tangutica_182484-182057
OP588116_Sedum_plumbizincicola_103872-104299
Schoepfia_48198-48625_cpDNA
Nelumbo_61518-61945_cpDNA
OZ124158_Buxus_sempervirens_290841-291268
Scurrula_44227-44654
Taxillus_43813-44240_cpDNA
Helixanthera_45843-46270_cpDNA
Dendrophthoe_42809-43236_cpDNA
Malania_57088-57515_cpDNA
CP144701_Vigna_mungo_315072-315499
Vigna_9638-9211_cpDNA
Leucaena_62495-62922_cpDNA
Acacia_59968-60395_cpDNA
MF683210_Ammopiptanthus_mongolicus_77572-77999
Millettia_6412-5985_cpDNA
Senna_60876-61303_cpDNA
PP230459_Delonix_regia_625145-624720
OQ933798_Suriana_maritima_334033-333615
Citrullus_57462-57889_cpDNA
Cucurbita_58721-59148_cpDNA
PQ164978_Benincasa_hispida_186118-185687
MT374097_Luffa_acutangula_151531-151958
OW176988_Potentilla_anserina_145992-146415
OX637593_Comarum_palustre_191714-192141
OZ232406_Rubus_idaeus_236810-237237
OZ121603_Filipendula_ulmaria_23291-22864
MT759783_Urtica_lobatifolia_51721-52147
OP022428_Prunus_sp._56774-57201
Malus_58684-59111_cpDNA
Ziziphus_58889-59316_cpDNA
OY340912_Alnus_glutinosa_504401-503974
OX422225_Sambucus_nigra_405847-406280
OZ005622_Myrica_gale_265029-265456
PP533608_Morella_rubra_368074-367647
Hevea_58296-58723
MT140634_Passiflora_edulis_508838-509265
OZ219169_Euphorbia_maculata_56206-56625
KJ414572_Licania_heteromorpha_3238-3663
KJ415047_Couepia_guianensis_3206-3631
Salix_100431-100004_cpDNA
Populus_56331-56758_cpDNA
JAIWQS010000261_Erythroxylum_novogranatense_107309-106882
OY756975_Euonymus_europaeus_144570-144143
OM952136_Fagonia_cretica_84624-85051
OK539778_Balanites_aegyptiaca_106269-106696
OY997597_Erodium_maritimum_170915-171342
MG736621_Citrus_sinensis_313860-313433
CM046829_Murraya_koenigii_294927-294500
MT759770_Zanthoxylum_bungeanum_57371-57798
PQ221923_Melicope_pteleifolia_547836-547409
PP099860_Melia_azedarach_69654-69227
MT759797_Heynea_velutina_58738-59156
CM021857_Mangifera_indica_493006-493433
MN057920_Spondias_mombin_117100-117527
OK539777_Commiphora_wightii_150586-150159
KT894204_Corchorus_capsularis_1952855-1952428
PQ072837_Tilia_amurensis_427653-427233
PP861158_Hopea_hainanensis_156328-156755
MT759787_Daphne_feddei_56482-56909
OR148386_Melastoma_dodecandrum_46083-45656
MT043351_Medinilla_magnifica_50808-51235
PP035762_Trapa_bicornis_252205-252632
Eucalyptus_58472-58899_cpDNA
ON729993_Santalum_album_62881-63308
Osyris_55723-56150_cpDNA
Viscum_47628-48055_cpDNA
Champereia_55745-56172_cpDNA
Erythropalum_56358-56785_cpDNA
PQ180366_Paeonia_ludlowii_89960-90387
Sapria himalayana individual 1
PV221057_Nekemias_grossedentata_171150-170720
Tetrastigma
KF754803_Amborella_trichopoda_3078924-3078513
0.04

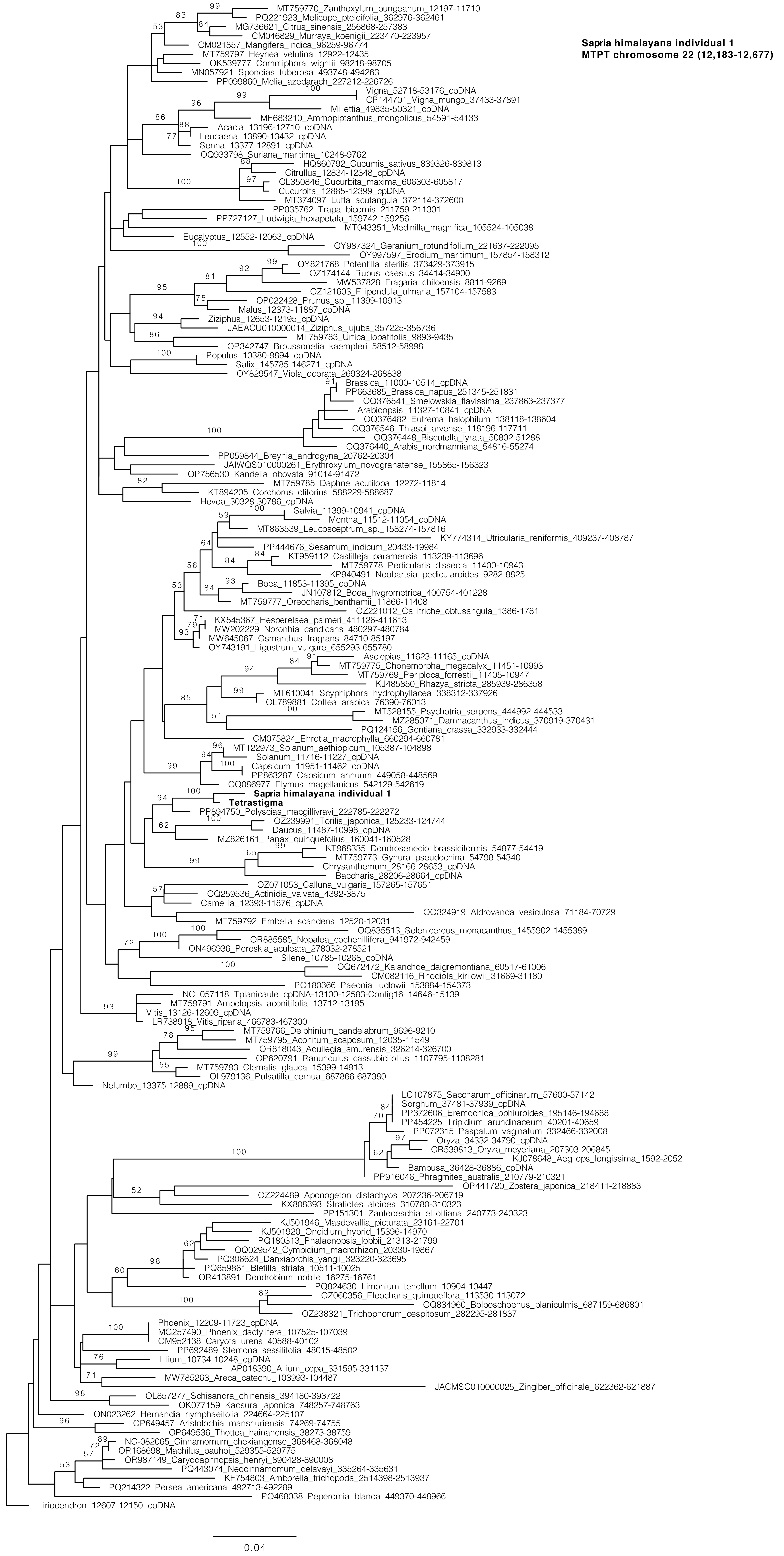
Sapria himalayana individual 1
MTPT chromosome 22 (12,183-12,677)
MT759770_Zanthoxylum_bungeanum_12197-11710
PQ221923_Melicope_pteleifolia_362976-362461
MG736621_Citrus_sinensis_256868-257383
CM046829_Murraya_koenigii_223470-223957
CM021857_Mangifera_indica_96259-96774
MT759797_Heynea_velutina_12922-12435
OK539777_Commiphora_wightii_98218-98705
MN057921_Spondias_tuberosa_493748-494263
PP099860_Melia_azedarach_227212-226726
Vigna_52718-53176_cpDNA
CP144701_Vigna_mungo_37433-37891
Millettia_49835-50321_cpDNA
MF683210_Ammopiptanthus_mongolicus_54591-54133
Acacia_13196-12710_cpDNA
Leucaena_13890-13432_cpDNA
Senna_13377-12891_cpDNA
OQ933798_Suriana_maritima_10248-9762
HQ860792_Cucumis_sativus_839326-839813
Citrullus_12834-12348_cpDNA
OL350846_Cucurbita_maxima_606303-605817
Cucurbita_12885-12399_cpDNA
MT374097_Luffa_acutangula_372114-372600
PP035762_Trapa_bicornis_211759-211301
PP727127_Ludwigia_hexapetala_159742-159256
MT043351_Medinilla_magnifica_105524-105038
Eucalyptus_12552-12063_cpDNA
OY987324_Geranium_rotundifolium_221637-222095
OY997597_Erodium_maritimum_157854-158312
OY821768_Potentilla_sterilis_373429-373915
OZ174144_Rubus_caesius_34414-34900
MW537828_Fragaria_chiloensis_8811-9269
OZ121603_Filipendula_ulmaria_157104-157583
OP022428_Prunus_sp._11399-10913
Malus_12373-11887_cpDNA
Ziziphus_12653-12195_cpDNA
JAEACU010000014_Ziziphus_jujuba_357225-356736
MT759783_Urtica_lobatifolia_9893-9435
OP342747_Broussonetia_kaempferi_58512-58998
Populus_10380-9894_cpDNA
Salix_145785-146271_cpDNA
OY829547_Viola_odorata_269324-268838
Brassica_11000-10514_cpDNA
PP663685_Brassica_napus_251345-251831
OQ376541_Smelowskia_flavissima_237863-237377
Arabidopsis_11327-10841_cpDNA
OQ376482_Eutrema_halophilum_138118-138604
OQ376546_Thlaspi_arvense_118196-117711
OQ376448_Biscutella_lyrata_50802-51288
OQ376440_Arabis_nordmanniana_54816-55274
PP059844_Breynia_androgyna_20762-20304
JAIWQS010000261_Erythroxylum_novogranatense_155865-156323
OP756530_Kandelia_obovata_91014-91472
MT759785_Daphne_acutiloba_12272-11814
KT894205_Corchorus_olitorius_588229-588687
Hevea_30328-30786_cpDNA
Salvia_11399-10941_cpDNA
Mentha_11512-11054_cpDNA
MT863539_Leucosceptrum_sp._158274-157816
KY774314_Utricularia_reniformis_409237-408787
PP444676_Sesamum_indicum_20433-19984
KT959112_Castilleja_paramensis_113239-113696
MT759778_Pedicularis_dissecta_11400-10943
KP940491_Neobartsia_pedicularoides_9282-8825
Boea_11853-11395_cpDNA
JN107812_Boea_hygrometrica_400754-401228
MT759777_Oreocharis_benthamii_11866-11408
OZ221012_Callitriche_obtusangula_1386-1781
KX545367_Hesperelaea_palmeri_411126-411613
MW202229_Noronhia_candicans_480297-480784
MW645067_Osmanthus_fragrans_84710-85197
OY743191_Ligustrum_vulgare_655293-655780
Asclepias_11623-11165_cpDNA
MT759775_Chonemorpha_megacalyx_11451-10993
MT759769_Periploca_forrestii_11405-10947
KJ485850_Rhazya_stricta_285939-286358
MT610041_Scyphiphora_hydrophyllacea_338312-337926
OL789881_Coffea_arabica_76390-76013
MT528155_Psychotria_serpens_444992-444533
MZ285071_Damnacanthus_indicus_370919-370431
PQ124156_Gentiana_crassa_332933-332444
CM075824_Ehretia_macrophylla_660294-660781
MT122973_Solanum_aethiopicum_105387-104898
Solanum_11716-11227_cpDNA
Capsicum_11951-11462_cpDNA
PP863287_Capsicum_annuum_449058-448569
OQ086977_Elymus_magellanicus_542129-542619
Sapria himalayana individual 1
Tetrastigma
PP894750_Polyscias_macgillivrayi_222785-222272
OZ239991_Torilis_japonica_125233-124744
Daucus_11487-10998_cpDNA
MZ826161_Panax_quinquefolius_160041-160528
KT968335_Dendrosenecio_brassiciformis_54877-54419
MT759773_Gynura_pseudochina_54798-54340
Chrysanthemum_28166-28653_cpDNA
Baccharis_28206-28664_cpDNA
OZ071053_Calluna_vulgaris_157265-157651
OQ259536_Actinidia_valvata_4392-3875
Camellia_12393-11876_cpDNA
OQ324919_Aldrovanda_vesiculosa_71184-70729
MT759792_Embelia_scandens_12520-12031
OQ835513_Selenicereus_monacanthus_1455902-1455389
OR885585_Nopalea_cochenillifera_941972-942459
ON496936_Pereskia_aculeata_278032-278521
Silene_10785-10268_cpDNA
OQ672472_Kalanchoe_daigremontiana_60517-61006
CM082116_Rhodiola_kirilowii_31669-31180
PQ180366_Paeonia_ludlowii_153884-154373
NC_057118_Tplanicaule_cpDNA-13100-12583-Contig16_14646-15139
MT759791_Ampelopsis_aconitifolia_13712-13195
Vitis_13126-12609_cpDNA
LR738918_Vitis_riparia_466783-467300
MT759766_Delphinium_candelabrum_9696-9210
MT759795_Aconitum_scaposum_12035-11549
OR818043_Aquilegia_amurensis_326214-326700
OP620791_Ranunculus_cassubicifolius_1107795-1108281
MT759793_Clematis_glauca_15399-14913
OL979136_Pulsatilla_cernua_687866-687380
Nelumbo_13375-12889_cpDNA
LC107875_Saccharum_officinarum_57600-57142
Sorghum_37481-37939_cpDNA
PP372606_Eremochloa_ophiuroides_195146-194688
PP454225_Tripidium_arundinaceum_40201-40659
PP072315_Paspalum_vaginatum_332466-332008
Oryza_34332-34790_cpDNA
OR539813_Oryza_meyeriana_207303-206845
KJ078648_Aegilops_longissima_1592-2052
Bambusa_36428-36886_cpDNA
PP916046_Phragmites_australis_210779-210321
OP441720_Zostera_japonica_218411-218883
OZ224489_Aponogeton_distachyos_207236-206719
KX808393_Stratiotes_aloides_310780-310323
PP151301_Zantedeschia_elliottiana_240773-240323
KJ501946_Masdevallia_picturata_23161-22701
KJ501920_Oncidium_hybrid_15396-14970
PQ180313_Phalaenopsis_lobbii_21313-21799
OQ029542_Cymbidium_macrorhizon_20330-19867
PQ306624_Danxiaorchis_yangii_323220-323695
PQ859861_Bletilla_striata_10511-10025
OR413891_Dendrobium_nobile_16275-16761
PQ824630_Limonium_tenellum_10904-10447
OZ060356_Eleocharis_quinqueflora_113530-113072
OQ834960_Bolboschoenus_planiculmis_687159-686801
OZ238321_Trichophorum_cespitosum_282295-281837
Phoenix_12209-11723_cpDNA
MG257490_Phoenix_dactylifera_107525-107039
OM952138_Caryota_urens_40588-40102
PP692489_Stemona_sessilifolia_48015-48502
Lilium_10734-10248_cpDNA
AP018390_Allium_cepa_331595-331137
MW785263_Areca_catechu_103993-104487
JACMSC010000025_Zingiber_officinale_622362-621887
OL857277_Schisandra_chinensis_394180-393722
OK077159_Kadsura_japonica_748257-748763
ON023262_Hernandia_nymphaeifolia_224664-225107
OP649457_Aristolochia_manshuriensis_74269-74755
OP649536_Thottea_hainanensis_38273-38759
NC-082065_Cinnamomum_chekiangense_368468-368048
OR168698_Machilus_pauhoi_529355-529775
OR987149_Caryodaphnopsis_henryi_890428-890008
PQ443074_Neocinnamomum_delavayi_335264-335631
KF754803_Amborella_trichopoda_2514398-2513937
PQ214322_Persea_americana_492713-492289
PQ468038_Peperomia_blanda_449370-448966
Liriodendron_12607-12150_cpDNA
0.04

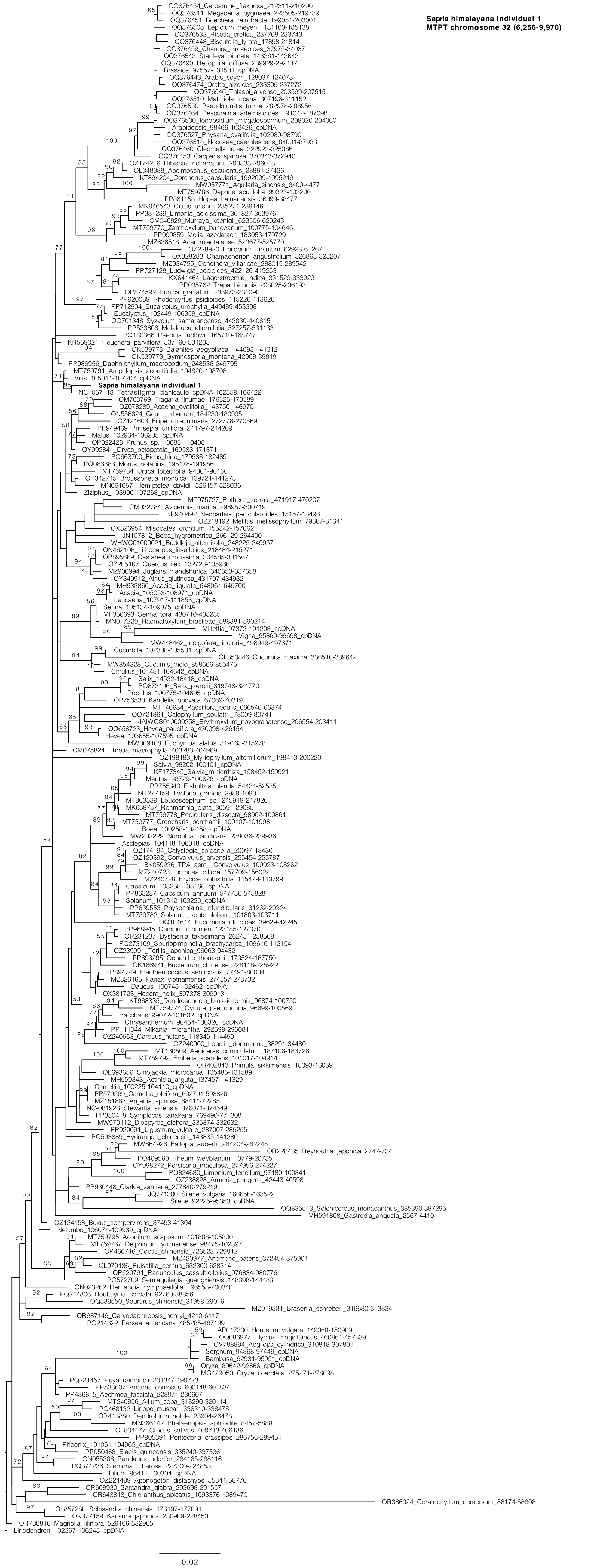

Sapria himalayana individual 1
MTPT chromosome 32 (6,256-9,970)
OQ376454_Cardamine_flexuosa_212311-210290
OQ376511_Megadenia_pygmaea_223505-219739
OQ376451_Boechera_retrofracta_199051-203001
OQ376505_Lepidium_meyenii_181183-185138
OQ376532_Ricotia_cretica_237708-233743
OQ376448_Biscutella_lyrata_17858-21814
OQ376459_Chamira_circaeoides_37975-34037
OQ376543_Stanleya_pinnata_146381-143643
OQ376490_Heliophila_diffusa_289929-292117
Brassica_97557-101501_cpDNA
OQ376443_Arabis_soyeri_128037-124073
OQ376474_Draba_aizoides_233305-237272
OQ376546_Thlaspi_arvense_203599-207515
OQ376510_Matthiola_incana_307196-311152
OQ376530_Pseudoturritis_turrita_282978-286956
OQ376464_Descurainia_artemisioides_191042-187098
OQ376500_Ionopsidium_megalospermum_208020-204060
Arabidopsis_98466-102426_cpDNA
OQ376527_Physaria_ovalifolia_102080-98790
OQ376518_Noccaea_caerulescens_84001-87933
OQ376460_Cleomella_lutea_322923-325386
OQ376453_Capparis_spinosa_370343-372940
OZ174216_Hibiscus_richardsonii_293833-296018
OL348388_Abelmoschus_esculentus_28861-27436
KT894204_Corchorus_capsularis_1992609-1995219
MW057771_Aquilaria_sinensis_8400-4477
MT759786_Daphne_acutiloba_99323-103200
PP861158_Hopea_hainanensis_36099-38477
MN946543_Citrus_unshiu_235271-239146
PP331239_Limonia_acidissima_361827-363976
CM046829_Murraya_koenigii_623506-620243
MT759770_Zanthoxylum_bungeanum_100775-104646
PP099859_Melia_azedarach_183053-179729
MZ636518_Acer_miaotaiense_523677-525770
OZ228920_Epilobium_hirsutum_62928-61267
OX328283_Chamaenerion_angustifolium_326868-325207
MZ934755_Oenothera_villaricae_288015-289542
PP727128_Ludwigia_peploides_422120-419253
KX641464_Lagerstroemia_indica_331529-333929
PP035762_Trapa_bicornis_208025-206193
OP874592_Punica_granatum_233973-231090
PP920089_Rhodomyrtus_psidioides_115226-113626
PP712904_Eucalyptus_urophylla_449489-453398
Eucalyptus_102449-106359_cpDNA
OQ701348_Syzygium_samarangense_443830-440815
PP533606_Melaleuca_alternifolia_527257-531133
PQ180366_Paeonia_ludlowii_165710-168747
KR559021_Heuchera_parviflora_537160-534203
OK539778_Balanites_aegyptiaca_144093-141312
OK539779_Gymnosporia_montana_42968-39819
PP986956_Daphniphyllum_macropodum_248536-249795
MT759791_Ampelopsis_aconitifolia_104820-108708
Vitis_105011-107207_cpDNA
Sapria himalayana individual 1
NC_057118_Tetrastigma_planicaule_cpDNA-102559-106422
OM763769_Fragaria_iinumae_176525-173589
OZ078289_Acaena_ovalifolia_143750-146970
ON556624_Geum_urbanum_184239-180995
OZ121603_Filipendula_ulmaria_272776-270569
PP949469_Prinsepia_uniflora_241797-244209
Malus_102964-106205_cpDNA
OP022428_Prunus_sp._100851-104081
OY992841_Dryas_octopetala_169583-171371
PQ663700_Ficus_hirta_179586-182489
PQ083383_Morus_notabilis_195178-191956
MT759784_Urtica_lobatifolia_94361-96156
OP342745_Broussonetia_monoica_139721-141273
MN061667_Hemiptelea_davidii_326157-328036
Ziziphus_103990-107268_cpDNA
MT075727_Rotheca_serrata_471917-470207
CM032784_Avicennia_marina_298957-300719
KP940492_Neobartsia_pedicularoides_15157-13496
OZ218192_Melittis_melissophyllum_79887-81641
OX326954_Misopates_orontium_155342-157062
JN107812_Boea_hygrometrica_266129-264400
WHWC01000021_Buddleja_alternifolia_248225-249957
ON462106_Lithocarpus_litseifolius_218484-215271
OP895669_Castanea_mollissima_304585-301567
OZ205167_Quercus_ilex_132723-135966
MZ900994_Juglans_mandshurica_340353-337658
OY340912_Alnus_glutinosa_431707-434932
MH933866_Acacia_ligulata_648061-645700
Acacia_105053-108971_cpDNA
Leucaena_107917-111853_cpDNA
Senna_105134-109075_cpDNA
MF358693_Senna_tora_430710-433285
MN017229_Haematoxylum_brasiletto_588381-590214
Millettia_97372-101203_cpDNA
Vigna_95860-99698_cpDNA
MW448462_Indigofera_tinctoria_498949-497371
Cucurbita_102308-105501_cpDNA
OL350846_Cucurbita_maxima_336510-339642
MW854328_Cucumis_melo_858666-855475
Citrullus_101451-104642_cpDNA
Salix_14532-18418_cpDNA
PQ873106_Salix_pierotii_319748-321770
Populus_100775-104695_cpDNA
OP756530_Kandelia_obovata_67069-70319
MT140634_Passiflora_edulis_666540-663741
OQ721861_Calophyllum_soulattri_78009-80741
JAIWQS010000258_Erythroxylum_novogranatense_206554-203411
OQ658723_Hevea_pauciflora_430098-426154
Hevea_103655-107595_cpDNA
MW009108_Euonymus_alatus_319163-315978
CM075824_Ehretia_macrophylla_403283-404969
OZ198183_Myriophyllum_alterniflorum_198413-200220
Salvia_98202-100101_cpDNA
KF177345_Salvia_miltiorrhiza_158452-159921
Mentha_98729-100628_cpDNA
PP755340_Elsholtzia_blanda_54434-52535
MT277159_Tectona_grandis_2989-1090
MT863539_Leucosceptrum_sp._245919-247826
MK658757_Rehmannia_elata_30591-29085
MT759778_Pedicularis_dissecta_98962-100861
MT759777_Oreocharis_benthamii_100107-101996
Boea_100258-102158_cpDNA
MW202229_Noronhia_candicans_238036-239936
Asclepias_104118-106018_cpDNA
OZ174194_Calystegia_soldanella_20097-18430
OZ120392_Convolvulus_arvensis_255454-253787
BK059236_TPA_asm__Convolvulus_109923-108262
MZ240723_Ipomoea_biflora_157709-156022
MZ240728_Erycibe_obtusifolia_115479-113799
Capsicum_103258-105166_cpDNA
PP863287_Capsicum_annuum_547736-545828
Solanum_101312-103220_cpDNA
PP639553_Physochlaina_infundibularis_31232-29324
MT759782_Solanum_septemlobum_101803-103711
OQ101614_Eucommia_ulmoides_39629-42245
PP968945_Cnidium_monnieri_123185-127070
OR231237_Dystaenia_takesimana_262451-258568
PQ273109_Spuriopimpinella_brachycarpa_109616-113154
OZ239991_Torilis_japonica_96063-94432
PP693295_Oenanthe_thomsonii_170524-167750
OK166971_Bupleurum_chinense_228118-225922
PP894749_Eleutherococcus_senticosus_77491-80004
MZ826165_Panax_vietnamensis_274857-278732
Daucus_100748-102462_cpDNA
OX381723_Hedera_helix_307378-309913
KT968335_Dendrosenecio_brassiciformis_96874-100750
MT759774_Gynura_pseudochina_96699-100569
Baccharis_99072-101602_cpDNA
Chrysanthemum_96454-100326_cpDNA
PP111044_Mikania_micrantha_292599-295081
OZ240663_Carduus_nutans_118345-114459
OZ240900_Lobelia_dortmanna_38291-34480
MT130509_Aegiceras_corniculatum_187106-183726
MT759792_Embelia_scandens_101017-104914
OR402843_Primula_sikkimensis_18093-16059
OL693656_Sinojackia_microcarpa_135485-131589
MH559343_Actinidia_arguta_137457-141329
Camellia_100225-104110_cpDNA
PP579569_Camellia_oleifera_602701-598826
MZ151883_Argania_spinosa_68411-72285
NC-081928_Stewartia_sinensis_376071-374549
PP350418_Symplocos_tanakana_769490-771308
MW970112_Diospyros_oleifera_335374-332632
PP920091_Ligustrum_vulgare_287007-285255
PQ593889_Hydrangea_chinensis_143835-141280
MW664926_Fallopia_aubertii_284204-282248
OR228435_Reynoutria_japonica_2747-734
PQ469560_Rheum_webbianum_18779-20735
OY998272_Persicaria_maculosa_277956-274227
PQ824630_Limonium_tenellum_97180-100341
OZ238828_Armeria_pungens_42443-40598
PP930448_Clarkia_xantiana_277840-279219
JQ771300_Silene_vulgaris_166656-163522
Silene_92225-95353_cpDNA
OQ835513_Selenicereus_monacanthus_385390-387295
MH591808_Gastrodia_angusta_2567-4410
OZ124158_Buxus_sempervirens_37453-41304
Nelumbo_106074-109939_cpDNA
MT759795_Aconitum_scaposum_101888-105800
MT759767_Delphinium_yunnanense_98475-102397
OP466716_Coptis_chinensis_726523-729912
MZ420977_Anemone_patens_372454-375901
OL979136_Pulsatilla_cernua_632300-628314
OP620791_Ranunculus_cassubicifolius_976834-980776
PQ572709_Semiaquilegia_guangxiensis_148398-144483
ON023262_Hernandia_nymphaeifolia_196558-200340
PQ214806_Houttuynia_cordata_92760-88856
OQ539550_Saururus_chinensis_31958-29016
MZ919331_Brasenia_schreberi_316630-313834
OR987149_Caryodaphnopsis_henryi_4210-6117
PQ214322_Persea_americana_485285-487199
AP017300_Hordeum_vulgare_149068-150909
OQ086977_Elymus_magellanicus_460861-457839
OV788894_Aegilops_cylindrica_310818-307801
Sorghum_94868-97449_cpDNA
Bambusa_92931-95951_cpDNA
Oryza_89642-92666_cpDNA
MG429050_Oryza_coarctata_275271-278098
PQ221457_Puya_raimondii_201347-199723
PP533607_Ananas_comosus_600148-601834
PP436815_Aechmea_fasciata_228971-230607
MT240856_Allium_cepa_318290-320114
PQ468132_Liriope_muscari_336310-338478
OR413880_Dendrobium_nobile_23904-26478
MN366142_Phalaenopsis_aphrodite_8457-5888
OL804177_Crocus_sativus_409713-406136
PP905391_Pontederia_crassipes_286756-289451
Phoenix_101061-104965_cpDNA
PP050468_Elaeis_guineensis_335240-337536
ON055386_Pandanus_odorifer_284165-288116
PQ374236_Stemona_tuberosa_227300-224853
Lilium_96411-100304_cpDNA
OZ224489_Aponogeton_distachyos_55841-58770
OR668930_Sarcandra_glabra_293698-291557
OR643818_Chloranthus_spicatus_1093376-1089470
OR366024_Ceratophyllum_demersum_86174-88808
OL857280_Schisandra_chinensis_173197-177091
OK077159_Kadsura_japonica_230909-228450
OR730816_Magnolia_liliiflora_529106-532965
Liriodendron_102367-106243_cpDNA
0.02

**Figure S9. Phylogenetic analyses of organellar repair, recombination, and replication (RRR) genes.** Maximum likelihood analyses were performed with RAxML. ML bootstrap values >50% are shown. The scale bar corresponds to substitutions per site.

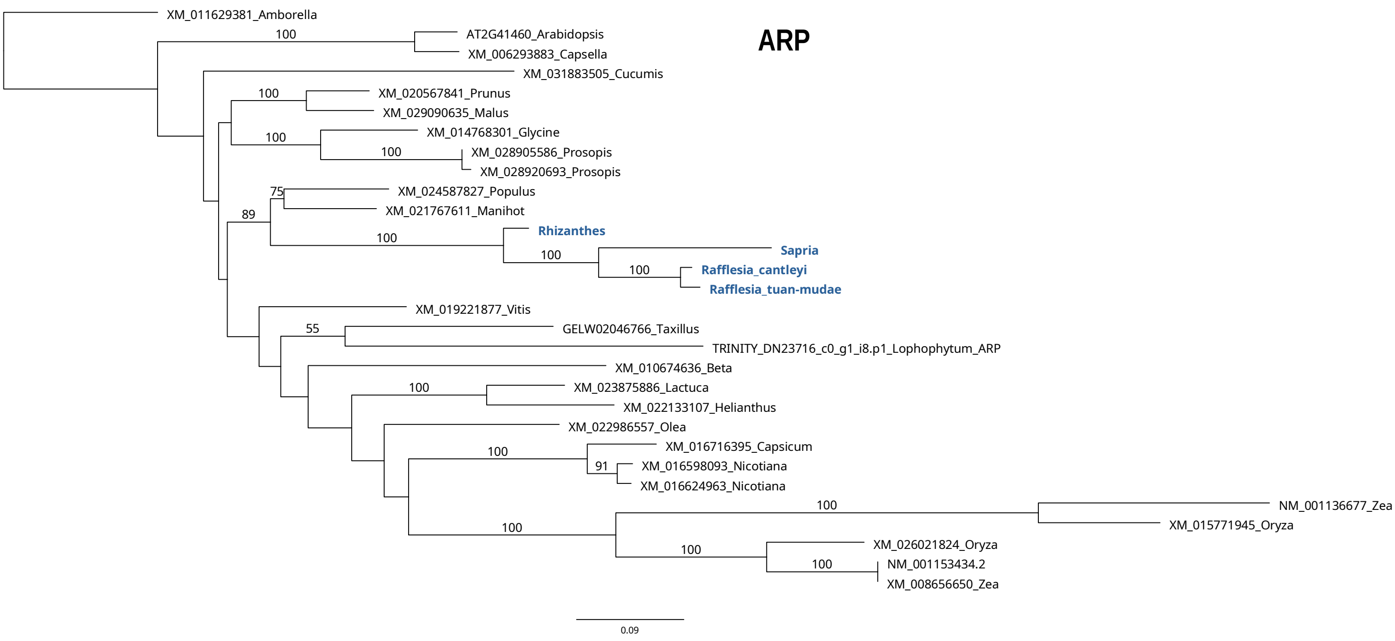


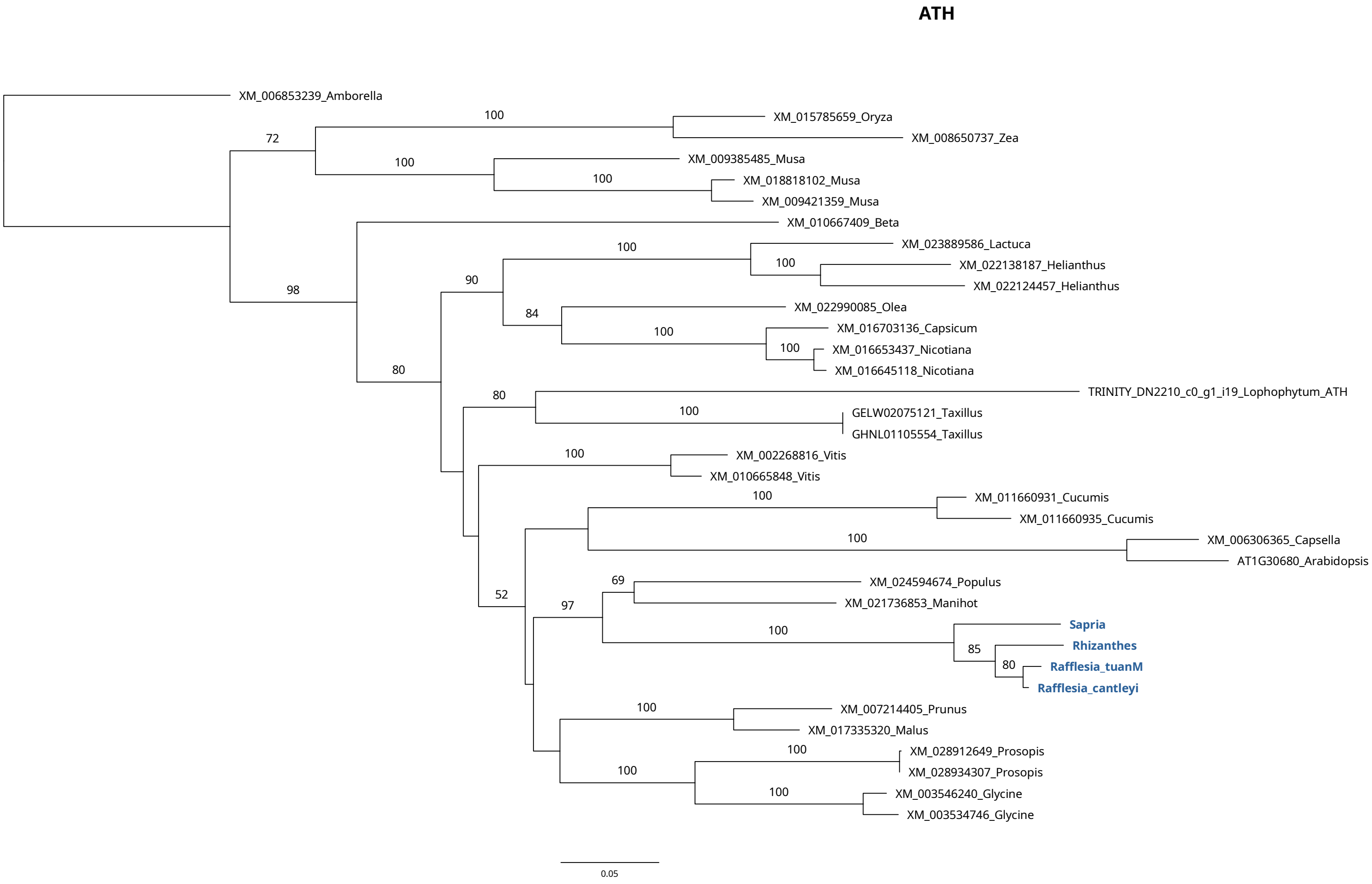

**FPG1**

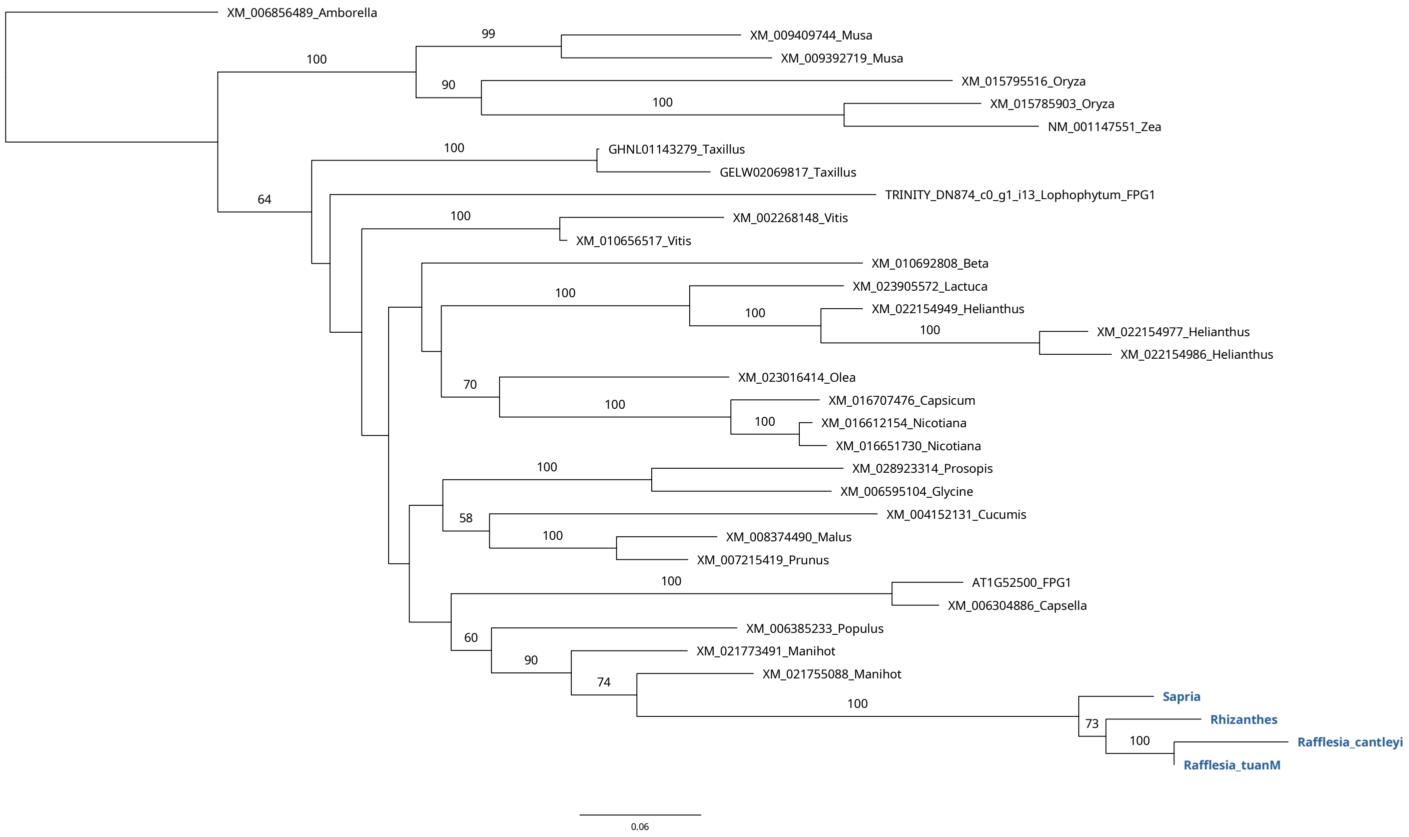
XM_006856489_Amborella
99
XM_009409744_Musa
100
XM_009392719_Musa
XM_015795516_Oryza
90
100
XM_015785903_Oryza
NM_001147551_Zea
100
GHNL01143279_Taxillus
GELW02069817_Taxillus
64
TRINITY_DN874_c0_g1_i13_Lophophytum_FPG1
100
XM_002268148_Vitis
XM_010656517_Vitis
XM_010692808_Beta
XM_023905572_Lactuca
100
100
XM_022154949_Helianthus
100
XM_022154977_Helianthus
XM_022154986_Helianthus
70
XM_023016414_Olea
100
XM_016707476_Capsicum
100
XM_016612154_Nicotiana
XM_016651730_Nicotiana
100
XM_028923314_Prosopis
XM_006595104_Glycine
58
XM_004152131_Cucumis
100
XM_008374490_Malus
XM_007215419_Prunus
100
AT1G52500_FPG1
XM_006304886_Capsella
XM_006385233_Populus
60
XM_021773491_Manihot
90
XM_021755088_Manihot
74
100
Sapria
Rhizanthes
73
100
Rafflesia_cantleyi
Rafflesia_tuanM
0.06

**LIG1**

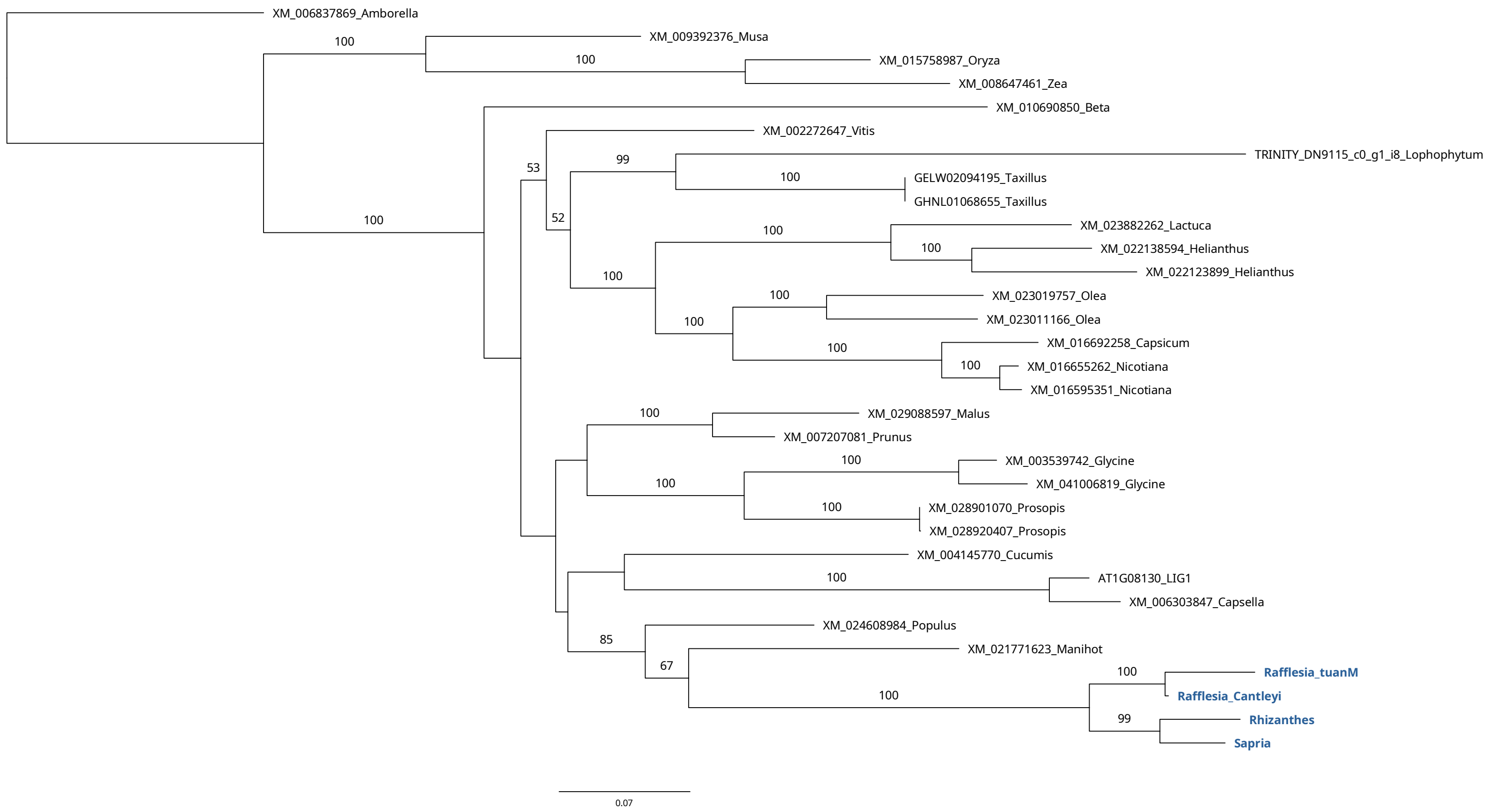
XM_006837869_Amborella
XM_009392376_Musa
100
100
XM_015758987_Oryza
XM_008647461_Zea
XM_010690850_Beta
XM_002272647_Vitis
53
99
TRINITY_DN9115_c0_g1_i8_Lophophytum
100
GELW02094195_Taxillus
GHNL01068655_Taxillus
100
52
100
XM_023882262_Lactuca
100
XM_022138594_Helianthus
XM_022123899_Helianthus
100
100
XM_023019757_Olea
XM_023011166_Olea
100
XM_016692258_Capsicum
100
XM_016655262_Nicotiana
XM_016595351_Nicotiana
100
XM_029088597_Malus
XM_007207081_Prunus
100
XM_003539742_Glycine
100
XM_041006819_Glycine
100
XM_028901070_Prosopis
XM_028920407_Prosopis
XM_004145770_Cucumis
100
AT1G08130_LIG1
XM_006303847_Capsella
XM_024608984_Populus
85
XM_021771623_Manihot
67
100
Rafflesia_tuanM
100
Rafflesia_Cantleyi
99
Rhizanthes
Sapria
0.07

#### GYRA

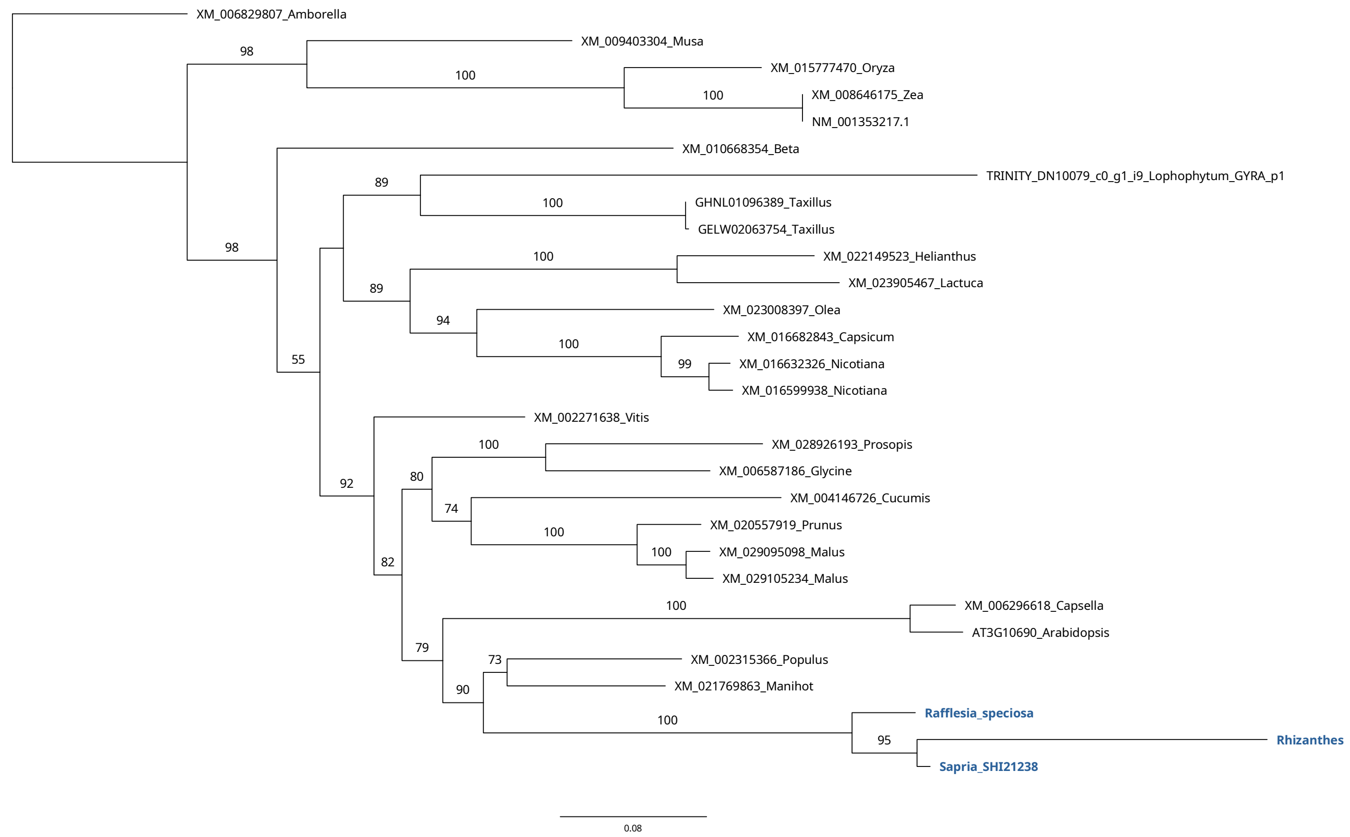
XM_006829807_Amborella
XM_009403304_Musa
XM_015777470_Oryza
XM_008646175_Zea
NM_001353217.1
XM_010668354_Beta
TRINITY_DN10079_c0_g1_i9_Lophophytum_GYRA_p1
GHNL01096389_Taxillus
GELW02063754_Taxillus
XM_022149523_Helianthus
XM_023905467_Lactuca
XM_023008397_Olea
XM_016682843_Capsicum
XM_016632326_Nicotiana
XM_016599938_Nicotiana
XM_002271638_Vitis
XM_028926193_Prosopis
XM_006587186_Glycine
XM_004146726_Cucumis
XM_020557919_Prunus
XM_029095098_Malus
XM_029105234_Malus
XM_006296618_Capsella
AT3G10690_Arabidopsis
XM_002315366_Populus
XM_021769863_Manihot
Rafflesia_speciosa
Rhizanthes
Sapria_SHI21238
0.08

#### GYRB

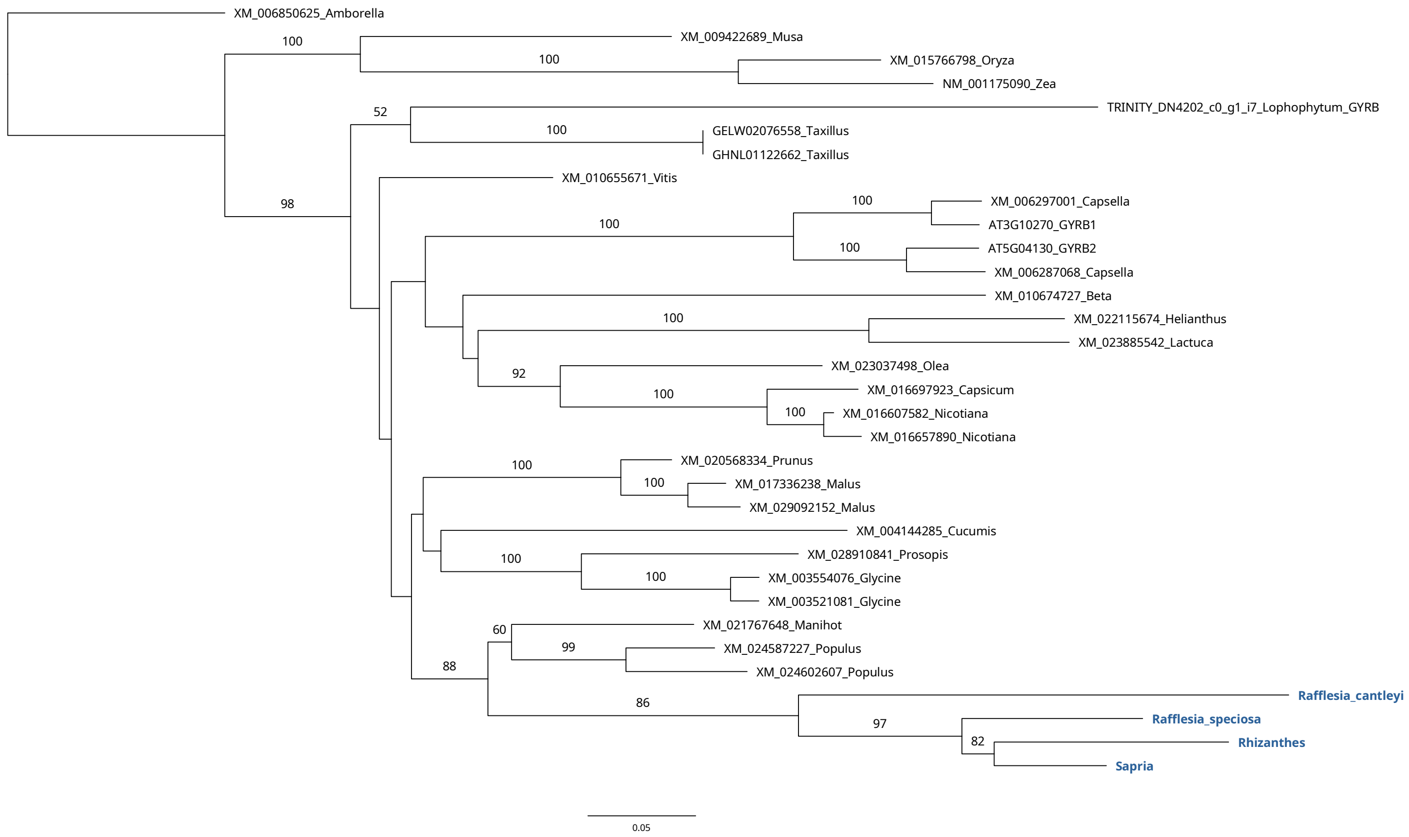
XM_006850625_Amborella
XM_009422689_Musa
XM_015766798_Oryza
NM_001175090_Zea
TRINITY_DN4202_c0_g1_i7_Lophophytum_GYRB
GELW02076558_Taxillus
GHNL01122662_Taxillus
XM_010655671_Vitis
XM_006297001_Capsella
AT3G10270_GYRB1
AT5G04130_GYRB2
XM_006287068_Capsella
XM_010674727_Beta
XM_022115674_Helianthus
XM_023885542_Lactuca
XM_023037498_Olea
XM_016697923_Capsicum
XM_016607582_Nicotiana
XM_016657890_Nicotiana
XM_020568334_Prunus
XM_017336238_Malus
XM_029092152_Malus
XM_004144285_Cucumis
XM_028910841_Prosopis
XM_003554076_Glycine
XM_003521081_Glycine
XM_021767648_Manihot
XM_024587227_Populus
XM_024602607_Populus
Rafflesia_cantleyi
Rafflesia_speciosa
Rhizanthes
Sapria
0.05

#### MSH1

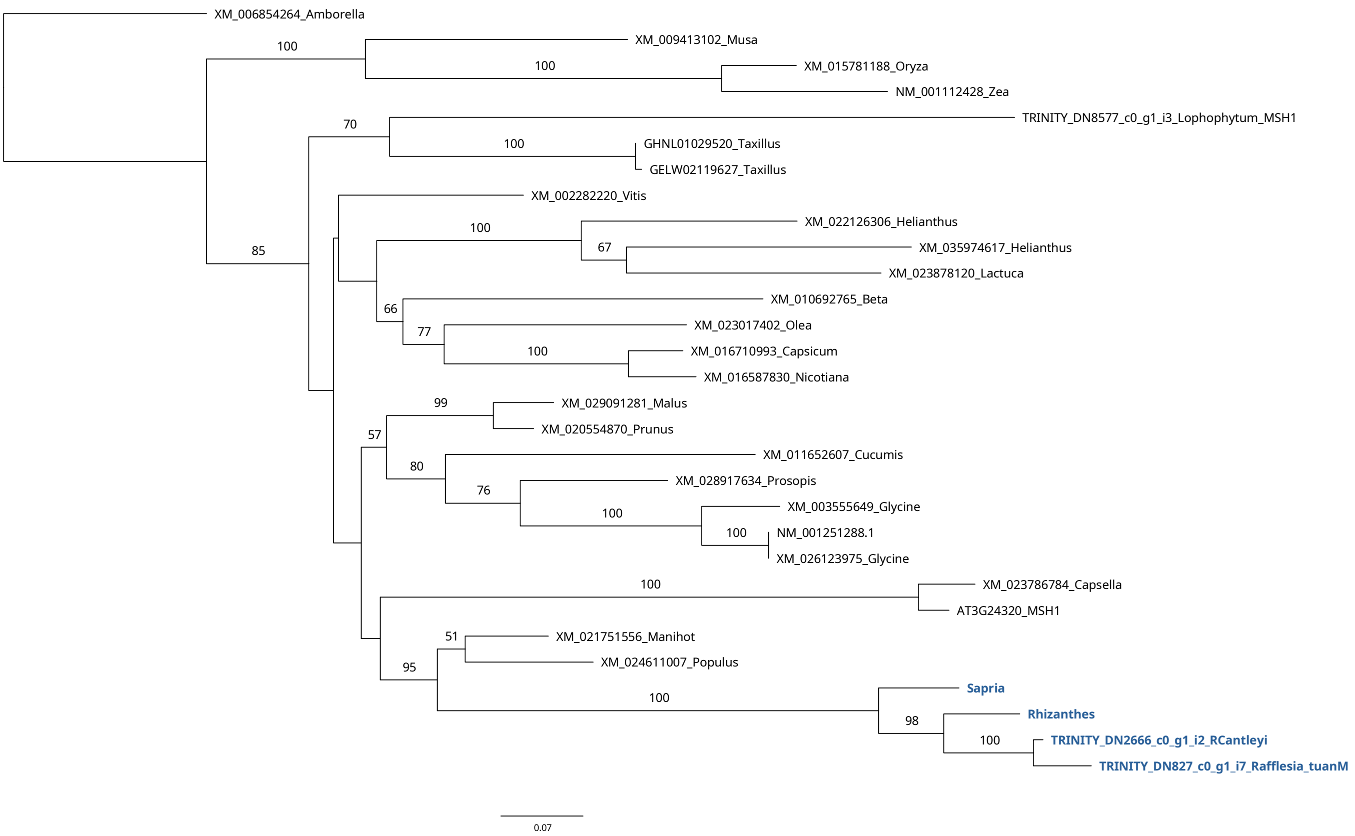
XM_006854264_Amborella
XM_009413102_Musa
XM_015781188_Oryza
NM_001112428_Zea
TRINITY_DN8577_c0_g1_i3_Lophophytum_MSH1
GHNL01029520_Taxillus
GELW02119627_Taxillus
XM_002282220_Vitis
XM_022126306_Helianthus
XM_035974617_Helianthus
XM_023878120_Lactuca
XM_010692765_Beta
XM_023017402_Olea
XM_016710993_Capsicum
XM_016587830_Nicotiana
XM_029091281_Malus
XM_020554870_Prunus
XM_011652607_Cucumis
XM_028917634_Prosopis
XM_003555649_Glycine
NM_001251288.1
XM_026123975_Glycine
XM_023786784_Capsella
AT3G24320_MSH1
XM_021751556_Manihot
XM_024611007_Populus
Sapria
Rhizanthes
TRINITY_DN2666_c0_g1_i2_RCantleyi
TRINITY_DN827_c0_g1_i7_Rafflesia_tuanM
100
100
70
100
85
100
67
66
77
100
99
57
80
76
100
100
100
51
95
100
98
100
0.07

#### NTH

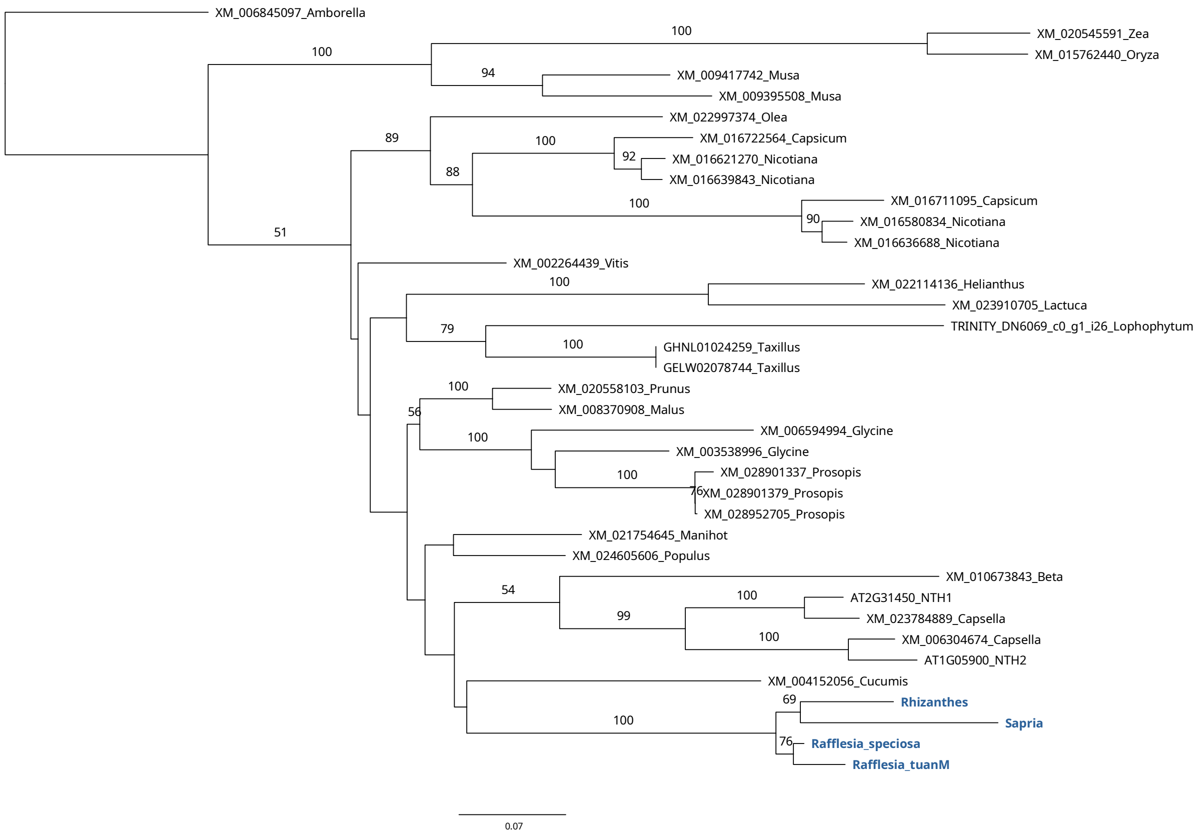
XM_006845097_Amborella
XM_020545591_Zea
XM_015762440_Oryza
XM_009417742_Musa
XM_009395508_Musa
XM_022997374_Olea
XM_016722564_Capsicum
XM_016621270_Nicotiana
XM_016639843_Nicotiana
XM_016711095_Capsicum
XM_016580834_Nicotiana
XM_016636688_Nicotiana
XM_002264439_Vitis
XM_022114136_Helianthus
XM_023910705_Lactuca
TRINITY_DN6069_c0_g1_i26_Lophophytum
GHNL01024259_Taxillus
GELW02078744_Taxillus
XM_020558103_Prunus
XM_008370908_Malus
XM_006594994_Glycine
XM_003538996_Glycine
XM_028901337_Prosopis
XM_028901379_Prosopis
XM_028952705_Prosopis
XM_021754645_Manihot
XM_024605606_Populus
XM_010673843_Beta
AT2G31450_NTH1
XM_023784889_Capsella
XM_006304674_Capsella
AT1G05900_NTH2
XM_004152056_Cucumis
Rhizanthes
Sapria
Rafflesia_speciosa
Rafflesia_tuanM
100
100
94
89
100
92
88
100
90
51
100
79
100
100
56
100
100
76
54
100
99
100
69
100
76
0.07

#### OEX

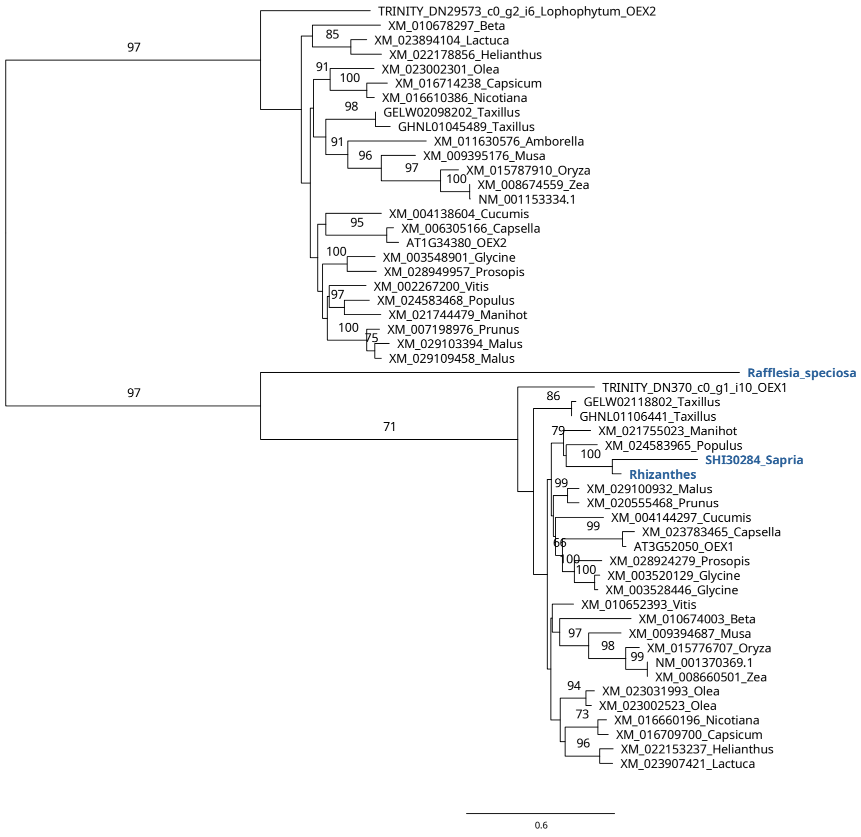

TRINITY_DN29573_c0_g2_i6_Lophophytum_OEX2
XM_010678297_Beta
XM_023894104_Lactuca
XM_022178856_Helianthus
XM_023002301_Olea
XM_016714238_Capsicum
XM_016610386_Nicotiana
GELW02098202_Taxillus
GHNL01045489_Taxillus
XM_011630576_Amborella
XM_009395176_Musa
XM_015787910_Oryza
XM_008674559_Zea
NM_001153334.1
XM_004138604_Cucumis
XM_006305166_Capsella
AT1G34380_OEX2
XM_003548901_Glycine
XM_028949957_Prosopis
XM_002267200_Vitis
XM_024583468_Populus
XM_021744479_Manihot
XM_007198976_Prunus
XM_029103394_Malus
XM_029109458_Malus
Rafflesia_speciosa
TRINITY_DN370_c0_g1_i10_OEX1
GELW02118802_Taxillus
GHNL01106441_Taxillus
XM_021755023_Manihot
XM_024583965_Populus
SHI30284_Sapria
Rhizanthes
XM_029100932_Malus
XM_020555468_Prunus
XM_004144297_Cucumis
XM_023783465_Capsella
AT3G52050_OEX1
XM_028924279_Prosopis
XM_003520129_Glycine
XM_003528446_Glycine
XM_010652393_Vitis
XM_010674003_Beta
XM_009394687_Musa
XM_015776707_Oryza
NM_001370369.1
XM_008660501_Zea
XM_023031993_Olea
XM_023002523_Olea
XM_016660196_Nicotiana
XM_016709700_Capsicum
XM_022153237_Helianthus
XM_023907421_Lactuca
0.6


#### OGG1

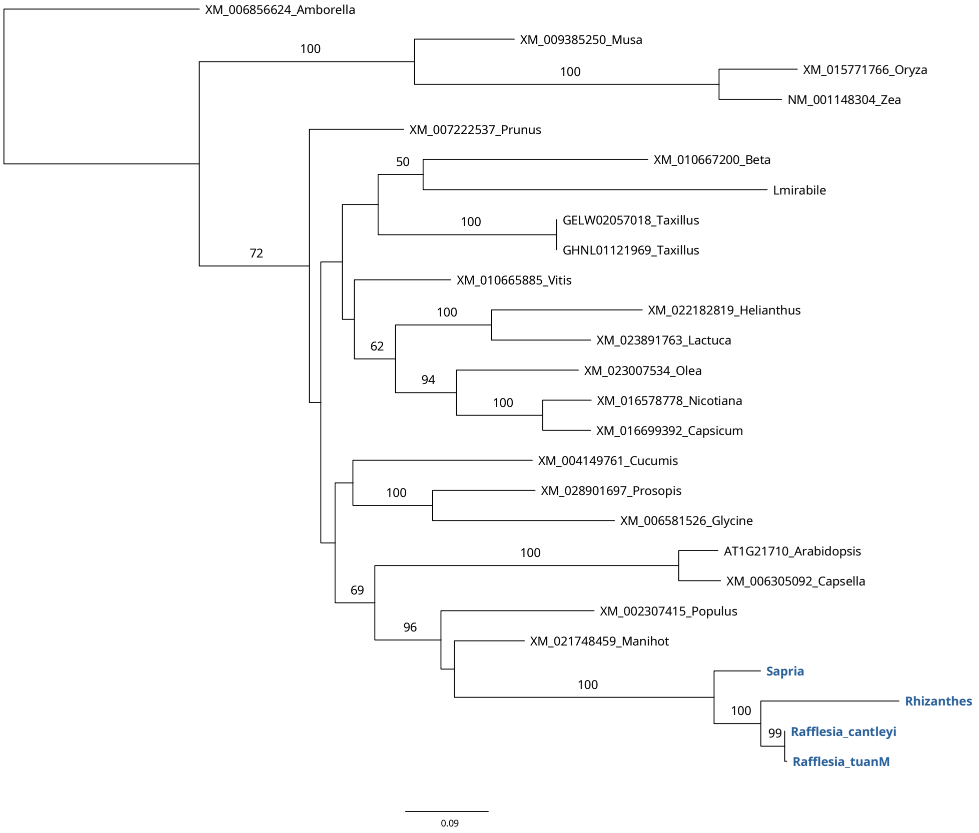

XM_006856624_Amborella
XM_009385250_Musa
XM_015771766_Oryza
NM_001148304_Zea
XM_007222537_Prunus
XM_010667200_Beta
Lmirabile
GELW02057018_Taxillus
GHNL01121969_Taxillus
XM_010665885_Vitis
XM_022182819_Helianthus
XM_023891763_Lactuca
XM_023007534_Olea
XM_016578778_Nicotiana
XM_016699392_Capsicum
XM_004149761_Cucumis
XM_028901697_Prosopis
XM_006581526_Glycine
AT1G21710_Arabidopsis
XM_006305092_Capsella
XM_002307415_Populus
XM_021748459_Manihot
Sapria
Rhizanthes
Rafflesia_cantleyi
Rafflesia_tuanM
0.09

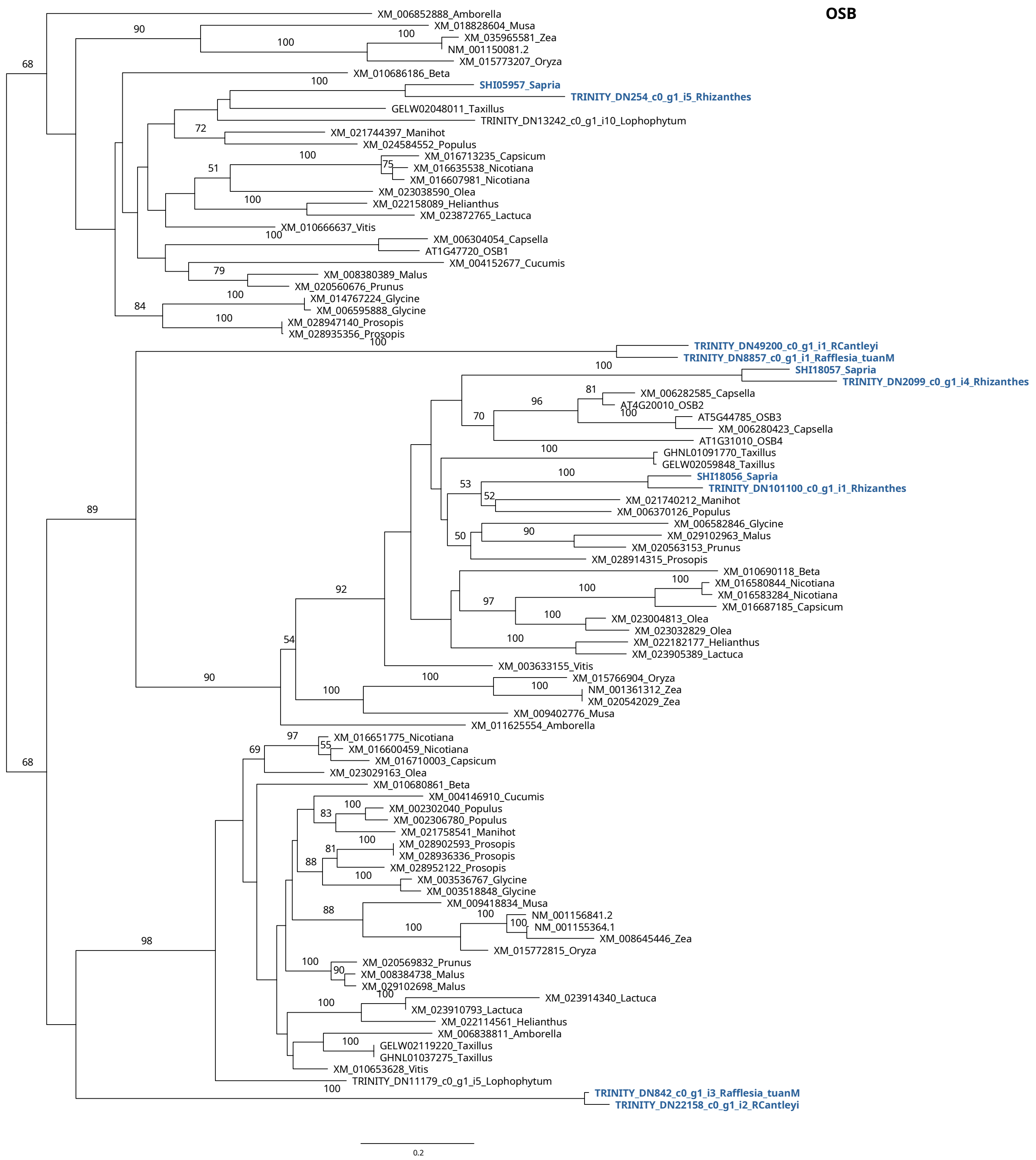
OSB
XM_006852888_Amborella
XM_018828604_Musa
XM_035965581_Zea
NM_001150081.2
XM_015773207_Oryza
XM_010686186_Beta
SHI05957_Sapria
TRINITY_DN254_c0_g1_i5_Rhizanthes
GELW02048011_Taxillus
TRINITY_DN13242_c0_g1_i10_Lophophytum
XM_021744397_Manihot
XM_024584552_Populus
XM_016713235_Capsicum
XM_016635538_Nicotiana
XM_016607981_Nicotiana
XM_023038590_Olea
XM_022158089_Helianthus
XM_023872765_Lactuca
XM_010666637_Vitis
XM_006304054_Capsella
AT1G47720_OSB1
XM_004152677_Cucumis
XM_008380389_Malus
XM_020560676_Prunus
XM_014767224_Glycine
XM_006595888_Glycine
XM_028947140_Prosopis
XM_028935356_Prosopis
TRINITY_DN49200_c0_g1_i1_RCantleyi
TRINITY_DN8857_c0_g1_i1_Rafflesia_tuanM
SHI18057_Sapria
TRINITY_DN2099_c0_g1_i4_Rhizanthes
XM_006282585_Capsella
AT4G20010_OSB2
AT5G44785_OSB3
XM_006280423_Capsella
AT1G31010_OSB4
GHNL01091770_Taxillus
GELW02059848_Taxillus
SHI18056_Sapria
TRINITY_DN101100_c0_g1_i1_Rhizanthes
XM_021740212_Manihot
XM_006370126_Populus
XM_006582846_Glycine
XM_029102963_Malus
XM_020563153_Prunus
XM_028914315_Prosopis
XM_010690118_Beta
XM_016580844_Nicotiana
XM_016583284_Nicotiana
XM_016687185_Capsicum
XM_023004813_Olea
XM_023032829_Olea
XM_022182177_Helianthus
XM_023905389_Lactuca
XM_003633155_Vitis
XM_015766904_Oryza
NM_001361312_Zea
XM_020542029_Zea
XM_009402776_Musa
XM_011625554_Amborella
XM_016651775_Nicotiana
XM_016600459_Nicotiana
XM_016710003_Capsicum
XM_023029163_Olea
XM_010680861_Beta
XM_004146910_Cucumis
XM_002302040_Populus
XM_002306780_Populus
XM_021758541_Manihot
XM_028902593_Prosopis
XM_028936336_Prosopis
XM_028952122_Prosopis
XM_003536767_Glycine
XM_003518848_Glycine
XM_009418834_Musa
NM_001156841.2
NM_001155364.1
XM_008645446_Zea
XM_015772815_Oryza
XM_020569832_Prunus
XM_008384738_Malus
XM_029102698_Malus
XM_023914340_Lactuca
XM_023910793_Lactuca
XM_022114561_Helianthus
XM_006838811_Amborella
GELW02119220_Taxillus
GHNL01037275_Taxillus
XM_010653628_Vitis
TRINITY_DN11179_c0_g1_i5_Lophophytum
TRINITY_DN842_c0_g1_i3_Rafflesia_tuanM
TRINITY_DN22158_c0_g1_i2_RCantleyi
0.2

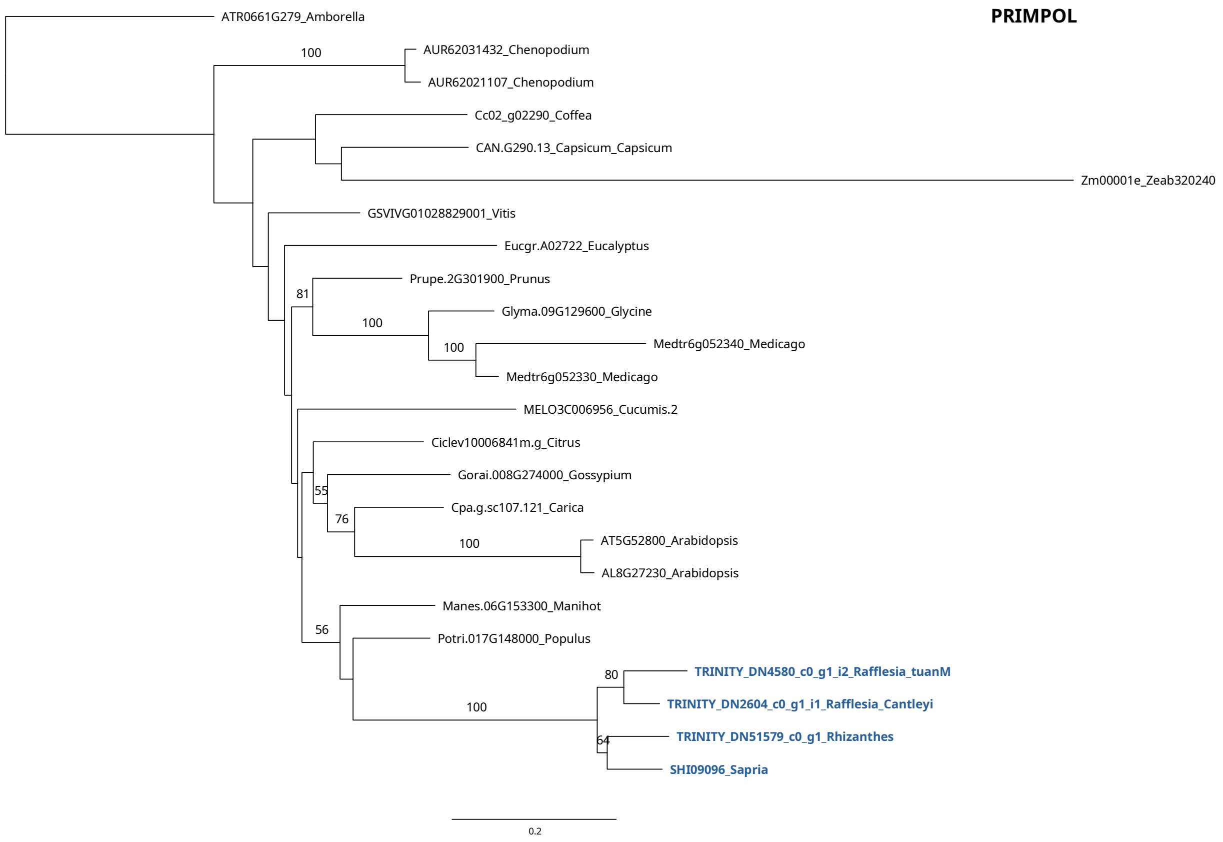
PRIMPOL
ATR0661G279_Amborella
AUR62031432_Chenopodium
AUR62021107_Chenopodium
Cc02_g02290_Coffea
CAN.G290.13_Capsicum_Capsicum
Zm00001e_Zeab320240
GSVIVG01028829001_Vitis
Eucgr.A02722_Eucalyptus
Prupe.2G301900_Prunus
Glyma.09G129600_Glycine
Medtr6g052340_Medicago
Medtr6g052330_Medicago
MELO3C006956_Cucumis.2
Ciclev10006841m.g_Citrus
Gorai.008G274000_Gossypium
Cpa.g.sc107.121_Carica
AT5G52800_Arabidopsis
AL8G27230_Arabidopsis
Manes.06G153300_Manihot
Potri.017G148000_Populus
TRINITY_DN4580_c0_g1_i2_Rafflesia_tuanM
TRINITY_DN2604_c0_g1_i1_Rafflesia_Cantleyi
TRINITY_DN51579_c0_g1_Rhizanthes
SHI09096_Sapria
0.2

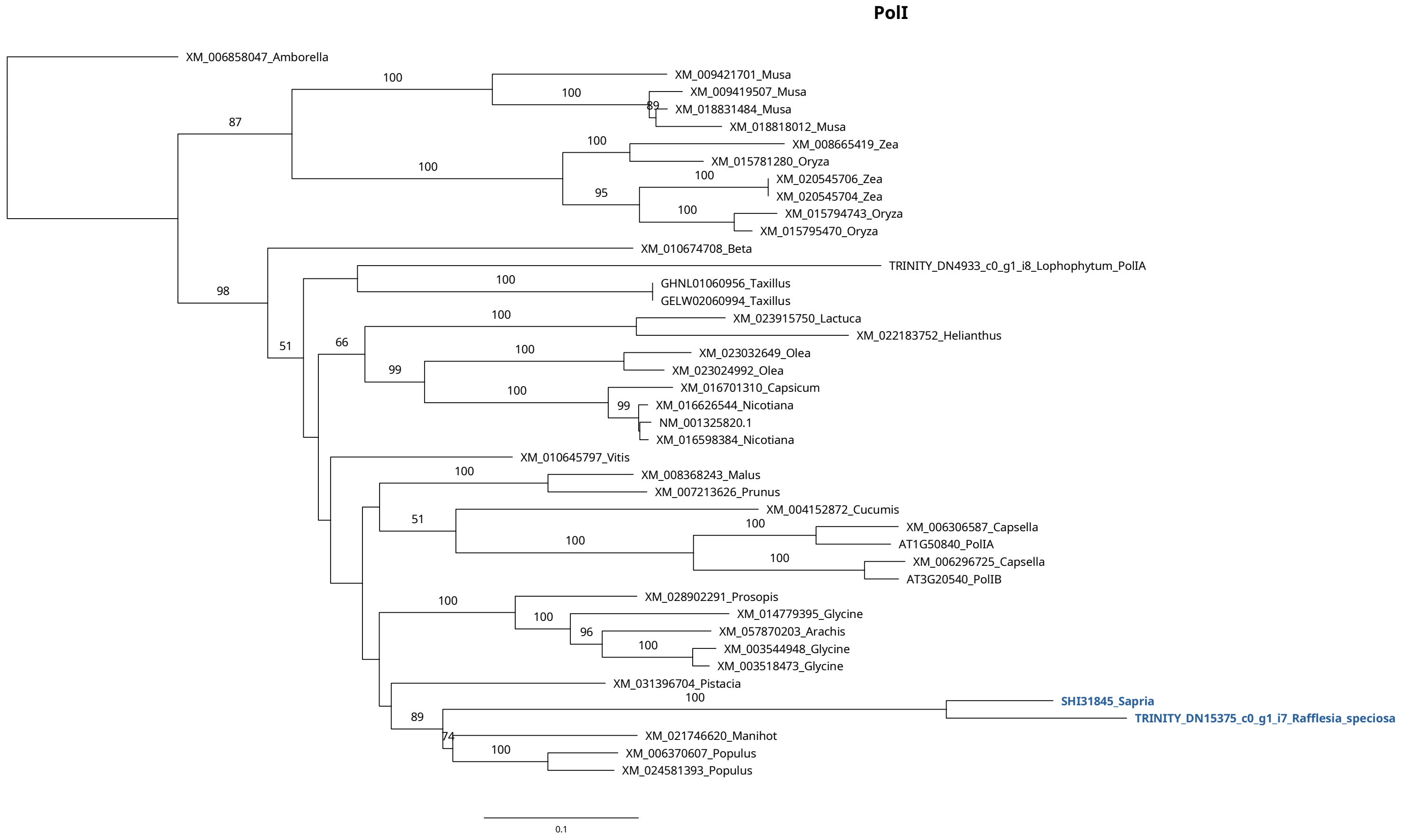
PolI
XM_006858047_Amborella
XM_009421701_Musa
XM_009419507_Musa
XM_018831484_Musa
XM_018818012_Musa
XM_008665419_Zea
XM_015781280_Oryza
XM_020545706_Zea
XM_020545704_Zea
XM_015794743_Oryza
XM_015795470_Oryza
XM_010674708_Beta
TRINITY_DN4933_c0_g1_i8_Lophophytum_PolIA
GHNL01060956_Taxillus
GELW02060994_Taxillus
XM_023915750_Lactuca
XM_022183752_Helianthus
XM_023032649_Olea
XM_023024992_Olea
XM_016701310_Capsicum
XM_016626544_Nicotiana
NM_001325820.1
XM_016598384_Nicotiana
XM_010645797_Vitis
XM_008368243_Malus
XM_007213626_Prunus
XM_004152872_Cucumis
XM_006306587_Capsella
AT1G50840_PolIA
XM_006296725_Capsella
AT3G20540_PolIB
XM_028902291_Prosopis
XM_014779395_Glycine
XM_057870203_Arachis
XM_003544948_Glycine
XM_003518473_Glycine
XM_031396704_Pistacia
SHI31845_Sapria
TRINITY_DN15375_c0_g1_i7_Rafflesia_speciosa
XM_021746620_Manihot
XM_006370607_Populus
XM_024581393_Populus
0.1

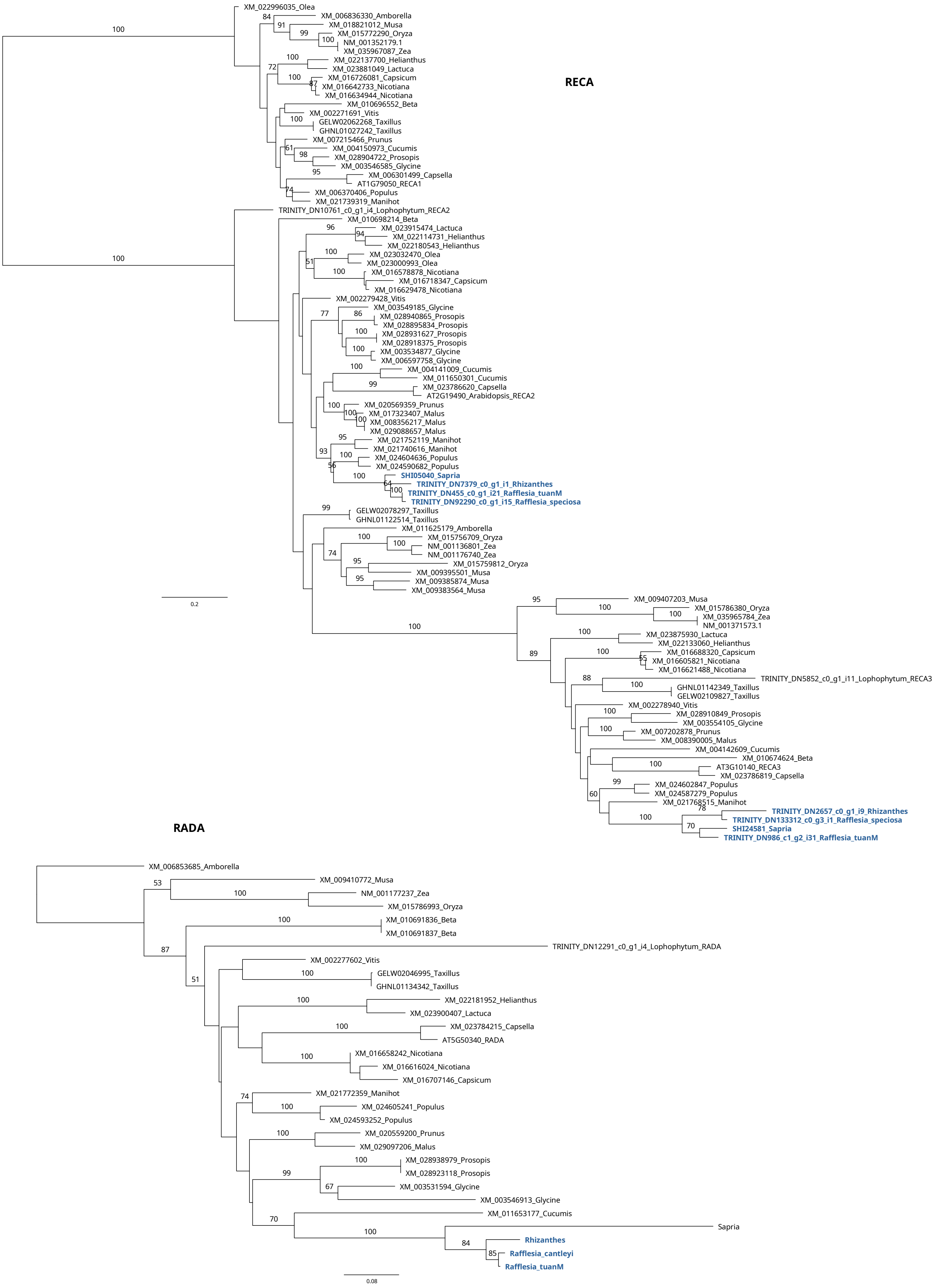
RECA
XM_022996035_Olea
XM_006836330_Amborella
XM_018821012_Musa
XM_015772290_Oryza
NM_001352179.1
XM_035967087_Zea
XM_022137700_Helianthus
XM_023881049_Lactuca
XM_016726081_Capsicum
XM_016642733_Nicotiana
XM_016634944_Nicotiana
XM_010696552_Beta
XM_002271691_Vitis
GELW02062268_Taxillus
GHNL01027242_Taxillus
XM_007215466_Prunus
XM_004150973_Cucumis
XM_028904722_Prosopis
XM_003546585_Glycine
XM_006301499_Capsella
AT1G79050_RECA1
XM_006370406_Populus
XM_021739319_Manihot
TRINITY_DN10761_c0_g1_i4_Lophophytum_RECA2
XM_010698214_Beta
XM_023915474_Lactuca
XM_022114731_Helianthus
XM_022180543_Helianthus
XM_023032470_Olea
XM_023000993_Olea
XM_016578878_Nicotiana
XM_016718347_Capsicum
XM_016629478_Nicotiana
XM_002279428_Vitis
XM_003549185_Glycine
XM_028940865_Prosopis
XM_028895834_Prosopis
XM_028931627_Prosopis
XM_028918375_Prosopis
XM_003534877_Glycine
XM_006597758_Glycine
XM_004141009_Cucumis
XM_011650301_Cucumis
XM_023786620_Capsella
AT2G19490_Arabidopsis_RECA2
XM_020569359_Prunus
XM_017323407_Malus
XM_008356217_Malus
XM_029088657_Malus
XM_021752119_Manihot
XM_021740616_Manihot
XM_024604636_Populus
XM_024590682_Populus
SHI05040_Sapria
TRINITY_DN7379_c0_g1_i1_Rhizanthes
TRINITY_DN455_c0_g1_i21_Rafflesia_tuanM
TRINITY_DN92290_c0_g1_i15_Rafflesia_speciosa
GELW02078297_Taxillus
GHNL01122514_Taxillus
XM_011625179_Amborella
XM_015756709_Oryza
NM_001136801_Zea
NM_001176740_Zea
XM_015759812_Oryza
XM_009395501_Musa
XM_009385874_Musa
XM_009383564_Musa
0.2
XM_009407203_Musa
XM_015786380_Oryza
XM_035965784_Zea
NM_001371573.1
XM_023875930_Lactuca
XM_022133060_Helianthus
XM_016688320_Capsicum
XM_016605821_Nicotiana
XM_016621488_Nicotiana
TRINITY_DN5852_c0_g1_i11_Lophophytum_RECA3
GHNL01142349_Taxillus
GELW02109827_Taxillus
XM_002278940_Vitis
XM_028910849_Prosopis
XM_003554105_Glycine
XM_007202878_Prunus
XM_008390005_Malus
XM_004142609_Cucumis
XM_010674624_Beta
AT3G10140_RECA3
XM_023786819_Capsella
XM_024602847_Populus
XM_024587279_Populus
XM_021768515_Manihot
TRINITY_DN2657_c0_g1_i9_Rhizanthes
TRINITY_DN133312_c0_g3_i1_Rafflesia_speciosa
SHI24581_Sapria
TRINITY_DN986_c1_g2_i31_Rafflesia_tuanM
RADA
XM_006853685_Amborella
XM_009410772_Musa
NM_001177237_Zea
XM_015786993_Oryza
XM_010691836_Beta
XM_010691837_Beta
TRINITY_DN12291_c0_g1_i4_Lophophytum_RADA
XM_002277602_Vitis
GELW02046995_Taxillus
GHNL01134342_Taxillus
XM_022181952_Helianthus
XM_023900407_Lactuca
XM_023784215_Capsella
AT5G50340_RADA
XM_016658242_Nicotiana
XM_016616024_Nicotiana
XM_016707146_Capsicum
XM_021772359_Manihot
XM_024605241_Populus
XM_024593252_Populus
XM_020559200_Prunus
XM_029097206_Malus
XM_028938979_Prosopis
XM_028923118_Prosopis
XM_003531594_Glycine
XM_003546913_Glycine
XM_011653177_Cucumis
Sapria
Rhizanthes
Rafflesia_cantleyi
Rafflesia_tuanM
0.08

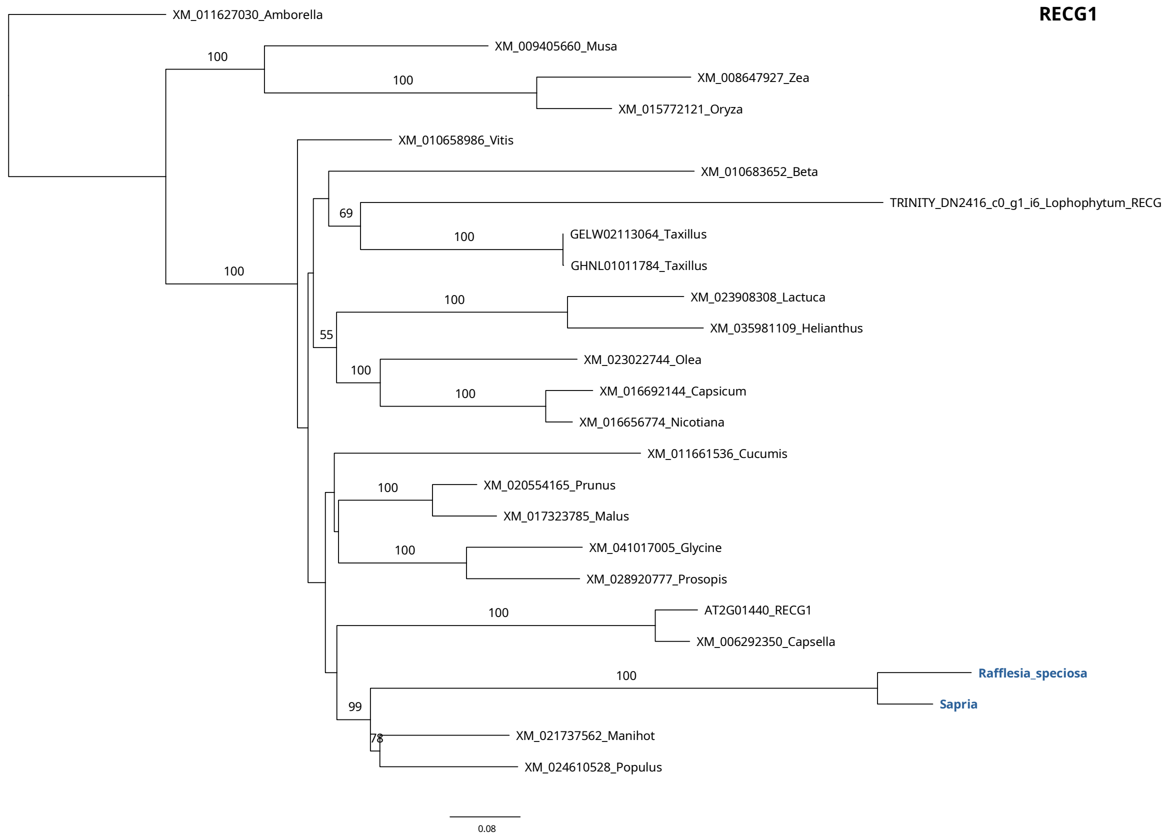
RECG1
XM_011627030_Amborella
XM_009405660_Musa
XM_008647927_Zea
XM_015772121_Oryza
XM_010658986_Vitis
XM_010683652_Beta
TRINITY_DN2416_c0_g1_i6_Lophophytum_RECG
GELW02113064_Taxillus
GHNL01011784_Taxillus
XM_023908308_Lactuca
XM_035981109_Helianthus
XM_023022744_Olea
XM_016692144_Capsicum
XM_016656774_Nicotiana
XM_011661536_Cucumis
XM_020554165_Prunus
XM_017323785_Malus
XM_041017005_Glycine
XM_028920777_Prosopis
AT2G01440_RECG1
XM_006292350_Capsella
Rafflesia_speciosa
Sapria
XM_021737562_Manihot
XM_024610528_Populus
100
100
100
69
100
100
55
100
100
100
100
100
100
99
78
0.08

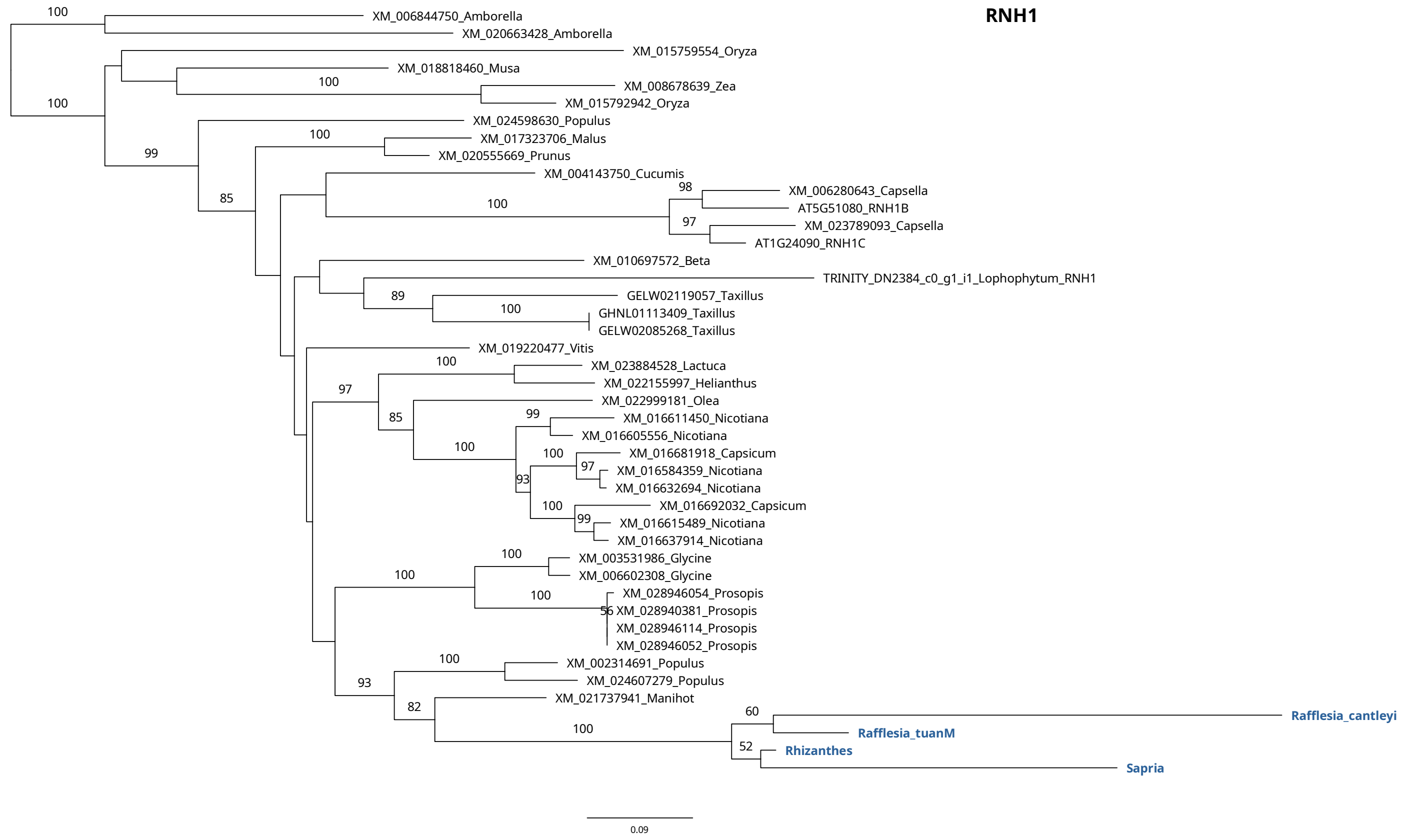
RNH1
XM_006844750_Amborella
XM_020663428_Amborella
XM_015759554_Oryza
XM_018818460_Musa
XM_008678639_Zea
XM_015792942_Oryza
XM_024598630_Populus
XM_017323706_Malus
XM_020555669_Prunus
XM_004143750_Cucumis
XM_006280643_Capsella
AT5G51080_RNH1B
XM_023789093_Capsella
AT1G24090_RNH1C
XM_010697572_Beta
TRINITY_DN2384_c0_g1_i1_Lophophytum_RNH1
GELW02119057_Taxillus
GHNL01113409_Taxillus
GELW02085268_Taxillus
XM_019220477_Vitis
XM_023884528_Lactuca
XM_022155997_Helianthus
XM_022999181_Olea
XM_016611450_Nicotiana
XM_016605556_Nicotiana
XM_016681918_Capsicum
XM_016584359_Nicotiana
XM_016632694_Nicotiana
XM_016692032_Capsicum
XM_016615489_Nicotiana
XM_016637914_Nicotiana
XM_003531986_Glycine
XM_006602308_Glycine
XM_028946054_Prosopis
XM_028940381_Prosopis
XM_028946114_Prosopis
XM_028946052_Prosopis
XM_002314691_Populus
XM_024607279_Populus
XM_021737941_Manihot
Rafflesia_cantleyi
Rafflesia_tuanM
Rhizanthes
Sapria
100
100
100
99
100
85
98
100
97
89
100
100
97
85
99
100
100
97
93
100
99
100
100
100
56
100
93
82
60
100
52
0.09

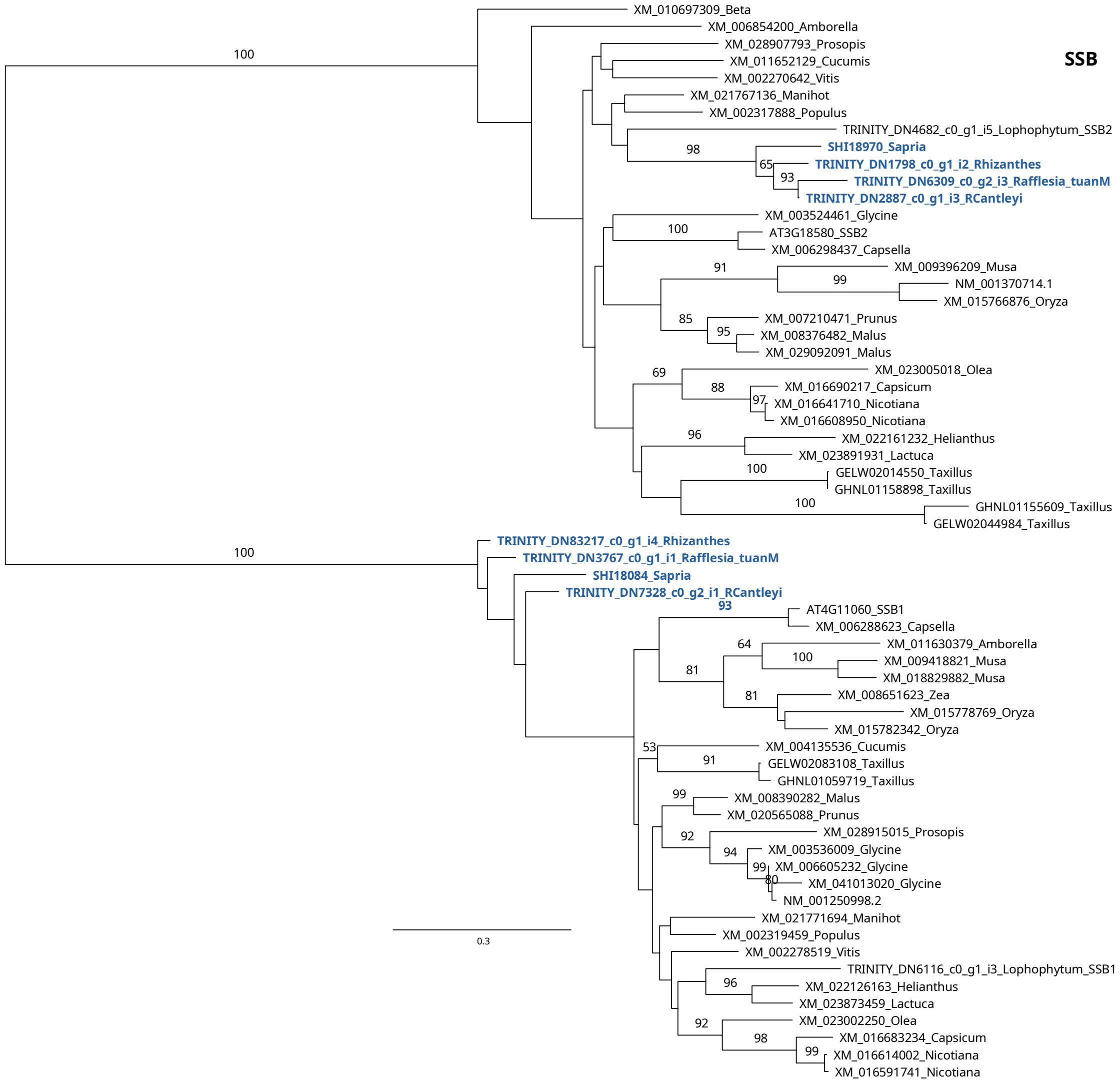
SSB
XM_010697309_Beta
XM_006854200_Amborella
XM_028907793_Prosopis
XM_011652129_Cucumis
XM_002270642_Vitis
XM_021767136_Manihot
XM_002317888_Populus
TRINITY_DN4682_c0_g1_i5_Lophophytum_SSB2
SHI18970_Sapria
TRINITY_DN1798_c0_g1_i2_Rhizanthes
TRINITY_DN6309_c0_g2_i3_Rafflesia_tuanM
TRINITY_DN2887_c0_g1_i3_RCantleyi
XM_003524461_Glycine
AT3G18580_SSB2
XM_006298437_Capsella
XM_009396209_Musa
NM_001370714.1
XM_015766876_Oryza
XM_007210471_Prunus
XM_008376482_Malus
XM_029092091_Malus
XM_023005018_Olea
XM_016690217_Capsicum
XM_016641710_Nicotiana
XM_016608950_Nicotiana
XM_022161232_Helianthus
XM_023891931_Lactuca
GELW02014550_Taxillus
GHNL01158898_Taxillus
GHNL01155609_Taxillus
GELW02044984_Taxillus
TRINITY_DN83217_c0_g1_i4_Rhizanthes
TRINITY_DN3767_c0_g1_i1_Rafflesia_tuanM
SHI18084_Sapria
TRINITY_DN7328_c0_g2_i1_RCantleyi
AT4G11060_SSB1
XM_006288623_Capsella
XM_011630379_Amborella
XM_009418821_Musa
XM_018829882_Musa
XM_008651623_Zea
XM_015778769_Oryza
XM_015782342_Oryza
XM_004135536_Cucumis
GELW02083108_Taxillus
GHNL01059719_Taxillus
XM_008390282_Malus
XM_020565088_Prunus
XM_028915015_Prosopis
XM_003536009_Glycine
XM_006605232_Glycine
XM_041013020_Glycine
NM_001250998.2
XM_021771694_Manihot
XM_002319459_Populus
XM_002278519_Vitis
TRINITY_DN6116_c0_g1_i3_Lophophytum_SSB1
XM_022126163_Helianthus
XM_023873459_Lactuca
XM_023002250_Olea
XM_016683234_Capsicum
XM_016614002_Nicotiana
XM_016591741_Nicotiana
0.3

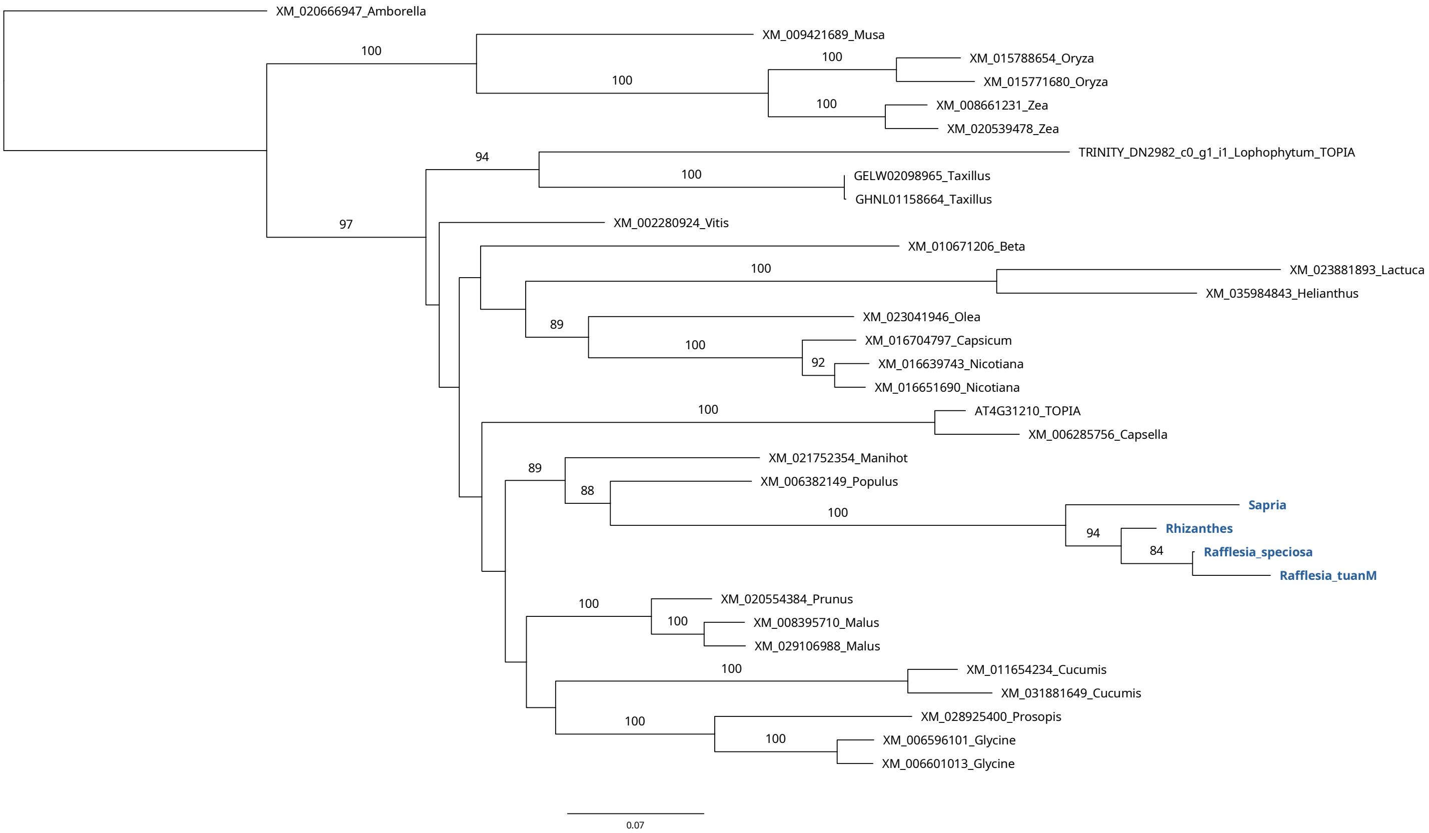
TOPIA
XM_020666947_Amborella
XM_009421689_Musa
XM_015788654_Oryza
XM_015771680_Oryza
XM_008661231_Zea
XM_020539478_Zea
TRINITY_DN2982_c0_g1_i1_Lophophytum_TOPIA
GELW02098965_Taxillus
GHNL01158664_Taxillus
XM_002280924_Vitis
XM_010671206_Beta
XM_023881893_Lactuca
XM_035984843_Helianthus
XM_023041946_Olea
XM_016704797_Capsicum
XM_016639743_Nicotiana
XM_016651690_Nicotiana
AT4G31210_TOPIA
XM_006285756_Capsella
XM_021752354_Manihot
XM_006382149_Populus
Sapria
Rhizanthes
Rafflesia_speciosa
Rafflesia_tuanM
XM_020554384_Prunus
XM_008395710_Malus
XM_029106988_Malus
XM_011654234_Cucumis
XM_031881649_Cucumis
XM_028925400_Prosopis
XM_006596101_Glycine
XM_006601013_Glycine
0.07

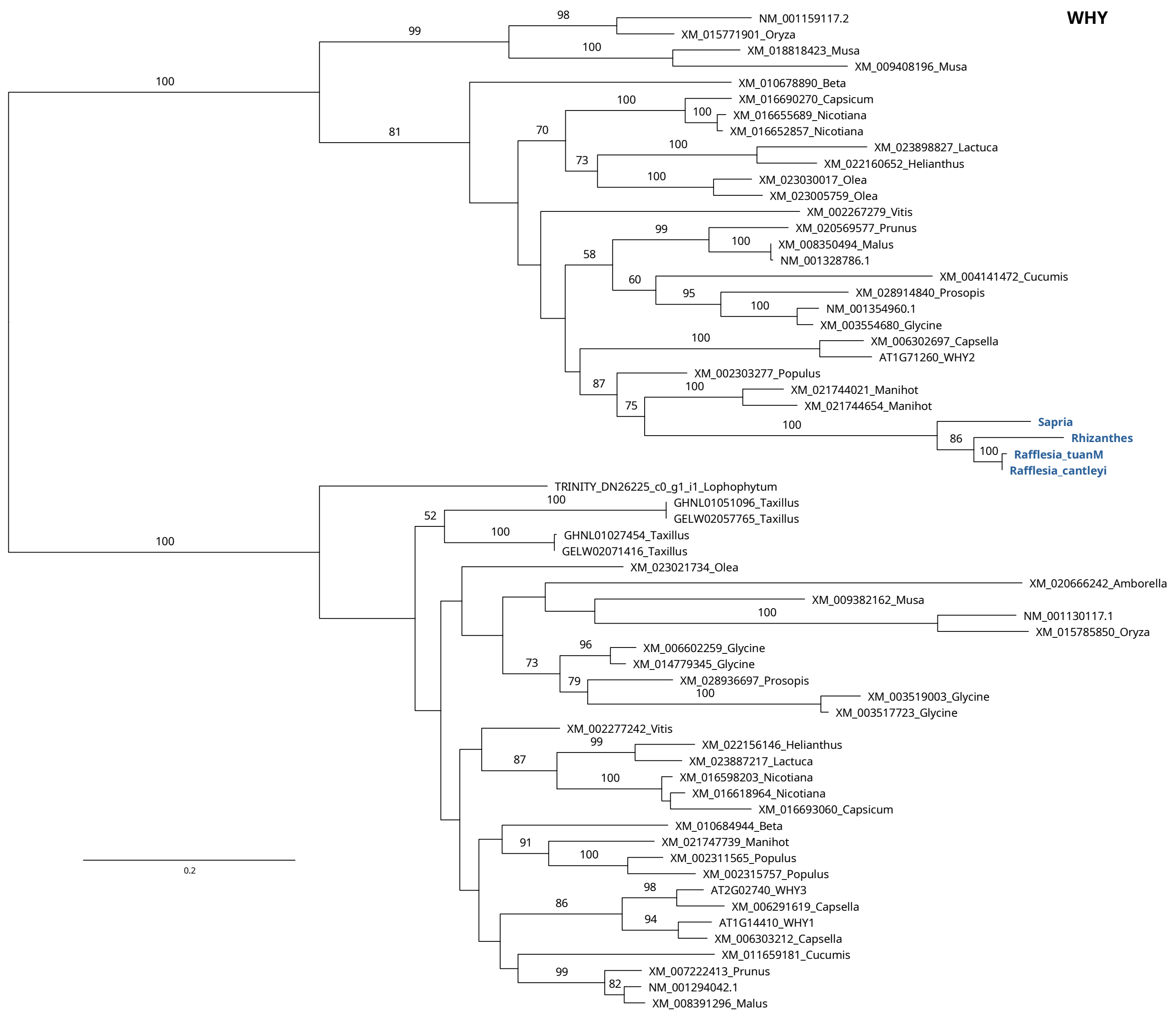
WHY
NM_001159117.2
XM_015771901_Oryza
XM_018818423_Musa
XM_009408196_Musa
XM_010678890_Beta
XM_016690270_Capsicum
XM_016655689_Nicotiana
XM_016652857_Nicotiana
XM_023898827_Lactuca
XM_022160652_Helianthus
XM_023030017_Olea
XM_023005759_Olea
XM_002267279_Vitis
XM_020569577_Prunus
XM_008350494_Malus
NM_001328786.1
XM_004141472_Cucumis
XM_028914840_Prosopis
NM_001354960.1
XM_003554680_Glycine
XM_006302697_Capsella
AT1G71260_WHY2
XM_002303277_Populus
XM_021744021_Manihot
XM_021744654_Manihot
Sapria
Rhizanthes
Rafflesia_tuanM
Rafflesia_cantleyi
TRINITY_DN26225_c0_g1_i1_Lophophytum
GHNL01051096_Taxillus
GELW02057765_Taxillus
GHNL01027454_Taxillus
GELW02071416_Taxillus
XM_023021734_Olea
XM_020666242_Amborella
XM_009382162_Musa
NM_001130117.1
XM_015785850_Oryza
XM_006602259_Glycine
XM_014779345_Glycine
XM_028936697_Prosopis
XM_003519003_Glycine
XM_003517723_Glycine
XM_002277242_Vitis
XM_022156146_Helianthus
XM_023887217_Lactuca
XM_016598203_Nicotiana
XM_016618964_Nicotiana
XM_016693060_Capsicum
XM_010684944_Beta
XM_021747739_Manihot
XM_002311565_Populus
XM_002315757_Populus
AT2G02740_WHY3
XM_006291619_Capsella
AT1G14410_WHY1
XM_006303212_Capsella
XM_011659181_Cucumis
XM_007222413_Prunus
NM_001294042.1
XM_008391296_Malus
0.2

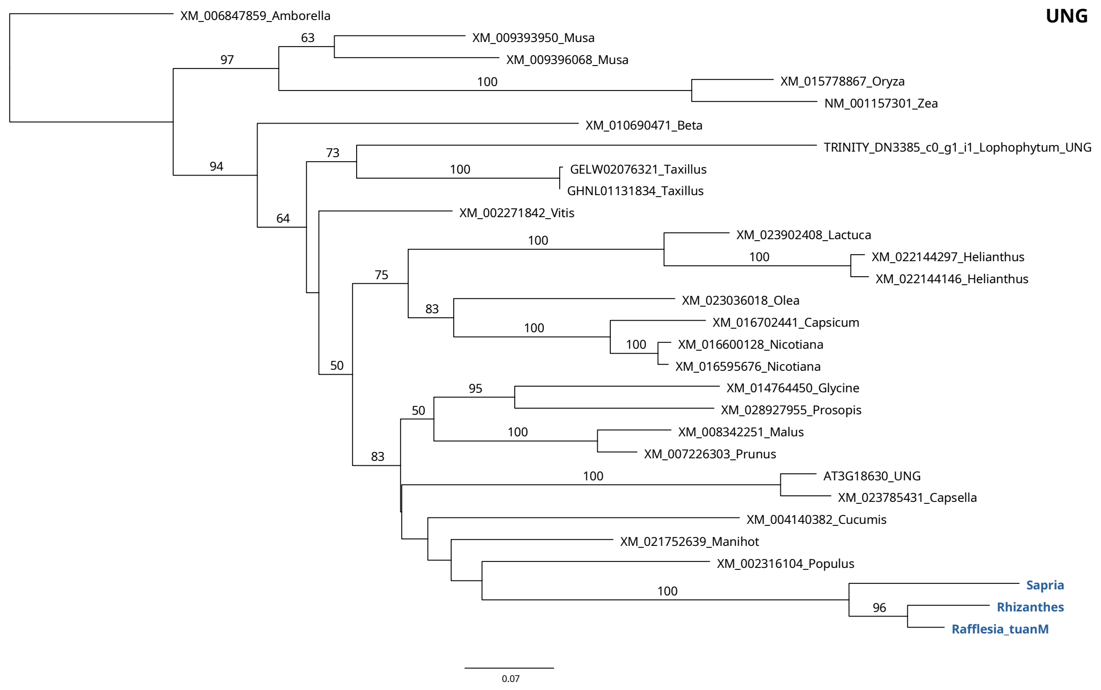
UNG
XM_006847859_Amborella
XM_009393950_Musa
XM_009396068_Musa
XM_015778867_Oryza
NM_001157301_Zea
XM_010690471_Beta
TRINITY_DN3385_c0_g1_i1_Lophophytum_UNG
GELW02076321_Taxillus
GHNL01131834_Taxillus
XM_002271842_Vitis
XM_023902408_Lactuca
XM_022144297_Helianthus
XM_022144146_Helianthus
XM_023036018_Olea
XM_016702441_Capsicum
XM_016600128_Nicotiana
XM_016595676_Nicotiana
XM_014764450_Glycine
XM_028927955_Prosopis
XM_008342251_Malus
XM_007226303_Prunus
AT3G18630_UNG
XM_023785431_Capsella
XM_004140382_Cucumis
XM_021752639_Manihot
XM_002316104_Populus
Sapria
Rhizanthes
Rafflesia_tuanM
0.07

**Figure S10. DNA read depth of the mitochondrial genomes calculated with Bowtie2 v.2.4.4 (parameters: --end-to-end --very-sensitive --no-contain --no-discordant --no-mixed).**

A) *Sapria himalayana* (Individual 1)

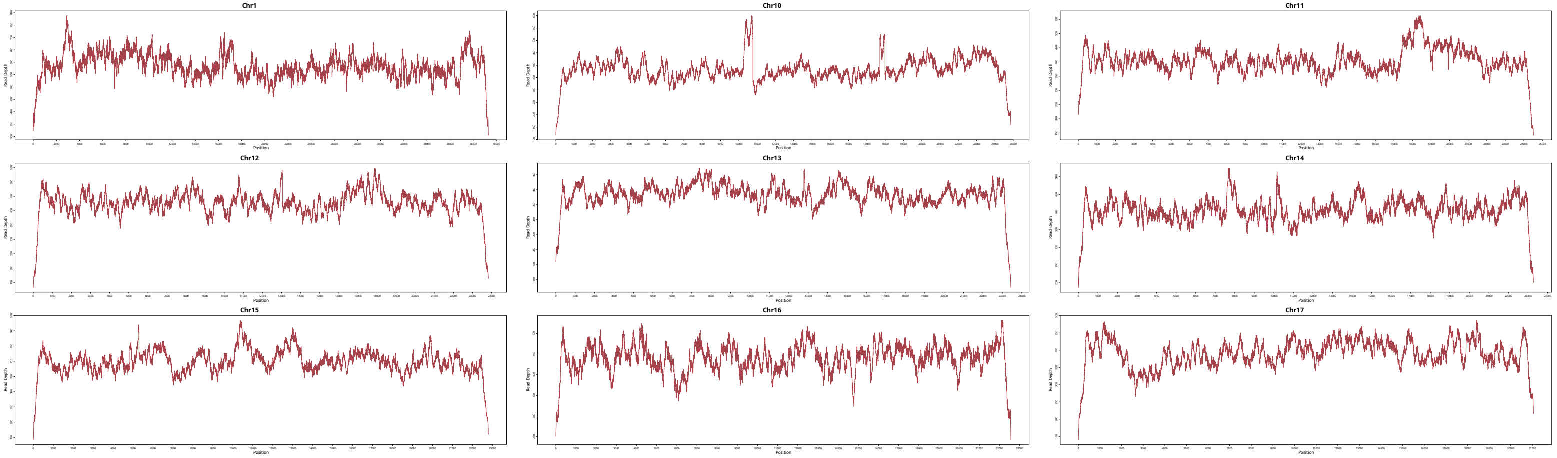

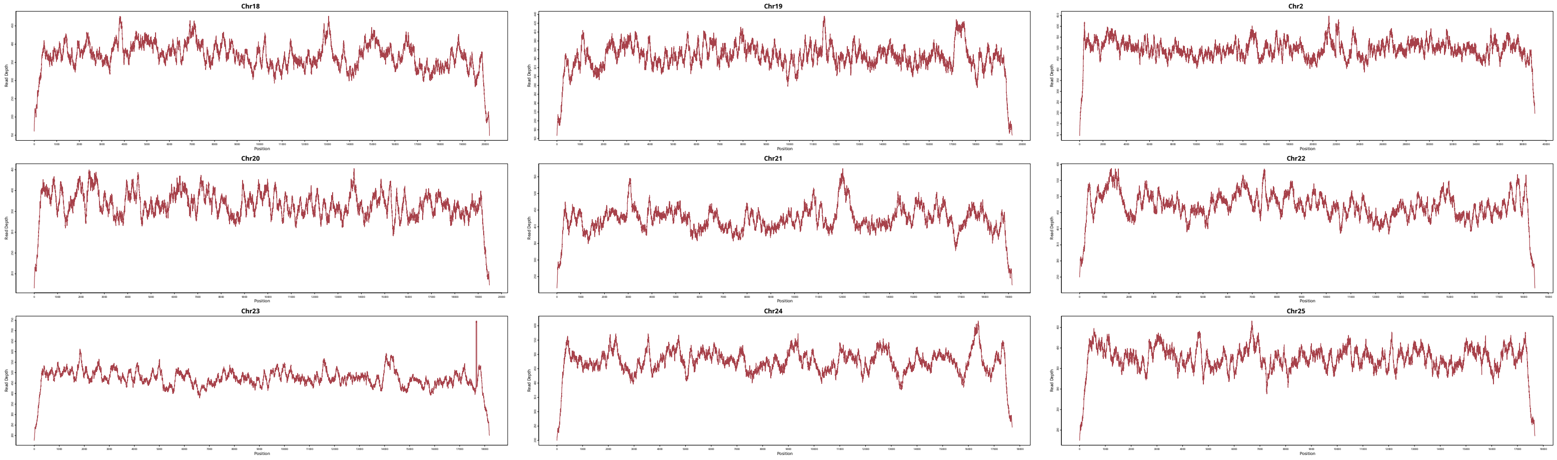

Chr18
Chr19
Chr2
Chr20
Chr21
Chr22
Chr23
Chr24
Chr25
Read Depth
Position

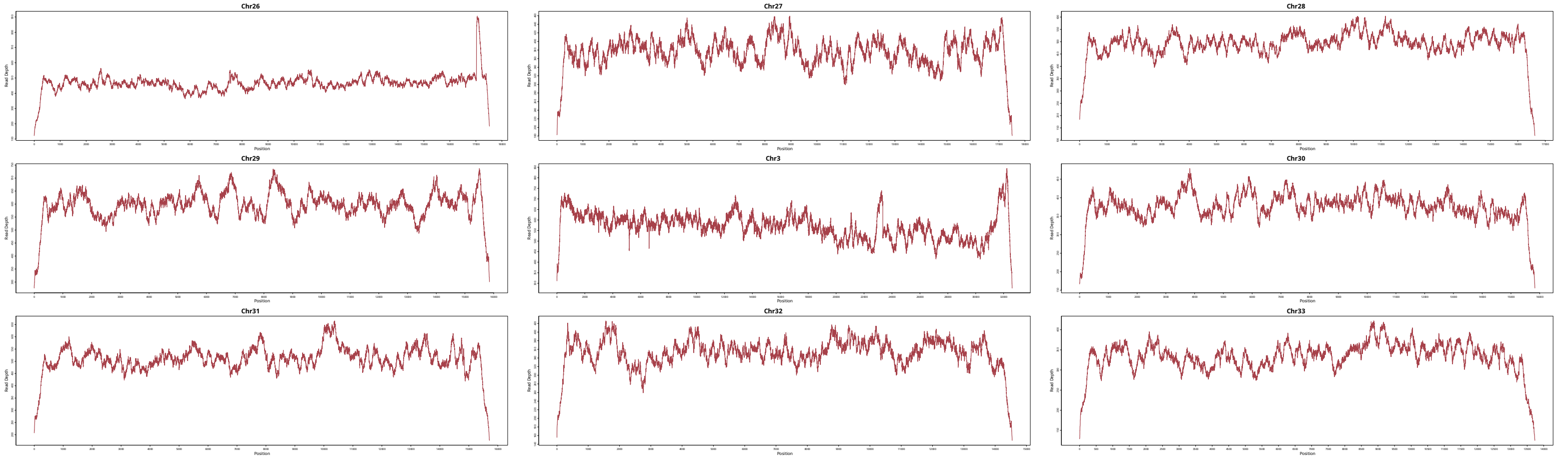
Chr26
Chr27
Chr28
Chr29
Chr3
Chr30
Chr31
Chr32
Chr33
Read Depth
Position

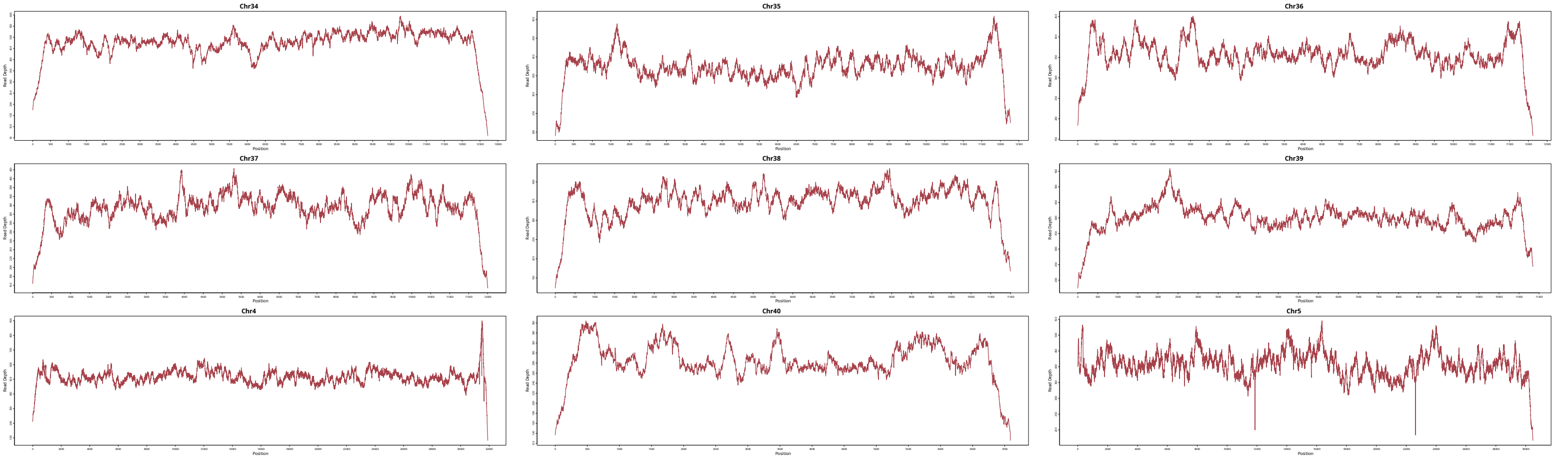
Chr34
Chr35
Chr36
Chr37
Chr38
Chr39
Chr4
Chr40
Chr5
Read Depth
Position

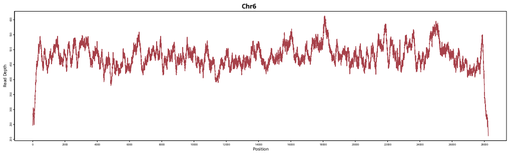
Chr6
Read Depth
Position

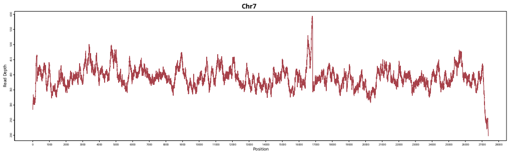
Chr7
Read Depth
Position

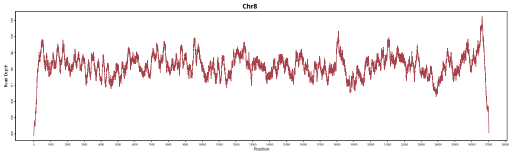
Chr8
Read Depth
Position

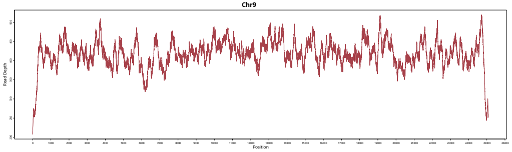
Chr9
Read Depth
Position

B) *Sapria himalayana* (Individual 2)

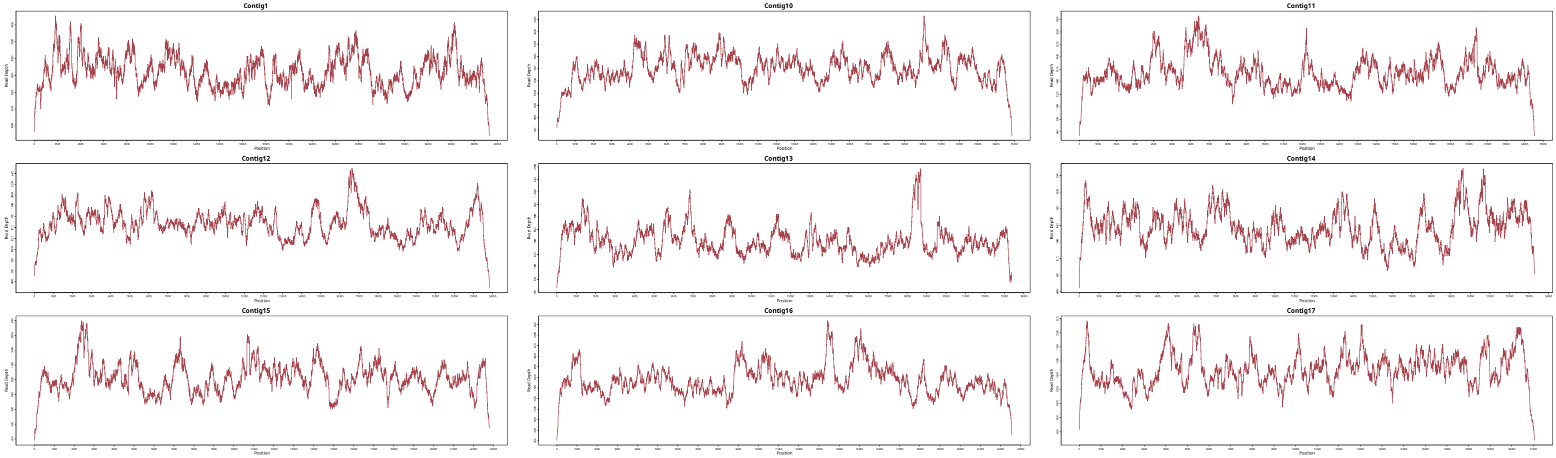

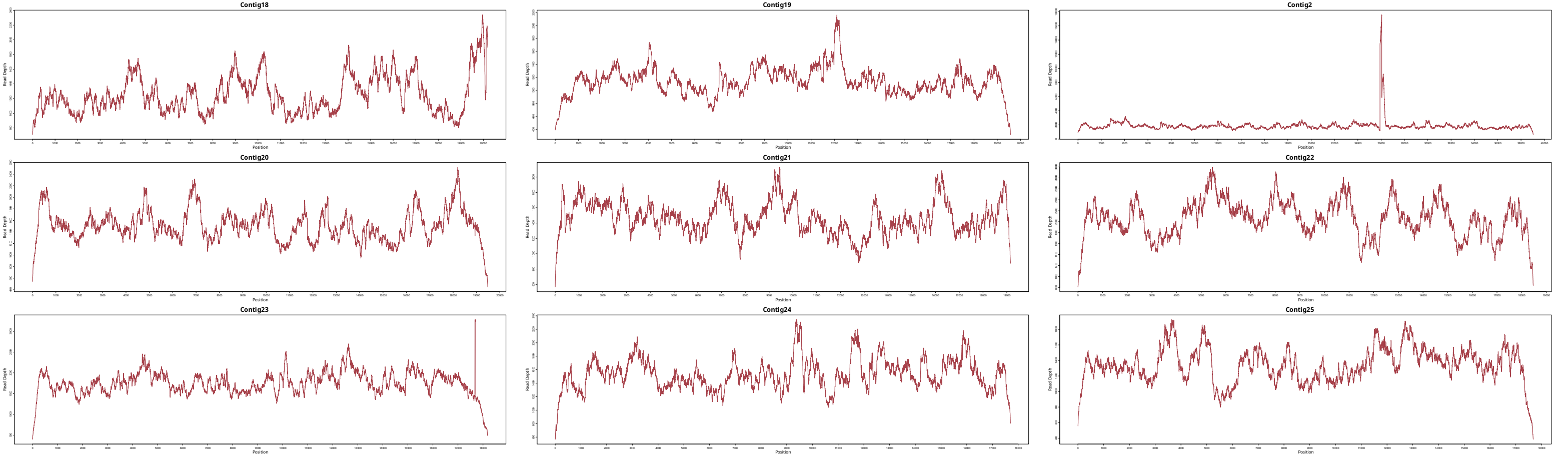
Contig18
Contig19
Contig2
Contig20
Contig21
Contig22
Contig23
Contig24
Contig25
Read Depth
Position

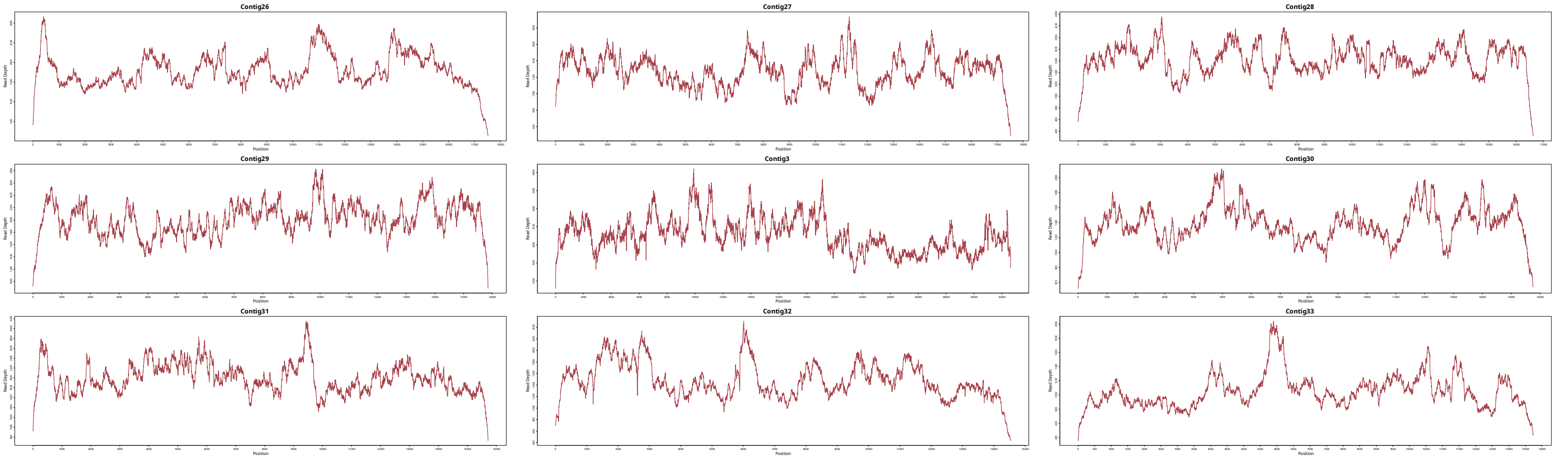
Contig26
Contig27
Contig28
Contig29
Contig3
Contig30
Contig31
Contig32
Contig33
Read Depth
Position

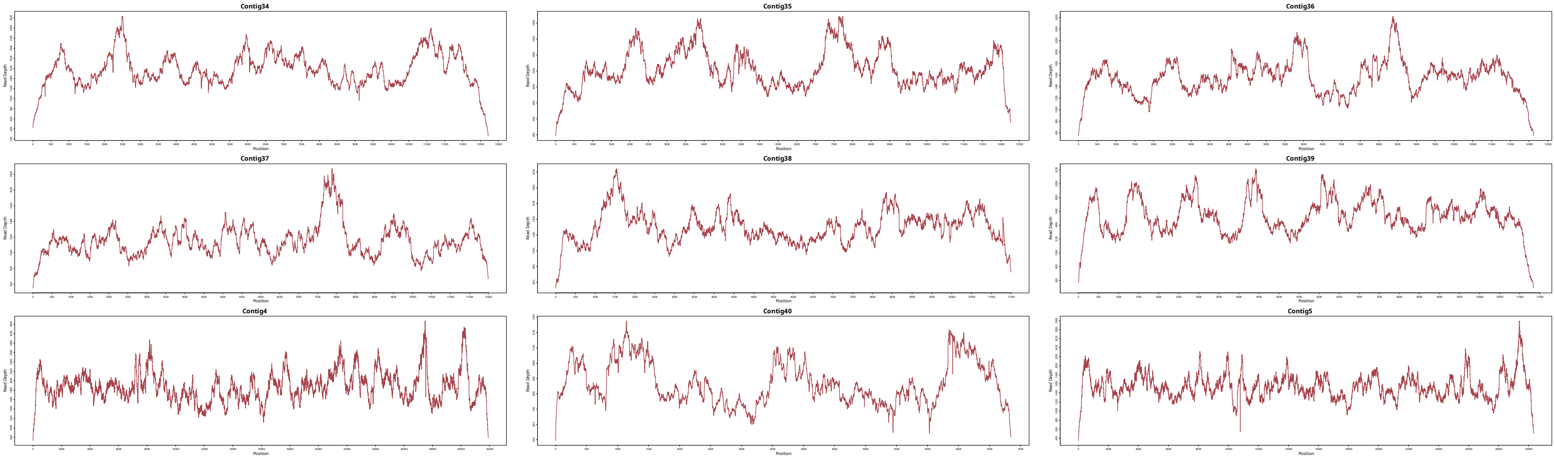
Contig34
Contig35
Contig36
Contig37
Contig38
Contig39
Contig4
Contig40
Contig5
Read Depth
Position

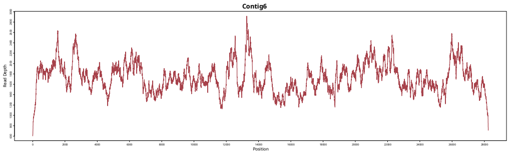
Contig6
Read Depth
Position

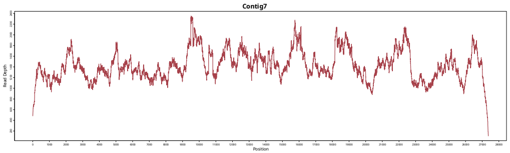
Contig7
Read Depth
Position

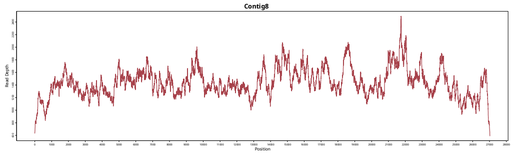
Contig8
Read Depth
Position

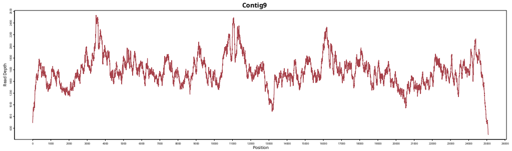
Contig9
Read Depth
Position

C) *Rhizanthes lowii*

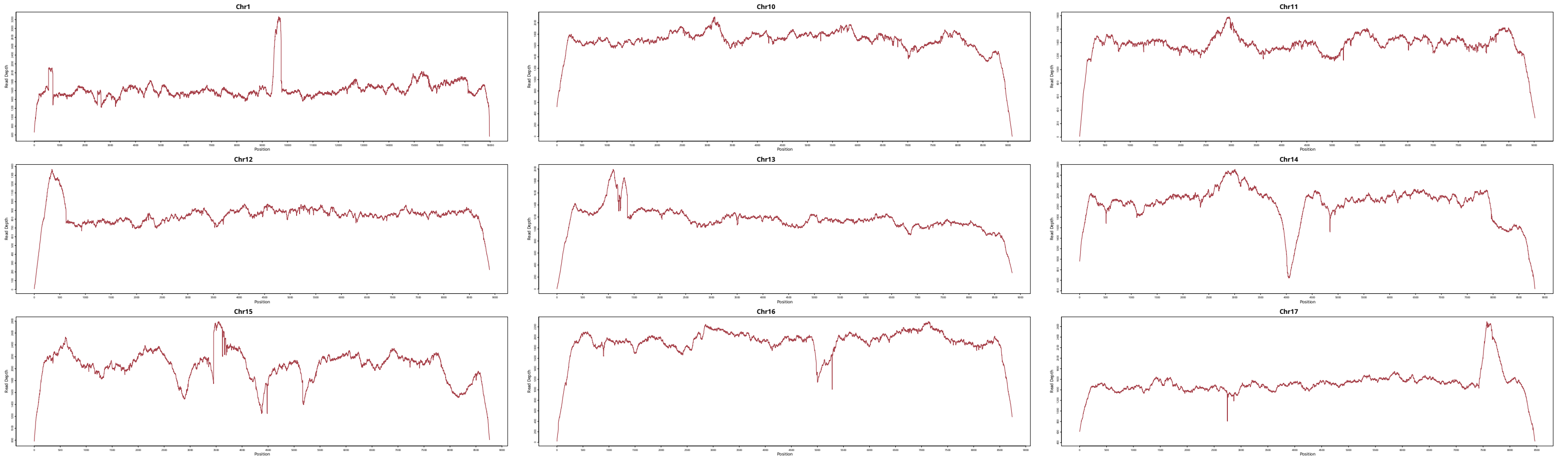

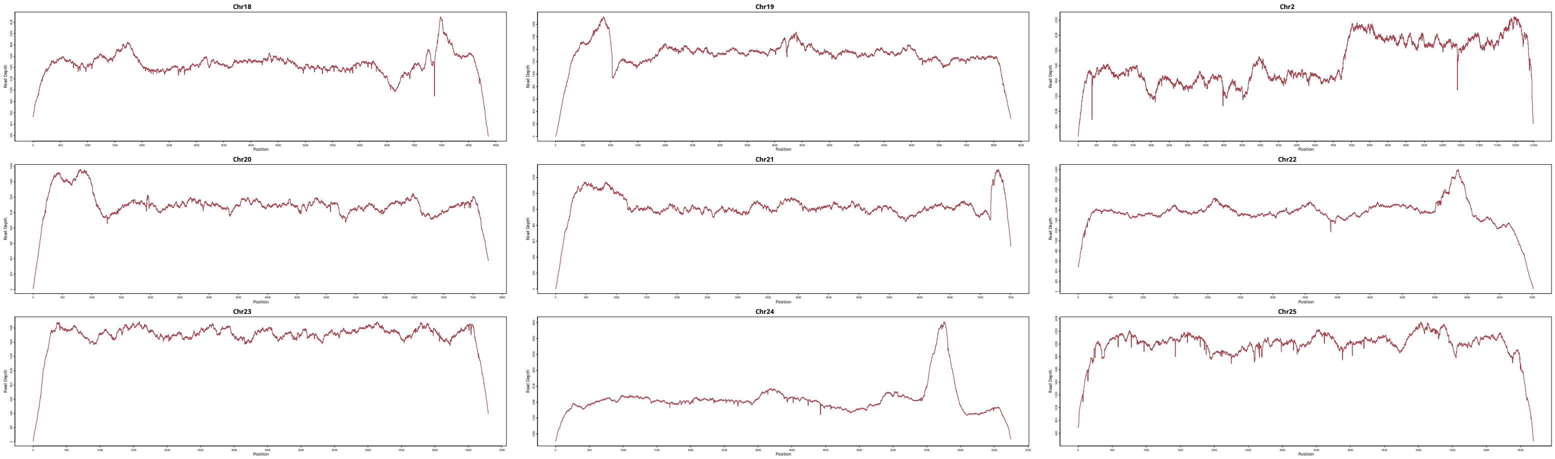
Chr18
Chr19
Chr2
Chr20
Chr21
Chr22
Chr23
Chr24
Chr25
Read Depth
Position

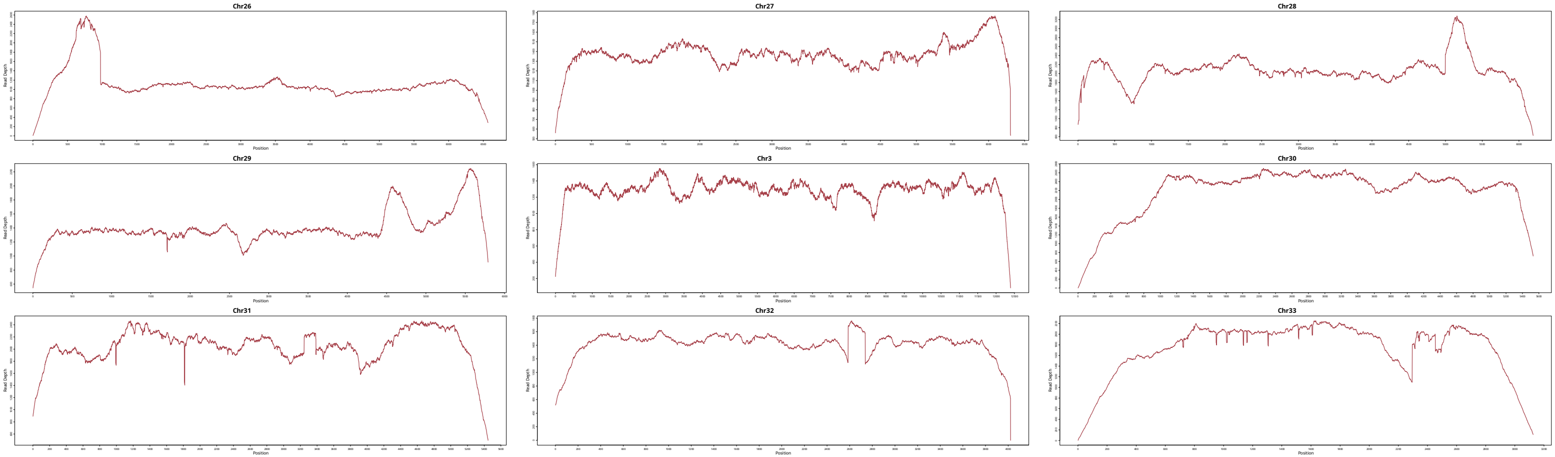

Chr26
Chr27
Chr28
Chr29
Chr3
Chr30
Chr31
Chr32
Chr33
Read Depth
Position

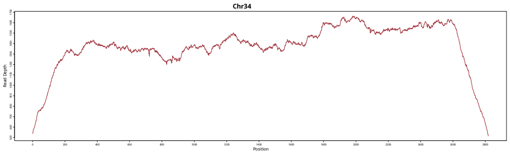
Chr34
Read Depth
Position

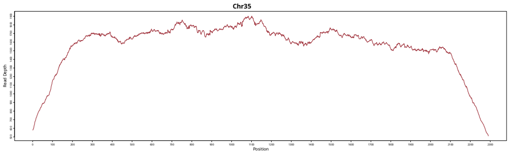
Chr35
Read Depth
Position

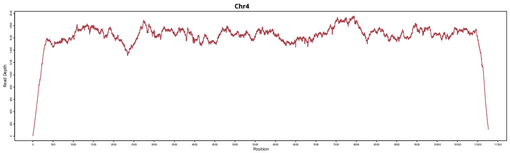
Chr4
Read Depth
Position

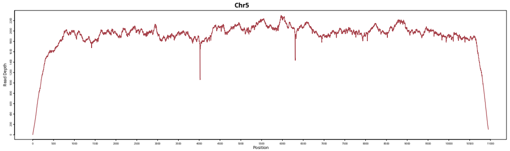
Chr5
Read Depth
Position

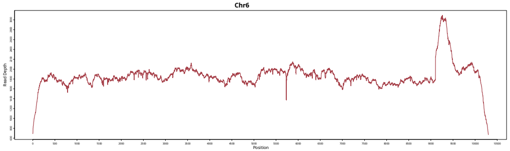
Chr6
Read Depth
Position

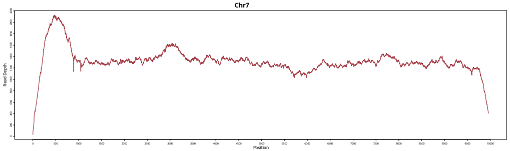
Chr7
Read Depth
Position

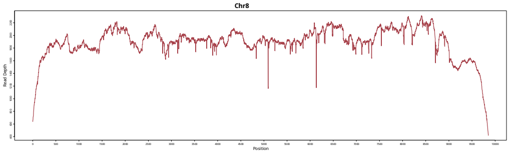
Chr8
Read Depth
Position

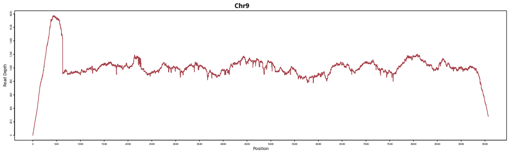
Chr9
Read Depth
Position

D) *Tetrastigma sp.* Individual 1

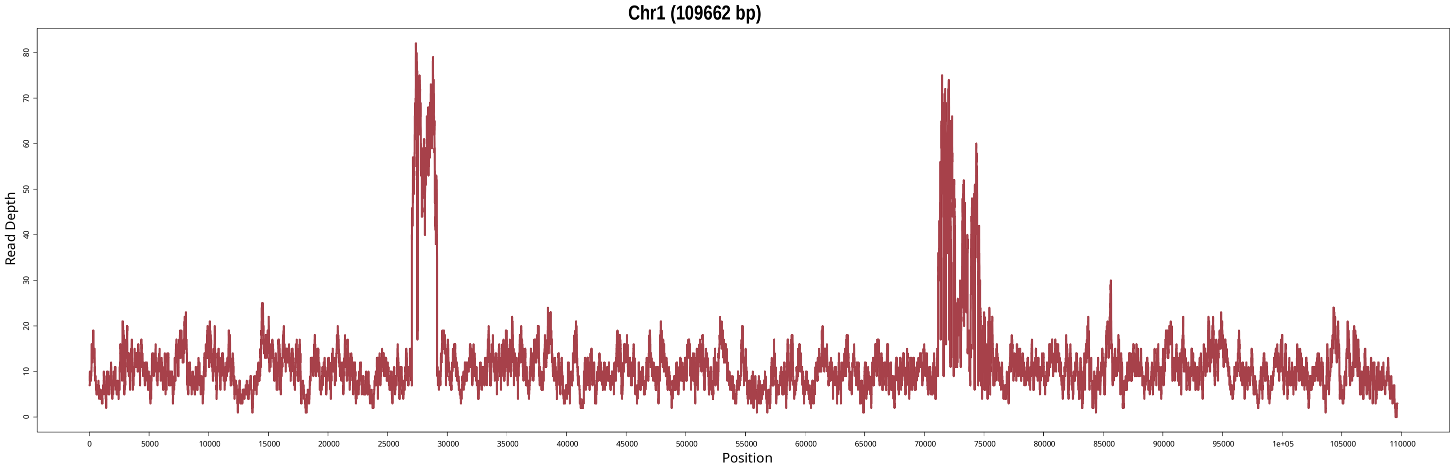

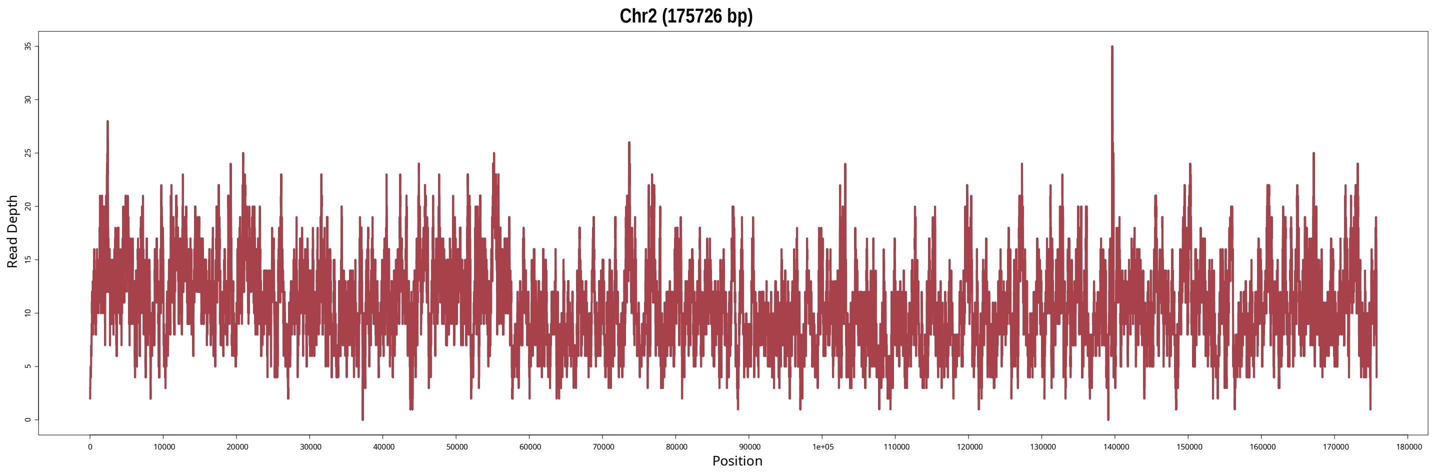
Chr2 (175726 bp)
Read Depth
0
5
10
15
20
25
30
35
0
10000
20000
30000
40000
50000
60000
70000
80000
90000
1e+05
110000
120000
130000
140000
150000
160000
170000
180000
Position

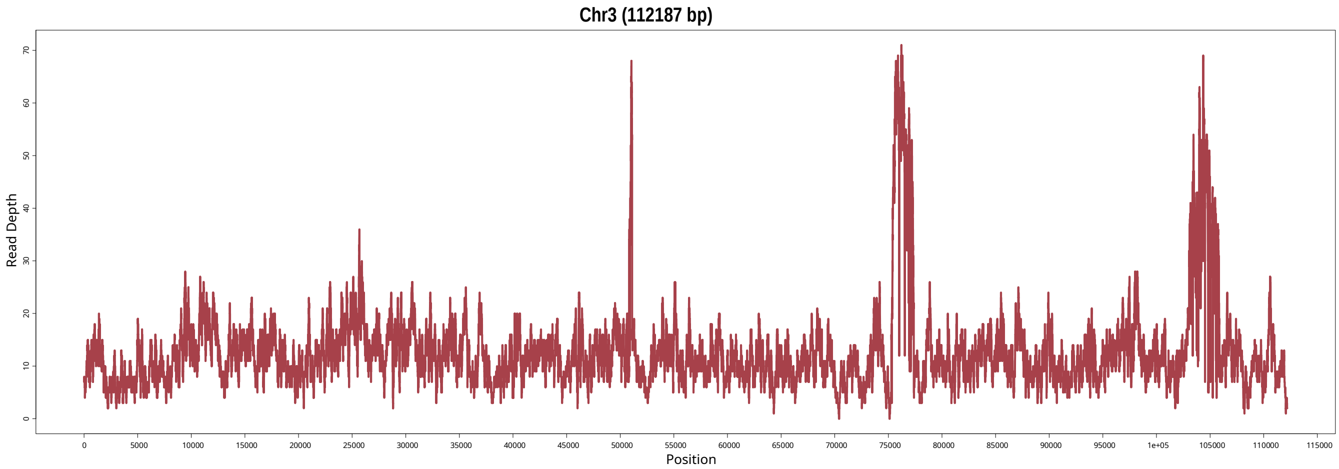
Chr3 (112187 bp)
Read Depth
0
10
20
30
40
50
60
70
0
5000
10000
15000
20000
25000
30000
35000
40000
45000
50000
55000
60000
65000
70000
75000
80000
85000
90000
95000
1e+05
105000
110000
115000
Position

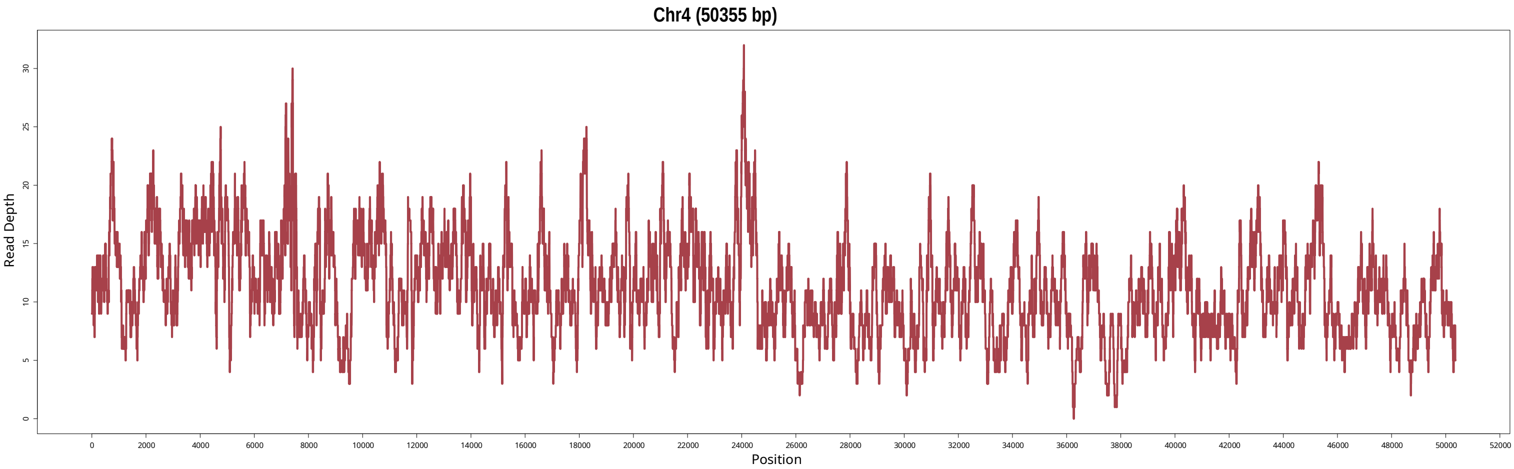
Chr4 (50355 bp)
Read Depth
Position
0
5
10
15
20
25
30
0
2000
4000
6000
8000
10000
12000
14000
16000
18000
20000
22000
24000
26000
28000
30000
32000
34000
36000
38000
40000
42000
44000
46000
48000
50000
52000

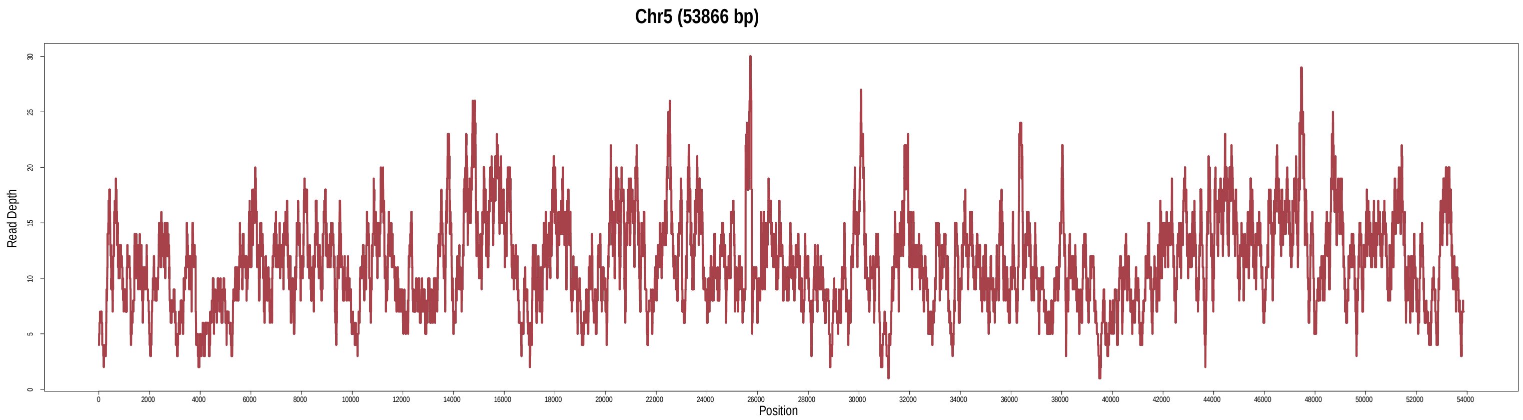

Chr5 (53866 bp)
Read Depth
Position
0
5
10
15
20
25
30
0
2000
4000
6000
8000
10000
12000
14000
16000
18000
20000
22000
24000
26000
28000
30000
32000
34000
36000
38000
40000
42000
44000
46000
48000
50000
52000
54000

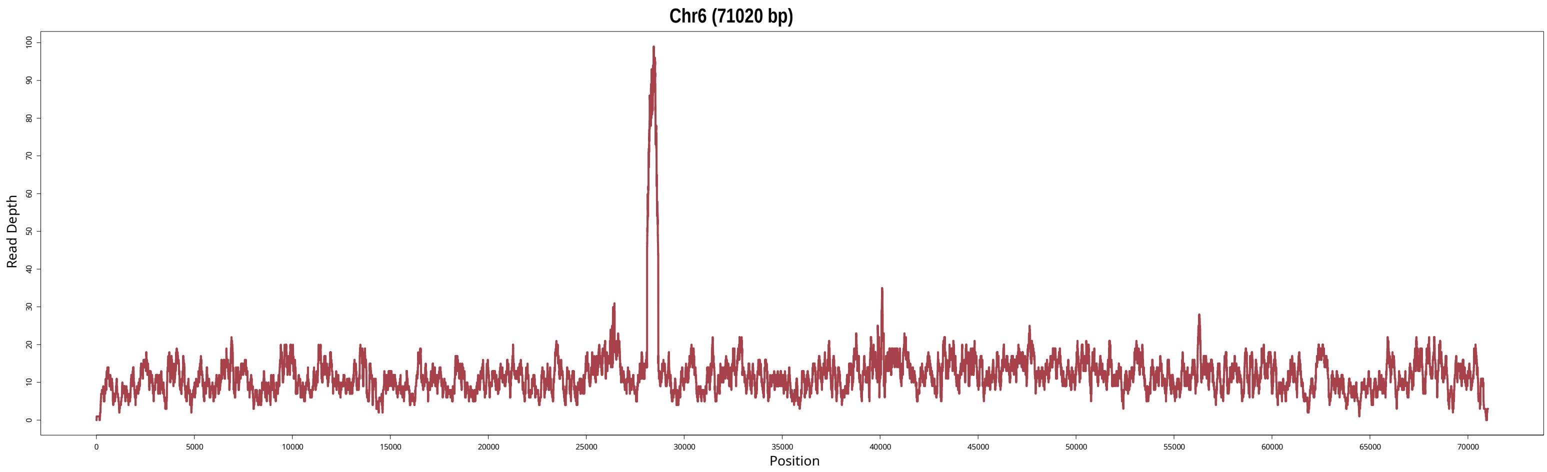

Chr6 (71020 bp)
Read Depth
Position
0
10
20
30
40
50
60
70
80
90
100
0
5000
10000
15000
20000
25000
30000
35000
40000
45000
50000
55000
60000
65000
70000

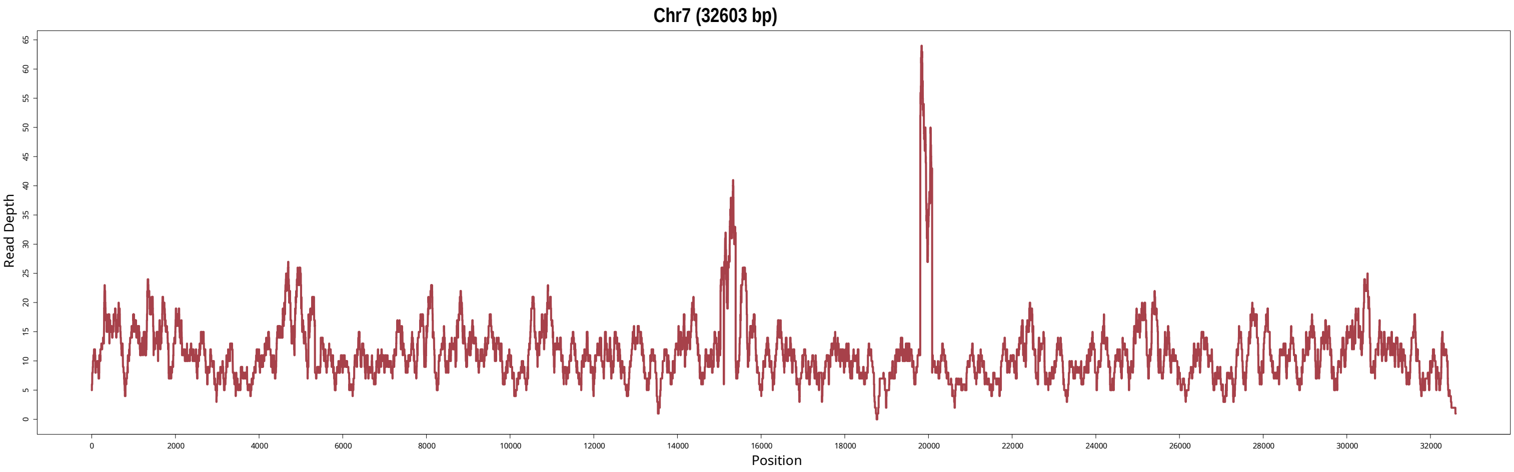

Chr7 (32603 bp)
Read Depth
0
5
10
15
20
25
30
35
40
45
50
55
60
65
Position
0
2000
4000
6000
8000
10000
12000
14000
16000
18000
20000
22000
24000
26000
28000
30000
32000

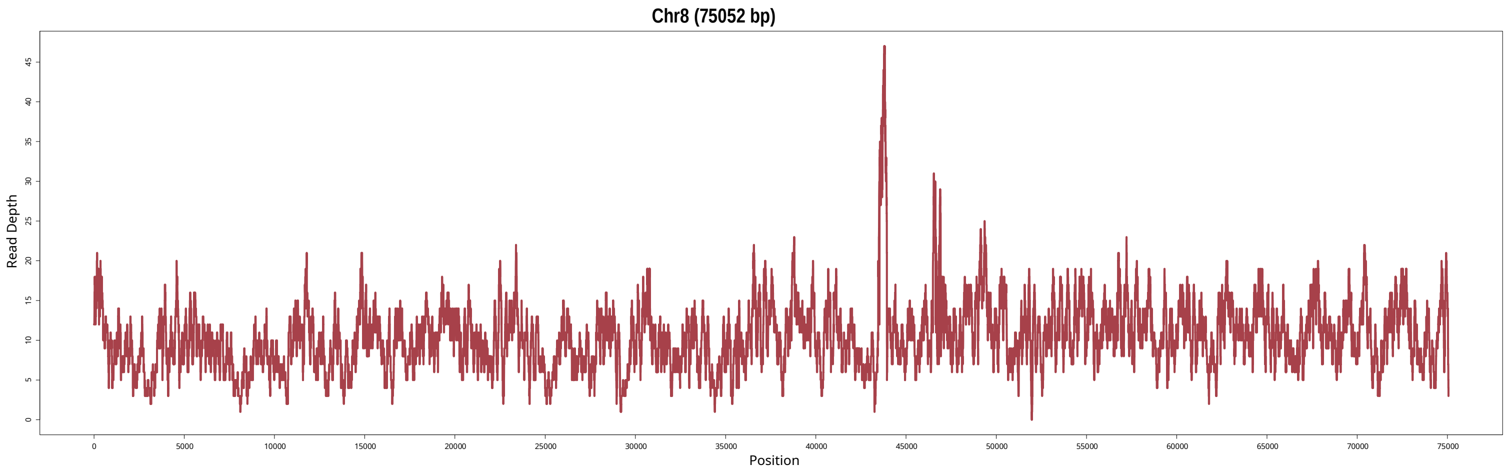

Chr8 (75052 bp)
Read Depth
Position
0
5
10
15
20
25
30
35
40
45
0
5000
10000
15000
20000
25000
30000
35000
40000
45000
50000
55000
60000
65000
70000
75000

E) *Tetrastigma sp.* Individual 2

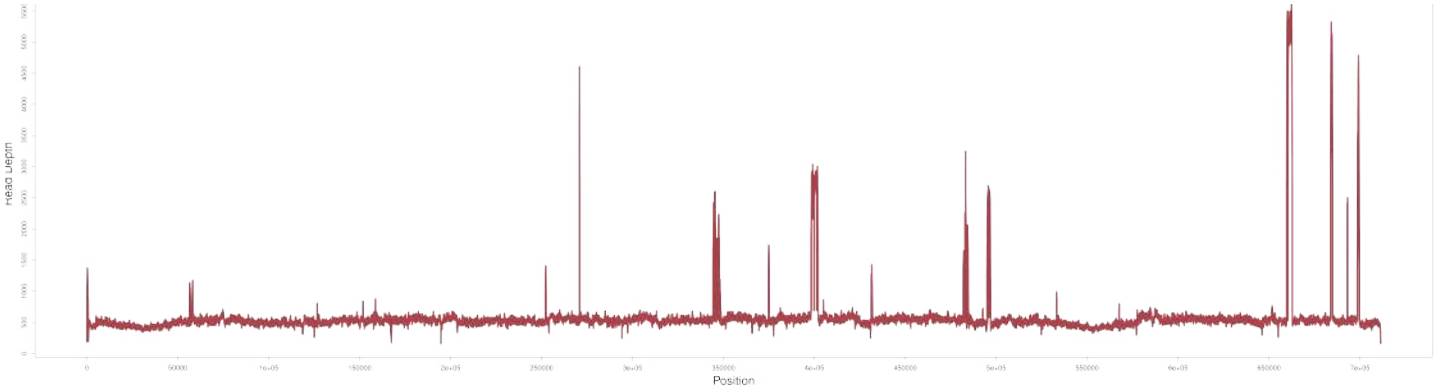

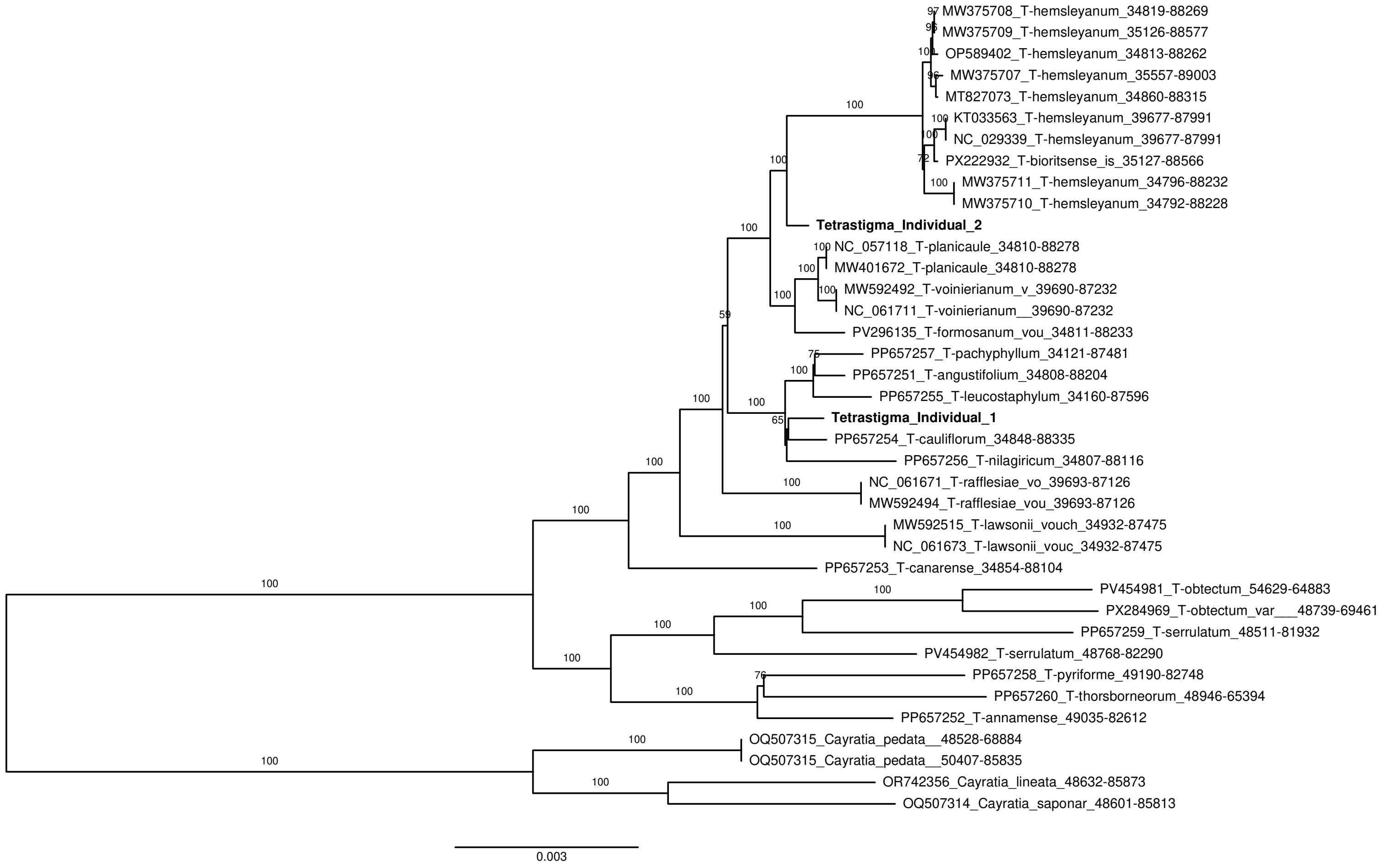


**Figure S11. Maximum-likelihood (ML) phylogenetic reconstruction of plastid regions from *Tetrastigma* (Vitaceae).** The phylogeny includes 38 *Tetrastigma* species, including the two individuals assembled in this study (*Tetrastigma* individual 1 and *Tetrastigma* individual 2). Four plastomes from *Cayratia* (Vitaceae) were used as outgroups. The alignment comprises approximately 37 kb of plastid DNA containing 43 plastid genes. Maximum-likelihood analyses were performed with RAxML. Branch support values were estimated from 1,000 rapid bootstrap replicates, and bootstrap values >50% are shown. The scale bar represents substitutions per site.
